# Tracing the Neanderthal–Modern Human hybrid zone using paleogenomic data

**DOI:** 10.64898/2026.01.06.697899

**Authors:** Lionel N. Di Santo, Claudio S. Quilodrán, Paola Cerrito, Mathias Currat

## Abstract

Interbreeding between closely related lineages is a pervasive evolutionary feature across taxa^1^, including modern humans, who admixed with Neanderthals and other archaic hominins^2,3^. Identifying where gene flow between these lineages happened can shed light on ecological and adaptive contexts in which introgressed genes were acquired. However, mapping ancient hybrid zones is challenging because post-admixture population dynamics obscure their genetic signatures. Here, we determine the conditions under which spatial patterns of introgressed DNA in ancient genomes allow the inference of the geographic location of past hybridization events during population expansions. Using computational simulations, we identify the conditions under which introgression-based statistics provide informative signals about past hybridization events. We show that the mean genomic proportion of introgressed DNA, which can be readily calculated from low-coverage ancient genomes, reveals a cline that increases with distance from the population expansion source until reaching the hybrid zone boundary and persists over time. Comparison with empirical Neanderthal introgression distribution from an extended Eurasian paleogenomic dataset supports a prolonged admixture pulse extending far beyond the Levant in western Eurasia. Our findings highlight the power of paleogenomics to reconstruct the spatiotemporal dynamics of species interactions.

## Introduction

The advent of genomics has revealed that admixture between lineages considered as distinct species is far more common in nature than previously assumed^1^, not only challenging the biological species concept^4^, but requiring biologists to explicitly define the concept of species they operationalize, depending on their research question. Although we use the word ‘species’ generically throughout this text for the sake of clarity, similar admixture processes occur across a broad range of taxonomic levels, from genera to subspecies, and even among populations. A major possible evolutionary consequence of interspecific hybridization is introgression, the incorporation of DNA from one species into the gene pool of another through repeated backcrossing^5^. Introgressed alleles can have negative consequences, including outbreeding depression^6^, genetic swamping^7^, or the introduction of maladaptive genetic variation^8^. Conversely, introgression can increase genetic diversity and introduce beneficial variants, thereby providing a substrate for new adaptations and other selective processes^9,10^. Introgression thus plays a central role in the evolution of species^11–14^, including modern humans (*Homo sapiens*), who hybridized with closely related archaic lineages, such as Neanderthals^2^, Denisovans^3^, and possibly others^15,16^. These past admixture events resulted in the introgression of archaic DNA segments into the genomes of modern humans, potentially contributing to adaptation to novel environments^17^, for example, through variants affecting the immune system^18,19^, skin pigmentation^20^, and high-altitude adaptation^21^. Insights into where hybridization occurred among these hominin lineages are therefore crucial for understanding the ecological and adaptive contexts underlying our own species’ evolutionary history.

Here, we focus on the complex relationship between modern humans and Neanderthals, two distinct lineages, hereafter considered as different species, which have diverged approximately 600 ka ago^22^. The skeletal remains from Apidima^23^, Misliya, Skhul, and Qafzeh^24^ provide evidence for early migrations of modern humans from Africa to the Levant and Europe. Without contributing significantly to present-day non-Africans, these events likely led to gene flow between Neanderthals and modern humans before 100 kya^25^ and, ultimately, to the replacement of Neanderthal mitochondrial and Y-chromosome lineages^26,27^. Nevertheless, the major contact with Neanderthals started around 60 kya during the main dispersal of modern humans out of Africa^28^. While there is ample evidence that the two lineages encountered each other and interbred during this period, the spatiotemporal dynamics of these interspecific interactions remain unclear. While between one and three admixture pulses have been estimated e.g.^20,29,30^, proposed hybridization zones between the two hominin lineages range from the Middle East^2,31^, to including Europe^32,33^, central Asia^34^, and a large area in western Eurasia^35^.

Identifying the area where interbreeding occurred (i.e. hybrid zone) is of major interest in evolutionary biology e.g.^36,37,38^ because it can provide a better understanding of the joint dynamics of gene flow, genetic drift, and natural selection where introgression occurs. Hybrid zones are typically identified by monitoring hybrid individuals^39^. However, this approach is inherently unsuitable for detecting historical hybrid zones, as field data documenting the spatial distribution and genetic composition of hybrid populations over multiple generations are rarely available^40^. Instead, abrupt shifts in allele frequencies^41,42^ and patterns of linkage disequilibrium^43–45^ can provide information on the distribution and dynamics of past hybrid zones. Yet, both approaches require large, well-sampled populations to produce reliable estimates, limiting their applicability to ancient DNA data and, therefore, their ability to investigate events deep in the past. Consequently, new statistical approaches are needed to fully exploit the rapidly growing field of paleogenomics, which now provides molecular time-series data and opens new opportunities to investigate ancient admixture events.

It has been recently shown that spatial patterns of introgression retain information about past population dynamics^46^ and behave largely as selectively neutral when examined at the genome-wide scale^47^. Studying genome-wide introgression patterns at the population level thus offers valuable perspectives on the history and dynamics of hybridization events. Statistics based on the Length of Introgressed Sequences (LIS) are widely used with modern genomic data to detect ancient admixture events^48,49^, and to estimate their timing^50–52^, duration^53^ and intensity^54^. This approach relies on the predictable dynamics of recombination at each backcrossing events, which progressively shortens the introgressed fragments over generations after admixture. As shown by a recent work^55^, a related measure, the intra-individual Variance of Introgressed Sequence length (VIS), may partially overcome the limitation imposed on the computation of LIS by the scarcity of ancient samples, as VIS can be estimated from a single genome. This makes VIS particularly useful for identifying the number and timing of admixture pulses from paleogenomic data, but its ability to detect hybrid zones has not been tested yet. Another introgression-based statistic, the Proportion of Introgression (PI), offers more limited resolution on the temporal dynamics of admixture than LIS and VIS^49,56^, except when computed from samples spanning the interval between admixture pulses^45^, a situation that is rarely encountered for ancient hybridization events.

Nevertheless, PI provides information about admixture history^49,50,57^, and it has proven especially useful for tracing spatial patterns left by the population dynamics of interbreeding groups when using paleogenomic data^46^. A notable example is the introgression signature left after a population range expansion involving hybridization, which can help infer the source and direction of the expansion^58,59^. This is important because population expansion is a recurrent evolutionary process in many plant and animal species e.g.^60^, including our own, often driven by climatic transitions e.g.^61^. In this study, we investigated how three introgression-based statistics, LIS, VIS, and PI, can help pinpoint where admixture between Neanderthals and modern humans likely occurred in Eurasia, a question that remains unresolved e.g.^35,62–64^. Our aim was to identify the conditions under which spatial patterns of genomic introgression can retrospectively reveal the geographic location of past admixture events, even in the absence of genetic data from the admixture period itself. Using population genomic simulations, we first characterized the behavior of each statistic under varying admixture pulse scenarios coupled with range expansion. We then compared these theoretical expectations with patterns of Neanderthal introgression in ancient human genomes inferred from an extended empirical genomic dataset to get insights into their hybrid zone.

## Results and discussion

### Introgression-based statistics are informative about past hybrid zones

We used computer simulations to model molecular data under various scenarios where an invasive species (e.g. modern humans) expands into the range of another local species (e.g. Neanderthals), resulting in interbreeding and eventual replacement of the pre-established local species. The range of each species was subdivided into demes, representing populations connected by *within-species* gene flow (W) to model their spatial dispersion. In areas where the ranges of both species overlap, demes from each species were also connected by *between-species* gene flow (B) to model admixture. We specifically simulated introgression from the local species into the expanding invasive species, assuming that interbreeding occurred within a confined hybrid zone covering only one-quarter of the invasive species’ total expansion range (Figure 1 and Methods). Our results show that PI in the invasive species tends to increase with distance from its expansion source, in accordance with a previous simulation study^59^ and empirical data^46^. This increasing trend is more pronounced under higher interspecific gene flow (compare B=0.01 to B=0.001 in Figure 2) and lower intraspecific gene flow (compare W=0.001 to W=0.01). An important observation is that the introgression cline occurs only within the hybrid zone, where the local species is present (Figure 2). Beyond this zone, PI levels remain stable over space and do not continue to increase. This is explained by the lack of local gene contribution outside the hybrid zone. Note, however, that our modelling framework, which includes a limited number of demes (maximum 20, see Figure 1), may underestimate the surfing effect^65^ of some introgressed alleles. Nonetheless, a previous spatially explicit simulation study using a high resolution of 10,000 demes revealed the same pattern of increasing PI within the hybrid zone^59^.

**Figure 1.**
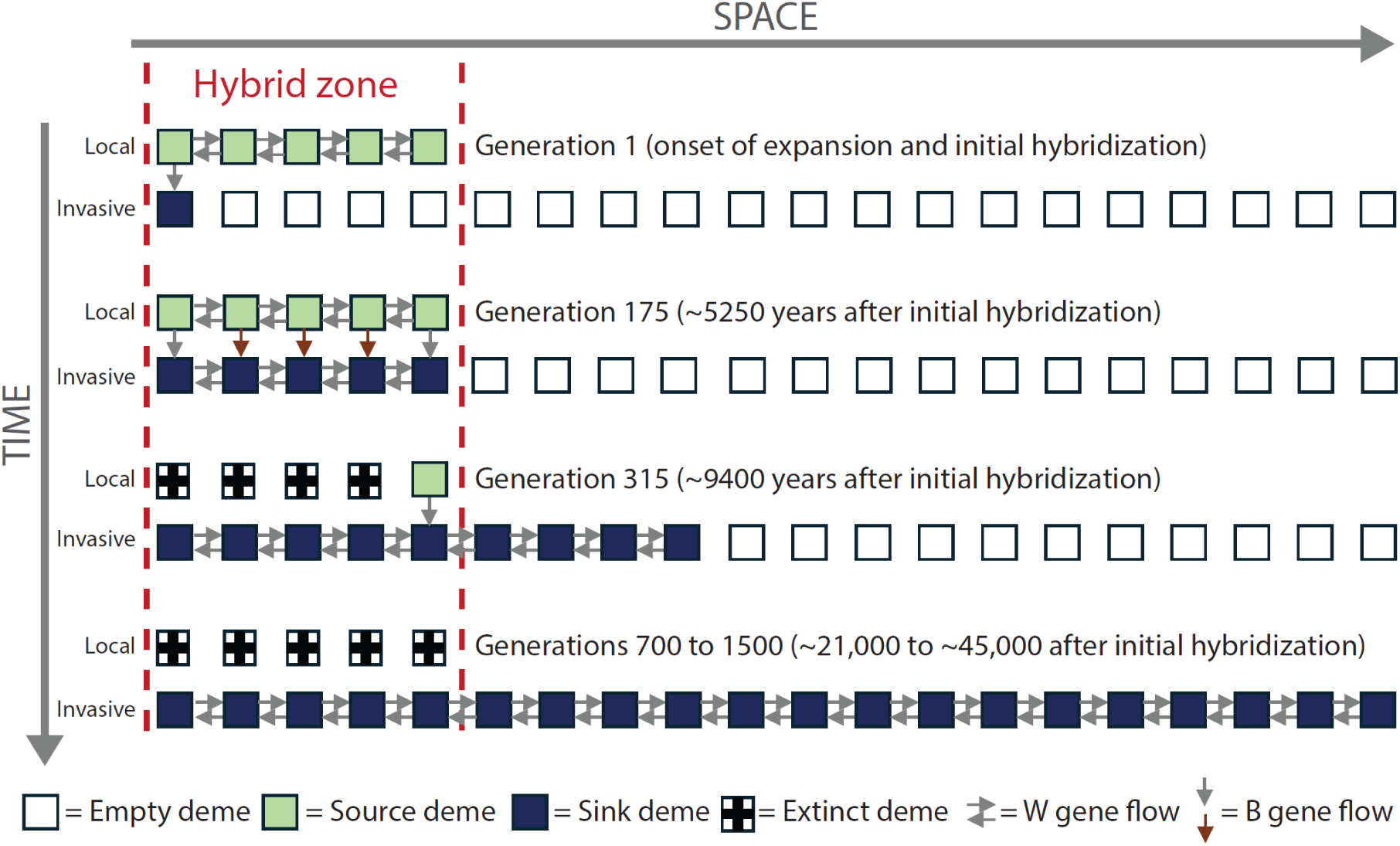
Schematic representation of the simulation framework. Each square represents a deme: the local species (e.g., Neanderthals) is shown at the top, and the invasive species (e.g., modern humans) at the bottom. Arrows indicate gene flow: W for intra-specific and B for inter-specific gene flow. Brown arrows appear only in Scenario 1 (continuous pulse).

**Figure 2.**
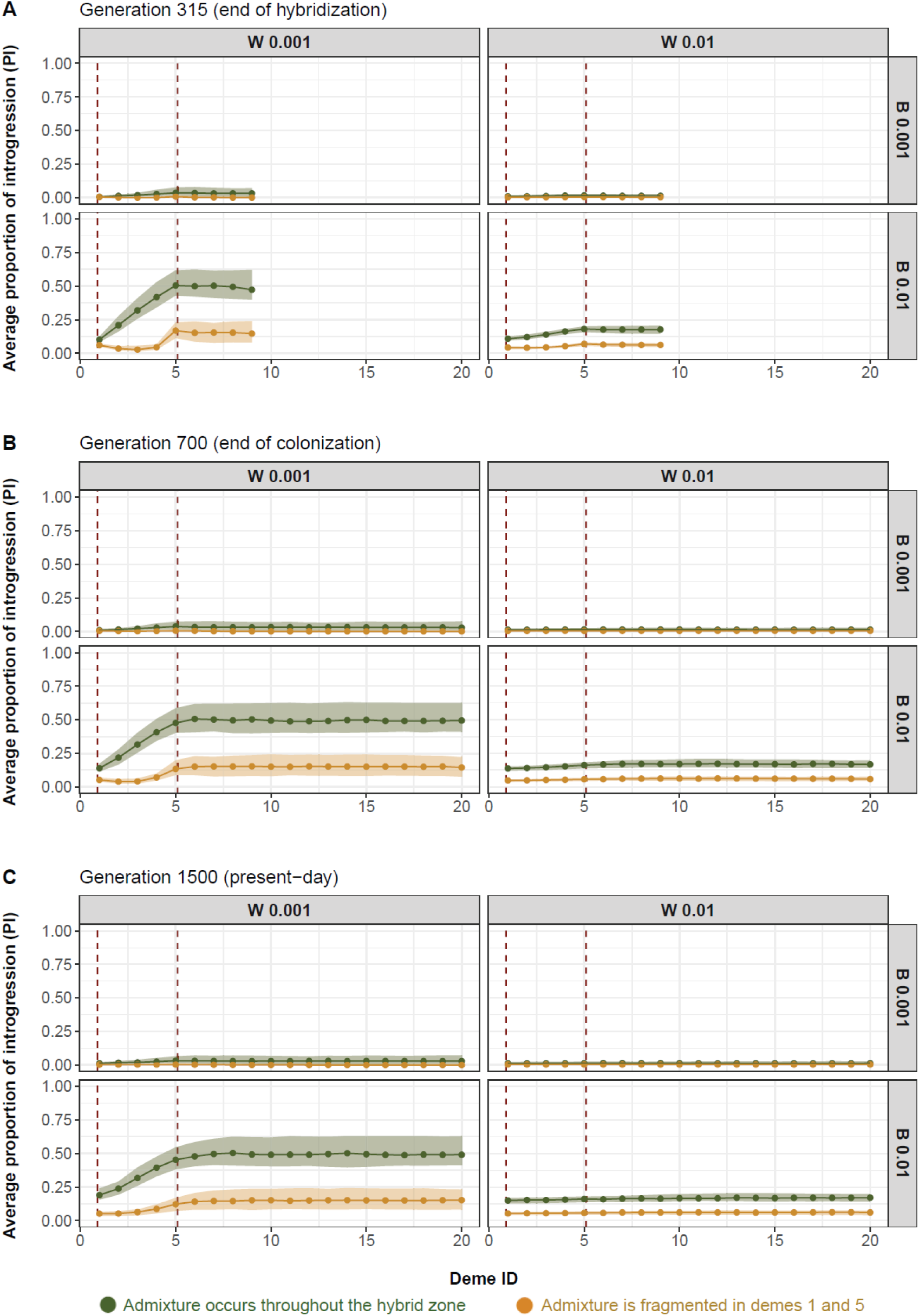
Simulated spatial introgression pattern measured by the Proportion of Introgression, PI (y-axis) after 350 (A), 700 (B), and 1500 generations (C) since the start of population expansion from deme 1 towards deme 20 (x-axis). The hybrid zone is delimited by dashed red lines. Each point represents the median of average PI across the 100 simulation replicates, and the shaded area the distribution of these averages bounded by the 25^th^ (lower) and 75^th^ (upper) quantiles. W stands for withing species gene flow, while B stands for between species gene flow.

Remarkably, the spatial pattern of PI remains highly stable over time: the clinal structure persists even 800 generations after the colonization event, and 1,200 generations after the hybridization event, although the exact boundary of the hybrid zone gradually becomes more diffuse (compare Figure 2A and Figure 2C). These patterns are observed under a “continuous” hybridization scenario (green curves in Figure 2), which involves a single, uninterrupted admixture pulse across space and time. In contrast, under a “discrete” scenario with two spatially separated admixture pulses (orange curve), PI no longer exhibits a clinal distribution. Instead, it displays one or two distinct peaks at the locations of admixture, with the second, more recent pulse producing a more pronounced peak than the first, older event. Upon closer inspection (Supplementary Figure S1), the patterns are evident immediately after the expansion in all scenarios. However, they fade more quickly over time when intraspecific gene flow (W) is greater and, though to a smaller degree, when interspecific gene flow (B) is lower. Moreover, the projected patterns are robust to sampling intensity, as sampling only one genome per deme still reproduces the same spatial structure (Supplementary Figure S2).

When examining VIS (Figure 3), the overall patterns under both scenarios resemble those observed for PI, though with some notable differences, particularly greater variance across simulation replicates. Among the three summary statistics studied, the VIS provides the clearest signal near the time of hybridization, with distinct peaks at the boundaries of the hybrid zone (Figure 3A). In the discrete admixture pulses scenario, the second pulse generates a more pronounced VIS peak than observed for the PI. However, as time progresses, these peaks fade, and the delineation of the hybrid zone blurs due to recombination breaking down long introgressed segments, levelling out their distribution within individuals. In the continuous hybridization scenario, the VIS forms an increasingly extended cline over time, spreading beyond the hybrid zone and making its boundary less distinct than with the PI (Figures 3B and 3C).

**Figure 3.**
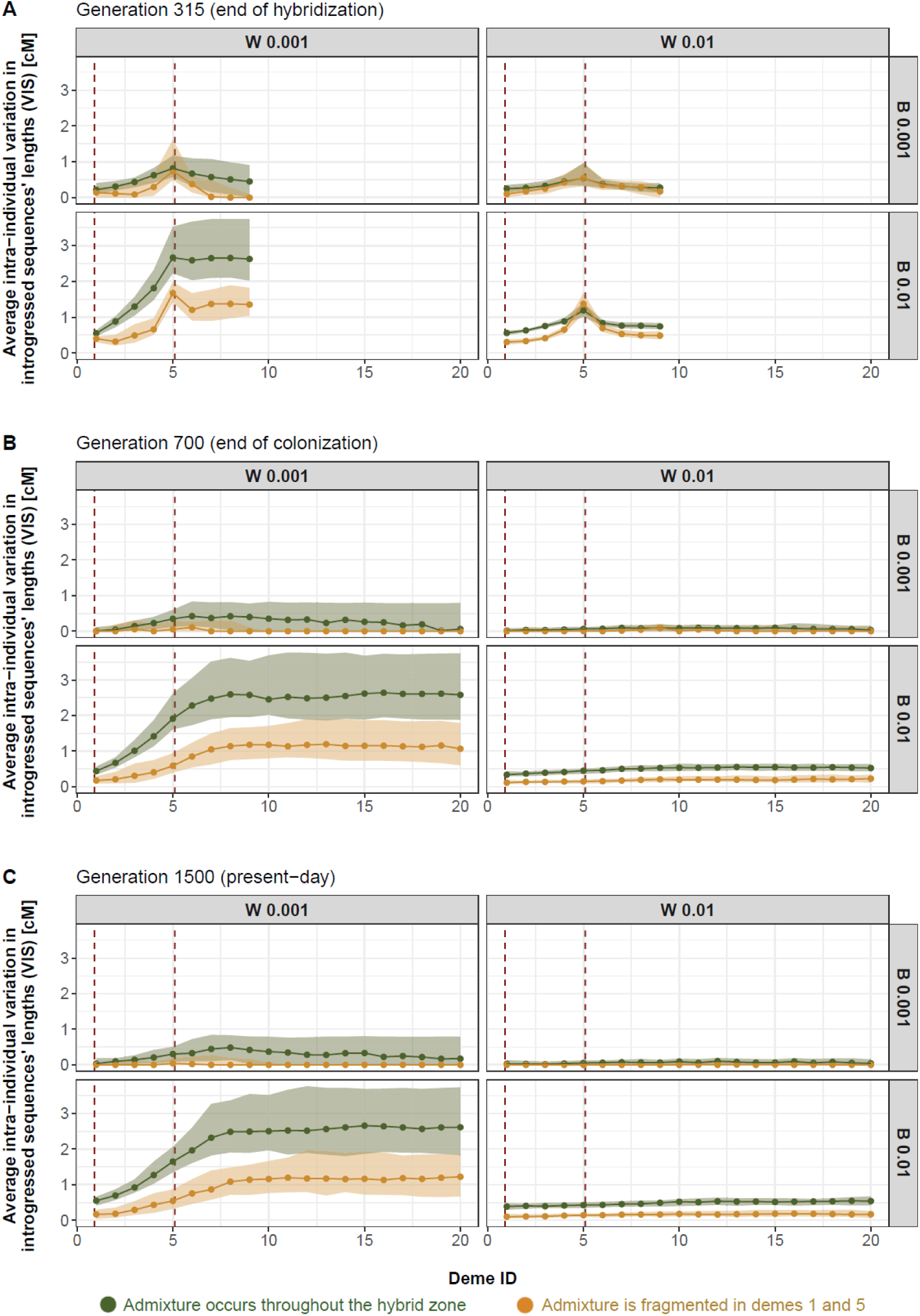
Simulated spatial introgression pattern measured by the intra-individual Variation in Introgressed Sequences’ length, VIS (y-axis) after 350 (A),700 (B), and 1500 generations (C) since the start of population expansion from deme 1 towards deme 20 (x-axis). The hybrid zone is delimited by dashed red lines. Each point represents the median of average VIS across the 100 simulation replicates, and the shaded area the distribution of these averages bounded by the 25^th^ (lower) and 75^th^ (upper) quantiles. W stands for withing species gene flow, while B stands for between species gene flow.

LIS displays patterns similar to those observed for VIS, but less pronounced, making it less effective for revealing spatial admixture signatures of past hybrid zones (Supplementary Figure S3). Moreover, a model artifact emerges when both inter- and intraspecific gene flow are low (B = W = 0.001). In this case, introgressed segments become extremely rare, and beyond deme 9, only single-base segments remain, causing LIS values to drop to zero. We therefore focus the subsequent analyses on VIS and PI.

### The proportion of introgression is an informative metric to detect past hybrid zones

The previous analysis demonstrated that the resulting spatial pattern of introgression from the local (i.e. Neanderthals) into the invasive (i.e. *H. sapiens*) species provides valuable insights into the location of interbreeding, even when only the invasive species remains. Notably, our results show that this spatial signal remains detectable over extended periods despite some blurring of its precise contours over time. Two of the introgression-based statistics analyzed (PI and VIS) exhibit similar spatiotemporal patterns and thus have strong potential to reveal the location of hybrid zones long after the admixture event. However, each metric has its nuances. VIS is most effective for detecting hybrid zones with data close in time to the admixture event and can be estimated from a single genome sequenced at sufficiently high coverage. Contrastingly, PI remains remarkably stable across generations and can thus provide a valuable means to locate hybrid zones with data post-dating the admixture event. Additionally, it has the advantage of being easily computed from low-coverage genomes, making it well-suited for detecting past hybrid zones using paleogenomic data.

### Continuous admixture generates lasting spatial introgression clines

Recent studies across diverse species indicate that admixture may occur in pulses rather than continuously^66,67^, which has opened the question about the number, timing, and spatial distribution of interbreeding between various hominins^16,25,30,55,66,68–71^. Our simulations show that introgression-based statistics can distinguish between a single continuous admixture pulse and two spatially distinct pulses at the edges of a hybrid zone, but only if the data are contemporaneous with, or shortly follow, the admixture events (Figures 2 and 3). Both statistics show that continuous pulses produce smooth clines increasing from the expansion source, whereas discrete pulses generate one or two weak peaks, with the signal being clearer for VIS than PI at the time of admixture. Over time, the signal from discrete pulses fades into a gradient that remains weaker than the one produced by a single continuous pulse, especially for VIS. Our results thus show that while multiple spatial admixture pulses can be inferred from introgression-based statistics using data contemporaneous with the admixture event, such inference becomes much more challenging at later time points.

### The conditions for the emergence of spatial introgression patterns

Furthermore, our simulations show that the introgression signatures of hybrid zones are only detectable under specific conditions, particularly when the rate of hybridization between interacting species remains low to moderate, as highlighted previously by Quilodrán, et al. ^59^. When reproductive isolation is strong, interspecific gene flow is minimal and leaves weak or undetectable genomic traces. Conversely, when hybridization is frequent, extensive introgression can obscure any spatial pattern. Our results further demonstrate a negative correlation between interspecific and intraspecific gene flow in shaping the spatial patterns of introgression, consistent with earlier findings^72^. An informative spatial introgression signal from past hybrid zones emerged only when the interspecific gene flow was sufficiently large and the intraspecific gene flow not too strong (Figures 2 and 3). In cases of increasing intraspecific gene flow, introgressed alleles become rapidly homogenized across populations, effectively erasing any spatial signal of hybridization and reducing the overall level of introgression^58,73^. Nevertheless, although the conditions required to detect a spatial pattern of introgression may be limited, numerous empirical cases exhibit introgression rates consistent with those simulated under our modelling framework. Notably, this is the case of the clinal pattern of Neanderthal ancestry in modern humans^46^, which empirically demonstrates that hybridization signals occurring during range expansion can persist and remain detectable even long after ancient admixture events. Such examples are not limited to hominins but can also be found in plants (e.g. wild tomatoes^74^, European white oaks^75^), and in animals (European wildcats^76^, canids^77^).

### Inferred Neanderthal-Modern human Hybrid zone covers a wide area in Western Eurasia

Based on the properties of introgression-based statistics described with our simulations, especially the cline stopping abruptly at the end of the hybrid zone, we used a large dataset of 4,147 paleogenomic samples older than 600 years ago^78^, distributed in more than 1,200 data points (Supplementary Table S1, see Methods for details) to infer the extent of the hybrid zone between Neanderthals and modern humans. Among the summary statistics evaluated, we selected PI because simulations showed that it provides more accurate results for data post-dating the admixture period than VIS and LIS. Indeed, although the oldest sample in the dataset dates to ∼44.4 kya, which is within the putative interbreeding period between modern humans and Neanderthals (∼40-54 kya,^31,79^), all other samples postdate this period. Moreover, PI can be computed from low-coverage data, thereby expanding the size of the datasets that can be used. We modelled PI using a Generalized Additive Model (GAM), using the Levant as the putative source of modern human expansion into Eurasia^80^. We defined the hybrid zone as the geographic region where Neanderthal introgression in human genomes increases spatially and then ceases, with the termination point marking its boundary.

The results show a gradual increase in PI with distance from the Levantine source, plateauing both westward toward Europe and eastward into Asia (Figure 4A), consistent with the expected pattern of a single continuous admixture pulse (Figure 2). Interestingly, the distance over which PI increases is similar in Europe and in Asia (∼3,900 *km*), in accordance with a previous study based on modern populations supporting a large hybrid zone in Asia^35^. The GAM regression shows a maximum around 3,900 *km* and a second, less pronounced peak near 9,000 *km* (Figure 4C), which likely reflects an artifact due to the limited number of observations at such great distances from the source of expansion (red dotted line in Figure 4A and yellow triangle in Figure 4B). We inferred that hybridization took place in a zone extending approximately between 0° and 69°E longitude and 29°N to 59°N latitude (Figure 4B). The inferred hybrid zone includes almost all known sites associated with Neanderthal fossils, spanning western Eurasia^81–83^, except those from the Altai region^82^. The hybrid zone inferred from paleogenomic data is more extensive than previous proposals based on modern genomic data^2^, or estimated with spatiotemporal species distribution models^62^, paleoclimatic simulations^63^, or ecological niche modeling^64^. Note that the Atlantic fringe, including western France and most of the Iberian Peninsula, is not included in the inferred hybrid zone, despite the well-documented Neanderthal presence e.g.^84,85^. This could be explained either by the absence of hybridization in this region or by the lack of precision of our approach, given the time lapse between most of the samples and the admixture event.

**Figure 4.**
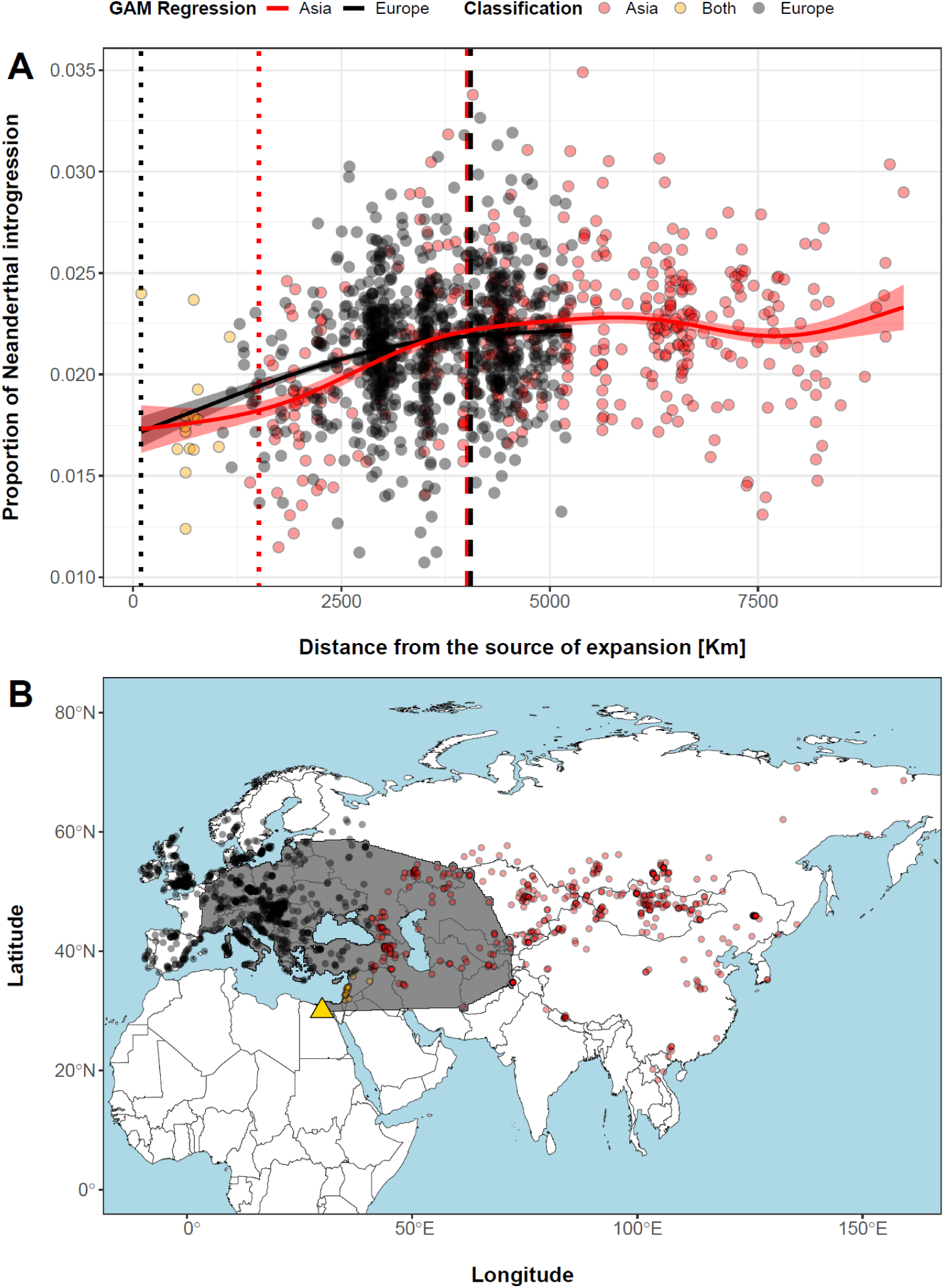
Detection of the historical hybrid zone between Neanderthals and modern humans. The analysis is based on 1,264 groups of paleogenomes older than 600 years retrieved from the AADR database^78^. A) Proportion of Neanderthal introgression (y axis) in European (black dots), Asian (red dots), and both groups (yellow dots) samples over their distance from the source of expansion (x axis). Vertical lines represent the inferred start (dotted lines) and end (dashed lines) of past European (black) and Asian (red) contact zones between the two hominin species (see Materials and Methods for details). Solid black and red lines represent GAM regressions of Neanderthal introgression over topographic distance along the European and Asian colonization routes, respectively, with shaded areas showing standard errors of regression lines. B) Geographic locations of European (black dots), Asian (red dots) and both (yellow dots) groups of paleogenomes over a map of Eurasia. The inferred hybrid zone is shown in grey and includes European and Asian samples located within regions where PI increases with distance from the source of modern human expansion into Eurasia (yellow triangle; arbitrarily positioned in Cairo).

To refine the spatial resolution of the hybrid zone, we thus analysed a subset of 32 genomes dating closer to the time of admixture (older than 10,000 years; Figure 5), which is expected to delineate the hybrid zone boundaries more precisely (Figure 2). The results are consistent with previous analyses by showing an increasing trend of introgression from the putative source of expansion of modern humans, compatible with a single admixture pulse. The inferred latitudinal range remains stable, while the longitudinal range shifts westward, notably including the western Atlantic fringe of Europe (reaching 2W° longitude) and spanning up to 70°E. Note that with this ancient subset, there was no data available neither from the Iberian Peninsula, nor from the Near East, which are therefore not included in the hybrid zone, despite the Near East being a widely recognized region of admixture^80^. The discrepancy between the western hybrid zone boundary inferred from the full dataset and that obtained from the subset of the most ancient genomes, which extends to the westernmost available data, encompassing all of France and southern Britain, may reflect the limited precision of boundary estimates resulting from the temporal distance to the admixture event. With all datasets (full and most ancient), the hybrid zone ends near 55°N –60°N to the north, around the Baltic Sea, consistent with ecological modelling of Neanderthal occurrence^86^, and the absence of archaeological evidence of Neanderthal presence beyond this latitude^87^

**Figure 5.**
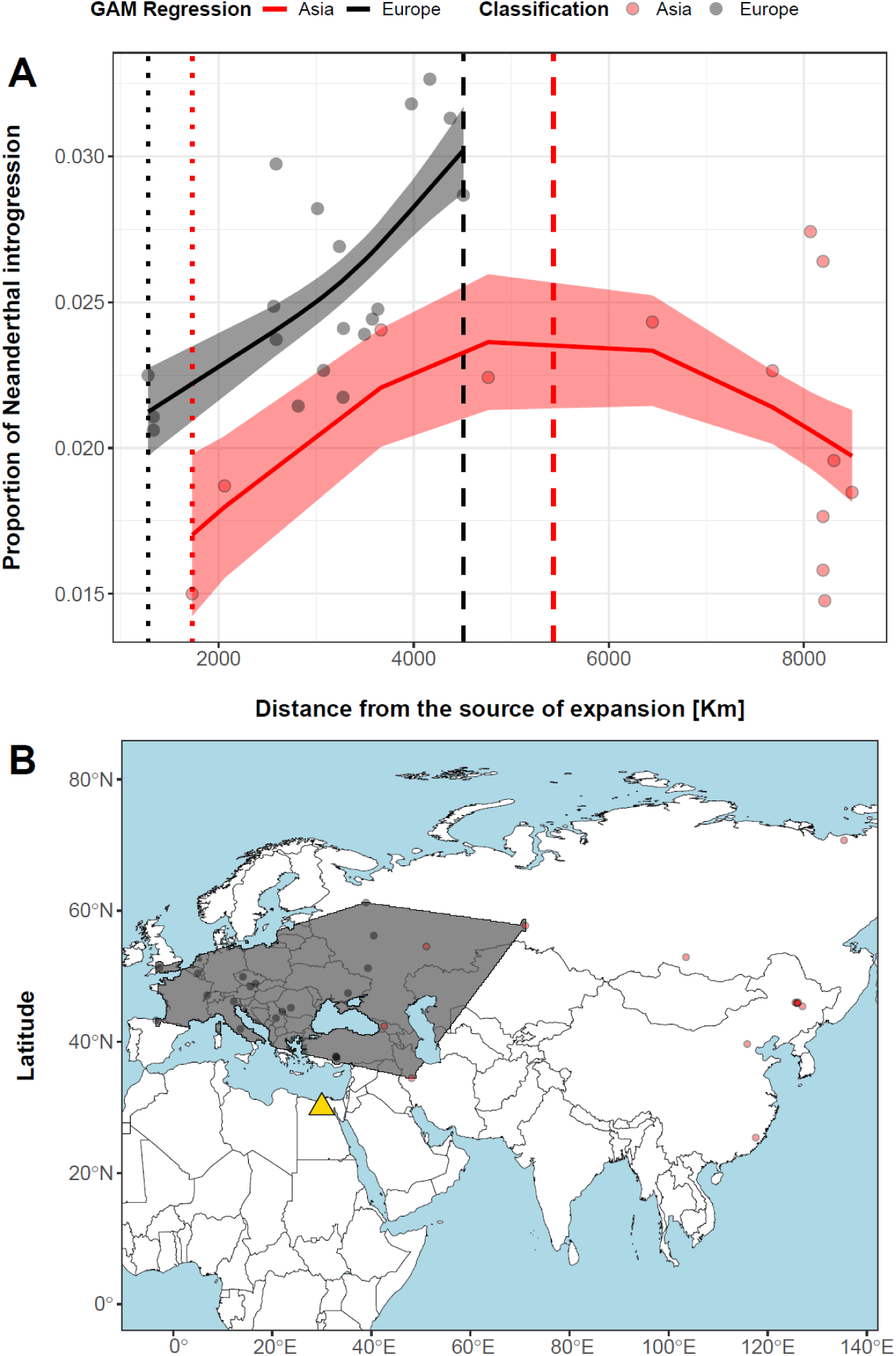
Detection of the historical hybrid zone between Neanderthals and modern humans. The analysis is based on 32 groups of paleogenomes older than 10,000 years retrieved from the AADR database ^78^. A) Proportion of Neanderthal introgression (y axis) in European (black dots), Asian (red dots), and both groups (yellow dots) samples over their distance from the source of expansion (x axis) Vertical lines represent the inferred start (dotted lines) and end (dashed lines) of past European (black) and Asian (red) contact zones between the two hominin species (see Materials and Methods for details). Solid black and red lines represent GAM regressions of Neanderthal introgression over topographic distance along the European and Asian colonization routes, respectively, with shaded areas showing standard errors of regression lines. B) Geographic locations of European (black dots) and Asian (red dots) groups of paleogenomes over a map of Eurasia. The inferred hybrid zone is shown in grey and includes European and Asian samples located within regions where PI increases with distance from the source of modern human expansion into Eurasia (yellow triangle; arbitrarily positioned in Cairo).

### Limits and perspectives of detecting past hybrid zones with introgression-based statistics

While our approach using introgression-based statistics offers a valuable framework for approximating the geographic zones where hybridization between closely related lineages occurred, it comes with several limitations. First, accurate inference depends on the availability of a sufficiently large and geographically well-distributed paleogenomic dataset, as well as specific conditions, most notably, a low to moderate rate of introgression. Second, the approach is applicable in scenarios involving population expansions with a putative source that can be reasonably inferred. Third, the accuracy of inferred boundaries diminishes as the temporal distance between the sampled genomes and the admixture event increases.

Although the approach can estimate the broad location of an admixture zone, it does not account for potential shifts in its position over time unless a spatiotemporally dense dataset from the period of admixture is available. Finally, the approach assumes that average levels of introgression are shaped primarily by neutral processes and demographic dynamics. This is a reasonable assumption in the case of Neanderthal introgression into humans when assessed at the genome level^46,47^, but the potential effects of selection on the spatiotemporal distribution of introgression remain to be investigated.

Future modelling studies could also explore whether the apparent absence of modern human DNA in late Neanderthals is solely the result of asymmetric introgression resulting from population dynamics during modern human range expansion out of Africa^35^ or difference in population sizes^88^, or whether other factors must be invoked, such as the limited genetic record of late Neanderthals or a social selective pressure against hybrids within Neanderthal groups^89^. It would therefore be particularly interesting to assess whether genetic models that incorporate sex-specific processes and social mechanisms can better reproduce the empirical spatial distribution of introgression across modern human and Neanderthal genomes, at autosomal but also sex-specific markers. Nonetheless, population dynamics provide the neutral baseline explanation for introgression patterns, to which other mechanisms can be added.

### Conclusion

The recent accumulation of paleogenomic data allow us to gain critical insights into the dynamics of historical species interactions and improve our understanding of present-day patterns of human genomic diversity. Inferring the spatial extent of past hybrid zones is particularly important, as these zones offer opportunities to study how populations adapt to new environments^36,90,91^, diverge, and potentially give rise to new species^11,92^. Pinpointing where and when admixture occurred in our own species may help identify introgressed DNA segments with functional effects, such as those influencing neurological, psychiatric, immunological, dermatological, and eating disorders e.g.^93,94^, conferring resistance or susceptibility to pathogens e.g.^95,96^ or facilitating adaptation to specific environmental conditions e.g.^18^. Our results support the hypothesis of a single hybridization pulse extending through both space and time, suggesting that as modern humans expanded their geographic range into Eurasia, they continued to interbreed with the Neanderthals they encountered along the way. This interpretation is consistent with two previous independent modelling studies, one emphasizing temporal dynamics^53^ and the other focusing on spatial patterns^35^.

Our identification of a broad, continuous spatial pulse of hybridization in Eurasia is relevant in light of recent evidence for differences in communicative mechanisms^97^ and vocal tract anatomy^98^ between the two lineages, implying that hybridization did not require shared communicative behaviors to succeed. It also contributes to explain the geographical disparity between known Neanderthal remains^81–83^ and Neanderthal ancestry among present-day populations^99–101^.

## Methods

### Genomic simulations

#### Model Overview

To evaluate genomic patterns of introgression inside and outside a hybrid zone following admixture between two divergent species, we used the R package ‘glads’^102^, an individual-based, forward-in-time simulation framework. We considered the evolution of two divergent interbreeding species composed of diploid individuals distributed across populations (demes). Every individual was defined by their genomic composition, sex, and population assignment (function *evolve()*, type = “dynamics”). The genomic background of each individual was modelled as a single pair of homologous chromosomes defined by their length (Mbps), the number and location of genetic markers (e.g., SNPs) along their sequence, biallelic genotypes at each genetic marker position, the mutation rate per base pair, and the chromosome-wide average recombination rate per base pair.

The simulation started with parametrized chromosomes and sexual identity of individuals within their respective population and species. The simulation began at generation *t* and proceeded by initiating reproduction among individuals within populations at *t*+1. The number of offsprings each mating pair can produce within a given generation (*O_t_*) was assigned from a Poisson distribution, with a mean (𝜆) that varies with generation-specific population densities (*N_t_*) and a density-dependent effect (𝜎*_𝑑em_*) to avoid exponential growth: 𝑂_𝑡_ = *Poisson* (𝜆) − 𝑁_𝑡_𝜎*_dem_*. The model proceeded in discrete time steps representing generations (*t*), with migrations from population *i* to *j* (𝑚*_ij_*) allowing gene flow within (W) each species. Populations from different species were organized as different grids of demes, potentially overlapping where the species co-occur. The probability of a newborn to migrate before it reaches sexual maturity was obtained from a uniform distribution. A value of zero means no migration, while a value of 0.5 means random migration and hence random reproduction between populations. Gene flow between species (B) was similarly regulated but also incorporated a density-dependence adjustment. The probability of admixture between individuals from different species increased or decreased proportionally to the respective population size of spatially co-occurring species. Similarly to W, a B value of zero means no admixture between species, while a value of 0.5 means random reproduction between species. More information about the framework can be obtained at www.glads.app.

#### Initialization

In our simulations, we considered two species, each one organized in a grid of demes distributed in a stepping-stone manner. Each deme represents a population within a species (Fig. 1). Because we analyzed unidirectional gene flow, we considered the local species as source and the invasive species as sink, the latter receiving introgressed DNA segments in its gene pool. The home range of the local species was characterized by five demes, while 20 demes were available for the invasive species. The five demes overlapping between species represented the hybridization zone between them. At the initial generation, the five source demes were already occupied at demographic equilibrium (∼100 individuals), while the invasive species started to colonize its grid of demes from the left corner deme with 100 initial individuals. For both species, the sex ratio within demes was assumed to be equal (probability of being a male = 0.5). The mean number of offspring (λ = 4) and the density-dependent demographic effect (𝜎_*dem*_ = 0.02) in the local species were selected for their propensity to stabilize population sizes around the initial value of 100 individuals, establishing a demographic equilibrium. Conversely, demographic values for the invasive species (λ = 3, 𝜎_*dem*_ = 0.001) were selected for their predisposition to gradually increase population sizes over time until reaching demographic equilibrium (∼1,200 individuals), effectively mimicking the natural process of colonization. In total, we simulated 1,500 generations, grossly corresponding to the timing of admixture between modern humans and Neanderthals^53^, so about 45,000 years considering a human generation time of 30 years^103^. Accordingly, we also based our genetic parameters on estimates from modern humans^104^, modelling the genomic background of individuals as a single pair of homologous chromosomes, each 150 Mbps in length, containing 201 evenly spaced SNPs and a recombination rate of 1.2 × 10⁻⁸ per base pair ^104^. Because the objective was tracking the fate of introgression and not reproducing a faithful evolutionary history, we set the mutation rate to zero and modelled fixed biallelic SNPs for both species, with allele 1 representing the invasive species and allele 2 representing the local species. We added a species-specific genetic marker at the end of each chromosome to track first-generation immigrants from one species into the grid of demes of the other species. These first-generation immigrants were removed when computing introgression-based summary statistics, as they would have introduced a bias. Note that the progeny of these individuals in subsequent generations was assigned a genetic marker corresponding to the grid of demes they were born into, so considered as introgressed individuals.

#### Simulation experiments

The simulation framework roughly represents the colonization of an area by an invasive species (e.g. *Homo sapiens*), with possible hybridization with a local species (e.g. Neanderthals) in only one quarter of the zone, which corresponds approximately to the area of Eurasia occupied by Neanderthals according to Currat and Excoffier ^35^. At the start of the simulation, all five local demes (in blue, Figure 1) were at demographic equilibrium, while the invasive species began its expansion from a single deme on the far left, initially occupied by 100 individuals (in green, Figure 1). Intra-specific gene flow (W) occurred at each generation between neighbouring demes of the same species arranged along a linear (vectorial) dimension. Inter-specific gene flow (B) occurred unidirectionally between demes of different species spatially co-locating, always from the local to the invasive demes. The first five demes on the left defined the hybrid zone, while the remaining 15 demes were exclusively occupied by the expanding invasive species. It took approximately 175 generations (e.g. ∼5,250 years, assuming a 30-year generation time) for the invasive species to fully colonize the hybrid zone, advancing at a rate of one deme every 35 generations. Afterwards, with each additional deme colonized beyond the hybrid zone, at the same pace, the leftmost local deme went extinct, reflecting the gradual retreat and disappearance of the local species following a period of cohabitation of 175 generations (e.g. ∼5,250 years) per deme. By generation 315 (e.g. ∼9,400 years after the onset of the invasive species’ expansion), the local species thus completely disappeared. The invasive species completed its expansion across the entire vector of 20 demes by generation 700 (e.g. ∼21,000 years), after which it remained at demographic equilibrium, maintaining intra-specific gene flow between neighbouring demes for an additional 800 generations (e.g. ∼24,000 years).

We simulated two scenarios of admixture: The first (continuous) scenario simulated a single pulse of hybridization that was continuous both spatially and temporally across all five demes within the hybrid zone. Although interspecific reproduction may occur at different times and in different demes, hybridization was uninterrupted, with no spatiotemporal fragmentation. The second (discrete) scenario simulated two separate pulses of hybridization occurring at opposite ends of the hybrid zone. In this case, interspecific gene flow is restricted to demes 1 and 5, with no hybridization occurring in demes 2 to 4. This setup captured a pattern of hybridization that was fragmented both spatially and temporally: deme 1 was colonized by the invasive species at the onset of the simulation, while deme 5 was not reached until 140 generations later. Finally, to explore the impact variation in within- and among- species gene flow may have on patterns of genomic summary statistics, we tested four combinations of interspecific (B=0.001, B = 0.01) and intraspecific (W=0.001, W = 0.01) migration values for each statistic. These specific values were selected based on exploratory simulations to enable meaningful comparisons of PI values, while avoiding extremes of either complete or absent introgression between the two species. We measured each statistic at three timepoints: (1) at the end of the hybridization phase (generation 315), (2) at the end of colonization (generation 700), and (3) at the end of simulation (generation 1,500). Each combination of admixture and migration scenario was replicated 100 times to account for the stochastic nature of simulations.

#### Estimation of population-level genomic summary statistics

Using simulation outputs from selected generations (315, 700, and 1,500), we estimated three genomic summary statistics for each of the 100 replicates of non-empty sink demes to describe genomic patterns of introgression. These statistics included: 1/ the average Length of Introgressed Segments (LIS); 2/ The average intra-individual Variation in Introgressed Segment lengths (VIS); 3/ The average Proportion of Introgression (PI). To estimate LIS, each homologous chromosome from individuals assigned to an invasive deme was scanned for continuous tracts of local alleles (allele 2). When at least one such allele was present, we identified consecutive introgressed alleles along a homolog as forming an introgressed segment. The positions of the first and last markers in each segment were recorded, and the segment’s length was calculated in base pairs. These values were then converted into centimorgans (cM) by multiplying them by 1.2 × 10⁻⁶. Deme-specific LIS was ultimately calculated as the average segment length (in cM) across all inferred introgressed segments and individuals. The second statistic, VIS, captured the within-individual variation in segment lengths. It was estimated first by computing the standard deviation of all introgressed segment lengths in each individual, and then by averaging these standard deviations across all individuals within an invasive deme. Notably, segments composed of only a single genetic marker were excluded from LIS and VIS calculations, as segment length in cM could not be determined for these cases. PI was computed for each individual as the total number of introgressed alleles across homologs divided by the total number of haploid markers (402). Deme-level PI was then estimated by averaging individual PI values across all individuals within an invasive deme. To avoid biases, genotypes of all first-generation immigrants from the local species were excluded from an invasive deme using the species-specific diagnostic allele. This allele was also removed from the genetic background of all individuals prior to estimating introgression-related statistics.

All simulation and analysis functions, including those for filtering immigrant individuals, removing the diagnostic allele, and computing summary statistics, were compiled into an R package named “companions4glads2”^55^. Finally, spatiotemporal dynamics of summary statistics across invasive demes were assessed by evaluating deme-specific distributions across all 100 simulation replicates per recorded generation. All analyses were conducted in R versions 4.3.1 (R Core Team 2023).

### Analysis of ancient genomes

#### Data collection and filtering

We evaluated whether simulated patterns of introgression could be detected in empirical data by using ancient genomes obtained from the Allen Ancient DNA Resource (AADR) version 54.1.p1 released on 03/06/2023^78^. Of all available genomes, we retained only those from *Homo sapiens* passing the database quality assessment score and with a number of SNPs ≥ 500,000. From the retained genomes, we then excluded redundant entries, keeping for each individual the version with the highest coverage and, when missing, the version with the highest number of SNP hits. In case multiple genomes for one individual were still present when selected based on coverage, the version with the most SNPs was kept. If these combined filters did not reduce the number of available genomes for an individual to one, then a genome among those remaining was chosen at random. Finally, because the focus was solely on genomic patterns of Neanderthal introgression into modern humans across Europe and Asia, ancient genomes distributed outside these regions were discarded. We considered a genome to belong to Eurasia when located on any landmass within a longitudinal range from −180.0 to 180.0 decimal degrees and a latitudinal range from 5.6 to 81.9 decimal degrees. These filters were applied (i) to retain genomes that maximize inference accuracy while minimizing data loss to maintain adequate spatiotemporal sampling, and (ii) to ensure the independence of sampled genomes. In total, we retained 4,147 genomes dated from 600 to ∼44,00 years ago (Supplementary Table S1). A lower threshold of 600 years was chosen so that all genomes considered predate the large-scale migrations associated with the onset of the colonial era. This database also includes 44 genomes older than 10,000 years. To separate the analysis according to the two main directions of *Homo sapiens* colonization from Africa, each sample was attributed either to the European group (*n* = 2,986), the Asian group (*n* = 1,113), or both (*n* = 48) based on their spatial location. Genomes located within a longitudinal range > 41 and < −120 decimal degrees were assigned to the Asian group, genomes located within a longitudinal range > -120 and < 41 decimal degrees were assigned to the European group, and genomes within a longitudinal range ≥ 30 and ≤ 41 decimal degrees and a latitudinal range ≥ 29 and ≤ 36 were assigned to both groups. Lastly, we selected high-coverage genomes from two Neanderthals (AltaiNeanderthal.DG and Vindija_snapAD.DG), one African *Homo sapiens* genome presumed to lack archaic introgression from Neanderthals (S_Dinka-1.DG), and one chimpanzee genome (Chimp.REF) to serve as references in F4-ratio calculations.

#### Estimation of genomic summary statistic

We estimated Neanderthal ancestry (denoted α) in each target genome using the F4-ratio method as described by Reich et al.^105^. This approach involves computing the ratio of two F4-statistics derived from a five-population framework, in which one population is assumed to be admixed between two others. To accounts for gene flow from European populations into northern and western Africa, we followed the correction proposed by Petr, et al. ^106^. Specifically, the Dinka population from eastern Africa was used as the reference (“C”) for the test genome (“X”) instead of Yoruba population. The Altai Neanderthal (“A”) was considered as the sister population to the Vindija Neanderthal (“B”), while the chimpanzee genome (“O”) acted as the outgroup. Genomic comparisons indicate that the Neanderthal ancestry in present-day non-Africans derives from a population more closely related to the Vindija (and Mezmaiskaya) Neanderthals than to the Altai Neanderthal^22,93^. The proportion of Vindija Neanderthal ancestry in genome X was then estimated using the formula: α = F4(A, O; X, C) / F4(A, O; B, C), where A =AltaiNeanderthal.DG, B = Vindija_snapAD.DG, C = S_Dinka-1.DG, O = Chimp.REF. Hereafter, we refer to this F4-ratio estimate as the measure of Neanderthal introgression into the test genome.

Because all genomes were not uniquely geographically located, we first grouped genomes with the same geographic coordinates together and considered the average across all genomes within a group as the measure of Neanderthal introgression. Although not identically dated, genomes within the vast majority of multi-genomes groups (51% of all groups) were relatively close in age (77% had a within-group standard deviation < 250 years and 95% had a within-group standard deviation < 1,000 years). Consequently, we believe grouping genomes with identical geographic coordinates is unlikely to have introduce any notable bias in estimates of introgression proportions. In total, from the initial 4,147 genomes, 1,264 groups, including 1,232 with genomes older than 600 years and 32 with genomes older than 10,000 years were used in subsequent analyses (Supplementary Table S1). To estimate how far from the putative out-of-Africa source a genome or group of genomes is located in Eurasia, we computed the shortest on-land topographic distance (in *km*) between each genome or group of genomes and the origin of *Homo sapiens* expansion using the R package “topoDistance”^107^. The origin was arbitrarily set in Cairo (yellow triangle in Figure 4B), as in Quilodrán, et al. ^46^. Topographic distance between three genomes belonging to two groups (gun011_noUDG.SG, I7332, and I7346) and the origin could not be estimated and were excluded from further analyses.

We used the Generalized Additive Model (GAM) framework implemented within the R package “mgcv”^108^ to investigate the non-linear relationships between an explanatory variable, here the topographic distance between each genome’s or group of genomes’ location and the source of the modern human expansion in Eurasia, and a response variable, here the proportion of Neanderthal ancestry within genomes or groups of genomes, measured as F4-ratios, as in Quilodrán, et al. ^46^. The optimal value for the basis dimension (*k* parameter), which affect the shape of the GAM curve, was determined for each continental group (Europe and Asia) by selecting the value minimizing the Akaike Information Criterion (AIC). We tested values of *k* ranging from 3 to 150. When no models showed a clear advantage, i.e. AIC values were identical for all *k* tested up to the fifth significant digit, we selected the smallest *k* value (*k* = 3). To establish the limits of the hybrid zone between Neanderthal and *Homo sapiens*, we used the R package “gratia”^109^ to compute first-order derivatives of GAM regression models and their 95% confidence intervals across 1000 equally spaced topographic distance values over the observed range of topographic distances. According to inferences drawn from simulations, we independently delineated the lower limit (the start) of the European and Asian hybrid zones at the smallest topographic distance from the origin with a significantly positive derivative (α = 0.05), indicating Neanderthal introgression started and was increasing. The upper limits (the end) of hybrid zones were set to the first topographic distance from the origin, greater than the lower limit, with a derivative non-significantly different from zero (α = 0.05), indicating Neanderthal introgression into modern human stopped. Based on these four values (two for each hybrid zone), we identified genomes or groups of genomes within the range bounded by inferred topographic distance limits and generated a spatial raster covering all these genomes using the R package “terra”^110^. This raster provided the estimate of the geographic location and distribution of the potential hybrid zone between modern human and Neanderthal based on introgression proportions. These analyses were conducted two times, once considering proportions of Vindija introgression in genomes or groups of genomes (i) older than 600 years (Figure 4) and (ii) older than 10,000 years (Figure 5). Note that we used the actual derivative estimate instead of the lower CI limit to identify hybrid zone boundaries when analyzing GAM models built with genomes older than 10,000 years due to their large confidence intervals resulting from the small number of available genomes (*n* = 32). Because of the uncertainty surrounding these derivatives, we recommend caution when interpreting hybrid zones inferred from genomes older than 10,000 years old. Note also that the hybrid zone shown in Figure 4B includes all Asian genomes up to a topographic distance of ∼4,000 km from the source (red dashed line), rather than only those within ∼1,500 km (red dotted line). This choice is justified because the GAM regression for Asian genomes matches the pattern expected under a discrete admixture model (see Figure 2), and excluding genomes closer to the source could bias the inferred spatial limits of the hybrid zone.

## Acknowledgments

This work was supported by the Swiss National Science Foundation grants 31003A_182577 and IZCO-0-240503 (to M.C.). We thank Pascale Gerbault, Kostas Kampourakis, Estella Poloni and Carel Van Schaik for their careful reading of an early version of the manuscript.

## Author contributions

Conceptualization and methodology: all authors; Project supervision and funding acquisition: M.C.; Data curation: L.N.D; Software: C.S.Q and L.N.D; Data analysis and simulations:

L.N.D. and P.C.; Original draft: M.C and L.N.D. Writing review and editing of the final version: all authors;

## Data and materials availability

All data needed to evaluate the conclusions of the study are present in the article and the supplementary materials. All genomes analysed in this study were already published, and references are provided in data S1. The version (0.1.5) of GLADS used for the study is available at www.glads.app. The complete set of functions used to estimate introgression summary statistics is available in the R package “companions4glads2” (version 2.1.5.9) published in Di Santo, *et al.* ^55^.

## Supplementary Table and Figures

**Table S1.** General information and proportion of Neanderthal introgression estimated on selected AADR’s ancient genomes.

**Figure S1.**
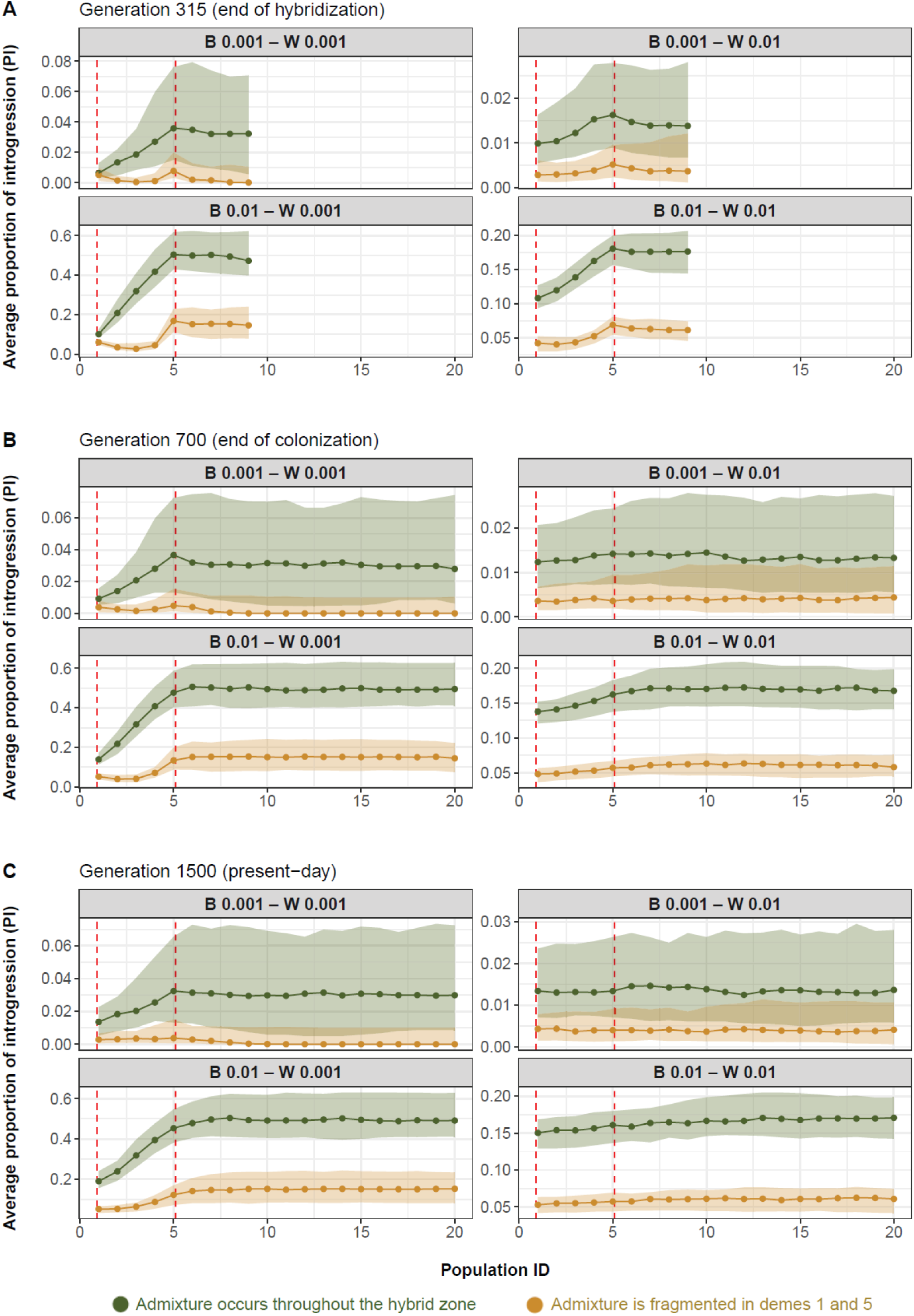
Simulated spatial introgression pattern measured by the Proportion of Introgression, PI (y-axis) after 350 (A), 700 (B), and 1500 generations (C) since the start of population expansion from deme 1 towards deme 20 (x-axis). The hybrid zone is delimited by dashed red lines. Each point represents the median of average PI across the 100 simulation replicates, and the shaded area the distribution of these averages bounded by the 25^th^ (lower) and 75^th^ (upper) quantiles. W stands for withing species gene flow, while B stands for between species gene flow. Note that, unlike Figure 2 in the main text, **this figure uses a free y-axis**; the scales therefore differ across panels, which better highlights the patterns.

**Figure S2.**
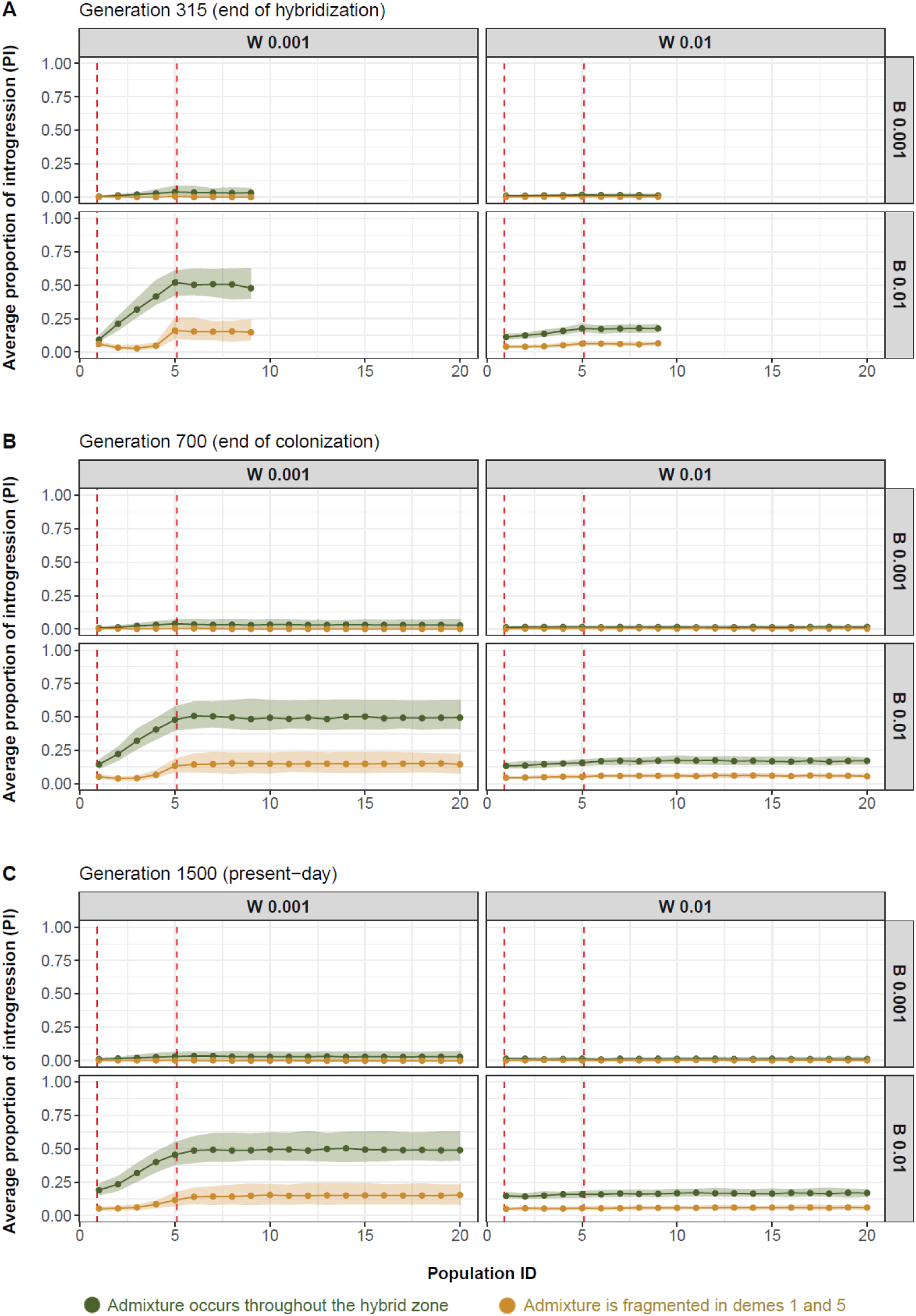
Simulated spatial introgression pattern measured by the Proportion of Introgression, PI (y-axis) after 350 (A), 700 (B), and 1500 generations (C) since the start of population expansion from deme 1 towards deme 20 (x-axis). The hybrid zone is delimited by dashed red lines. Each point represents the median of average PI across the 100 simulation replicates, and the shaded area the distribution of these averages bounded by the 25^th^ (lower) and 75^th^ (upper) quantiles. W stands for withing species gene flow, while B stands for between species gene flow. Note that, in contrast to Figure 2 in the main text, the pattern shown here is based on **sampling a single genome per deme** rather than using all genomes within each deme.

**Figure S3.**
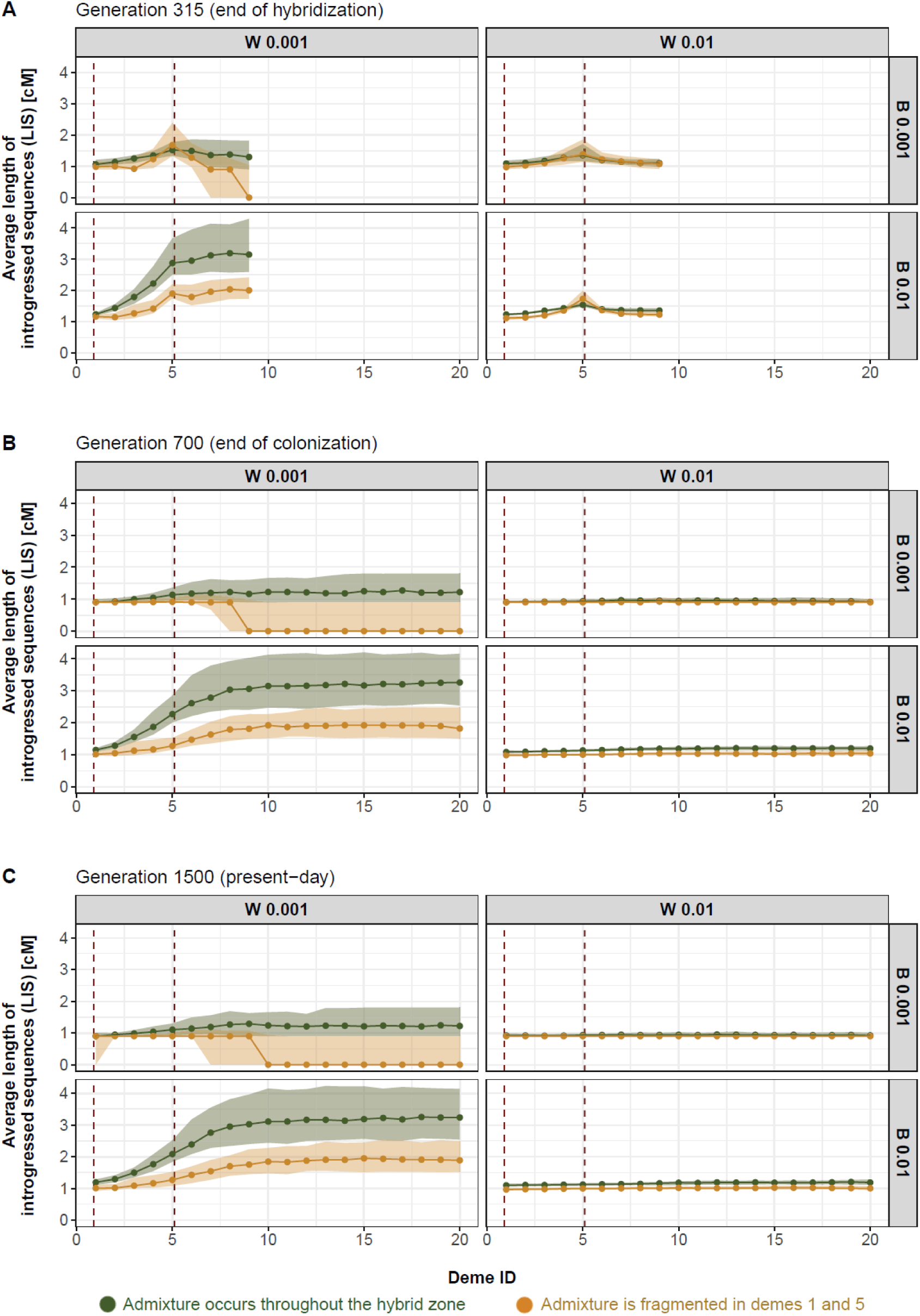
Simulated spatial introgression pattern measured by the Length of Introgressed Sequences, LIS (y-axis) after 350 (A), 700 (B), and 1500 generations (C) since the start of population expansion from deme 1 towards deme 20 (x-axis). The hybrid zone is delimited by dashed red lines. Each point represents the median of average LIS across the 100 simulation replicates, and the shaded area the distribution of these averages bounded by the 25^th^ (lower) and 75^th^ (upper) quantiles. W stands for withing species gene flow, while B stands for between species gene flow.

