## Supplementary Table S1 for "Tracing the Neanderthal–Modern Human hybrid zone using paleogenomic data"

**Table S1.** General information and proportion of neanderthal introgression ($PI_{Vindija}$) estimated using the Vindija reference on selected AADR's ancient genomes.

[1] Each ancient genome analyzed was paired with a group ID according to its geographic coordinates (Longitude, Latitude). When identical, ancient genomes were assigned the same group ID. For the inference of putative hybrid zones, introgression proportions were averaged per group and groups instead of individual genomes

| Genome ID (AADR) | Group ID[1] | Continent | Country | Distance to the origin of expansion [Km] | Longitude | Latitude | Average age of the genome (years before 1950) | $PI_{Vindija}$ |
|---|---|---|---|---|---|---|---|---|
| UstIshim_snpAD.DG | 407 | Asia | Russia | 4763.8 | 71.10 | 57.70 | 44366 | 0.0224 |
| Tianyuan | 399 | Asia | China | 7677.7 | 115.87 | 39.66 | 39565 | 0.0227 |
| Kostenki14 | 978 | Europe | Russia | 3076.2 | 39.30 | 51.23 | 38052 | 0.0227 |
| GoyetQ116-1_noUDG | 544 | Europe | Belgium | 3976.3 | 5.01 | 50.45 | 35208 | 0.0318 |
| Sunghir2_noUDG.SG | 1147 | Europe | Russia | 3576.6 | 40.50 | 56.18 | 34629 | 0.0220 |
| Sunghir3_noUDG.SG | 1147 | Europe | Russia | 3576.6 | 40.50 | 56.18 | 34517 | 0.0252 |
| PM1 | 1045 | Europe | Romania | 2589.6 | 23.75 | 45.19 | 34415 | 0.0297 |
| Sunghir4_noUDG.SG | 1147 | Europe | Russia | 3576.6 | 40.50 | 56.18 | 34323 | 0.0274 |
| ZKU002_noUDG.SG | 1264 | Europe | Czech Republic | 3495.2 | 14.07 | 49.92 | 34054 | 0.0239 |
| NE20_noUDG | 332 | Asia | China | 8307.2 | 127.03 | 45.40 | 33592 | 0.0196 |
| Sunghir1_noUDG.SG | 1147 | Europe | Russia | 3576.6 | 40.50 | 56.18 | 33209 | 0.0230 |
| Yana_old_noUDG.SG | 417 | Asia | Russia | 8066.8 | 135.42 | 70.72 | 31850 | 0.0282 |
| Yana_old2_noUDG.SG | 417 | Asia | Russia | 8066.8 | 135.42 | 70.72 | 31850 | 0.0266 |
| I2483_noUDG | 797 | Europe | Austria | 3279.7 | 15.59 | 48.41 | 30850 | 0.0241 |
| Vestonice16 | 1178 | Europe | Czech Republic | 3275.6 | 16.65 | 48.88 | 29800 | 0.0217 |
| MA1_noUDG.SG | 322 | Asia | Russia | 6446.8 | 103.50 | 52.90 | 24320 | 0.0243 |
| NE56_noUDG | 340 | Asia | China | 8195.5 | 126.15 | 46.00 | 19427 | 0.0264 |
| ElMiron_d | 530 | Europe | Spain | 4374.2 | -3.45 | 43.26 | 18775 | 0.0313 |
| ZBC_IPB001.B-C0101_Luk2-Pinarbasi | 1262 | Europe | Turkey | 1277.7 | 33.03 | 37.48 | 15411 | 0.0225 |
| NE34_noUDG | 336 | Asia | China | 8195.5 | 125.58 | 45.98 | 14521 | 0.0176 |
| Villabruna_noUDG | 1180 | Europe | Italy | 3238.7 | 12.21 | 46.15 | 14021 | 0.0269 |
| Bichon_noUDG.SG | 483 | Europe | Switzerland | 3632.1 | 6.87 | 47.10 | 13698 | 0.0248 |
| SATP_noUDG.SG | 375 | Asia | Georgia | 2061.3 | 42.59 | 42.38 | 13282 | 0.0187 |

| | | | | | | | | |
|---|---|---|---|---|---|---|---|---|
| PES001_noUDG.SG | 1041 | Europe | Russia | 4165.3 | 38.91 | 61.23 | 12711 | 0.0326 |
| R11.SG | 1062 | Europe | Italy | 3014.0 | 13.54 | 41.96 | 11868 | 0.0310 |
| Qihe3 | 351 | Asia | China | 8493.2 | 117.60 | 25.40 | 11564 | 0.0185 |
| Sidelkino_noUDG.SG | 384 | Asia | Russia | 3665.1 | 51.11 | 54.53 | 11321 | 0.0241 |
| I5241 | 885 | Europe | Serbia | 2590.5 | 22.01 | 44.60 | 11198 | 0.0281 |
| I5235 | 885 | Europe | Serbia | 2590.5 | 22.01 | 44.60 | 10835 | 0.0220 |
| I5240 | 885 | Europe | Serbia | 2590.5 | 22.01 | 44.60 | 10805 | 0.0217 |
| I5244 | 885 | Europe | Serbia | 2590.5 | 22.01 | 44.60 | 10785 | 0.0265 |
| R7.SG | 1062 | Europe | Italy | 3014.0 | 13.54 | 41.96 | 10706 | 0.0254 |
| I1819 | 715 | Europe | Ukraine | 2816.3 | 35.28 | 47.43 | 10658 | 0.0231 |
| NE3_noUDG | 334 | Asia | China | 8213.7 | 126.00 | 45.85 | 10628 | 0.0148 |
| NE4_noUDG | 339 | Asia | China | 8195.5 | 125.85 | 45.93 | 10557 | 0.0158 |
| I5242 | 885 | Europe | Serbia | 2590.5 | 22.01 | 44.60 | 10530 | 0.0241 |
| I5239 | 885 | Europe | Serbia | 2590.5 | 22.01 | 44.60 | 10333 | 0.0223 |
| I6767_noUDG.SG | 926 | Europe | United Kingdom | 4509.5 | -2.77 | 51.28 | 10280 | 0.0287 |
| VLASA32.SG | 1252 | Europe | Serbia | 2565.0 | 20.74 | 43.61 | 10223 | 0.0249 |
| ZHJ_BON024.A0101_Luk84 | 1263 | Europe | Turkey | 1331.7 | 32.87 | 37.75 | 10190 | 0.0206 |
| I1954 | 168 | Asia | Iran | 1729.2 | 48.12 | 34.45 | 10162 | 0.0150 |
| I1763 | 715 | Europe | Ukraine | 2816.3 | 35.28 | 47.43 | 10081 | 0.0198 |
| Bon002_noUDG.WGC | 488 | Europe | Turkey | 1331.7 | 32.86 | 37.75 | 10078 | 0.0211 |
| I5236 | 885 | Europe | Serbia | 2590.5 | 22.01 | 44.60 | 10038 | 0.0214 |
| I5238 | 885 | Europe | Serbia | 2590.5 | 22.01 | 44.60 | 9993 | 0.0269 |
| I1947 | 168 | Asia | Iran | 1729.2 | 48.12 | 34.45 | 9992 | 0.0157 |
| AH1_noUDG.SG | 2 | Asia | Iran | 1813.8 | 48.37 | 34.19 | 9925 | 0.0147 |
| AH2_noUDG.SG | 2 | Asia | Iran | 1813.8 | 48.37 | 34.19 | 9921 | 0.0219 |
| AH4_noUDG.SG | 2 | Asia | Iran | 1813.8 | 48.37 | 34.19 | 9918 | 0.0174 |
| ZHAG_BON004.A0101_Luk10 | 1263 | Europe | Turkey | 1331.7 | 32.87 | 37.75 | 9900 | 0.0210 |
| ZKO_BON001.A0101_Luk7 | 1263 | Europe | Turkey | 1331.7 | 32.87 | 37.75 | 9900 | 0.0194 |
| ZMOJ_BON014.A0101_Luk21 | 1263 | Europe | Turkey | 1331.7 | 32.87 | 37.75 | 9900 | 0.0209 |
| Ne30_genotyping_noUDG | 335 | Asia | China | 8270.0 | 126.23 | 45.96 | 9896 | 0.0165 |
| I1290 | 168 | Asia | Iran | 1729.2 | 48.12 | 34.45 | 9806 | 0.0118 |

| | | | | | | | | |
|---|---|---|---|---|---|---|---|---|
| Kolyma_River_noUDG.SG | 303 | Asia | Russia | 9030.1 | 159.10 | 68.60 | 9775 | 0.0255 |
| I4871 | 869 | Europe | Serbia | 2544.2 | 22.05 | 44.53 | 9750 | 0.0243 |
| Bianbian_noUDG | 23 | Asia | China | 8078.1 | 118.50 | 36.10 | 9709 | 0.0195 |
| KK1_noUDG.SG | 296 | Asia | Georgia | 2064.0 | 43.28 | 42.28 | 9678 | 0.0210 |
| I5234 | 885 | Europe | Serbia | 2590.5 | 22.01 | 44.60 | 9500 | 0.0243 |
| I5237 | 885 | Europe | Serbia | 2590.5 | 22.01 | 44.60 | 9500 | 0.0179 |
| I5409 | 885 | Europe | Serbia | 2590.5 | 22.01 | 44.60 | 9500 | 0.0209 |
| Hum2_noUDG.SG | 560 | Europe | Norway | 4499.6 | 7.74 | 58.06 | 9364 | 0.0241 |
| I4630 | 865 | Europe | Latvia | 3691.3 | 25.13 | 56.28 | 9218 | 0.0217 |
| WC1_noUDG.SG | 411 | Asia | Iran | 1636.2 | 47.11 | 34.61 | 9214 | 0.0207 |
| VLASA7.SG | 1252 | Europe | Serbia | 2565.0 | 20.74 | 43.61 | 9170 | 0.0219 |
| BS | 37 | Asia | China | 7974.4 | 117.90 | 36.50 | 9125 | 0.0213 |
| Chan_noUDG.SG | 504 | Europe | Spain | 4731.0 | -7.03 | 42.73 | 9109 | 0.0296 |
| R15.SG | 1062 | Europe | Italy | 3014.0 | 13.54 | 41.96 | 9086 | 0.0249 |
| I5436 | 893 | Europe | Romania | 2590.5 | 22.61 | 44.63 | 9025 | 0.0302 |
| Ne35_genotyping_noUDG | 337 | Asia | China | 8195.5 | 125.95 | 45.91 | 8990 | 0.0211 |
| Klein7.SG | 876 | Europe | Austria | 3275.6 | 16.59 | 48.66 | 8970 | 0.0229 |
| I4881 | 869 | Europe | Serbia | 2544.2 | 22.05 | 44.53 | 8869 | 0.0225 |
| M96_noUDG.SG | 893 | Europe | Romania | 2590.5 | 22.61 | 44.63 | 8825 | 0.0198 |
| irk007_noUDG.SG | 272 | Asia | Russia | 6589.0 | 105.85 | 53.99 | 8801 | 0.0220 |
| I4916 | 872 | Europe | Serbia | 2590.5 | 22.30 | 44.64 | 8763 | 0.0262 |
| I4870 | 869 | Europe | Serbia | 2544.2 | 22.05 | 44.53 | 8740 | 0.0239 |
| Dushan4_1 | 107 | Asia | China | 7636.3 | 107.13 | 23.60 | 8707 | 0.0205 |
| OC_noUDG.SG | 868 | Europe | Romania | 2544.2 | 22.72 | 44.52 | 8704 | 0.0214 |
| I0867 | 425 | Both | Israel | 531.4 | 35.17 | 31.79 | 8700 | 0.0163 |
| I4582 | 868 | Europe | Romania | 2544.2 | 22.72 | 44.52 | 8618 | 0.0218 |
| BDB001 | 480 | Europe | Germany | 3710.4 | 12.07 | 51.30 | 8600 | 0.0240 |
| I5411 | 893 | Europe | Romania | 2590.5 | 22.61 | 44.63 | 8600 | 0.0223 |
| AKT16.SG | 458 | Europe | Turkey | 1828.6 | 28.77 | 40.17 | 8563 | 0.0235 |
| I4877 | 869 | Europe | Serbia | 2544.2 | 22.05 | 44.53 | 8505 | 0.0249 |
| I4874 | 869 | Europe | Serbia | 2544.2 | 22.05 | 44.53 | 8499 | 0.0220 |

| | | | | | | | | |
|---|---|---|---|---|---|---|---|---|
| I4550 | 865 | Europe | Latvia | 3691.3 | 25.13 | 56.28 | 8489 | 0.0222 |
| I4875 | 869 | Europe | Serbia | 2544.2 | 22.05 | 44.53 | 8458 | 0.0164 |
| I4876 | 869 | Europe | Serbia | 2544.2 | 22.05 | 44.53 | 8455 | 0.0224 |
| I0061 | 567 | Europe | Russia | 4291.4 | 35.65 | 61.65 | 8450 | 0.0243 |
| irk00x_noUDG.SG | 273 | Asia | Russia | 7241.5 | 115.87 | 55.67 | 8419 | 0.0244 |
| Yumin_noUDG | 418 | Asia | China | 7494.0 | 114.20 | 42.00 | 8375 | 0.0216 |
| Bar25.SG | 478 | Europe | Turkey | 1775.5 | 29.61 | 40.30 | 8316 | 0.0234 |
| I4632 | 865 | Europe | Latvia | 3691.3 | 25.13 | 56.28 | 8316 | 0.0289 |
| Rev5_noUDG.SG | 1120 | Europe | Greece | 2255.2 | 20.92 | 39.49 | 8316 | 0.0178 |
| I0736 | 580 | Europe | Turkey | 1775.5 | 29.57 | 40.30 | 8300 | 0.0227 |
| I1096 | 580 | Europe | Turkey | 1775.5 | 29.57 | 40.30 | 8300 | 0.0156 |
| I1099 | 580 | Europe | Turkey | 1775.5 | 29.57 | 40.30 | 8300 | 0.0221 |
| I1101 | 580 | Europe | Turkey | 1775.5 | 29.57 | 40.30 | 8300 | 0.0165 |
| I1103 | 580 | Europe | Turkey | 1775.5 | 29.57 | 40.30 | 8300 | 0.0176 |
| I1097 | 580 | Europe | Turkey | 1775.5 | 29.57 | 40.30 | 8282 | 0.0133 |
| I1098 | 580 | Europe | Turkey | 1775.5 | 29.57 | 40.30 | 8282 | 0.0193 |
| Spiginas4 | 1141 | Europe | Lithuania | 3744.2 | 22.42 | 55.77 | 8276 | 0.0263 |
| I1583.DG | 478 | Europe | Turkey | 1775.5 | 29.61 | 40.30 | 8273 | 0.0184 |
| Bar31_noUDG.SG | 478 | Europe | Turkey | 1775.5 | 29.61 | 40.30 | 8272 | 0.0140 |
| I0744 | 580 | Europe | Turkey | 1775.5 | 29.57 | 40.30 | 8270 | 0.0162 |
| cta016_noUDG.SG | 68 | Asia | Russia | 7535.3 | 118.94 | 50.02 | 8264 | 0.0279 |
| I1581_enhanced | 478 | Europe | Turkey | 1775.5 | 29.61 | 40.30 | 8252 | 0.0161 |
| I0745 | 580 | Europe | Turkey | 1775.5 | 29.57 | 40.30 | 8243 | 0.0178 |
| Nea3.SG | 1016 | Europe | Greece | 2230.1 | 22.25 | 40.58 | 8218 | 0.0219 |
| I1580_enhanced | 478 | Europe | Turkey | 1775.5 | 29.61 | 40.30 | 8205 | 0.0191 |
| NE39_noUDG | 338 | Asia | China | 8195.5 | 125.90 | 45.91 | 8196 | 0.0234 |
| I5402 | 872 | Europe | Serbia | 2590.5 | 22.30 | 44.64 | 8156 | 0.0295 |
| I1960 | 211 | Asia | Russia | 4537.3 | 69.34 | 56.04 | 8116 | 0.0289 |
| I4915 | 872 | Europe | Serbia | 2590.5 | 22.30 | 44.64 | 8115 | 0.0245 |
| I1736 | 715 | Europe | Ukraine | 2816.3 | 35.28 | 47.43 | 8110 | 0.0250 |
| I0854_enhanced | 478 | Europe | Turkey | 1775.5 | 29.61 | 40.30 | 8104 | 0.0206 |

| | | | | | | | | |
|---|---|---|---|---|---|---|---|---|
| I0708 | 580 | Europe | Turkey | 1775.5 | 29.57 | 40.30 | 8101 | 0.0203 |
| Nea2.SG | 1016 | Europe | Greece | 2230.1 | 22.25 | 40.58 | 8101 | 0.0212 |
| I1734 | 715 | Europe | Ukraine | 2816.3 | 35.28 | 47.43 | 8100 | 0.0240 |
| I0707 | 580 | Europe | Turkey | 1775.5 | 29.57 | 40.30 | 8096 | 0.0193 |
| LEPE52.SG | 988 | Europe | Serbia | 2590.5 | 22.03 | 44.56 | 8093 | 0.0216 |
| I4914 | 872 | Europe | Serbia | 2590.5 | 22.30 | 44.64 | 8088 | 0.0217 |
| I0709 | 580 | Europe | Turkey | 1775.5 | 29.57 | 40.30 | 8085 | 0.0211 |
| I1579_enhanced | 580 | Europe | Turkey | 1775.5 | 29.57 | 40.30 | 8072 | 0.0139 |
| Bar8_noUDG.SG | 478 | Europe | Turkey | 1775.5 | 29.61 | 40.30 | 8064 | 0.0228 |
| I4917 | 872 | Europe | Serbia | 2590.5 | 22.30 | 44.64 | 8058 | 0.0245 |
| Loschbour.DG | 991 | Europe | Luxembourg | 3854.3 | 6.40 | 49.81 | 8025 | 0.0242 |
| I1585_enhanced | 580 | Europe | Turkey | 1775.5 | 29.57 | 40.30 | 8021 | 0.0176 |
| I5233 | 885 | Europe | Serbia | 2590.5 | 22.01 | 44.60 | 8001 | 0.0178 |
| I5407 | 892 | Europe | Serbia | 2544.2 | 22.03 | 44.55 | 8000 | 0.0220 |
| R2.SG | 1062 | Europe | Italy | 3014.0 | 13.54 | 41.96 | 7981 | 0.0219 |
| I4596 | 865 | Europe | Latvia | 3691.3 | 25.13 | 56.28 | 7970 | 0.0217 |
| I4432 | 865 | Europe | Latvia | 3691.3 | 25.13 | 56.28 | 7958 | 0.0272 |
| DA245_noUDG.SG | 87 | Asia | Russia | 6444.1 | 103.70 | 51.70 | 7944 | 0.0178 |
| LEPE48.SG | 988 | Europe | Serbia | 2590.5 | 22.03 | 44.56 | 7944 | 0.0185 |
| I0746 | 580 | Europe | Turkey | 1775.5 | 29.57 | 40.30 | 7928 | 0.0196 |
| irk051_noUDG.SG | 90 | Asia | Russia | 6514.2 | 104.25 | 52.29 | 7922 | 0.0265 |
| I4351_noUDG | 215 | Asia | Iran | 1633.2 | 45.47 | 36.99 | 7920 | 0.0190 |
| I4241 | 215 | Asia | Iran | 1633.2 | 45.47 | 36.99 | 7904 | 0.0177 |
| I5232 | 885 | Europe | Serbia | 2590.5 | 22.01 | 44.60 | 7901 | 0.0175 |
| I0698 | 579 | Europe | Bulgaria | 2168.7 | 25.75 | 42.10 | 7900 | 0.0208 |
| I5427 | 832 | Europe | Greece | 2105.7 | 22.38 | 36.64 | 7883 | 0.0191 |
| FNO001 | 116 | Asia | Russia | 6653.5 | 106.77 | 52.05 | 7881 | 0.0205 |
| La368.SG | 315 | Asia | Laos | 7554.5 | 104.48 | 18.40 | 7872 | 0.0131 |
| I0723_enhanced | 581 | Europe | Turkey | 1775.5 | 29.65 | 40.26 | 7870 | 0.0178 |
| I4873 | 869 | Europe | Serbia | 2544.2 | 22.05 | 44.53 | 7867 | 0.0197 |
| I3948 | 844 | Europe | Croatia | 2810.3 | 16.65 | 43.59 | 7863 | 0.0229 |

| | | | | | | | | |
|---|---|---|---|---|---|---|---|---|
| I0843 | 576 | Europe | Spain | 4552.3 | -5.38 | 42.91 | 7858 | 0.0337 |
| DA249_noUDG.SG | 87 | Asia | Russia | 6444.1 | 103.70 | 51.70 | 7840 | 0.0236 |
| I3947 | 844 | Europe | Croatia | 2810.3 | 16.65 | 43.59 | 7834 | 0.0224 |
| Xiaogao_noUDG | 414 | Asia | China | 7898.0 | 117.60 | 37.90 | 7827 | 0.0186 |
| I4882 | 869 | Europe | Serbia | 2544.2 | 22.05 | 44.53 | 7825 | 0.0266 |
| I4880 | 869 | Europe | Serbia | 2544.2 | 22.05 | 44.53 | 7813 | 0.0299 |
| I0585 | 576 | Europe | Spain | 4552.3 | -5.38 | 42.91 | 7811 | 0.0301 |
| I4878 | 869 | Europe | Serbia | 2544.2 | 22.05 | 44.53 | 7803 | 0.0241 |
| I3433 | 844 | Europe | Croatia | 2810.3 | 16.65 | 43.59 | 7800 | 0.0168 |
| I0676 | 578 | Europe | North Macedonia | 2383.7 | 21.35 | 41.90 | 7792 | 0.0174 |
| I4349_noUDG | 215 | Asia | Iran | 1633.2 | 45.47 | 36.99 | 7747 | 0.0234 |
| I10055 | 584 | Europe | Croatia | 2930.8 | 17.64 | 45.45 | 7727 | 0.0202 |
| I11134 | 602 | Europe | Bulgaria | 2209.4 | 25.88 | 43.16 | 7725 | 0.0153 |
| I11135 | 602 | Europe | Bulgaria | 2209.4 | 25.88 | 43.16 | 7725 | 0.0148 |
| I17981 | 602 | Europe | Bulgaria | 2209.4 | 25.88 | 43.16 | 7725 | 0.0175 |
| DA246_noUDG.SG | 87 | Asia | Russia | 6444.1 | 103.70 | 51.70 | 7723 | 0.0198 |
| R3.SG | 1062 | Europe | Italy | 3014.0 | 13.54 | 41.96 | 7716 | 0.0218 |
| I0015 | 561 | Europe | Sweden | 4310.4 | 15.05 | 58.53 | 7710 | 0.0214 |
| DA247_noUDG.SG | 87 | Asia | Russia | 6444.1 | 103.70 | 51.70 | 7689 | 0.0209 |
| I1878 | 741 | Europe | Hungary | 2919.0 | 18.70 | 46.21 | 7687 | 0.0154 |
| I0014 | 561 | Europe | Sweden | 4310.4 | 15.05 | 58.53 | 7686 | 0.0263 |
| Latvia_HG2_noUDG.SG | 865 | Europe | Latvia | 3691.3 | 25.13 | 56.28 | 7677 | 0.0260 |
| I4551 | 865 | Europe | Latvia | 3691.3 | 25.13 | 56.28 | 7670 | 0.0295 |
| I1507 | 681 | Europe | Hungary | 2958.7 | 20.72 | 47.56 | 7661 | 0.0201 |
| NEO235_noUDG.SG | 343 | Asia | Russia | 9012.2 | 135.40 | 44.50 | 7659 | 0.0216 |
| NEO236_noUDG.SG | 343 | Asia | Russia | 9012.2 | 135.40 | 44.50 | 7650 | 0.0263 |
| DA248_noUDG.SG | 87 | Asia | Russia | 6444.1 | 103.70 | 51.70 | 7646 | 0.0221 |
| I4439 | 865 | Europe | Latvia | 3691.3 | 25.13 | 56.28 | 7646 | 0.0229 |
| MTT001 | 327 | Asia | Azerbaijan | 2035.1 | 45.83 | 40.94 | 7638 | 0.0148 |
| I4595 | 865 | Europe | Latvia | 3691.3 | 25.13 | 56.28 | 7637 | 0.0278 |
| R10.SG | 1062 | Europe | Italy | 3014.0 | 13.54 | 41.96 | 7628 | 0.0225 |

| | | | | | | | | |
|---|---|---|---|---|---|---|---|---|
| I0017 | 561 | Europe | Sweden | 4310.4 | 15.05 | 58.53 | 7624 | 0.0265 |
| I2532 | 803 | Europe | Romania | 2454.9 | 27.01 | 45.47 | 7614 | 0.0127 |
| I2529_enhanced | 579 | Europe | Bulgaria | 2168.7 | 25.75 | 42.10 | 7610 | 0.0252 |
| I1880 | 742 | Europe | Hungary | 2919.0 | 18.58 | 45.99 | 7600 | 0.0227 |
| I4918 | 873 | Europe | Serbia | 2596.3 | 21.08 | 44.49 | 7600 | 0.0303 |
| I0012 | 561 | Europe | Sweden | 4310.4 | 15.05 | 58.53 | 7598 | 0.0229 |
| I1508 | 682 | Europe | Hungary | 2913.1 | 21.53 | 47.32 | 7591 | 0.0211 |
| DA341_noUDG.SG | 90 | Asia | Russia | 6514.2 | 104.25 | 52.29 | 7577 | 0.0190 |
| I1877_noUDG | 741 | Europe | Hungary | 2919.0 | 18.70 | 46.21 | 7567 | 0.0198 |
| I0011 | 561 | Europe | Sweden | 4310.4 | 15.05 | 58.53 | 7564 | 0.0252 |
| I2794 | 830 | Europe | Hungary | 2913.5 | 20.40 | 47.19 | 7564 | 0.0207 |
| I5072 | 877 | Europe | Croatia | 2810.3 | 16.46 | 43.35 | 7545 | 0.0202 |
| STAR1.SG | 1145 | Europe | Serbia | 2690.8 | 20.57 | 44.82 | 7538 | 0.0176 |
| I11696 | 160 | Asia | Mongolia | 6379.3 | 102.70 | 49.39 | 7528 | 0.0220 |
| Asp6.SG | 472 | Europe | Austria | 3275.6 | 16.50 | 48.59 | 7528 | 0.0213 |
| I11698 | 160 | Asia | Mongolia | 6379.3 | 102.70 | 49.39 | 7520 | 0.0257 |
| CBT018 | 501 | Europe | Turkey | 1468.5 | 34.62 | 40.02 | 7516 | 0.0230 |
| I13179 | 182 | Asia | Mongolia | 6449.9 | 103.28 | 49.53 | 7515 | 0.0220 |
| NEO240_noUDG.SG | 343 | Asia | Russia | 9012.2 | 135.40 | 44.50 | 7508 | 0.0238 |
| I13698 | 182 | Asia | Mongolia | 6449.9 | 103.28 | 49.53 | 7504 | 0.0295 |
| I2521 | 602 | Europe | Bulgaria | 2209.4 | 25.88 | 43.16 | 7503 | 0.0182 |
| I11697 | 160 | Asia | Mongolia | 6379.3 | 102.70 | 49.39 | 7500 | 0.0214 |
| VC3-2.SG | 1174 | Europe | Serbia | 2690.8 | 20.62 | 44.76 | 7484 | 0.0174 |
| R9.SG | 1062 | Europe | Italy | 3014.0 | 13.54 | 41.96 | 7481 | 0.0241 |
| I14000 | 182 | Asia | Mongolia | 6449.9 | 103.28 | 49.53 | 7480 | 0.0192 |
| I4434 | 865 | Europe | Latvia | 3691.3 | 25.13 | 56.28 | 7441 | 0.0234 |
| I0633_noUDG | 577 | Europe | Serbia | 2690.8 | 19.75 | 44.90 | 7424 | 0.0193 |
| brn008_noUDG.SG | 35 | Asia | Russia | 7343.5 | 116.99 | 52.23 | 7379 | 0.0251 |
| POT002 | 350 | Asia | Azerbaijan | 2069.9 | 48.65 | 39.52 | 7378 | 0.0157 |
| DA250_noUDG.SG | 87 | Asia | Russia | 6444.1 | 103.70 | 51.70 | 7377 | 0.0187 |
| I2533 | 804 | Europe | Romania | 2448.3 | 23.90 | 44.27 | 7371 | 0.0191 |

| | | | | | | | | |
|---|---|---|---|---|---|---|---|---|
| I4553 | 865 | Europe | Latvia | 3691.3 | 25.13 | 56.28 | 7370 | 0.0239 |
| I4552 | 865 | Europe | Latvia | 3691.3 | 25.13 | 56.28 | 7369 | 0.0264 |
| PEN003 | 1040 | Europe | France | 3583.2 | 7.51 | 43.81 | 7363 | 0.0080 |
| PEN001_real1 | 1040 | Europe | France | 3583.2 | 7.51 | 43.81 | 7361 | 0.0199 |
| I2937 | 832 | Europe | Greece | 2105.7 | 22.38 | 36.64 | 7360 | 0.0241 |
| PEN001_real2 | 1040 | Europe | France | 3583.2 | 7.51 | 43.81 | 7359 | 0.0238 |
| Baojianshan5_M1 | 18 | Asia | China | 7577.5 | 106.85 | 22.34 | 7350 | 0.0230 |
| I2380 | 782 | Europe | Hungary | 3003.9 | 20.58 | 47.78 | 7350 | 0.0218 |
| I4114 | 857 | Europe | Ukraine | 2963.6 | 33.76 | 48.91 | 7350 | 0.0212 |
| I1738 | 714 | Europe | Ukraine | 2908.7 | 35.08 | 48.13 | 7348 | 0.0186 |
| CB13_noUDG.SG | 498 | Europe | Spain | 4140.4 | 1.89 | 41.37 | 7345 | 0.0252 |
| DA252_noUDG.SG | 87 | Asia | Russia | 6444.1 | 103.70 | 51.70 | 7317 | 0.0208 |
| R18.SG | 1088 | Europe | Italy | 3094.7 | 13.03 | 43.72 | 7271 | 0.0192 |
| I4438 | 865 | Europe | Latvia | 3691.3 | 25.13 | 56.28 | 7248 | 0.0190 |
| DA253_noUDG.SG | 87 | Asia | Russia | 6444.1 | 103.70 | 51.70 | 7240 | 0.0215 |
| DA362_noUDG.SG | 87 | Asia | Russia | 6444.1 | 103.70 | 51.70 | 7230 | 0.0230 |
| I1732 | 714 | Europe | Ukraine | 2908.7 | 35.08 | 48.13 | 7218 | 0.0212 |
| R17.SG | 1088 | Europe | Italy | 3094.7 | 13.03 | 43.72 | 7216 | 0.0157 |
| I0409 | 531 | Europe | Spain | 4074.4 | 0.50 | 42.50 | 7210 | 0.0247 |
| R6.SG | 1062 | Europe | Italy | 3014.0 | 13.54 | 41.96 | 7202 | 0.0325 |
| I15818 | 686 | Europe | Czech Republic | 3537.1 | 13.88 | 50.59 | 7200 | 0.0145 |
| I7952 | 946 | Europe | Czech Republic | 3495.2 | 14.25 | 50.08 | 7200 | 0.0193 |
| UzOO77 | 1171 | Europe | Russia | 4337.5 | 35.36 | 62.05 | 7200 | 0.0199 |
| I1895 | 745 | Europe | Hungary | 2878.2 | 18.28 | 45.60 | 7195 | 0.0163 |
| I0412 | 531 | Europe | Spain | 4074.4 | 0.50 | 42.50 | 7177 | 0.0218 |
| DER032 | 519 | Europe | Gernamy | 3796.2 | 10.91 | 51.87 | 7172 | 0.0223 |
| Stuttgart.DG | 562 | Europe | Germany | 3623.0 | 9.18 | 48.78 | 7168 | 0.0230 |
| I6358 | 251 | Asia | Mongolia | 7363.6 | 115.10 | 48.29 | 7167 | 0.0145 |
| I7126 | 619 | Europe | Romania | 2914.2 | 22.40 | 47.75 | 7162 | 0.0213 |
| I2739 | 741 | Europe | Hungary | 2919.0 | 18.70 | 46.21 | 7158 | 0.0231 |
| I5205 | 884 | Europe | Austria | 3275.6 | 16.47 | 48.58 | 7150 | 0.0262 |

| | | | | | | | | |
|---|---|---|---|---|---|---|---|---|
| I5206 | 884 | Europe | Austria | 3275.6 | 16.47 | 48.58 | 7150 | 0.0228 |
| I5207 | 884 | Europe | Austria | 3275.6 | 16.47 | 48.58 | 7150 | 0.0235 |
| I5208 | 884 | Europe | Austria | 3275.6 | 16.47 | 48.58 | 7150 | 0.0202 |
| NE1_noUDG.SG | 620 | Europe | Hungary | 2958.8 | 21.15 | 47.85 | 7146 | 0.0182 |
| I0413 | 531 | Europe | Spain | 4074.4 | 0.50 | 42.50 | 7143 | 0.0210 |
| mur_noUDG.SG | 1006 | Europe | Spain | 4828.8 | -4.30 | 37.54 | 7127 | 0.0264 |
| I0025 | 562 | Europe | Germany | 3623.0 | 9.18 | 48.78 | 7125 | 0.0169 |
| I0026 | 562 | Europe | Germany | 3623.0 | 9.18 | 48.78 | 7125 | 0.0151 |
| I1550 | 563 | Europe | Germany | 3796.2 | 11.05 | 51.90 | 7125 | 0.0160 |
| I26738 | 816 | Europe | Croatia | 2837.4 | 18.06 | 45.15 | 7125 | 0.0158 |
| I2384 | 783 | Europe | Hungary | 2958.8 | 21.43 | 47.86 | 7114 | 0.0170 |
| I0410 | 531 | Europe | Spain | 4074.4 | 0.50 | 42.50 | 7106 | 0.0206 |
| I1498 | 676 | Europe | Hungary | 2913.1 | 21.59 | 47.52 | 7094 | 0.0225 |
| Canes_noUDG.SG | 497 | Europe | Spain | 4538.4 | -4.72 | 43.36 | 7089 | 0.0296 |
| Dil16.SG | 520 | Europe | Germany | 3570.0 | 10.54 | 48.59 | 7085 | 0.0259 |
| Herx.SG | 553 | Europe | Germany | 3712.2 | 8.21 | 49.14 | 7078 | 0.0214 |
| I7399 | 934 | Europe | Croatia | 2784.8 | 18.78 | 45.30 | 7078 | 0.0236 |
| I1499 | 679 | Europe | Hungary | 3004.4 | 21.17 | 48.52 | 7077 | 0.0197 |
| I0054 | 565 | Europe | Germany | 3756.0 | 11.53 | 51.66 | 7074 | 0.0245 |
| I4440 | 865 | Europe | Latvia | 3691.3 | 25.13 | 56.28 | 7074 | 0.0230 |
| R1014_oEEF.SG | 1063 | Europe | Italy | 2990.8 | 13.29 | 41.37 | 7074 | 0.0260 |
| DER022 | 519 | Europe | Gernamy | 3796.2 | 10.91 | 51.87 | 7058 | 0.0209 |
| I1500 | 680 | Europe | Hungary | 2867.4 | 20.83 | 47.17 | 7058 | 0.0154 |
| I1505 | 620 | Europe | Hungary | 2958.8 | 21.15 | 47.85 | 7057 | 0.0174 |
| LBR005 | 985 | Europe | France | 3583.2 | 6.99 | 43.70 | 7051 | 0.0198 |
| I7021 | 266 | Asia | Mongolia | 7224.9 | 113.93 | 48.00 | 7050 | 0.0248 |
| I4196 | 860 | Europe | Hungary | 3004.9 | 18.91 | 47.50 | 7050 | 0.0230 |
| DER024 | 519 | Europe | Gernamy | 3796.2 | 10.91 | 51.87 | 7047 | 0.0215 |
| I0046 | 563 | Europe | Germany | 3796.2 | 11.05 | 51.90 | 7041 | 0.0191 |
| I1496 | 675 | Europe | Hungary | 2913.5 | 19.83 | 47.17 | 7040 | 0.0215 |
| NE19_noUDG | 331 | Asia | China | 8195.5 | 125.75 | 45.96 | 7039 | 0.0199 |

| | | | | | | | | |
|---|---|---|---|---|---|---|---|---|
| I7759 | 197 | Asia | Russia | 6446.8 | 103.66 | 52.92 | 7035 | 0.0281 |
| I4628 | 865 | Europe | Latvia | 3691.3 | 25.13 | 56.28 | 7033 | 0.0272 |
| I0659 | 563 | Europe | Germany | 3796.2 | 11.05 | 51.90 | 7031 | 0.0278 |
| DER009.SG | 519 | Europe | Gernamy | 3796.2 | 10.91 | 51.87 | 7028 | 0.0243 |
| OBN006 | 1027 | Europe | France | 3695.3 | 7.48 | 48.46 | 7026 | 0.0203 |
| I1904 | 746 | Europe | Hungary | 2919.0 | 18.70 | 46.20 | 7010 | 0.0180 |
| I2377 | 780 | Europe | Hungary | 2958.8 | 21.18 | 48.00 | 7001 | 0.0260 |
| I5077 | 817 | Europe | Croatia | 2826.1 | 18.75 | 45.55 | 7001 | 0.0204 |
| LBR002 | 985 | Europe | France | 3583.2 | 6.99 | 43.70 | 6991 | 0.0155 |
| I2379 | 781 | Europe | Hungary | 2958.8 | 21.00 | 47.86 | 6984 | 0.0258 |
| Ess7.SG | 534 | Europe | Germany | 3465.3 | 12.27 | 48.64 | 6975 | 0.0242 |
| DER010 | 519 | Europe | Gernamy | 3796.2 | 10.91 | 51.87 | 6966 | 0.0182 |
| I5069 | 876 | Europe | Austria | 3275.6 | 16.59 | 48.66 | 6961 | 0.0210 |
| DER002.SG | 519 | Europe | Gernamy | 3796.2 | 10.91 | 51.87 | 6950 | 0.0173 |
| DER007 | 519 | Europe | Gernamy | 3796.2 | 10.91 | 51.87 | 6950 | 0.0208 |
| DER012 | 519 | Europe | Gernamy | 3796.2 | 10.91 | 51.87 | 6950 | 0.0184 |
| DER013 | 519 | Europe | Gernamy | 3796.2 | 10.91 | 51.87 | 6950 | 0.0167 |
| DER015 | 519 | Europe | Gernamy | 3796.2 | 10.91 | 51.87 | 6950 | 0.0137 |
| DER023 | 519 | Europe | Gernamy | 3796.2 | 10.91 | 51.87 | 6950 | 0.0200 |
| DER028 | 519 | Europe | Gernamy | 3796.2 | 10.91 | 51.87 | 6950 | 0.0150 |
| DER033 | 519 | Europe | Gernamy | 3796.2 | 10.91 | 51.87 | 6950 | 0.0150 |
| DER034 | 519 | Europe | Gernamy | 3796.2 | 10.91 | 51.87 | 6950 | 0.0190 |
| DER036 | 519 | Europe | Gernamy | 3796.2 | 10.91 | 51.87 | 6950 | 0.0195 |
| I11933 | 620 | Europe | Hungary | 2958.8 | 21.15 | 47.85 | 6950 | 0.0216 |
| I0100 | 563 | Europe | Germany | 3796.2 | 11.05 | 51.90 | 6944 | 0.0219 |
| XW-M1R18_noUDG | 415 | Asia | China | 7447.8 | 110.98 | 34.56 | 6927 | 0.0244 |
| IUO001 | 284 | Asia | Russia | 6661.3 | 106.26 | 54.87 | 6905 | 0.0231 |
| NE29_noUDG | 333 | Asia | China | 8270.0 | 126.37 | 45.93 | 6877 | 0.0213 |
| KAG001 | 288 | Asia | Russia | 6589.0 | 105.89 | 53.95 | 6856 | 0.0249 |
| PG2001 | 348 | Asia | Russia | 2254.5 | 43.35 | 43.82 | 6850 | 0.0233 |
| I4063 | 855 | Europe | Italy | 3070.3 | 12.92 | 37.72 | 6832 | 0.0250 |

| | | | | | | | | |
|---|---|---|---|---|---|---|---|---|
| I3355 | 164 | Asia | Russia | 8782.2 | 131.28 | 42.79 | 6821 | 0.0180 |
| I4062 | 855 | Europe | Italy | 3070.3 | 12.92 | 37.72 | 6815 | 0.0186 |
| I4065 | 855 | Europe | Italy | 3070.3 | 12.92 | 37.72 | 6815 | 0.0170 |
| LBR004 | 985 | Europe | France | 3583.2 | 6.99 | 43.70 | 6797 | 0.0186 |
| I0122 | 125 | Asia | Russia | 3323.9 | 48.08 | 52.35 | 6788 | 0.0289 |
| kol6_noUDG.SG | 975 | Europe | Czech Republic | 3449.6 | 15.20 | 50.03 | 6775 | 0.0261 |
| Veibri4 | 1175 | Europe | Estonia | 3852.9 | 26.46 | 58.20 | 6760 | 0.0292 |
| LBR001 | 985 | Europe | France | 3583.2 | 6.99 | 43.70 | 6740 | 0.0280 |
| I4064 | 855 | Europe | Italy | 3070.3 | 12.92 | 37.72 | 6718 | 0.0194 |
| I4441 | 865 | Europe | Latvia | 3691.3 | 25.13 | 56.28 | 6718 | 0.0223 |
| HGO-26.SG | 554 | Europe | Hungary | 2868.4 | 20.52 | 46.39 | 6700 | 0.0238 |
| I0449 | 574 | Europe | Hungary | 2868.4 | 20.28 | 46.36 | 6700 | 0.0210 |
| OBN010 | 1027 | Europe | France | 3695.3 | 7.48 | 48.46 | 6700 | 0.0203 |
| VM-33.SG | 1254 | Europe | Hungary | 2867.4 | 21.24 | 47.08 | 6700 | 0.0267 |
| GRG027 | 545 | Europe | France | 3887.4 | 3.56 | 47.97 | 6690 | 0.0263 |
| I4305 | 862 | Europe | France | 3635.6 | 5.86 | 43.46 | 6680 | 0.0201 |
| LBR003 | 985 | Europe | France | 3583.2 | 6.99 | 43.70 | 6680 | 0.0147 |
| I1662 | 205 | Asia | Iran | 1729.2 | 47.96 | 34.50 | 6677 | 0.0142 |
| I1190 | 164 | Asia | Russia | 8782.2 | 131.28 | 42.79 | 6675 | 0.0180 |
| GRG028 | 545 | Europe | France | 3887.4 | 3.56 | 47.97 | 6671 | 0.0220 |
| KGH6.SG | 966 | Europe | Ireland | 4900.0 | -8.33 | 52.60 | 6658 | 0.0219 |
| I4304 | 862 | Europe | France | 3635.6 | 5.86 | 43.46 | 6637 | 0.0194 |
| POP07_noUDG.SG | 1051 | Europe | Croatia | 2878.2 | 18.57 | 45.75 | 6635 | 0.0184 |
| TU919_SX33 | 1168 | Europe | France | 3640.9 | 7.68 | 48.55 | 6625 | 0.0112 |
| I2431 | 788 | Europe | Bulgaria | 2209.4 | 26.72 | 43.10 | 6624 | 0.0221 |
| I4303 | 862 | Europe | France | 3635.6 | 5.86 | 43.46 | 6624 | 0.0196 |
| GRG057 | 545 | Europe | France | 3887.4 | 3.56 | 47.97 | 6615 | 0.0214 |
| I4111 | 857 | Europe | Ukraine | 2963.6 | 33.76 | 48.91 | 6593 | 0.0234 |
| STB002 | 392 | Asia | Russia | 6652.4 | 106.05 | 53.15 | 6587 | 0.0217 |
| I26741 | 817 | Europe | Croatia | 2826.1 | 18.75 | 45.55 | 6585 | 0.0190 |
| I5078 | 817 | Europe | Croatia | 2826.1 | 18.75 | 45.55 | 6571 | 0.0190 |

| | | | | | | | | |
|---|---|---|---|---|---|---|---|---|
| Donkalnis6 | 523 | Europe | Lithuania | 3744.2 | 22.42 | 55.81 | 6569 | 0.0156 |
| I0634_noUDG | 577 | Europe | Serbia | 2690.8 | 19.75 | 44.90 | 6560 | 0.0213 |
| brn003_noUDG.SG | 34 | Asia | Russia | 7142.2 | 113.21 | 52.07 | 6550 | 0.0280 |
| I1903 | 746 | Europe | Hungary | 2919.0 | 18.70 | 46.20 | 6550 | 0.0171 |
| OBN011 | 1027 | Europe | France | 3695.3 | 7.48 | 48.46 | 6549 | 0.0193 |
| OBN003 | 1027 | Europe | France | 3695.3 | 7.48 | 48.46 | 6544 | 0.0168 |
| OBN008 | 1027 | Europe | France | 3695.3 | 7.48 | 48.46 | 6543 | 0.0230 |
| I1661 | 205 | Asia | Iran | 1729.2 | 47.96 | 34.50 | 6539 | 0.0207 |
| FLR004 | 537 | Europe | France | 4224.4 | -0.38 | 49.19 | 6512 | 0.0187 |
| I3708 | 832 | Europe | Greece | 2105.7 | 22.38 | 36.64 | 6500 | 0.0162 |
| POP14_noUDG.SG | 1051 | Europe | Croatia | 2878.2 | 18.57 | 45.75 | 6488 | 0.0216 |
| I1131 | 577 | Europe | Serbia | 2690.8 | 19.75 | 44.90 | 6482 | 0.0216 |
| I2056 | 760 | Europe | Russia | 2256.7 | 40.22 | 44.25 | 6466 | 0.0169 |
| ANY-4027.SG | 467 | Europe | Hungary | 2870.8 | 18.80 | 46.22 | 6450 | 0.0172 |
| GRG003 | 545 | Europe | France | 3887.4 | 3.56 | 47.97 | 6450 | 0.0252 |
| GRG008 | 545 | Europe | France | 3887.4 | 3.56 | 47.97 | 6450 | 0.0227 |
| GRG015 | 545 | Europe | France | 3887.4 | 3.56 | 47.97 | 6450 | 0.0186 |
| GRG019 | 545 | Europe | France | 3887.4 | 3.56 | 47.97 | 6450 | 0.0243 |
| GRG021 | 545 | Europe | France | 3887.4 | 3.56 | 47.97 | 6450 | 0.0246 |
| GRG022 | 545 | Europe | France | 3887.4 | 3.56 | 47.97 | 6450 | 0.0166 |
| GRG025 | 545 | Europe | France | 3887.4 | 3.56 | 47.97 | 6450 | 0.0210 |
| GRG035 | 545 | Europe | France | 3887.4 | 3.56 | 47.97 | 6450 | 0.0180 |
| GRG041 | 545 | Europe | France | 3887.4 | 3.56 | 47.97 | 6450 | 0.0233 |
| GRG043 | 545 | Europe | France | 3887.4 | 3.56 | 47.97 | 6450 | 0.0221 |
| GRG052 | 545 | Europe | France | 3887.4 | 3.56 | 47.97 | 6450 | 0.0149 |
| I10545 | 594 | Europe | Turkey | 1775.5 | 29.30 | 40.48 | 6450 | 0.0229 |
| I14190 | 609 | Europe | Czech Republic | 3495.2 | 14.36 | 50.05 | 6450 | 0.0196 |
| POP02_noUDG.SG | 1051 | Europe | Croatia | 2878.2 | 18.57 | 45.75 | 6450 | 0.0186 |
| POP04_noUDG.SG | 1051 | Europe | Croatia | 2878.2 | 18.57 | 45.75 | 6450 | 0.0118 |
| POP05_noUDG.SG | 1051 | Europe | Croatia | 2878.2 | 18.57 | 45.75 | 6450 | 0.0223 |
| POP06_noUDG.SG | 1051 | Europe | Croatia | 2878.2 | 18.57 | 45.75 | 6450 | 0.0215 |

| | | | | | | | | |
|---|---|---|---|---|---|---|---|---|
| POP08_noUDG.SG | 1051 | Europe | Croatia | 2878.2 | 18.57 | 45.75 | 6450 | 0.0231 |
| POP09_noUDG.SG | 1051 | Europe | Croatia | 2878.2 | 18.57 | 45.75 | 6450 | 0.0234 |
| POP12_noUDG.SG | 1051 | Europe | Croatia | 2878.2 | 18.57 | 45.75 | 6450 | 0.0135 |
| POP13_noUDG.SG | 1051 | Europe | Croatia | 2878.2 | 18.57 | 45.75 | 6450 | 0.0247 |
| POP16_noUDG.SG | 1051 | Europe | Croatia | 2878.2 | 18.57 | 45.75 | 6450 | 0.0182 |
| POP19_noUDG.SG | 1051 | Europe | Croatia | 2878.2 | 18.57 | 45.75 | 6450 | 0.0208 |
| POP24_noUDG.SG | 1051 | Europe | Croatia | 2878.2 | 18.57 | 45.75 | 6450 | 0.0239 |
| POP27_noUDG.SG | 1051 | Europe | Croatia | 2878.2 | 18.57 | 45.75 | 6450 | 0.0188 |
| POP36_noUDG.SG | 1051 | Europe | Croatia | 2878.2 | 18.57 | 45.75 | 6450 | 0.0224 |
| I2430 | 786 | Europe | Bulgaria | 2161.8 | 26.98 | 43.06 | 6448 | 0.0162 |
| I7197 | 871 | Europe | Czech Republic | 3495.2 | 14.37 | 50.05 | 6438 | 0.0215 |
| FLR001 | 537 | Europe | France | 4224.4 | -0.38 | 49.19 | 6414 | 0.0125 |
| GRG016 | 545 | Europe | France | 3887.4 | 3.56 | 47.97 | 6407 | 0.0239 |
| OBN009 | 1027 | Europe | France | 3695.3 | 7.48 | 48.46 | 6398 | 0.0220 |
| OBN007 | 1027 | Europe | France | 3695.3 | 7.48 | 48.46 | 6377 | 0.0287 |
| FLR003 | 537 | Europe | France | 4224.4 | -0.38 | 49.19 | 6376 | 0.0158 |
| I4894 | 870 | Europe | Czech Republic | 3495.2 | 14.46 | 50.12 | 6371 | 0.0227 |
| I1495 | 675 | Europe | Hungary | 2913.5 | 19.83 | 47.17 | 6367 | 0.0213 |
| TU877_SX11 | 1166 | Europe | Switzerland | 3632.1 | 7.25 | 47.01 | 6355 | 0.0236 |
| Klei10_noUDG.SG | 969 | Europe | Greece | 2217.7 | 21.86 | 40.43 | 6348 | 0.0275 |
| Pal7_noUDG.SG | 1034 | Europe | Greece | 2206.0 | 22.01 | 39.75 | 6348 | 0.0234 |
| I4893 | 870 | Europe | Czech Republic | 3495.2 | 14.46 | 50.12 | 6345 | 0.0257 |
| TU875_SX9 | 1166 | Europe | Switzerland | 3632.1 | 7.25 | 47.01 | 6338 | 0.0171 |
| NE9_noUDG | 342 | Asia | China | 8195.5 | 125.80 | 46.01 | 6331 | 0.0190 |
| FLR007 | 537 | Europe | France | 4224.4 | -0.38 | 49.19 | 6326 | 0.0141 |
| I2427 | 787 | Europe | Bulgaria | 2209.4 | 26.77 | 43.06 | 6326 | 0.0216 |
| POP33_noUDG.SG | 1051 | Europe | Croatia | 2878.2 | 18.57 | 45.75 | 6324 | 0.0192 |
| FLR005 | 537 | Europe | France | 4224.4 | -0.38 | 49.19 | 6301 | 0.0190 |
| I2424 | 786 | Europe | Bulgaria | 2161.8 | 26.98 | 43.06 | 6288 | 0.0135 |
| I2793 | 830 | Europe | Hungary | 2913.5 | 20.40 | 47.19 | 6273 | 0.0244 |
| N22_noUDG.SG | 1012 | Europe | Poland | 3539.2 | 18.90 | 52.61 | 6263 | 0.0198 |

| | | | | | | | | |
|---|---|---|---|---|---|---|---|---|
| Spiginas1 | 1141 | Europe | Lithuania | 3744.2 | 22.42 | 55.77 | 6263 | 0.0227 |
| N25_noUDG.SG | 1013 | Europe | Poland | 3539.2 | 18.80 | 52.61 | 6250 | 0.0215 |
| N27_noUDG.SG | 1013 | Europe | Poland | 3539.2 | 18.80 | 52.61 | 6250 | 0.0197 |
| N36_noUDG.SG | 1014 | Europe | Poland | 3232.5 | 21.38 | 50.67 | 6250 | 0.0199 |
| VJ1001 | 409 | Asia | Russia | 2254.5 | 43.16 | 44.02 | 6230 | 0.0215 |
| PRI001 | 1054 | Europe | France | 4155.8 | -0.48 | 46.15 | 6218 | 0.0191 |
| I13838 | 653 | Europe | Albania | 2315.6 | 20.98 | 40.67 | 6200 | 0.0192 |
| I4088 | 619 | Europe | Romania | 2914.2 | 22.40 | 47.75 | 6191 | 0.0214 |
| I1907 | 749 | Europe | Hungary | 3098.7 | 17.36 | 47.64 | 6183 | 0.0214 |
| N26_noUDG.SG | 1014 | Europe | Poland | 3232.5 | 21.38 | 50.67 | 6177 | 0.0219 |
| I4437 | 865 | Europe | Latvia | 3691.3 | 25.13 | 56.28 | 6164 | 0.0294 |
| I1634 | 192 | Asia | Armenia | 1945.9 | 45.20 | 39.73 | 6156 | 0.0225 |
| I11929 | 620 | Europe | Hungary | 2958.8 | 21.15 | 47.85 | 6150 | 0.0195 |
| Kretuonas2 | 981 | Europe | Lithuania | 3562.8 | 26.10 | 55.26 | 6150 | 0.0258 |
| Kretuonas4 | 981 | Europe | Lithuania | 3562.8 | 26.10 | 55.26 | 6150 | 0.0232 |
| VLI004 | 1253 | Europe | Czech Republic | 3537.1 | 14.45 | 50.37 | 6141 | 0.0205 |
| N31_noUDG.SG | 1013 | Europe | Poland | 3539.2 | 18.80 | 52.61 | 6130 | 0.0228 |
| I17914 | 718 | Europe | Serbia | 2823.1 | 20.27 | 46.02 | 6125 | 0.0136 |
| I17915 | 718 | Europe | Serbia | 2823.1 | 20.27 | 46.02 | 6125 | 0.0246 |
| I4436 | 865 | Europe | Latvia | 3691.3 | 25.13 | 56.28 | 6109 | 0.0226 |
| I13840 | 653 | Europe | Albania | 2315.6 | 20.98 | 40.67 | 6102 | 0.0216 |
| I7550 | 936 | Europe | Spain | 5078.6 | -6.21 | 36.44 | 6100 | 0.0177 |
| I8134 | 936 | Europe | Spain | 5078.6 | -6.21 | 36.44 | 6100 | 0.0173 |
| I1631 | 192 | Asia | Armenia | 1945.9 | 45.20 | 39.73 | 6099 | 0.0246 |
| I10293 | 584 | Europe | Croatia | 2930.8 | 17.64 | 45.45 | 6093 | 0.0254 |
| I7129 | 619 | Europe | Romania | 2914.2 | 22.40 | 47.75 | 6090 | 0.0221 |
| PG2004 | 348 | Asia | Russia | 2254.5 | 43.35 | 43.82 | 6088 | 0.0220 |
| GRG032 | 545 | Europe | France | 3887.4 | 3.56 | 47.97 | 6088 | 0.0229 |
| I15623 | 619 | Europe | Romania | 2914.2 | 22.40 | 47.75 | 6088 | 0.0135 |
| I7132 | 619 | Europe | Romania | 2914.2 | 22.40 | 47.75 | 6086 | 0.0189 |
| I7042 | 930 | Europe | Hungary | 3004.9 | 19.05 | 47.43 | 6086 | 0.0198 |

| | | | | | | | | |
|---|---|---|---|---|---|---|---|---|
| TUC007 | 1169 | Europe | Czech Republic | 3495.2 | 14.27 | 50.14 | 6080 | 0.0178 |
| I7130 | 619 | Europe | Romania | 2914.2 | 22.40 | 47.75 | 6078 | 0.0213 |
| I7133 | 619 | Europe | Romania | 2914.2 | 22.40 | 47.75 | 6078 | 0.0188 |
| I1632 | 192 | Asia | Armenia | 1945.9 | 45.20 | 39.73 | 6076 | 0.0249 |
| NER001 | 1017 | Europe | Czech Republic | 3495.2 | 14.52 | 50.27 | 6076 | 0.0276 |
| I7136 | 619 | Europe | Romania | 2914.2 | 22.40 | 47.75 | 6074 | 0.0208 |
| N28_noUDG.SG | 1013 | Europe | Poland | 3539.2 | 18.80 | 52.61 | 6074 | 0.0250 |
| I4627 | 865 | Europe | Latvia | 3691.3 | 25.13 | 56.28 | 6071 | 0.0226 |
| I10045 | 584 | Europe | Croatia | 2930.8 | 17.64 | 45.45 | 6050 | 0.0227 |
| I10046 | 584 | Europe | Croatia | 2930.8 | 17.64 | 45.45 | 6050 | 0.0241 |
| I10047 | 584 | Europe | Croatia | 2930.8 | 17.64 | 45.45 | 6050 | 0.0168 |
| I10048 | 584 | Europe | Croatia | 2930.8 | 17.64 | 45.45 | 6050 | 0.0223 |
| I10049 | 584 | Europe | Croatia | 2930.8 | 17.64 | 45.45 | 6050 | 0.0205 |
| I10050 | 584 | Europe | Croatia | 2930.8 | 17.64 | 45.45 | 6050 | 0.0194 |
| I10053 | 584 | Europe | Croatia | 2930.8 | 17.64 | 45.45 | 6050 | 0.0149 |
| I10054 | 584 | Europe | Croatia | 2930.8 | 17.64 | 45.45 | 6050 | 0.0244 |
| I10056 | 584 | Europe | Croatia | 2930.8 | 17.64 | 45.45 | 6050 | 0.0219 |
| I10057 | 584 | Europe | Croatia | 2930.8 | 17.64 | 45.45 | 6050 | 0.0228 |
| I10061 | 584 | Europe | Croatia | 2930.8 | 17.64 | 45.45 | 6050 | 0.0218 |
| I10062 | 584 | Europe | Croatia | 2930.8 | 17.64 | 45.45 | 6050 | 0.0179 |
| I10063 | 584 | Europe | Croatia | 2930.8 | 17.64 | 45.45 | 6050 | 0.0255 |
| I10064 | 584 | Europe | Croatia | 2930.8 | 17.64 | 45.45 | 6050 | 0.0240 |
| I10065 | 584 | Europe | Croatia | 2930.8 | 17.64 | 45.45 | 6050 | 0.0154 |
| I10068 | 584 | Europe | Croatia | 2930.8 | 17.64 | 45.45 | 6050 | 0.0252 |
| I10069 | 584 | Europe | Croatia | 2930.8 | 17.64 | 45.45 | 6050 | 0.0229 |
| I10070 | 584 | Europe | Croatia | 2930.8 | 17.64 | 45.45 | 6050 | 0.0157 |
| I10072 | 584 | Europe | Croatia | 2930.8 | 17.64 | 45.45 | 6050 | 0.0300 |
| I10271 | 584 | Europe | Croatia | 2930.8 | 17.64 | 45.45 | 6050 | 0.0170 |
| I10273 | 584 | Europe | Croatia | 2930.8 | 17.64 | 45.45 | 6050 | 0.0178 |
| I10294 | 584 | Europe | Croatia | 2930.8 | 17.64 | 45.45 | 6050 | 0.0263 |
| I10297 | 584 | Europe | Croatia | 2930.8 | 17.64 | 45.45 | 6050 | 0.0275 |

| | | | | | | | | |
|---|---|---|---|---|---|---|---|---|
| I10299 | 584 | Europe | Croatia | 2930.8 | 17.64 | 45.45 | 6050 | 0.0228 |
| I10303 | 584 | Europe | Croatia | 2930.8 | 17.64 | 45.45 | 6050 | 0.0237 |
| I1908 | 750 | Europe | Hungary | 3011.4 | 17.24 | 46.71 | 6050 | 0.0213 |
| I4189 | 859 | Europe | Hungary | 2870.8 | 18.73 | 46.20 | 6050 | 0.0221 |
| I5766 | 243 | Asia | Russia | 4467.8 | 65.36 | 57.36 | 6049 | 0.0255 |
| I16444 | 693 | Europe | Channel Islands | 4383.6 | -2.51 | 49.50 | 6048 | 0.0240 |
| I7128 | 619 | Europe | Romania | 2914.2 | 22.40 | 47.75 | 6046 | 0.0235 |
| I7187 | 657 | Europe | Czech Republic | 3495.2 | 14.22 | 50.15 | 6044 | 0.0241 |
| I2783 | 827 | Europe | Hungary | 2870.8 | 19.05 | 46.33 | 6010 | 0.0193 |
| I7127 | 619 | Europe | Romania | 2914.2 | 22.40 | 47.75 | 6006 | 0.0228 |
| SRA62.SG | 1142 | Europe | Ireland | 4945.5 | -8.34 | 54.30 | 6001 | 0.0261 |
| Latvia_MN2_noUDG.SG | 865 | Europe | Latvia | 3691.3 | 25.13 | 56.28 | 5997 | 0.0189 |
| FLR002 | 537 | Europe | France | 4224.4 | -0.38 | 49.19 | 5972 | 0.0168 |
| I7137 | 619 | Europe | Romania | 2914.2 | 22.40 | 47.75 | 5972 | 0.0235 |
| I4435 | 865 | Europe | Latvia | 3691.3 | 25.13 | 56.28 | 5972 | 0.0192 |
| irk030_noUDG.SG | 277 | Asia | Russia | 6592.3 | 105.21 | 54.37 | 5969 | 0.0230 |
| WGM35_noUDG | 413 | Asia | China | 7690.3 | 113.40 | 34.82 | 5950 | 0.0220 |
| I0644 | 424 | Both | Israel | 629.1 | 35.33 | 32.97 | 5950 | 0.0235 |
| I1152 | 424 | Both | Israel | 629.1 | 35.33 | 32.97 | 5950 | 0.0134 |
| I1160 | 424 | Both | Israel | 629.1 | 35.33 | 32.97 | 5950 | 0.0130 |
| I1166 | 424 | Both | Israel | 629.1 | 35.33 | 32.97 | 5950 | 0.0136 |
| I1168 | 424 | Both | Israel | 629.1 | 35.33 | 32.97 | 5950 | 0.0207 |
| I1169 | 424 | Both | Israel | 629.1 | 35.33 | 32.97 | 5950 | 0.0167 |
| I1178 | 424 | Both | Israel | 629.1 | 35.33 | 32.97 | 5950 | 0.0193 |
| I11902 | 619 | Europe | Romania | 2914.2 | 22.40 | 47.75 | 5950 | 0.0194 |
| I11906 | 619 | Europe | Romania | 2914.2 | 22.40 | 47.75 | 5950 | 0.0177 |
| I14157 | 619 | Europe | Romania | 2914.2 | 22.40 | 47.75 | 5950 | 0.0232 |
| I14158 | 619 | Europe | Romania | 2914.2 | 22.40 | 47.75 | 5950 | 0.0241 |
| I14159 | 619 | Europe | Romania | 2914.2 | 22.40 | 47.75 | 5950 | 0.0227 |
| I14160 | 619 | Europe | Romania | 2914.2 | 22.40 | 47.75 | 5950 | 0.0191 |
| I14161 | 619 | Europe | Romania | 2914.2 | 22.40 | 47.75 | 5950 | 0.0281 |

| | | | | | | | | |
|---|---|---|---|---|---|---|---|---|
| I14162 | 619 | Europe | Romania | 2914.2 | 22.40 | 47.75 | 5950 | 0.0244 |
| I14163 | 619 | Europe | Romania | 2914.2 | 22.40 | 47.75 | 5950 | 0.0239 |
| I14164 | 619 | Europe | Romania | 2914.2 | 22.40 | 47.75 | 5950 | 0.0239 |
| I14165 | 619 | Europe | Romania | 2914.2 | 22.40 | 47.75 | 5950 | 0.0202 |
| I14166 | 619 | Europe | Romania | 2914.2 | 22.40 | 47.75 | 5950 | 0.0130 |
| I15616 | 619 | Europe | Romania | 2914.2 | 22.40 | 47.75 | 5950 | 0.0165 |
| I15617 | 619 | Europe | Romania | 2914.2 | 22.40 | 47.75 | 5950 | 0.0201 |
| I15618 | 619 | Europe | Romania | 2914.2 | 22.40 | 47.75 | 5950 | 0.0246 |
| I15619 | 619 | Europe | Romania | 2914.2 | 22.40 | 47.75 | 5950 | 0.0229 |
| I15620 | 619 | Europe | Romania | 2914.2 | 22.40 | 47.75 | 5950 | 0.0204 |
| I15621 | 619 | Europe | Romania | 2914.2 | 22.40 | 47.75 | 5950 | 0.0154 |
| I15622 | 619 | Europe | Romania | 2914.2 | 22.40 | 47.75 | 5950 | 0.0180 |
| I18114 | 619 | Europe | Romania | 2914.2 | 22.40 | 47.75 | 5950 | 0.0194 |
| I18115 | 619 | Europe | Romania | 2914.2 | 22.40 | 47.75 | 5950 | 0.0211 |
| I18116 | 619 | Europe | Romania | 2914.2 | 22.40 | 47.75 | 5950 | 0.0234 |
| I20806 | 619 | Europe | Romania | 2914.2 | 22.40 | 47.75 | 5950 | 0.0210 |
| I20809 | 619 | Europe | Romania | 2914.2 | 22.40 | 47.75 | 5950 | 0.0152 |
| I20810 | 619 | Europe | Romania | 2914.2 | 22.40 | 47.75 | 5950 | 0.0160 |
| I7131 | 619 | Europe | Romania | 2914.2 | 22.40 | 47.75 | 5950 | 0.0147 |
| I7135 | 619 | Europe | Romania | 2914.2 | 22.40 | 47.75 | 5950 | 0.0215 |
| I20502 | 759 | Europe | Croatia | 2837.4 | 18.40 | 45.31 | 5950 | 0.0203 |
| LugarCanto41_noUDG.SG | 993 | Europe | Portugal | 5010.4 | -8.82 | 39.41 | 5950 | 0.0213 |
| LugarCanto42_noUDG.SG | 993 | Europe | Portugal | 5010.4 | -8.82 | 39.41 | 5950 | 0.0206 |
| LugarCanto44_noUDG.SG | 993 | Europe | Portugal | 5010.4 | -8.82 | 39.41 | 5950 | 0.0211 |
| I4554 | 865 | Europe | Latvia | 3691.3 | 25.13 | 56.28 | 5946 | 0.0194 |
| I1407 | 192 | Asia | Armenia | 1945.9 | 45.20 | 39.73 | 5875 | 0.0248 |
| I3709 | 832 | Europe | Greece | 2105.7 | 22.38 | 36.64 | 5860 | 0.0173 |
| I14167 | 657 | Europe | Czech Republic | 3495.2 | 14.22 | 50.15 | 5850 | 0.0255 |
| I14168 | 657 | Europe | Czech Republic | 3495.2 | 14.22 | 50.15 | 5850 | 0.0198 |
| I14169 | 657 | Europe | Czech Republic | 3495.2 | 14.22 | 50.15 | 5850 | 0.0220 |
| I14170 | 657 | Europe | Czech Republic | 3495.2 | 14.22 | 50.15 | 5850 | 0.0249 |

| | | | | | | | | |
|---|---|---|---|---|---|---|---|---|
| I14171 | 657 | Europe | Czech Republic | 3495.2 | 14.22 | 50.15 | 5850 | 0.0216 |
| I14172 | 657 | Europe | Czech Republic | 3495.2 | 14.22 | 50.15 | 5850 | 0.0213 |
| I14173 | 657 | Europe | Czech Republic | 3495.2 | 14.22 | 50.15 | 5850 | 0.0172 |
| I14174 | 657 | Europe | Czech Republic | 3495.2 | 14.22 | 50.15 | 5850 | 0.0286 |
| I14175 | 657 | Europe | Czech Republic | 3495.2 | 14.22 | 50.15 | 5850 | 0.0234 |
| I16121 | 657 | Europe | Czech Republic | 3495.2 | 14.22 | 50.15 | 5850 | 0.0192 |
| I16122 | 657 | Europe | Czech Republic | 3495.2 | 14.22 | 50.15 | 5850 | 0.0291 |
| I1674 | 205 | Asia | Iran | 1729.2 | 47.96 | 34.50 | 5831 | 0.0141 |
| I1665 | 205 | Asia | Iran | 1729.2 | 47.96 | 34.50 | 5819 | 0.0195 |
| SUC008 | 1093 | Europe | Italy | 3570.8 | 8.44 | 40.81 | 5817 | 0.0191 |
| ELT006_merged | 531 | Europe | Spain | 4074.4 | 0.50 | 42.50 | 5809 | 0.0162 |
| I6760_noUDG.SG | 922 | Europe | United Kingdom | 4467.2 | -1.85 | 51.84 | 5781 | 0.0250 |
| I1584_noUDG | 687 | Europe | Turkey | 1775.5 | 29.56 | 40.30 | 5770 | 0.0160 |
| ART042 | 470 | Europe | Turkey | 1427.6 | 38.36 | 38.38 | 5767 | 0.0172 |
| ELT002_merged | 531 | Europe | Spain | 4074.4 | 0.50 | 42.50 | 5753 | 0.0167 |
| PN10_PN113.SG | 1047 | Europe | Ireland | 4982.1 | -9.14 | 53.05 | 5746 | 0.0230 |
| PN04.SG | 1047 | Europe | Ireland | 4982.1 | -9.14 | 53.05 | 5742 | 0.0252 |
| I3920 | 832 | Europe | Greece | 2105.7 | 22.38 | 36.64 | 5734 | 0.0187 |
| PN05.SG | 1047 | Europe | Ireland | 4982.1 | -9.14 | 53.05 | 5729 | 0.0221 |
| I7186 | 657 | Europe | Czech Republic | 3495.2 | 14.22 | 50.15 | 5713 | 0.0250 |
| PN107.SG | 1047 | Europe | Ireland | 4982.1 | -9.14 | 53.05 | 5705 | 0.0232 |
| I0406 | 573 | Europe | Spain | 4367.0 | -2.33 | 41.25 | 5700 | 0.0201 |
| I0407 | 573 | Europe | Spain | 4367.0 | -2.33 | 41.25 | 5700 | 0.0263 |
| I6762_noUDG.SG | 924 | Europe | United Kingdom | 4382.1 | -1.45 | 51.56 | 5700 | 0.0192 |
| I0408 | 573 | Europe | Spain | 4367.0 | -2.33 | 41.25 | 5686 | 0.0264 |
| I2788 | 829 | Europe | Hungary | 2913.5 | 20.00 | 47.19 | 5685 | 0.0231 |
| I1565 | 684 | Europe | Germany | 3943.7 | 7.55 | 51.36 | 5674 | 0.0254 |
| I16165 | 691 | Europe | Italy | 3570.9 | 8.64 | 40.41 | 5671 | 0.0246 |
| VERT035 | 1177 | Europe | Ukraine | 2837.9 | 25.85 | 48.78 | 5671 | 0.0226 |
| prs002_noUDG.SG | 1055 | Europe | Ireland | 4945.5 | -8.56 | 54.25 | 5665 | 0.0230 |
| I7191 | 657 | Europe | Czech Republic | 3495.2 | 14.22 | 50.15 | 5657 | 0.0254 |

| | | | | | | | | |
|---|---|---|---|---|---|---|---|---|
| I3005 | 835 | Europe | United Kingdom | 4509.5 | -2.75 | 51.32 | 5657 | 0.0154 |
| ERM001 | 113 | Asia | Mongolia | 6311.2 | 101.31 | 48.51 | 5653 | 0.0210 |
| I2789 | 829 | Europe | Hungary | 2913.5 | 20.00 | 47.19 | 5650 | 0.0162 |
| I6751_noUDG.SG | 918 | Europe | United Kingdom | 4423.8 | -1.73 | 51.09 | 5650 | 0.0244 |
| VERT031 | 1177 | Europe | Ukraine | 2837.9 | 25.85 | 48.78 | 5646 | 0.0179 |
| VERT100B | 1177 | Europe | Ukraine | 2837.9 | 25.85 | 48.78 | 5646 | 0.0246 |
| VERT104B | 1177 | Europe | Ukraine | 2837.9 | 25.85 | 48.78 | 5642 | 0.0207 |
| I2633 | 814 | Europe | United Kingdom | 5167.4 | -3.60 | 58.53 | 5639 | 0.0236 |
| TRPTN_613_noUDG.SG | 1164 | Europe | United Kingdom | 4389.8 | 0.10 | 52.17 | 5639 | 0.0183 |
| I4089 | 619 | Europe | Romania | 2914.2 | 22.40 | 47.75 | 5636 | 0.0176 |
| KOB002 | 974 | Europe | Czech Republic | 3449.6 | 15.23 | 50.01 | 5630 | 0.0245 |
| I2790 | 829 | Europe | Hungary | 2913.5 | 20.00 | 47.19 | 5625 | 0.0211 |
| I7193 | 657 | Europe | Czech Republic | 3495.2 | 14.22 | 50.15 | 5621 | 0.0238 |
| DA342_noUDG.SG | 91 | Asia | Russia | 6449.2 | 103.37 | 53.19 | 5617 | 0.0235 |
| HOL001 | 557 | Europe | Czech Republic | 3495.2 | 14.29 | 50.20 | 5613 | 0.0237 |
| I7188 | 657 | Europe | Czech Republic | 3495.2 | 14.22 | 50.15 | 5610 | 0.0254 |
| ASH3.SG | 471 | Europe | Ireland | 4900.0 | -8.19 | 52.93 | 5609 | 0.0254 |
| I3356 | 164 | Asia | Russia | 8782.2 | 131.28 | 42.79 | 5605 | 0.0237 |
| I5359 | 886 | Europe | United Kingdom | 4509.5 | -3.31 | 51.45 | 5600 | 0.0207 |
| VERT028 | 1177 | Europe | Ukraine | 2837.9 | 25.85 | 48.78 | 5600 | 0.0229 |
| VERT029 | 1177 | Europe | Ukraine | 2837.9 | 25.85 | 48.78 | 5600 | 0.0234 |
| VERT030 | 1177 | Europe | Ukraine | 2837.9 | 25.85 | 48.78 | 5600 | 0.0278 |
| VERT033 | 1177 | Europe | Ukraine | 2837.9 | 25.85 | 48.78 | 5600 | 0.0225 |
| VERT103B | 1177 | Europe | Ukraine | 2837.9 | 25.85 | 48.78 | 5600 | 0.0211 |
| VERT105B | 1177 | Europe | Ukraine | 2837.9 | 25.85 | 48.78 | 5600 | 0.0217 |
| VERT107 | 1177 | Europe | Ukraine | 2837.9 | 25.85 | 48.78 | 5600 | 0.0246 |
| HMF32_noUDG | 123 | Asia | China | 7980.8 | 121.99 | 44.04 | 5594 | 0.0202 |
| VERT015 | 1177 | Europe | Ukraine | 2837.9 | 25.85 | 48.78 | 5589 | 0.0203 |
| PN13.SG | 1047 | Europe | Ireland | 4982.1 | -9.14 | 53.05 | 5572 | 0.0192 |
| I7134 | 619 | Europe | Romania | 2914.2 | 22.40 | 47.75 | 5569 | 0.0185 |
| irk040.SG | 278 | Asia | Russia | 6449.2 | 103.39 | 53.22 | 5559 | 0.0217 |

| | | | | | | | | |
|---|---|---|---|---|---|---|---|---|
| PN112.SG | 1047 | Europe | Ireland | 4982.1 | -9.14 | 53.05 | 5558 | 0.0206 |
| I7192 | 657 | Europe | Czech Republic | 3495.2 | 14.22 | 50.15 | 5552 | 0.0209 |
| I15823 | 609 | Europe | Czech Republic | 3495.2 | 14.36 | 50.05 | 5550 | 0.0174 |
| I14176 | 657 | Europe | Czech Republic | 3495.2 | 14.22 | 50.15 | 5550 | 0.0280 |
| I6677 | 916 | Europe | Czech Republic | 3537.1 | 13.78 | 50.55 | 5550 | 0.0188 |
| PN02.SG | 1047 | Europe | Ireland | 4982.1 | -9.14 | 53.05 | 5546 | 0.0198 |
| PB357.SG | 1035 | Europe | Ireland | 4968.5 | -9.11 | 52.99 | 5545 | 0.0211 |
| prs016_noUDG.SG | 1055 | Europe | Ireland | 4945.5 | -8.56 | 54.25 | 5539 | 0.0197 |
| CBT016 | 500 | Europe | Turkey | 1468.5 | 34.59 | 40.02 | 5538 | 0.0218 |
| I3068 | 631 | Europe | United Kingdom | 4557.3 | -1.64 | 53.08 | 5533 | 0.0190 |
| I6747_noUDG.SG | 631 | Europe | United Kingdom | 4557.3 | -1.64 | 53.08 | 5528 | 0.0172 |
| I7194 | 657 | Europe | Czech Republic | 3495.2 | 14.22 | 50.15 | 5527 | 0.0239 |
| I1593 | 684 | Europe | Germany | 3943.7 | 7.55 | 51.36 | 5527 | 0.0210 |
| I6759 | 921 | Europe | United Kingdom | 4620.3 | -2.29 | 54.07 | 5527 | 0.0247 |
| I4910 | 240 | Asia | Tajikistan | 3611.2 | 67.46 | 39.51 | 5526 | 0.0213 |
| CBT004 | 500 | Europe | Turkey | 1468.5 | 34.59 | 40.02 | 5526 | 0.0156 |
| CBT013 | 500 | Europe | Turkey | 1468.5 | 34.59 | 40.02 | 5526 | 0.0135 |
| I7189 | 657 | Europe | Czech Republic | 3495.2 | 14.22 | 50.15 | 5526 | 0.0207 |
| I6746_noUDG.SG | 835 | Europe | United Kingdom | 4509.5 | -2.75 | 51.32 | 5526 | 0.0217 |
| irk034_noUDG.SG | 278 | Asia | Russia | 6449.2 | 103.39 | 53.22 | 5525 | 0.0246 |
| CBT015 | 500 | Europe | Turkey | 1468.5 | 34.59 | 40.02 | 5525 | 0.0136 |
| I6761 | 923 | Europe | United Kingdom | 4296.4 | -0.11 | 50.83 | 5525 | 0.0198 |
| I2635 | 814 | Europe | United Kingdom | 5167.4 | -3.60 | 58.53 | 5523 | 0.0225 |
| I2791 | 829 | Europe | Hungary | 2913.5 | 20.00 | 47.19 | 5523 | 0.0263 |
| BRZ002 | 492 | Europe | Czech Republic | 3537.1 | 13.74 | 50.36 | 5521 | 0.0187 |
| CBT014 | 500 | Europe | Turkey | 1468.5 | 34.59 | 40.02 | 5518 | 0.0181 |
| CH448.SG | 503 | Europe | Ireland | 4911.4 | -7.03 | 54.05 | 5517 | 0.0179 |
| I3135 | 837 | Europe | United Kingdom | 4959.2 | -5.48 | 56.40 | 5517 | 0.0235 |
| I5366 | 887 | Europe | United Kingdom | 4296.4 | -0.40 | 50.86 | 5516 | 0.0243 |
| ASH1.SG | 471 | Europe | Ireland | 4900.0 | -8.19 | 52.93 | 5501 | 0.0246 |
| I6757 | 920 | Europe | United Kingdom | 4620.3 | -2.27 | 54.08 | 5493 | 0.0250 |

| | | | | | | | | |
|---|---|---|---|---|---|---|---|---|
| N19_noUDG.SG | 1011 | Europe | Poland | 3539.2 | 18.96 | 52.62 | 5490 | 0.0194 |
| PB1794.SG | 1035 | Europe | Ireland | 4968.5 | -9.11 | 52.99 | 5486 | 0.0251 |
| CBT010 | 500 | Europe | Turkey | 1468.5 | 34.59 | 40.02 | 5480 | 0.0138 |
| CBT011 | 500 | Europe | Turkey | 1468.5 | 34.59 | 40.02 | 5480 | 0.0194 |
| CBT017 | 500 | Europe | Turkey | 1468.5 | 34.59 | 40.02 | 5480 | 0.0189 |
| PB768.SG | 1035 | Europe | Ireland | 4968.5 | -9.11 | 52.99 | 5475 | 0.0229 |
| PB443.SG | 1035 | Europe | Ireland | 4968.5 | -9.11 | 52.99 | 5474 | 0.0251 |
| BG72.SG | 482 | Europe | Ireland | 4831.6 | -6.82 | 52.81 | 5467 | 0.0206 |
| N20_noUDG.SG | 1011 | Europe | Poland | 3539.2 | 18.96 | 52.62 | 5467 | 0.0227 |
| I10542 | 594 | Europe | Turkey | 1775.5 | 29.30 | 40.48 | 5461 | 0.0170 |
| I3134 | 837 | Europe | United Kingdom | 4959.2 | -5.48 | 56.40 | 5461 | 0.0127 |
| PN03.SG | 1047 | Europe | Ireland | 4982.1 | -9.14 | 53.05 | 5460 | 0.0148 |
| N18_noUDG.SG | 1011 | Europe | Poland | 3539.2 | 18.96 | 52.62 | 5459 | 0.0200 |
| PN06.SG | 1047 | Europe | Ireland | 4982.1 | -9.14 | 53.05 | 5459 | 0.0193 |
| DA345_noUDG.SG | 91 | Asia | Russia | 6449.2 | 103.37 | 53.19 | 5454 | 0.0257 |
| AY2003 | 15 | Asia | Russia | 2447.9 | 43.26 | 45.69 | 5452 | 0.0170 |
| I4085 | 225 | Asia | Turkmenistan | 2702.2 | 58.23 | 37.87 | 5450 | 0.0200 |
| I4086 | 225 | Asia | Turkmenistan | 2702.2 | 58.23 | 37.87 | 5450 | 0.0101 |
| I4087 | 225 | Asia | Turkmenistan | 2702.2 | 58.23 | 37.87 | 5450 | 0.0178 |
| I4635 | 230 | Asia | Turkmenistan | 2630.8 | 56.25 | 38.35 | 5450 | 0.0119 |
| DolmenAnsiao96B_noUDG.SG | 522 | Europe | Portugal | 4910.4 | -8.43 | 39.92 | 5450 | 0.0190 |
| I1563 | 684 | Europe | Germany | 3943.7 | 7.55 | 51.36 | 5448 | 0.0207 |
| CBT001 | 500 | Europe | Turkey | 1468.5 | 34.59 | 40.02 | 5446 | 0.0168 |
| I3133 | 837 | Europe | United Kingdom | 4959.2 | -5.48 | 56.40 | 5446 | 0.0205 |
| PB672.SG | 1035 | Europe | Ireland | 4968.5 | -9.11 | 52.99 | 5442 | 0.0239 |
| PN16.SG | 1047 | Europe | Ireland | 4982.1 | -9.14 | 53.05 | 5442 | 0.0248 |
| PB2031.SG | 1035 | Europe | Ireland | 4968.5 | -9.11 | 52.99 | 5431 | 0.0206 |
| I5079 | 881 | Europe | Croatia | 2930.8 | 17.64 | 45.49 | 5421 | 0.0266 |
| TRPTN_611_noUDG.SG | 1164 | Europe | United Kingdom | 4389.8 | 0.10 | 52.17 | 5418 | 0.0249 |
| ARM001 | 9 | Asia | Armenia | 1927.6 | 43.72 | 40.87 | 5417 | 0.0180 |
| I7272 | 933 | Europe | Czech Republic | 3495.2 | 14.16 | 50.19 | 5417 | 0.0211 |

| | | | | | | | | |
|---|---|---|---|---|---|---|---|---|
| CBT005 | 500 | Europe | Turkey | 1468.5 | 34.59 | 40.02 | 5415 | 0.0201 |
| ART022 | 470 | Europe | Turkey | 1427.6 | 38.36 | 38.38 | 5413 | 0.0210 |
| PB581.SG | 1035 | Europe | Ireland | 4968.5 | -9.11 | 52.99 | 5413 | 0.0147 |
| PN07.SG | 1047 | Europe | Ireland | 4982.1 | -9.14 | 53.05 | 5410 | 0.0252 |
| PB1327.SG | 1035 | Europe | Ireland | 4968.5 | -9.11 | 52.99 | 5407 | 0.0267 |
| ANN1.SG | 465 | Europe | Ireland | 4900.0 | -8.46 | 52.68 | 5405 | 0.0226 |
| prs013_noUDG.SG | 1055 | Europe | Ireland | 4945.5 | -8.56 | 54.25 | 5403 | 0.0273 |
| ARD2.SG | 468 | Europe | Ireland | 4900.0 | -8.15 | 52.93 | 5401 | 0.0252 |
| I7950 | 558 | Europe | Czech Republic | 3495.2 | 14.28 | 50.08 | 5398 | 0.0204 |
| prs009_noUDG.SG | 1055 | Europe | Ireland | 4945.5 | -8.56 | 54.25 | 5398 | 0.0264 |
| AY2001 | 15 | Asia | Russia | 2447.9 | 43.26 | 45.69 | 5397 | 0.0250 |
| irk075_noUDG.SG | 283 | Asia | Russia | 6652.4 | 106.76 | 53.02 | 5397 | 0.0187 |
| SA6001 | 373 | Asia | Russia | 2447.9 | 43.99 | 45.73 | 5397 | 0.0184 |
| I15648 | 685 | Europe | Czech Republic | 3495.2 | 14.16 | 49.99 | 5397 | 0.0140 |
| I2605 | 811 | Europe | United Kingdom | 4382.1 | -0.67 | 51.49 | 5397 | 0.0166 |
| I3136 | 837 | Europe | United Kingdom | 4959.2 | -5.48 | 56.40 | 5397 | 0.0151 |
| BOT2016_noUDG.SG | 30 | Asia | Kazakhstan | 4280.2 | 67.65 | 53.31 | 5396 | 0.0202 |
| PSS4693.SG | 1058 | Europe | France | 3900.5 | 3.60 | 48.52 | 5396 | 0.0261 |
| R24.SG | 1093 | Europe | Italy | 3570.8 | 8.44 | 40.81 | 5396 | 0.0251 |
| I4110 | 857 | Europe | Ukraine | 2963.6 | 33.76 | 48.91 | 5394 | 0.0103 |
| PB186.SG | 1035 | Europe | Ireland | 4968.5 | -9.11 | 52.99 | 5393 | 0.0247 |
| GB_noUDG.SG | 540 | Europe | Romania | 2776.7 | 23.52 | 46.79 | 5389 | 0.0238 |
| I16092 | 690 | Europe | Czech Republic | 3537.1 | 13.79 | 50.38 | 5385 | 0.0197 |
| I6266 | 908 | Europe | Russia | 2256.7 | 40.39 | 44.39 | 5385 | 0.0188 |
| ART024 | 470 | Europe | Turkey | 1427.6 | 38.36 | 38.38 | 5384 | 0.0214 |
| PN12.SG | 1047 | Europe | Ireland | 4982.1 | -9.14 | 53.05 | 5383 | 0.0271 |
| BOT14_noUDG.SG | 30 | Asia | Kazakhstan | 4280.2 | 67.65 | 53.31 | 5337 | 0.0206 |
| I6753_noUDG.SG | 919 | Europe | United Kingdom | 4311.6 | 0.37 | 51.32 | 5305 | 0.0187 |
| GNM1007.SG | 542 | Europe | Ireland | 5011.8 | -9.63 | 53.97 | 5288 | 0.0246 |
| ART018 | 470 | Europe | Turkey | 1427.6 | 38.36 | 38.38 | 5260 | 0.0224 |
| ART014 | 470 | Europe | Turkey | 1427.6 | 38.36 | 38.38 | 5255 | 0.0160 |

| | | | | | | | | |
|---|---|---|---|---|---|---|---|---|
| ART032 | 470 | Europe | Turkey | 1427.6 | 38.36 | 38.38 | 5254 | 0.0180 |
| BLSM27S_noUDG | 29 | Asia | China | 8011.1 | 120.27 | 41.41 | 5250 | 0.0185 |
| ART015 | 470 | Europe | Turkey | 1427.6 | 38.36 | 38.38 | 5212 | 0.0164 |
| TGM009 | 1159 | Europe | Germany | 3791.8 | 11.97 | 52.54 | 5208 | 0.0188 |
| bal004_noUDG.SG | 476 | Europe | United Kingdom | 5070.7 | -3.91 | 57.76 | 5186 | 0.0286 |
| I2369 | 778 | Europe | Hungary | 3004.9 | 19.04 | 47.62 | 5186 | 0.0158 |
| I7554 | 937 | Europe | United Kingdom | 5189.1 | -3.25 | 58.99 | 5182 | 0.0169 |
| hem005.SG | 552 | Europe | Sweden | 4252.3 | 18.57 | 57.22 | 5178 | 0.0226 |
| I4483 | 236 | Asia | Turkey | 1496.4 | 41.87 | 37.54 | 5176 | 0.0111 |
| I2752 | 826 | Europe | Hungary | 3007.4 | 17.73 | 46.78 | 5175 | 0.0244 |
| I2785 | 828 | Europe | Hungary | 3003.9 | 19.92 | 47.68 | 5175 | 0.0194 |
| PB675.SG | 1035 | Europe | Ireland | 4968.5 | -9.11 | 52.99 | 5171 | 0.0229 |
| I2371 | 779 | Europe | Hungary | 3004.9 | 19.17 | 47.32 | 5169 | 0.0209 |
| ART017 | 470 | Europe | Turkey | 1427.6 | 38.36 | 38.38 | 5166 | 0.0135 |
| ROUQFF.SG | 1132 | Europe | France | 3799.7 | 3.92 | 43.59 | 5166 | 0.0215 |
| 2H10.SG | 438 | Europe | France | 3914.4 | 4.00 | 48.86 | 5165 | 0.0218 |
| I11501 | 156 | Asia | Kazakhstan | 3543.2 | 58.51 | 49.49 | 5164 | 0.0232 |
| SIJ002 | 1133 | Europe | Russia | 2399.5 | 39.91 | 45.05 | 5164 | 0.0209 |
| SIJ003 | 1133 | Europe | Russia | 2399.5 | 39.91 | 45.05 | 5164 | 0.0180 |
| ARM002 | 9 | Asia | Armenia | 1927.6 | 43.72 | 40.87 | 5163 | 0.0203 |
| ART026 | 470 | Europe | Turkey | 1427.6 | 38.36 | 38.38 | 5163 | 0.0235 |
| ROUQHH.SG | 1132 | Europe | France | 3799.7 | 3.92 | 43.59 | 5163 | 0.0270 |
| BLSM45_noUDG | 29 | Asia | China | 8011.1 | 120.27 | 41.41 | 5162 | 0.0266 |
| I1658 | 204 | Asia | Armenia | 1882.1 | 43.87 | 40.38 | 5162 | 0.0131 |
| 2H11.SG | 438 | Europe | France | 3914.4 | 4.00 | 48.86 | 5162 | 0.0238 |
| SA6004 | 373 | Asia | Russia | 2447.9 | 43.99 | 45.73 | 5159 | 0.0223 |
| ART012 | 470 | Europe | Turkey | 1427.6 | 38.36 | 38.38 | 5159 | 0.0252 |
| ROUQV.SG | 1132 | Europe | France | 3799.7 | 3.92 | 43.59 | 5159 | 0.0148 |
| 2H17.SG | 438 | Europe | France | 3914.4 | 4.00 | 48.86 | 5157 | 0.0194 |
| I0172 | 564 | Europe | Germany | 3710.4 | 11.68 | 51.42 | 5155 | 0.0257 |
| ans008_noUDG.SG | 466 | Europe | Sweden | 4300.0 | 18.14 | 57.51 | 5154 | 0.0256 |

| | | | | | | | | |
|---|---|---|---|---|---|---|---|---|
| HOL004 | 557 | Europe | Czech Republic | 3495.2 | 14.29 | 50.20 | 5154 | 0.0204 |
| I2520 | 602 | Europe | Bulgaria | 2209.4 | 25.88 | 43.16 | 5153 | 0.0246 |
| NG10.SG | 1018 | Europe | Ireland | 4828.4 | -6.48 | 53.69 | 5151 | 0.0235 |
| MK5001 | 324 | Asia | Russia | 2254.5 | 43.52 | 43.91 | 5150 | 0.0177 |
| MK5004 | 324 | Asia | Russia | 2254.5 | 43.52 | 43.91 | 5150 | 0.0195 |
| I3085 | 813 | Europe | United Kingdom | 5139.6 | -2.92 | 58.74 | 5149 | 0.0271 |
| I5076 | 880 | Europe | Portugal | 5083.7 | -8.59 | 37.20 | 5146 | 0.0195 |
| hem004.SG | 552 | Europe | Sweden | 4252.3 | 18.57 | 57.22 | 5145 | 0.0198 |
| BOT15_noUDG.SG | 30 | Asia | Kazakhstan | 4280.2 | 67.65 | 53.31 | 5143 | 0.0221 |
| HOL002 | 557 | Europe | Czech Republic | 3495.2 | 14.29 | 50.20 | 5140 | 0.0260 |
| I2978 | 813 | Europe | United Kingdom | 5139.6 | -2.92 | 58.74 | 5140 | 0.0176 |
| I2934 | 813 | Europe | United Kingdom | 5139.6 | -2.92 | 58.74 | 5138 | 0.0172 |
| GNM1076.SG | 542 | Europe | Ireland | 5011.8 | -9.63 | 53.97 | 5135 | 0.0197 |
| irk017_noUDG.SG | 274 | Asia | Russia | 6589.0 | 105.79 | 54.00 | 5132 | 0.0241 |
| irk071_noUDG.SG | 282 | Asia | Russia | 6720.1 | 106.96 | 53.15 | 5132 | 0.0227 |
| bally_noUDG.SG | 477 | Europe | Ireland | 4869.8 | -5.96 | 54.54 | 5131 | 0.0240 |
| I1838 | 730 | Europe | Spain | 4402.7 | -2.70 | 42.63 | 5126 | 0.0229 |
| SA6002 | 1133 | Europe | Russia | 2399.5 | 39.91 | 45.05 | 5126 | 0.0230 |
| I1657 | 203 | Asia | Armenia | 1882.1 | 43.89 | 40.39 | 5123 | 0.0198 |
| I2370 | 779 | Europe | Hungary | 3004.9 | 19.17 | 47.32 | 5122 | 0.0232 |
| I2366 | 778 | Europe | Hungary | 3004.9 | 19.04 | 47.62 | 5118 | 0.0165 |
| 1H06.SG | 438 | Europe | France | 3914.4 | 4.00 | 48.86 | 5115 | 0.0309 |
| I2935 | 813 | Europe | United Kingdom | 5139.6 | -2.92 | 58.74 | 5115 | 0.0204 |
| ans014_noUDG.SG | 466 | Europe | Sweden | 4300.0 | 18.14 | 57.51 | 5111 | 0.0286 |
| por002_noUDG.SG | 474 | Europe | Spain | 4402.7 | -3.52 | 42.35 | 5111 | 0.0241 |
| I2979 | 813 | Europe | United Kingdom | 5139.6 | -2.92 | 58.74 | 5104 | 0.0204 |
| I5118 | 883 | Europe | Hungary | 2958.8 | 20.89 | 47.81 | 5100 | 0.0182 |
| I6766 | 925 | Europe | United Kingdom | 5119.1 | -4.00 | 57.91 | 5100 | 0.0191 |
| I11028 | 153 | Asia | Uzbekistan | 3507.3 | 67.00 | 37.67 | 5099 | 0.0148 |
| I5119 | 883 | Europe | Hungary | 2958.8 | 20.89 | 47.81 | 5097 | 0.0222 |
| I2367 | 778 | Europe | Hungary | 3004.9 | 19.04 | 47.62 | 5085 | 0.0223 |

| | | | | | | | | |
|---|---|---|---|---|---|---|---|---|
| I2753 | 826 | Europe | Hungary | 3007.4 | 17.73 | 46.78 | 5085 | 0.0220 |
| I0443 | 131 | Asia | Russia | 3576.4 | 50.66 | 53.64 | 5084 | 0.0229 |
| I0429 | 131 | Asia | Russia | 3576.4 | 50.66 | 53.64 | 5074 | 0.0173 |
| I2606 | 811 | Europe | United Kingdom | 4382.1 | -0.67 | 51.49 | 5065 | 0.0227 |
| I2069 | 213 | Asia | Russia | 5163.3 | 84.81 | 50.91 | 5061 | 0.0229 |
| ans017_noUDG.SG | 466 | Europe | Sweden | 4300.0 | 18.14 | 57.51 | 5053 | 0.0222 |
| I12485 | 166 | Asia | Turkmenistan | 2964.6 | 61.03 | 37.19 | 5050 | 0.0191 |
| I12486 | 166 | Asia | Turkmenistan | 2964.6 | 61.03 | 37.19 | 5050 | 0.0184 |
| CabecoArruda122A_noUDG.SG | 493 | Europe | Portugal | 5028.8 | -8.66 | 39.11 | 5050 | 0.0207 |
| I11248 | 610 | Europe | Spain | 4306.1 | -2.22 | 43.09 | 5050 | 0.0178 |
| I11249 | 610 | Europe | Spain | 4306.1 | -2.22 | 43.09 | 5050 | 0.0185 |
| I11301 | 610 | Europe | Spain | 4306.1 | -2.22 | 43.09 | 5050 | 0.0229 |
| I2441 | 789 | Europe | Poland | 3579.4 | 17.88 | 52.85 | 5050 | 0.0251 |
| I7602 | 942 | Europe | Spain | 4306.1 | -2.25 | 43.09 | 5050 | 0.0190 |
| I7603 | 942 | Europe | Spain | 4306.1 | -2.25 | 43.09 | 5050 | 0.0209 |
| I7604 | 942 | Europe | Spain | 4306.1 | -2.25 | 43.09 | 5050 | 0.0231 |
| I7606 | 942 | Europe | Spain | 4306.1 | -2.25 | 43.09 | 5050 | 0.0256 |
| I7489 | 134 | Asia | Russia | 3487.5 | 50.99 | 52.91 | 5047 | 0.0190 |
| ROUQEE.SG | 1132 | Europe | France | 3799.7 | 3.92 | 43.59 | 5047 | 0.0258 |
| NGZ1.SG | 1019 | Europe | Ireland | 4828.4 | -6.48 | 53.70 | 5024 | 0.0209 |
| I11736 | 161 | Asia | Kazakhstan | 3222.8 | 49.34 | 50.77 | 5021 | 0.0238 |
| I4259 | 230 | Asia | Turkmenistan | 2630.8 | 56.25 | 38.35 | 5021 | 0.0204 |
| I1497 | 675 | Europe | Hungary | 2913.5 | 19.83 | 47.17 | 5019 | 0.0186 |
| SHT001 | 382 | Asia | Mongolia | 6258.0 | 100.76 | 46.31 | 5009 | 0.0195 |
| I0444 | 141 | Asia | Russia | 3571.7 | 51.22 | 53.18 | 5003 | 0.0246 |
| I8726 | 155 | Asia | Iran | 3027.5 | 61.40 | 30.65 | 5000 | 0.0230 |
| I4634 | 230 | Asia | Turkmenistan | 2630.8 | 56.25 | 38.35 | 4998 | 0.0176 |
| I3138 | 837 | Europe | United Kingdom | 4959.2 | -5.48 | 56.40 | 4998 | 0.0229 |
| I6221 | 244 | Asia | Mongolia | 6252.4 | 100.82 | 46.40 | 4996 | 0.0262 |
| SHT002 | 382 | Asia | Mongolia | 6258.0 | 100.76 | 46.31 | 4996 | 0.0244 |
| I4615 | 236 | Asia | Turkey | 1496.4 | 41.87 | 37.54 | 4991 | 0.0227 |

| | | | | | | | | |
|---|---|---|---|---|---|---|---|---|
| atp016_noUDG.SG | 474 | Europe | Spain | 4402.7 | -3.52 | 42.35 | 4971 | 0.0255 |
| Aes4 | 453 | Europe | Switzerland | 3569.2 | 7.60 | 47.47 | 4970 | 0.0193 |
| I2180 | 772 | Europe | Bulgaria | 2161.8 | 27.02 | 43.07 | 4959 | 0.0240 |
| N38_noUDG.SG | 1012 | Europe | Poland | 3539.2 | 18.90 | 52.61 | 4959 | 0.0164 |
| I3276 | 733 | Europe | Spain | 4402.7 | -2.57 | 42.57 | 4955 | 0.0090 |
| I1917 | 751 | Europe | Ukraine | 2998.4 | 33.95 | 48.99 | 4955 | 0.0197 |
| I5277 | 241 | Asia | Russia | 5152.2 | 85.56 | 50.76 | 4952 | 0.0183 |
| 1H13.SG | 438 | Europe | France | 3914.4 | 4.00 | 48.86 | 4950 | 0.0160 |
| I2433 | 789 | Europe | Poland | 3579.4 | 17.88 | 52.85 | 4950 | 0.0225 |
| MX210 | 1008 | Europe | Switzerland | 3569.2 | 7.66 | 47.26 | 4950 | 0.0213 |
| TU910_SX22 | 1167 | Europe | Switzerland | 3514.3 | 9.29 | 47.45 | 4949 | 0.0160 |
| I0357 | 131 | Asia | Russia | 3576.4 | 50.66 | 53.64 | 4942 | 0.0212 |
| I5742 | 906 | Europe | Turkey | 1516.5 | 35.56 | 40.69 | 4942 | 0.0128 |
| I2071 | 214 | Asia | Russia | 5245.6 | 85.97 | 51.50 | 4940 | 0.0213 |
| TUC001 | 1169 | Europe | Czech Republic | 3495.2 | 14.27 | 50.14 | 4940 | 0.0190 |
| TUC003 | 1169 | Europe | Czech Republic | 3495.2 | 14.27 | 50.14 | 4940 | 0.0158 |
| TUC004 | 1169 | Europe | Czech Republic | 3495.2 | 14.27 | 50.14 | 4940 | 0.0249 |
| TUC005 | 1169 | Europe | Czech Republic | 3495.2 | 14.27 | 50.14 | 4940 | 0.0164 |
| I11531 | 156 | Asia | Kazakhstan | 3543.2 | 58.51 | 49.49 | 4937 | 0.0214 |
| vbj012_noUDG.SG | 1173 | Europe | Sweden | 4300.0 | 18.70 | 57.58 | 4933 | 0.0256 |
| TUC002 | 1169 | Europe | Czech Republic | 3495.2 | 14.27 | 50.14 | 4932 | 0.0200 |
| I16425 | 693 | Europe | Channel Islands | 4383.6 | -2.51 | 49.50 | 4929 | 0.0223 |
| Gokhem2_noUDG.SG | 543 | Europe | Sweden | 4318.3 | 13.41 | 58.17 | 4928 | 0.0239 |
| CAK533.SG | 495 | Europe | Ireland | 4945.5 | -8.38 | 54.06 | 4926 | 0.0217 |
| I6669 | 230 | Asia | Turkmenistan | 2630.8 | 56.25 | 38.35 | 4924 | 0.0170 |
| I3432 | 843 | Europe | Portugal | 4883.6 | -7.55 | 38.39 | 4924 | 0.0268 |
| I5743 | 906 | Europe | Turkey | 1516.5 | 35.56 | 40.69 | 4924 | 0.0187 |
| I11732 | 156 | Asia | Kazakhstan | 3543.2 | 58.51 | 49.49 | 4921 | 0.0209 |
| Aes3 | 453 | Europe | Switzerland | 3569.2 | 7.60 | 47.47 | 4919 | 0.0153 |
| I11734 | 156 | Asia | Kazakhstan | 3543.2 | 58.51 | 49.49 | 4918 | 0.0231 |
| I4565 | 867 | Europe | Spain | 4188.1 | 1.06 | 41.29 | 4915 | 0.0248 |

| | | | | | | | | |
|---|---|---|---|---|---|---|---|---|
| I10494 | 596 | Europe | Romania | 2402.8 | 26.01 | 44.90 | 4914 | 0.0218 |
| I2165 | 771 | Europe | Bulgaria | 2168.7 | 25.50 | 42.13 | 4910 | 0.0226 |
| I14326 | 658 | Europe | United Kingdom | 4552.7 | -0.28 | 54.11 | 4908 | 0.0197 |
| VLI076 | 1253 | Europe | Czech Republic | 3537.1 | 14.45 | 50.37 | 4906 | 0.0189 |
| I1843 | 733 | Europe | Spain | 4402.7 | -2.57 | 42.57 | 4904 | 0.0221 |
| RISE1166_noUDG.SG | 1121 | Europe | Poland | 3229.3 | 20.58 | 50.17 | 4904 | 0.0209 |
| I5269 | 241 | Asia | Russia | 5152.2 | 85.56 | 50.76 | 4902 | 0.0176 |
| Aes13 | 453 | Europe | Switzerland | 3569.2 | 7.60 | 47.47 | 4902 | 0.0172 |
| I10287 | 585 | Europe | Spain | 4140.4 | 1.57 | 41.44 | 4900 | 0.0227 |
| vbj006_noUDG.SG | 1173 | Europe | Sweden | 4300.0 | 18.70 | 57.58 | 4898 | 0.0279 |
| CAK532.SG | 495 | Europe | Ireland | 4945.5 | -8.38 | 54.06 | 4892 | 0.0187 |
| I5279 | 242 | Asia | Russia | 5152.2 | 85.56 | 50.75 | 4891 | 0.0234 |
| Damgaard2018Yamnaya_noUDG.SG | 103 | Asia | Kazakhstan | 4459.0 | 75.85 | 49.13 | 4890 | 0.0220 |
| Aes11 | 453 | Europe | Switzerland | 3569.2 | 7.60 | 47.47 | 4885 | 0.0221 |
| TUC006 | 1169 | Europe | Czech Republic | 3495.2 | 14.27 | 50.14 | 4879 | 0.0197 |
| Aes17 | 453 | Europe | Switzerland | 3569.2 | 7.60 | 47.47 | 4878 | 0.0240 |
| I2933 | 813 | Europe | United Kingdom | 5139.6 | -2.92 | 58.74 | 4877 | 0.0215 |
| I5273 | 242 | Asia | Russia | 5152.2 | 85.56 | 50.75 | 4872 | 0.0251 |
| atp12-1420_noUDG.SG | 474 | Europe | Spain | 4402.7 | -3.52 | 42.35 | 4868 | 0.0207 |
| vbj018_noUDG.SG | 1173 | Europe | Sweden | 4300.0 | 18.70 | 57.58 | 4868 | 0.0252 |
| I5429 | 843 | Europe | Portugal | 4883.6 | -7.55 | 38.39 | 4866 | 0.0186 |
| I8725 | 155 | Asia | Iran | 3027.5 | 61.40 | 30.65 | 4862 | 0.0138 |
| I13180 | 183 | Asia | Mongolia | 5419.2 | 88.83 | 49.30 | 4862 | 0.0269 |
| I5272 | 241 | Asia | Russia | 5152.2 | 85.56 | 50.76 | 4861 | 0.0219 |
| RA62 | 1119 | Europe | Switzerland | 3645.6 | 7.38 | 47.31 | 4853 | 0.0202 |
| I0370 | 132 | Asia | Russia | 3658.7 | 58.18 | 51.27 | 4850 | 0.0219 |
| I11112 | 154 | Asia | Russia | 5648.2 | 91.47 | 53.87 | 4850 | 0.0179 |
| I6715 | 219 | Asia | Russia | 5041.2 | 83.91 | 48.83 | 4850 | 0.0178 |
| RISE548_noUDG.SG | 363 | Asia | Russia | 2544.6 | 43.70 | 46.54 | 4850 | 0.0214 |
| ZO2002 | 421 | Asia | Russia | 2403.3 | 42.61 | 45.66 | 4850 | 0.0144 |
| I3269 | 730 | Europe | Spain | 4402.7 | -2.70 | 42.63 | 4850 | 0.0255 |

| | | | | | | | | |
|---|---|---|---|---|---|---|---|---|
| I2105 | 768 | Europe | Ukraine | 2822.2 | 37.15 | 48.22 | 4850 | 0.0217 |
| LU339_noUDG.SG | 987 | Europe | Portugal | 4761.4 | -6.93 | 41.71 | 4849 | 0.0196 |
| I0231 | 126 | Asia | Russia | 3492.8 | 49.47 | 53.09 | 4844 | 0.0162 |
| I11462 | 155 | Asia | Iran | 3027.5 | 61.40 | 30.65 | 4844 | 0.0162 |
| I2977 | 813 | Europe | United Kingdom | 5139.6 | -2.92 | 58.74 | 4843 | 0.0248 |
| Aes20 | 453 | Europe | Switzerland | 3569.2 | 7.60 | 47.47 | 4842 | 0.0234 |
| Plinkaigalis242 | 1043 | Europe | Lithuania | 3663.4 | 23.65 | 55.41 | 4842 | 0.0244 |
| R4.SG | 1062 | Europe | Italy | 3014.0 | 13.54 | 41.96 | 4841 | 0.0263 |
| Aes24 | 453 | Europe | Switzerland | 3569.2 | 7.60 | 47.47 | 4839 | 0.0219 |
| Aes18 | 453 | Europe | Switzerland | 3569.2 | 7.60 | 47.47 | 4838 | 0.0252 |
| R5.SG | 1062 | Europe | Italy | 3014.0 | 13.54 | 41.96 | 4832 | 0.0227 |
| ajv28_noUDG.SG | 456 | Europe | Sweden | 4252.3 | 18.20 | 57.28 | 4829 | 0.0219 |
| RISE1171_noUDG.SG | 1121 | Europe | Poland | 3229.3 | 20.58 | 50.17 | 4826 | 0.0235 |
| Aes19 | 453 | Europe | Switzerland | 3569.2 | 7.60 | 47.47 | 4825 | 0.0201 |
| Aes21 | 453 | Europe | Switzerland | 3569.2 | 7.60 | 47.47 | 4825 | 0.0180 |
| I1846 | 733 | Europe | Spain | 4402.7 | -2.57 | 42.57 | 4825 | 0.0286 |
| I1978 | 733 | Europe | Spain | 4402.7 | -2.57 | 42.57 | 4825 | 0.0193 |
| I12977 | 179 | Asia | Mongolia | 5419.2 | 88.71 | 49.36 | 4822 | 0.0284 |
| NEO232_noUDG.SG | 197 | Asia | Russia | 6446.8 | 103.66 | 52.92 | 4812 | 0.0226 |
| I7642 | 944 | Europe | Spain | 4532.3 | -0.49 | 38.70 | 4809 | 0.0217 |
| RA61 | 1119 | Europe | Switzerland | 3645.6 | 7.38 | 47.31 | 4804 | 0.0253 |
| I2510 | 602 | Europe | Bulgaria | 2209.4 | 25.88 | 43.16 | 4794 | 0.0173 |
| I0438 | 139 | Asia | Russia | 3531.9 | 50.38 | 53.38 | 4785 | 0.0170 |
| RISE1161_noUDG.SG | 1121 | Europe | Poland | 3229.3 | 20.58 | 50.17 | 4785 | 0.0243 |
| ILK003 | 953 | Europe | Ukraine | 2894.8 | 27.69 | 49.56 | 4781 | 0.0208 |
| I1526 | 197 | Asia | Russia | 6446.8 | 103.66 | 52.92 | 4779 | 0.0205 |
| vbj013_noUDG.SG | 1173 | Europe | Sweden | 4300.0 | 18.70 | 57.58 | 4778 | 0.0269 |
| G218M5-2.SG | 117 | Asia | China | 4909.4 | 83.54 | 43.90 | 4775 | 0.0252 |
| ILK001 | 953 | Europe | Ukraine | 2894.8 | 27.69 | 49.56 | 4773 | 0.0197 |
| RISE1241_noUDG.SG | 1122 | Europe | Poland | 3232.5 | 21.70 | 50.60 | 4773 | 0.0274 |
| RISE1162_noUDG.SG | 1121 | Europe | Poland | 3229.3 | 20.58 | 50.17 | 4765 | 0.0195 |

| | | | | | | | | |
|---|---|---|---|---|---|---|---|---|
| VLI008 | 1253 | Europe | Czech Republic | 3537.1 | 14.45 | 50.37 | 4759 | 0.0233 |
| RISE511_noUDG.SG | 152 | Asia | Russia | 5674.0 | 90.92 | 54.36 | 4757 | 0.0233 |
| VPR001 | 1258 | Europe | Czech Republic | 3495.2 | 14.32 | 50.16 | 4756 | 0.0196 |
| RISE1165_noUDG.SG | 1121 | Europe | Poland | 3229.3 | 20.58 | 50.17 | 4754 | 0.0197 |
| RISE1170_noUDG.SG | 1121 | Europe | Poland | 3229.3 | 20.58 | 50.17 | 4754 | 0.0147 |
| I7779 | 197 | Asia | Russia | 6446.8 | 103.66 | 52.92 | 4750 | 0.0229 |
| CDP001 | 502 | Europe | Spain | 4483.0 | -0.34 | 38.78 | 4750 | 0.0263 |
| CLL004 | 507 | Europe | Spain | 4581.0 | -0.86 | 38.63 | 4750 | 0.0236 |
| CLL006 | 507 | Europe | Spain | 4581.0 | -0.86 | 38.63 | 4750 | 0.0243 |
| I11605 | 613 | Europe | Portugal | 5010.4 | -9.30 | 39.23 | 4750 | 0.0223 |
| KO1016 | 972 | Europe | Czech Republic | 3449.6 | 15.17 | 50.03 | 4750 | 0.0150 |
| PLZ001 | 1044 | Europe | Spain | 4581.0 | -0.87 | 38.64 | 4750 | 0.0241 |
| RISE1169_noUDG.SG | 1121 | Europe | Poland | 3229.3 | 20.58 | 50.17 | 4747 | 0.0173 |
| atp002_noUDG.SG | 474 | Europe | Spain | 4402.7 | -3.52 | 42.35 | 4746 | 0.0206 |
| VLI007 | 1253 | Europe | Czech Republic | 3537.1 | 14.45 | 50.37 | 4739 | 0.0244 |
| RISE1249_noUDG.SG | 1123 | Europe | Poland | 3186.9 | 21.40 | 50.20 | 4737 | 0.0216 |
| RISE1163_noUDG.SG | 1121 | Europe | Poland | 3229.3 | 20.58 | 50.17 | 4736 | 0.0244 |
| I5884 | 857 | Europe | Ukraine | 2963.6 | 33.76 | 48.91 | 4735 | 0.0264 |
| ILK002 | 953 | Europe | Ukraine | 2894.8 | 27.69 | 49.56 | 4734 | 0.0293 |
| I7671 | 269 | Asia | Russia | 3531.9 | 50.67 | 53.17 | 4729 | 0.0176 |
| I5733 | 905 | Europe | Turkey | 1699.7 | 27.14 | 38.42 | 4729 | 0.0140 |
| I10565 | 152 | Asia | Russia | 5674.0 | 90.92 | 54.36 | 4726 | 0.0175 |
| I3387 | 152 | Asia | Russia | 5674.0 | 90.92 | 54.36 | 4725 | 0.0187 |
| RISE1247_noUDG.SG | 1123 | Europe | Poland | 3186.9 | 21.40 | 50.20 | 4725 | 0.0231 |
| RISE1248_noUDG.SG | 1123 | Europe | Poland | 3186.9 | 21.40 | 50.20 | 4725 | 0.0223 |
| RISE1159_noUDG.SG | 1121 | Europe | Poland | 3229.3 | 20.58 | 50.17 | 4724 | 0.0262 |
| Aes23 | 453 | Europe | Switzerland | 3569.2 | 7.60 | 47.47 | 4722 | 0.0254 |
| RISE1160_noUDG.SG | 1121 | Europe | Poland | 3229.3 | 20.58 | 50.17 | 4721 | 0.0286 |
| RISE1167_noUDG.SG | 1121 | Europe | Poland | 3229.3 | 20.58 | 50.17 | 4721 | 0.0209 |
| I0440 | 140 | Asia | Russia | 3531.9 | 50.39 | 53.38 | 4720 | 0.0262 |
| irk036_noUDG.SG | 279 | Asia | Russia | 6514.2 | 104.26 | 52.29 | 4720 | 0.0223 |

| | | | | | | | | |
|---|---|---|---|---|---|---|---|---|
| STD002 | 1146 | Europe | Czech Republic | 3537.1 | 13.96 | 50.61 | 4720 | 0.0225 |
| VLI092 | 1253 | Europe | Czech Republic | 3537.1 | 14.45 | 50.37 | 4719 | 0.0200 |
| GW1001 | 121 | Asia | Russia | 2254.5 | 43.13 | 44.03 | 4718 | 0.0146 |
| VLI011 | 1253 | Europe | Czech Republic | 3537.1 | 14.45 | 50.37 | 4718 | 0.0223 |
| VLI067 | 1253 | Europe | Czech Republic | 3537.1 | 14.45 | 50.37 | 4718 | 0.0202 |
| RK1001 | 370 | Asia | Russia | 2359.3 | 41.12 | 45.54 | 4717 | 0.0158 |
| I2924 | 217 | Asia | Iran | 2407.9 | 54.38 | 36.15 | 4715 | 0.0217 |
| I2925 | 217 | Asia | Iran | 2407.9 | 54.38 | 36.15 | 4715 | 0.0213 |
| VLI071 | 1253 | Europe | Czech Republic | 3537.1 | 14.45 | 50.37 | 4715 | 0.0255 |
| DA358_noUDG.SG | 93 | Asia | Russia | 6720.1 | 106.96 | 53.18 | 4713 | 0.0185 |
| RISE547_noUDG.SG | 363 | Asia | Russia | 2544.6 | 43.70 | 46.54 | 4713 | 0.0201 |
| TOU001 | 1162 | Europe | Czech Republic | 3449.6 | 14.72 | 50.17 | 4712 | 0.0161 |
| I3951 | 219 | Asia | Russia | 5041.2 | 83.91 | 48.83 | 4711 | 0.0234 |
| I4481 | 236 | Asia | Turkey | 1496.4 | 41.87 | 37.54 | 4711 | 0.0193 |
| CAK530.SG | 495 | Europe | Ireland | 4945.5 | -8.38 | 54.06 | 4711 | 0.0185 |
| BZK002 | 43 | Asia | Russia | 5845.8 | 92.85 | 55.99 | 4709 | 0.0185 |
| I6294 | 128 | Asia | Russia | 3576.4 | 50.57 | 53.59 | 4709 | 0.0280 |
| I2923 | 217 | Asia | Iran | 2407.9 | 54.38 | 36.15 | 4707 | 0.0178 |
| I3950.SG | 219 | Asia | Russia | 5041.2 | 83.91 | 48.83 | 4707 | 0.0216 |
| poz81_noUDG.SG | 1053 | Europe | Poland | 3576.9 | 17.55 | 52.30 | 4702 | 0.0223 |
| vbj004_noUDG.SG | 1173 | Europe | Sweden | 4300.0 | 18.70 | 57.58 | 4702 | 0.0292 |
| CAK531.SG | 495 | Europe | Ireland | 4945.5 | -8.38 | 54.06 | 4701 | 0.0259 |
| I14812 | 186 | Asia | Armenia | 1927.6 | 43.95 | 40.78 | 4700 | 0.0174 |
| I4478 | 236 | Asia | Turkey | 1496.4 | 41.87 | 37.54 | 4700 | 0.0191 |
| Ajvide58_noUDG.SG | 457 | Europe | Sweden | 4252.3 | 18.21 | 57.29 | 4700 | 0.0232 |
| VOR004_noUDG.SG | 1256 | Europe | Russia | 3752.9 | 39.55 | 57.53 | 4698 | 0.0272 |
| DRO001 | 524 | Europe | Czech Republic | 3582.7 | 13.41 | 50.44 | 4697 | 0.0130 |
| I0432 | 138 | Asia | Russia | 3576.4 | 50.67 | 53.66 | 4695 | 0.0305 |
| BLS002 | 485 | Europe | Czech Republic | 3537.1 | 13.85 | 50.35 | 4692 | 0.0213 |
| I8569 | 951 | Europe | Spain | 4335.0 | 0.28 | 40.30 | 4692 | 0.0241 |
| I3954 | 219 | Asia | Russia | 5041.2 | 83.91 | 48.83 | 4690 | 0.0157 |

| | | | | | | | | |
|---|---|---|---|---|---|---|---|---|
| I0371 | 128 | Asia | Russia | 3576.4 | 50.57 | 53.59 | 4678 | 0.0189 |
| BLS001 | 485 | Europe | Czech Republic | 3537.1 | 13.85 | 50.35 | 4675 | 0.0269 |
| vbj001_noUDG.SG | 1173 | Europe | Sweden | 4300.0 | 18.70 | 57.58 | 4675 | 0.0254 |
| RISE1168_noUDG.SG | 1121 | Europe | Poland | 3229.3 | 20.58 | 50.17 | 4672 | 0.0228 |
| LYG001 | 319 | Asia | Russia | 2254.5 | 43.21 | 44.05 | 4670 | 0.0222 |
| I3952 | 219 | Asia | Russia | 5041.2 | 83.91 | 48.83 | 4667 | 0.0253 |
| I8296 | 197 | Asia | Russia | 6446.8 | 103.66 | 52.92 | 4665 | 0.0284 |
| RISE1172_noUDG.SG | 1121 | Europe | Poland | 3229.3 | 20.58 | 50.17 | 4664 | 0.0179 |
| vbj008_noUDG.SG | 1173 | Europe | Sweden | 4300.0 | 18.70 | 57.58 | 4663 | 0.0191 |
| I1281 | 630 | Europe | Spain | 4417.6 | -3.50 | 42.33 | 4653 | 0.0256 |
| PNL002 | 1048 | Europe | Czech Republic | 3404.5 | 15.81 | 50.25 | 4653 | 0.0219 |
| I11752 | 162 | Asia | Russia | 5034.1 | 83.06 | 51.31 | 4650 | 0.0212 |
| I3388.DG | 219 | Asia | Russia | 5041.2 | 83.91 | 48.83 | 4650 | 0.0265 |
| I8197 | 949 | Europe | Spain | 4878.1 | -4.24 | 37.24 | 4650 | 0.0205 |
| I8198 | 949 | Europe | Spain | 4878.1 | -4.24 | 37.24 | 4650 | 0.0281 |
| I8199 | 949 | Europe | Spain | 4878.1 | -4.24 | 37.24 | 4650 | 0.0206 |
| TRM006 | 1163 | Europe | Czech Republic | 3537.1 | 13.99 | 50.64 | 4644 | 0.0179 |
| ART011 | 470 | Europe | Turkey | 1427.6 | 38.36 | 38.38 | 4640 | 0.0186 |
| SSGM16.SG | 390 | Asia | China | 5167.8 | 85.92 | 46.82 | 4637 | 0.0278 |
| I4619 | 236 | Asia | Turkey | 1496.4 | 41.87 | 37.54 | 4635 | 0.0243 |
| Aes25 | 453 | Europe | Switzerland | 3569.2 | 7.60 | 47.47 | 4635 | 0.0218 |
| RISE1173_noUDG.SG | 1121 | Europe | Poland | 3229.3 | 20.58 | 50.17 | 4635 | 0.0199 |
| C1707 | 49 | Asia | China | 5253.4 | 86.87 | 47.70 | 4632 | 0.0238 |
| NEO298_noUDG.SG | 197 | Asia | Russia | 6446.8 | 103.66 | 52.92 | 4630 | 0.0262 |
| POP39_noUDG.SG | 1051 | Europe | Croatia | 2878.2 | 18.57 | 45.75 | 4626 | 0.0167 |
| I6712 | 152 | Asia | Russia | 5674.0 | 90.92 | 54.36 | 4625 | 0.0163 |
| I13957 | 189 | Asia | Mongolia | 5419.2 | 88.71 | 49.34 | 4625 | 0.0303 |
| ART010 | 470 | Europe | Turkey | 1427.6 | 38.36 | 38.38 | 4625 | 0.0157 |
| PRE001 | 1048 | Europe | Czech Republic | 3404.5 | 15.81 | 50.25 | 4625 | 0.0149 |
| IVA001_noUDG.SG | 958 | Europe | Russia | 3658.4 | 36.07 | 55.82 | 4624 | 0.0307 |
| VLI079 | 1253 | Europe | Czech Republic | 3537.1 | 14.45 | 50.37 | 4623 | 0.0182 |

| | | | | | | | | |
|---|---|---|---|---|---|---|---|---|
| I5740 | 906 | Europe | Turkey | 1516.5 | 35.56 | 40.69 | 4621 | 0.0095 |
| TRM003 | 1163 | Europe | Czech Republic | 3537.1 | 13.99 | 50.64 | 4620 | 0.0194 |
| LSC002.SG | 992 | Europe | Italy | 2990.8 | 13.24 | 41.43 | 4619 | 0.0264 |
| oll007_noUDG.SG | 1030 | Europe | Sweden | 4044.7 | 14.07 | 56.01 | 4617 | 0.0194 |
| I7780 | 197 | Asia | Russia | 6446.8 | 103.66 | 52.92 | 4615 | 0.0279 |
| HAN002_noUDG.SG | 550 | Europe | Russia | 3646.5 | 35.90 | 55.64 | 4605 | 0.0290 |
| STB001 | 391 | Asia | Russia | 6596.5 | 105.13 | 54.80 | 4604 | 0.0224 |
| I11474 | 155 | Asia | Iran | 3027.5 | 61.40 | 30.65 | 4600 | 0.0110 |
| I11476 | 155 | Asia | Iran | 3027.5 | 61.40 | 30.65 | 4600 | 0.0218 |
| I1272 | 630 | Europe | Spain | 4417.6 | -3.50 | 42.33 | 4595 | 0.0299 |
| SM-SGDLM6_noUDG | 386 | Asia | China | 7293.5 | 110.31 | 38.57 | 4589 | 0.0226 |
| I8364 | 949 | Europe | Spain | 4878.1 | -4.24 | 37.24 | 4588 | 0.0271 |
| I8365 | 949 | Europe | Spain | 4878.1 | -4.24 | 37.24 | 4588 | 0.0215 |
| I3447 | 142 | Asia | Kazakhstan | 4598.6 | 79.36 | 45.13 | 4577 | 0.0155 |
| I6695 | 916 | Europe | Czech Republic | 3537.1 | 13.78 | 50.55 | 4575 | 0.0257 |
| I6696 | 917 | Europe | Czech Republic | 3537.1 | 14.07 | 50.54 | 4575 | 0.0285 |
| MA2210_noUDG.SG | 995 | Europe | Turkey | 1321.9 | 34.30 | 38.63 | 4575 | 0.0237 |
| MA2212_noUDG.SG | 995 | Europe | Turkey | 1321.9 | 34.30 | 38.63 | 4575 | 0.0227 |
| MA2213_noUDG.SG | 995 | Europe | Turkey | 1321.9 | 34.30 | 38.63 | 4575 | 0.0177 |
| TORTE.SG | 541 | Europe | France | 3895.7 | 3.06 | 43.15 | 4574 | 0.0216 |
| I3949.DG | 219 | Asia | Russia | 5041.2 | 83.91 | 48.83 | 4569 | 0.0256 |
| GLZ001 | 119 | Asia | Russia | 6514.2 | 104.25 | 52.28 | 4556 | 0.0226 |
| ART009 | 470 | Europe | Turkey | 1427.6 | 38.36 | 38.38 | 4553 | 0.0116 |
| I6671 | 230 | Asia | Turkmenistan | 2630.8 | 56.25 | 38.35 | 4550 | 0.0193 |
| ETM010 | 423 | Both | Syria | 1028.7 | 36.80 | 35.80 | 4550 | 0.0149 |
| I1300 | 630 | Europe | Spain | 4417.6 | -3.50 | 42.33 | 4550 | 0.0231 |
| I5838 | 630 | Europe | Spain | 4417.6 | -3.50 | 42.33 | 4550 | 0.0179 |
| WD-WT1H16_noUDG | 412 | Asia | China | 7727.7 | 113.37 | 34.18 | 4536 | 0.0219 |
| KO1002 | 972 | Europe | Czech Republic | 3449.6 | 15.17 | 50.03 | 4533 | 0.0257 |
| MX190 | 1007 | Europe | Switzerland | 3569.2 | 8.22 | 47.26 | 4530 | 0.0177 |
| NAU001_noUDG.SG | 1015 | Europe | Russia | 3752.9 | 39.63 | 57.59 | 4526 | 0.0294 |

| | | | | | | | | |
|---|---|---|---|---|---|---|---|---|
| MX191 | 1007 | Europe | Switzerland | 3569.2 | 8.22 | 47.26 | 4523 | 0.0205 |
| I1392 | 655 | Europe | France | 3645.6 | 7.52 | 47.56 | 4516 | 0.0201 |
| NAU002_noUDG.SG | 1015 | Europe | Russia | 3752.9 | 39.63 | 57.59 | 4516 | 0.0260 |
| HAN004_noUDG.SG | 550 | Europe | Russia | 3646.5 | 35.90 | 55.64 | 4511 | 0.0263 |
| RK4002 | 371 | Asia | Russia | 2359.3 | 41.06 | 45.54 | 4510 | 0.0173 |
| MX198 | 1007 | Europe | Switzerland | 3569.2 | 8.22 | 47.26 | 4509 | 0.0199 |
| NIK008AB_noUDG.SG | 1021 | Europe | Russia | 3752.9 | 39.67 | 57.53 | 4508 | 0.0224 |
| I14689 | 669 | Europe | Albania | 2438.6 | 20.39 | 42.02 | 4505 | 0.0202 |
| HAL001_noUDG.SG | 549 | Europe | Russia | 3704.5 | 39.62 | 57.32 | 4504 | 0.0230 |
| CAK68.SG | 495 | Europe | Ireland | 4945.5 | -8.38 | 54.06 | 4503 | 0.0263 |
| I11456_enhanced | 155 | Asia | Iran | 3027.5 | 61.40 | 30.65 | 4500 | 0.0205 |
| I11478 | 155 | Asia | Iran | 3027.5 | 61.40 | 30.65 | 4500 | 0.0202 |
| I8728_enhanced | 155 | Asia | Iran | 3027.5 | 61.40 | 30.65 | 4500 | 0.0178 |
| STD001 | 1146 | Europe | Czech Republic | 3537.1 | 13.96 | 50.61 | 4500 | 0.0136 |
| C2034 | 52 | Asia | China | 5177.6 | 86.27 | 47.97 | 4498 | 0.0176 |
| Gyvakarai1_10bp_noUDG | 547 | Europe | Lithuania | 3643.5 | 24.91 | 55.92 | 4495 | 0.0247 |
| MX196 | 1007 | Europe | Switzerland | 3569.2 | 8.22 | 47.26 | 4495 | 0.0202 |
| RISE94_noUDG.SG | 1128 | Europe | Sweden | 4044.7 | 14.23 | 56.02 | 4487 | 0.0260 |
| KUR001 | 189 | Asia | Mongolia | 5419.2 | 88.71 | 49.34 | 4483 | 0.0237 |
| ber1M_noUDG.SG | 481 | Europe | Sweden | 4310.4 | 15.63 | 58.42 | 4483 | 0.0220 |
| VLI010 | 1253 | Europe | Czech Republic | 3537.1 | 14.45 | 50.37 | 4483 | 0.0234 |
| EBA2_noUDG.SG | 110 | Asia | Kazakhstan | 4733.4 | 76.73 | 52.63 | 4480 | 0.0235 |
| I15132 | 193 | Asia | Armenia | 2026.2 | 45.08 | 41.03 | 4479 | 0.0122 |
| I6714.SG | 219 | Asia | Russia | 5041.2 | 83.91 | 48.83 | 4479 | 0.0229 |
| LJM3_noUDG | 318 | Asia | China | 6588.7 | 101.80 | 36.51 | 4479 | 0.0213 |
| I2683 | 800 | Europe | Turkey | 1488.0 | 30.71 | 37.92 | 4479 | 0.0183 |
| OHR002 | 1029 | Europe | Czech Republic | 3495.2 | 14.34 | 50.05 | 4477 | 0.0195 |
| I1635 | 200 | Asia | Armenia | 2035.1 | 45.12 | 40.65 | 4476 | 0.0219 |
| GBVPL.SG | 541 | Europe | France | 3895.7 | 3.06 | 43.15 | 4474 | 0.0231 |
| I12978 | 170 | Asia | Mongolia | 5552.5 | 91.57 | 46.12 | 4472 | 0.0185 |
| I0103 | 564 | Europe | Germany | 3710.4 | 11.68 | 51.42 | 4472 | 0.0245 |

| | | | | | | | | |
|---|---|---|---|---|---|---|---|---|
| I2630 | 813 | Europe | United Kingdom | 5139.6 | -2.92 | 58.74 | 4471 | 0.0180 |
| ETM012 | 423 | Both | Syria | 1028.7 | 36.80 | 35.80 | 4470 | 0.0166 |
| PRU002 | 1056 | Europe | Czech Republic | 3495.2 | 14.31 | 50.08 | 4470 | 0.0206 |
| TIS002 | 1160 | Europe | Czech Republic | 3495.2 | 14.54 | 50.28 | 4470 | 0.0189 |
| irk068_noUDG.SG | 281 | Asia | Russia | 6589.0 | 105.68 | 54.01 | 4468 | 0.0205 |
| LD1174_noUDG.SG | 986 | Europe | Spain | 4878.1 | -4.42 | 37.41 | 4466 | 0.0249 |
| I7335 | 197 | Asia | Russia | 6446.8 | 103.66 | 52.92 | 4465 | 0.0236 |
| I1633 | 200 | Asia | Armenia | 2035.1 | 45.12 | 40.65 | 4465 | 0.0164 |
| RISE1164_noUDG.SG | 1121 | Europe | Poland | 3229.3 | 20.58 | 50.17 | 4463 | 0.0214 |
| I12957 | 170 | Asia | Mongolia | 5552.5 | 91.57 | 46.12 | 4462 | 0.0238 |
| I6713 | 152 | Asia | Russia | 5674.0 | 90.92 | 54.36 | 4450 | 0.0228 |
| I14788 | 670 | Europe | Turkey | 1180.9 | 37.17 | 36.69 | 4450 | 0.0127 |
| I14794 | 670 | Europe | Turkey | 1180.9 | 37.17 | 36.69 | 4450 | 0.0060 |
| I14797 | 670 | Europe | Turkey | 1180.9 | 37.17 | 36.69 | 4450 | 0.0136 |
| I14798 | 670 | Europe | Turkey | 1180.9 | 37.17 | 36.69 | 4450 | 0.0215 |
| VLI020 | 1253 | Europe | Czech Republic | 3537.1 | 14.45 | 50.37 | 4446 | 0.0197 |
| RISE718_noUDG.SG | 369 | Asia | Russia | 5674.0 | 91.51 | 54.37 | 4434 | 0.0259 |
| I2932 | 813 | Europe | United Kingdom | 5139.6 | -2.92 | 58.74 | 4433 | 0.0206 |
| GLZ002 | 119 | Asia | Russia | 6514.2 | 104.25 | 52.28 | 4430 | 0.0267 |
| PRU003 | 1056 | Europe | Czech Republic | 3495.2 | 14.31 | 50.08 | 4427 | 0.0268 |
| MA973_noUDG.SG | 996 | Europe | Estonia | 3852.9 | 26.15 | 58.33 | 4416 | 0.0218 |
| BOL003_noUDG.SG | 487 | Europe | Russia | 3844.9 | 36.63 | 57.74 | 4414 | 0.0216 |
| I1277 | 630 | Europe | Spain | 4417.6 | -3.50 | 42.33 | 4403 | 0.0209 |
| N44_noUDG.SG | 1011 | Europe | Poland | 3539.2 | 18.96 | 52.62 | 4400 | 0.0212 |
| pcw040_noUDG.SG | 1036 | Europe | Poland | 3103.6 | 22.86 | 49.92 | 4400 | 0.0179 |
| I0108 | 569 | Europe | Germany | 3756.0 | 11.54 | 51.45 | 4399 | 0.0183 |
| MX188 | 1007 | Europe | Switzerland | 3569.2 | 8.22 | 47.26 | 4399 | 0.0178 |
| KON004 | 976 | Europe | Czech Republic | 3537.1 | 13.66 | 50.56 | 4398 | 0.0226 |
| I5748 | 856 | Europe | Netherlands | 4151.8 | 5.10 | 52.73 | 4383 | 0.0170 |
| RISE552_noUDG.SG | 364 | Asia | Russia | 2544.6 | 43.33 | 46.62 | 4382 | 0.0243 |
| LD270_noUDG.SG | 987 | Europe | Portugal | 4761.4 | -6.93 | 41.71 | 4380 | 0.0234 |

| | | | | | | | | |
|---|---|---|---|---|---|---|---|---|
| N45_noUDG.SG | 1011 | Europe | Poland | 3539.2 | 18.96 | 52.62 | 4375 | 0.0206 |
| N47_noUDG.SG | 1011 | Europe | Poland | 3539.2 | 18.96 | 52.62 | 4375 | 0.0164 |
| N49_noUDG.SG | 1011 | Europe | Poland | 3539.2 | 18.96 | 52.62 | 4375 | 0.0312 |
| I7207 | 678 | Europe | Czech Republic | 3537.1 | 14.07 | 50.41 | 4370 | 0.0246 |
| EHU002 | 528 | Europe | Spain | 4402.7 | -3.48 | 42.42 | 4367 | 0.0184 |
| I0104 | 564 | Europe | Germany | 3710.4 | 11.68 | 51.42 | 4358 | 0.0186 |
| I1730 | 427 | Both | Jordan | 627.3 | 35.98 | 31.99 | 4356 | 0.0191 |
| I2495 | 800 | Europe | Turkey | 1488.0 | 30.71 | 37.92 | 4355 | 0.0208 |
| SEC002 | 1137 | Europe | Italy | 3521.6 | 8.60 | 40.79 | 4352 | 0.0195 |
| MX197 | 1007 | Europe | Switzerland | 3569.2 | 8.22 | 47.26 | 4351 | 0.0118 |
| I0374 | 133 | Asia | Russia | 2970.9 | 44.67 | 49.97 | 4350 | 0.0226 |
| irk025_noUDG.SG | 276 | Asia | Russia | 6446.8 | 103.52 | 52.98 | 4350 | 0.0240 |
| I0459 | 575 | Europe | Spain | 4484.7 | -3.75 | 42.40 | 4350 | 0.0212 |
| CAH005 | 494 | Europe | Czech Republic | 3582.7 | 13.37 | 50.37 | 4349 | 0.0249 |
| SEC005 | 1137 | Europe | Italy | 3521.6 | 8.60 | 40.79 | 4348 | 0.0193 |
| I2514 | 217 | Asia | Iran | 2407.9 | 54.38 | 36.15 | 4347 | 0.0145 |
| SUC006 | 1093 | Europe | Italy | 3570.8 | 8.44 | 40.81 | 4347 | 0.0198 |
| UNTA58_68Sk2 | 1170 | Europe | Germany | 3480.3 | 10.89 | 48.32 | 4347 | 0.0170 |
| VK531_noUDG.SG | 1240 | Europe | Norway | 5247.1 | 18.00 | 69.47 | 4347 | 0.0237 |
| I1706 | 427 | Both | Jordan | 627.3 | 35.98 | 31.99 | 4345 | 0.0161 |
| VLI016 | 1253 | Europe | Czech Republic | 3537.1 | 14.45 | 50.37 | 4345 | 0.0234 |
| ART001 | 470 | Europe | Turkey | 1427.6 | 38.36 | 38.38 | 4342 | 0.0193 |
| KOP003 | 977 | Europe | Czech Republic | 3449.6 | 15.21 | 50.01 | 4342 | 0.0141 |
| irk061_noUDG.SG | 280 | Asia | Russia | 6589.0 | 105.84 | 53.97 | 4341 | 0.0242 |
| I14340 | 193 | Asia | Armenia | 2026.2 | 45.08 | 41.03 | 4340 | 0.0148 |
| I13599 | 186 | Asia | Armenia | 1927.6 | 43.95 | 40.78 | 4337 | 0.0192 |
| SEC004 | 1137 | Europe | Italy | 3521.6 | 8.60 | 40.79 | 4337 | 0.0259 |
| GBVPK.SG | 541 | Europe | France | 3895.7 | 3.06 | 43.15 | 4336 | 0.0214 |
| VLI017 | 1253 | Europe | Czech Republic | 3537.1 | 14.45 | 50.37 | 4336 | 0.0171 |
| I5379 | 890 | Europe | United Kingdom | 4423.8 | -2.02 | 50.92 | 4333 | 0.0170 |
| I7278 | 933 | Europe | Czech Republic | 3495.2 | 14.16 | 50.19 | 4333 | 0.0303 |

| | | | | | | | | |
|---|---|---|---|---|---|---|---|---|
| KO1005 | 972 | Europe | Czech Republic | 3449.6 | 15.17 | 50.03 | 4331 | 0.0199 |
| PRU001 | 1056 | Europe | Czech Republic | 3495.2 | 14.31 | 50.08 | 4331 | 0.0249 |
| SEC006 | 1137 | Europe | Italy | 3521.6 | 8.60 | 40.79 | 4331 | 0.0250 |
| I14677 | 668 | Europe | Italy | 3521.6 | 8.59 | 40.79 | 4329 | 0.0247 |
| DA337_noUDG.SG | 87 | Asia | Russia | 6444.1 | 103.70 | 51.70 | 4328 | 0.0249 |
| I11735 | 161 | Asia | Kazakhstan | 3222.8 | 49.34 | 50.77 | 4328 | 0.0207 |
| TU908_SX21 | 1167 | Europe | Switzerland | 3514.3 | 9.29 | 47.45 | 4327 | 0.0130 |
| I0118 | 571 | Europe | Germany | 3710.4 | 11.63 | 51.45 | 4325 | 0.0228 |
| I7208 | 678 | Europe | Czech Republic | 3537.1 | 14.07 | 50.41 | 4324 | 0.0204 |
| SUC007 | 1093 | Europe | Italy | 3570.8 | 8.44 | 40.81 | 4321 | 0.0207 |
| I5367 | 888 | Europe | United Kingdom | 5038.8 | -6.46 | 56.68 | 4312 | 0.0217 |
| I5243 | 885 | Europe | Serbia | 2590.5 | 22.01 | 44.60 | 4306 | 0.0210 |
| Olsund | 1031 | Europe | Sweden | 4543.7 | 17.00 | 61.66 | 4306 | 0.0182 |
| pcw362_noUDG.SG | 1038 | Europe | Poland | 3150.3 | 23.66 | 50.46 | 4305 | 0.0178 |
| I0111 | 569 | Europe | Germany | 3756.0 | 11.54 | 51.45 | 4303 | 0.0246 |
| I8745 | 133 | Asia | Russia | 2970.9 | 44.67 | 49.97 | 4300 | 0.0189 |
| RISE672_noUDG.SG | 361 | Asia | Russia | 5568.9 | 90.21 | 53.16 | 4300 | 0.0303 |
| RISE680_noUDG.SG | 368 | Asia | Russia | 5581.2 | 90.36 | 53.71 | 4300 | 0.0224 |
| RISE685_noUDG.SG | 368 | Asia | Russia | 5581.2 | 90.36 | 53.71 | 4300 | 0.0254 |
| I3529 | 778 | Europe | Hungary | 3004.9 | 19.04 | 47.62 | 4300 | 0.0202 |
| I4178 | 825 | Europe | Hungary | 3004.9 | 19.02 | 47.38 | 4300 | 0.0250 |
| I7044 | 930 | Europe | Hungary | 3004.9 | 19.05 | 47.43 | 4300 | 0.0173 |
| pcw361_noUDG.SG | 1038 | Europe | Poland | 3150.3 | 23.66 | 50.46 | 4300 | 0.0282 |
| I10345 | 586 | Europe | France | 3649.9 | 6.37 | 44.48 | 4299 | 0.0206 |
| I2365 | 777 | Europe | Hungary | 3004.9 | 19.05 | 47.60 | 4299 | 0.0216 |
| I15942 | 688 | Europe | Italy | 3570.8 | 8.32 | 40.63 | 4297 | 0.0129 |
| JCKM1-1_noUDG | 286 | Asia | China | 6662.8 | 102.53 | 36.92 | 4296 | 0.0209 |
| I0049 | 564 | Europe | Germany | 3710.4 | 11.68 | 51.42 | 4295 | 0.0180 |
| I13467 | 641 | Europe | Czech Republic | 3495.2 | 14.34 | 50.04 | 4289 | 0.0274 |
| I7289 | 678 | Europe | Czech Republic | 3537.1 | 14.07 | 50.41 | 4289 | 0.0233 |
| ETM023 | 423 | Both | Syria | 1028.7 | 36.80 | 35.80 | 4275 | 0.0176 |

| | | | | | | | | |
|---|---|---|---|---|---|---|---|---|
| CAH010 | 494 | Europe | Czech Republic | 3582.7 | 13.37 | 50.37 | 4275 | 0.0218 |
| RISE664_noUDG.SG | 367 | Asia | Russia | 5648.2 | 91.03 | 53.55 | 4270 | 0.0212 |
| I2786 | 825 | Europe | Hungary | 3004.9 | 19.02 | 47.38 | 4270 | 0.0202 |
| I7209 | 678 | Europe | Czech Republic | 3537.1 | 14.07 | 50.41 | 4269 | 0.0246 |
| PRU004 | 1056 | Europe | Czech Republic | 3495.2 | 14.31 | 50.08 | 4264 | 0.0158 |
| I2787 | 825 | Europe | Hungary | 3004.9 | 19.02 | 47.38 | 4254 | 0.0203 |
| irk022_noUDG.SG | 275 | Asia | Russia | 6446.8 | 103.42 | 52.99 | 4251 | 0.0234 |
| HOP003 | 558 | Europe | Czech Republic | 3495.2 | 14.28 | 50.08 | 4250 | 0.0270 |
| I13028 | 638 | Europe | Netherlands | 4077.7 | 4.87 | 51.87 | 4250 | 0.0184 |
| I1381 | 652 | Europe | France | 3819.7 | 6.10 | 49.15 | 4250 | 0.0187 |
| I2417 | 702 | Europe | United Kingdom | 4423.8 | -1.77 | 51.17 | 4250 | 0.0151 |
| I7195 | 871 | Europe | Czech Republic | 3495.2 | 14.37 | 50.05 | 4250 | 0.0254 |
| I7196 | 871 | Europe | Czech Republic | 3495.2 | 14.37 | 50.05 | 4250 | 0.0185 |
| I7200 | 871 | Europe | Czech Republic | 3495.2 | 14.37 | 50.05 | 4250 | 0.0190 |
| I7201 | 871 | Europe | Czech Republic | 3495.2 | 14.37 | 50.05 | 4250 | 0.0234 |
| I7202 | 871 | Europe | Czech Republic | 3495.2 | 14.37 | 50.05 | 4250 | 0.0216 |
| I4950 | 874 | Europe | United Kingdom | 4439.0 | -1.81 | 51.29 | 4250 | 0.0249 |
| I5512 | 895 | Europe | United Kingdom | 4439.0 | -1.76 | 51.24 | 4250 | 0.0213 |
| I5513 | 896 | Europe | United Kingdom | 4423.8 | -1.77 | 51.22 | 4250 | 0.0241 |
| I5521 | 897 | Europe | Germany | 3480.3 | 10.89 | 48.33 | 4250 | 0.0196 |
| I5523 | 898 | Europe | Germany | 3417.1 | 12.71 | 48.66 | 4250 | 0.0232 |
| I5524 | 898 | Europe | Germany | 3417.1 | 12.71 | 48.66 | 4250 | 0.0226 |
| I5525 | 898 | Europe | Germany | 3417.1 | 12.71 | 48.66 | 4250 | 0.0199 |
| I5531 | 899 | Europe | Germany | 3517.4 | 11.33 | 48.71 | 4250 | 0.0205 |
| KOP002 | 977 | Europe | Czech Republic | 3449.6 | 15.21 | 50.01 | 4250 | 0.0103 |
| Log02.SG | 990 | Europe | Greece | 2206.0 | 21.82 | 39.98 | 4250 | 0.0197 |
| Log04.SG | 990 | Europe | Greece | 2206.0 | 21.82 | 39.98 | 4250 | 0.0179 |
| Pta08.SG | 1059 | Europe | Greece | 1797.6 | 26.11 | 35.20 | 4250 | 0.0137 |
| I2450 | 790 | Europe | United Kingdom | 4397.4 | -1.30 | 51.68 | 4248 | 0.0207 |
| I14678 | 668 | Europe | Italy | 3521.6 | 8.59 | 40.79 | 4247 | 0.0228 |
| I14792 | 670 | Europe | Turkey | 1180.9 | 37.17 | 36.69 | 4247 | 0.0155 |

| | | | | | | | | |
|---|---|---|---|---|---|---|---|---|
| I7212 | 678 | Europe | Czech Republic | 3537.1 | 14.07 | 50.41 | 4247 | 0.0126 |
| I2741 | 825 | Europe | Hungary | 3004.9 | 19.02 | 47.38 | 4245 | 0.0224 |
| I0461 | 575 | Europe | Spain | 4484.7 | -3.75 | 42.40 | 4241 | 0.0229 |
| HOP001 | 558 | Europe | Czech Republic | 3495.2 | 14.28 | 50.08 | 4240 | 0.0219 |
| RK4001 | 371 | Asia | Russia | 2359.3 | 41.06 | 45.54 | 4238 | 0.0212 |
| I2418 | 702 | Europe | United Kingdom | 4423.8 | -1.77 | 51.17 | 4237 | 0.0219 |
| pcw260_noUDG.SG | 1037 | Europe | Poland | 3274.4 | 20.57 | 50.36 | 4237 | 0.0266 |
| SEC001 | 1137 | Europe | Italy | 3521.6 | 8.60 | 40.79 | 4237 | 0.0216 |
| MX199 | 1007 | Europe | Switzerland | 3569.2 | 8.22 | 47.26 | 4235 | 0.0179 |
| I7213 | 678 | Europe | Czech Republic | 3537.1 | 14.07 | 50.41 | 4231 | 0.0189 |
| I2459 | 702 | Europe | United Kingdom | 4423.8 | -1.77 | 51.17 | 4231 | 0.0209 |
| I7045 | 930 | Europe | Hungary | 3004.9 | 19.05 | 47.43 | 4231 | 0.0193 |
| I5385 | 891 | Europe | United Kingdom | 5139.6 | -3.40 | 58.36 | 4230 | 0.0132 |
| I27380 | 822 | Europe | United Kingdom | 4352.6 | -0.90 | 50.87 | 4228 | 0.0194 |
| I6581 | 914 | Europe | Poland | 3315.8 | 18.10 | 50.09 | 4228 | 0.0174 |
| RISE1281_noUDG.SG | 1124 | Europe | Denmark | 4214.4 | 10.82 | 56.50 | 4226 | 0.0271 |
| I1549 | 566 | Europe | Germany | 3796.2 | 10.91 | 51.82 | 4225 | 0.0203 |
| MX195 | 1007 | Europe | Switzerland | 3569.2 | 8.22 | 47.26 | 4225 | 0.0125 |
| C2048 | 53 | Asia | China | 5552.6 | 90.75 | 46.48 | 4224 | 0.0228 |
| I0112 | 570 | Europe | Germany | 3796.2 | 11.14 | 51.79 | 4224 | 0.0266 |
| HOP004 | 558 | Europe | Czech Republic | 3495.2 | 14.28 | 50.08 | 4222 | 0.0263 |
| I7286 | 678 | Europe | Czech Republic | 3537.1 | 14.07 | 50.41 | 4221 | 0.0175 |
| I2364 | 777 | Europe | Hungary | 3004.9 | 19.05 | 47.60 | 4220 | 0.0177 |
| I8561 | 950 | Europe | Italy | 2968.4 | 14.08 | 37.95 | 4212 | 0.0196 |
| I4246 | 861 | Europe | Spain | 4482.6 | -3.50 | 40.44 | 4209 | 0.0152 |
| I15940 | 688 | Europe | Italy | 3570.8 | 8.32 | 40.63 | 4203 | 0.0079 |
| C1699 | 48 | Asia | China | 5177.6 | 86.42 | 48.06 | 4200 | 0.0294 |
| C1700 | 48 | Asia | China | 5177.6 | 86.42 | 48.06 | 4200 | 0.0195 |
| C1701 | 48 | Asia | China | 5177.6 | 86.42 | 48.06 | 4200 | 0.0237 |
| C1702 | 48 | Asia | China | 5177.6 | 86.42 | 48.06 | 4200 | 0.0186 |
| C1704 | 48 | Asia | China | 5177.6 | 86.42 | 48.06 | 4200 | 0.0237 |

| | | | | | | | | |
|---|---|---|---|---|---|---|---|---|
| N4b2_noUDG.SG | 329 | Asia | Russia | 8170.2 | 132.33 | 62.07 | 4200 | 0.0242 |
| CovaMoura364_noUDG.SG | 509 | Europe | Portugal | 5047.7 | -9.22 | 38.75 | 4200 | 0.0203 |
| CovaMoura9B_noUDG.SG | 509 | Europe | Portugal | 5047.7 | -9.22 | 38.75 | 4200 | 0.0222 |
| I7205 | 678 | Europe | Czech Republic | 3537.1 | 14.07 | 50.41 | 4200 | 0.0233 |
| I7214 | 678 | Europe | Czech Republic | 3537.1 | 14.07 | 50.41 | 4200 | 0.0238 |
| I7282 | 678 | Europe | Czech Republic | 3537.1 | 14.07 | 50.41 | 4200 | 0.0158 |
| I7283 | 678 | Europe | Czech Republic | 3537.1 | 14.07 | 50.41 | 4200 | 0.0192 |
| I7287 | 678 | Europe | Czech Republic | 3537.1 | 14.07 | 50.41 | 4200 | 0.0219 |
| I7290 | 678 | Europe | Czech Republic | 3537.1 | 14.07 | 50.41 | 4200 | 0.0248 |
| I5023 | 875 | Europe | Germany | 3417.1 | 13.02 | 48.69 | 4200 | 0.0208 |
| I5519 | 897 | Europe | Germany | 3480.3 | 10.89 | 48.33 | 4200 | 0.0163 |
| I5833 | 907 | Europe | Germany | 3417.1 | 12.75 | 48.84 | 4200 | 0.0241 |
| I5834 | 907 | Europe | Germany | 3417.1 | 12.75 | 48.84 | 4200 | 0.0151 |
| I5835 | 907 | Europe | Germany | 3417.1 | 12.75 | 48.84 | 4200 | 0.0170 |
| I6590 | 907 | Europe | Germany | 3417.1 | 12.75 | 48.84 | 4200 | 0.0212 |
| I6591 | 907 | Europe | Germany | 3417.1 | 12.75 | 48.84 | 4200 | 0.0150 |
| I6624 | 907 | Europe | Germany | 3417.1 | 12.75 | 48.84 | 4200 | 0.0238 |
| I7251 | 933 | Europe | Czech Republic | 3495.2 | 14.16 | 50.19 | 4200 | 0.0212 |
| I7269 | 933 | Europe | Czech Republic | 3495.2 | 14.16 | 50.19 | 4200 | 0.0198 |
| I7275 | 933 | Europe | Czech Republic | 3495.2 | 14.16 | 50.19 | 4200 | 0.0183 |
| I7276 | 933 | Europe | Czech Republic | 3495.2 | 14.16 | 50.19 | 4200 | 0.0224 |
| I7279 | 933 | Europe | Czech Republic | 3495.2 | 14.16 | 50.19 | 4200 | 0.0224 |
| I7280 | 933 | Europe | Czech Republic | 3495.2 | 14.16 | 50.19 | 4200 | 0.0233 |
| I15941 | 688 | Europe | Italy | 3570.8 | 8.32 | 40.63 | 4196 | 0.0201 |
| EKA1_noUDG.SG | 529 | Europe | Estonia | 3901.2 | 26.73 | 58.37 | 4195 | 0.0247 |
| I1382 | 652 | Europe | France | 3819.7 | 6.10 | 49.15 | 4195 | 0.0248 |
| I4521 | 426 | Both | Israel | 629.1 | 35.02 | 32.58 | 4194 | 0.0129 |
| I7281 | 641 | Europe | Czech Republic | 3495.2 | 14.34 | 50.04 | 4194 | 0.0212 |
| WEHR_1192SkA | 1260 | Europe | Germany | 3480.3 | 10.81 | 48.25 | 4193 | 0.0214 |
| SUC004 | 1093 | Europe | Italy | 3570.8 | 8.44 | 40.81 | 4185 | 0.0229 |
| I7211 | 678 | Europe | Czech Republic | 3537.1 | 14.07 | 50.41 | 4183 | 0.0198 |

| | | | | | | | | |
|---|---|---|---|---|---|---|---|---|
| I0059 | 566 | Europe | Germany | 3796.2 | 10.91 | 51.82 | 4179 | 0.0225 |
| I4945 | 870 | Europe | Czech Republic | 3495.2 | 14.46 | 50.12 | 4177 | 0.0263 |
| I3589 | 849 | Europe | Germany | 3465.3 | 12.53 | 48.88 | 4175 | 0.0146 |
| I3594 | 849 | Europe | Germany | 3465.3 | 12.53 | 48.88 | 4175 | 0.0276 |
| I3601 | 849 | Europe | Germany | 3465.3 | 12.53 | 48.88 | 4175 | 0.0210 |
| ISC001 | 957 | Europe | Italy | 3570.8 | 8.54 | 40.66 | 4175 | 0.0156 |
| I6582 | 914 | Europe | Poland | 3315.8 | 18.10 | 50.09 | 4171 | 0.0191 |
| kra001_noUDG.SG | 305 | Asia | Russia | 6049.2 | 95.82 | 56.19 | 4170 | 0.0176 |
| I4885 | 870 | Europe | Czech Republic | 3495.2 | 14.46 | 50.12 | 4170 | 0.0254 |
| HUGO_168 | 559 | Europe | Germany | 3480.3 | 10.90 | 48.33 | 4167 | 0.0215 |
| KO1004 | 972 | Europe | Czech Republic | 3449.6 | 15.17 | 50.03 | 4166 | 0.0199 |
| PDA005 | 1039 | Europe | Czech Republic | 3495.2 | 14.49 | 50.15 | 4164 | 0.0215 |
| I4896 | 871 | Europe | Czech Republic | 3495.2 | 14.37 | 50.05 | 4163 | 0.0262 |
| KO1003 | 972 | Europe | Czech Republic | 3449.6 | 15.17 | 50.03 | 4160 | 0.0231 |
| I4229 | 509 | Europe | Portugal | 5047.7 | -9.22 | 38.75 | 4158 | 0.0290 |
| I5514 | 871 | Europe | Czech Republic | 3495.2 | 14.37 | 50.05 | 4155 | 0.0161 |
| I6579 | 913 | Europe | Poland | 3446.8 | 17.07 | 51.04 | 4155 | 0.0182 |
| I0244 | 128 | Asia | Russia | 3576.4 | 50.57 | 53.59 | 4154 | 0.0280 |
| I4139 | 858 | Europe | Czech Republic | 3495.2 | 14.32 | 50.17 | 4150 | 0.0208 |
| I6775 | 928 | Europe | United Kingdom | 4495.0 | -3.08 | 51.12 | 4150 | 0.0261 |
| I6778 | 929 | Europe | United Kingdom | 4423.8 | -1.81 | 51.15 | 4150 | 0.0121 |
| R27.SG | 1093 | Europe | Italy | 3570.8 | 8.44 | 40.81 | 4150 | 0.0187 |
| R29.SG | 1093 | Europe | Italy | 3570.8 | 8.44 | 40.81 | 4150 | 0.0247 |
| VLI024 | 1253 | Europe | Czech Republic | 3537.1 | 14.45 | 50.37 | 4150 | 0.0229 |
| VLI026 | 1253 | Europe | Czech Republic | 3537.1 | 14.45 | 50.37 | 4150 | 0.0209 |
| VLI029 | 1253 | Europe | Czech Republic | 3537.1 | 14.45 | 50.37 | 4150 | 0.0211 |
| I6674 | 230 | Asia | Turkmenistan | 2630.8 | 56.25 | 38.35 | 4148 | 0.0176 |
| I0113 | 570 | Europe | Germany | 3796.2 | 11.14 | 51.79 | 4146 | 0.0225 |
| SUC005 | 1093 | Europe | Italy | 3570.8 | 8.44 | 40.81 | 4146 | 0.0141 |
| I6583 | 915 | Europe | Poland | 3446.8 | 17.02 | 50.96 | 4138 | 0.0232 |
| I12786 | 633 | Europe | United Kingdom | 4467.2 | -1.69 | 51.70 | 4137 | 0.0227 |

| | | | | | | | | |
|---|---|---|---|---|---|---|---|---|
| C3344 | 55 | Asia | China | 5172.4 | 85.87 | 47.44 | 4131 | 0.0189 |
| I11442 | 611 | Europe | Italy | 3057.3 | 13.48 | 37.91 | 4131 | 0.0182 |
| I4889 | 870 | Europe | Czech Republic | 3495.2 | 14.46 | 50.12 | 4131 | 0.0183 |
| I4891 | 870 | Europe | Czech Republic | 3495.2 | 14.46 | 50.12 | 4131 | 0.0206 |
| PDA003 | 1039 | Europe | Czech Republic | 3495.2 | 14.49 | 50.15 | 4131 | 0.0183 |
| I6537 | 911 | Europe | Poland | 3315.8 | 18.21 | 50.11 | 4130 | 0.0184 |
| I2453 | 792 | Europe | United Kingdom | 4405.4 | -0.35 | 52.67 | 4122 | 0.0215 |
| EHU001 | 528 | Europe | Spain | 4402.7 | -3.48 | 42.42 | 4121 | 0.0203 |
| I3123 | 611 | Europe | Italy | 3057.3 | 13.48 | 37.91 | 4121 | 0.0203 |
| I6774 | 927 | Europe | United Kingdom | 4296.4 | -0.12 | 50.86 | 4121 | 0.0204 |
| I2568 | 809 | Europe | United Kingdom | 4814.3 | -2.44 | 55.97 | 4113 | 0.0216 |
| I6531 | 910 | Europe | Poland | 3315.8 | 18.22 | 50.23 | 4113 | 0.0203 |
| EBA1_noUDG.SG | 109 | Asia | Kazakhstan | 4533.8 | 74.66 | 51.63 | 4108 | 0.0221 |
| I16424 | 632 | Europe | United Kingdom | 4624.9 | -5.00 | 50.54 | 4106 | 0.0249 |
| I4895 | 871 | Europe | Czech Republic | 3495.2 | 14.37 | 50.05 | 4106 | 0.0263 |
| HUGO_190 | 559 | Europe | Germany | 3480.3 | 10.90 | 48.33 | 4103 | 0.0254 |
| I11159 | 609 | Europe | Czech Republic | 3495.2 | 14.36 | 50.05 | 4100 | 0.0210 |
| I20182 | 757 | Europe | Bulgaria | 2071.2 | 26.33 | 41.73 | 4100 | 0.0145 |
| I6623 | 861 | Europe | Spain | 4482.6 | -3.50 | 40.44 | 4100 | 0.0211 |
| PMI001 | 1046 | Europe | Czech Republic | 3495.2 | 14.54 | 50.16 | 4096 | 0.0226 |
| I0117 | 564 | Europe | Germany | 3710.4 | 11.68 | 51.42 | 4092 | 0.0223 |
| I2443 | 752 | Europe | United Kingdom | 4397.4 | -1.31 | 51.80 | 4091 | 0.0212 |
| I11443 | 611 | Europe | Italy | 3057.3 | 13.48 | 37.91 | 4090 | 0.0276 |
| PLTM310_noUDG | 349 | Asia | China | 7833.7 | 114.92 | 33.71 | 4088 | 0.0228 |
| I4886 | 870 | Europe | Czech Republic | 3495.2 | 14.46 | 50.12 | 4088 | 0.0233 |
| MX193 | 1007 | Europe | Switzerland | 3569.2 | 8.22 | 47.26 | 4087 | 0.0259 |
| I2596 | 702 | Europe | United Kingdom | 4423.8 | -1.77 | 51.17 | 4086 | 0.0202 |
| BNL005 | 486 | Europe | Czech Republic | 3495.2 | 14.65 | 50.17 | 4084 | 0.0203 |
| RISE98_noUDG.SG | 1130 | Europe | Sweden | 4060.0 | 13.45 | 55.38 | 4082 | 0.0260 |
| SUC002 | 1093 | Europe | Italy | 3570.8 | 8.44 | 40.81 | 4081 | 0.0285 |
| I2597 | 702 | Europe | United Kingdom | 4423.8 | -1.77 | 51.17 | 4080 | 0.0157 |

| | | | | | | | | |
|---|---|---|---|---|---|---|---|---|
| I11526 | 157 | Asia | Kyrgyzstan | 4314.5 | 75.89 | 41.43 | 4079 | 0.0190 |
| SUA002 | 956 | Europe | Italy | 3620.3 | 9.24 | 39.87 | 4076 | 0.0154 |
| I20997 | 767 | Europe | United Kingdom | 4656.2 | -3.11 | 54.16 | 4075 | 0.0179 |
| I3122 | 611 | Europe | Italy | 3057.3 | 13.48 | 37.91 | 4073 | 0.0244 |
| I2604 | 648 | Europe | United Kingdom | 4352.6 | -1.36 | 51.15 | 4073 | 0.0220 |
| I7210 | 678 | Europe | Czech Republic | 3537.1 | 14.07 | 50.41 | 4071 | 0.0217 |
| I4887 | 870 | Europe | Czech Republic | 3495.2 | 14.46 | 50.12 | 4071 | 0.0155 |
| BNL006 | 486 | Europe | Czech Republic | 3495.2 | 14.65 | 50.17 | 4070 | 0.0225 |
| RISE516_noUDG.SG | 361 | Asia | Russia | 5568.9 | 90.21 | 53.16 | 4065 | 0.0314 |
| I3874 | 586 | Europe | France | 3649.9 | 6.37 | 44.48 | 4065 | 0.0184 |
| RISE674_noUDG.SG | 361 | Asia | Russia | 5568.9 | 90.21 | 53.16 | 4064 | 0.0187 |
| I3256 | 607 | Europe | United Kingdom | 4389.8 | 0.11 | 52.17 | 4063 | 0.0272 |
| I1784 | 207 | Asia | Turkmenistan | 3066.1 | 62.03 | 38.21 | 4060 | 0.0196 |
| MX189 | 1007 | Europe | Switzerland | 3569.2 | 8.22 | 47.26 | 4058 | 0.0116 |
| PDA002 | 1039 | Europe | Czech Republic | 3495.2 | 14.49 | 50.15 | 4057 | 0.0163 |
| I2457 | 702 | Europe | United Kingdom | 4423.8 | -1.77 | 51.17 | 4056 | 0.0189 |
| I14676 | 668 | Europe | Italy | 3521.6 | 8.59 | 40.79 | 4053 | 0.0200 |
| PDA004 | 1039 | Europe | Czech Republic | 3495.2 | 14.49 | 50.15 | 4053 | 0.0128 |
| I2452 | 791 | Europe | United Kingdom | 4512.9 | -1.56 | 52.85 | 4052 | 0.0210 |
| MX288 | 1009 | Europe | Germany | 3550.6 | 8.87 | 47.77 | 4051 | 0.0227 |
| SM-SGDLM27_noUDG | 386 | Asia | China | 7293.5 | 110.31 | 38.57 | 4050 | 0.0251 |
| I5750 | 856 | Europe | Netherlands | 4151.8 | 5.10 | 52.73 | 4050 | 0.0216 |
| I6777 | 929 | Europe | United Kingdom | 4423.8 | -1.81 | 51.15 | 4050 | 0.0179 |
| PLTM311_noUDG | 349 | Asia | China | 7833.7 | 114.92 | 33.71 | 4049 | 0.0205 |
| I2445 | 752 | Europe | United Kingdom | 4397.4 | -1.31 | 51.80 | 4048 | 0.0238 |
| I11737 | 161 | Asia | Kazakhstan | 3222.8 | 49.34 | 50.77 | 4046 | 0.0216 |
| I1767 | 716 | Europe | United Kingdom | 4635.1 | -1.31 | 54.52 | 4045 | 0.0280 |
| KNE002 | 971 | Europe | Czech Republic | 3495.2 | 14.27 | 50.12 | 4043 | 0.0287 |
| I4354_noUDG | 115 | Asia | Iran | 1677.7 | 45.46 | 37.01 | 4042 | 0.0135 |
| I2454 | 793 | Europe | United Kingdom | 4389.8 | 0.03 | 52.34 | 4042 | 0.0291 |
| I13027 | 637 | Europe | Netherlands | 4077.7 | 4.86 | 51.89 | 4039 | 0.0210 |

| | | | | | | | | |
|---|---|---|---|---|---|---|---|---|
| I7199 | 871 | Europe | Czech Republic | 3495.2 | 14.37 | 50.05 | 4039 | 0.0181 |
| I4888 | 870 | Europe | Czech Republic | 3495.2 | 14.46 | 50.12 | 4038 | 0.0190 |
| KBD002 | 289 | Asia | Russia | 2207.8 | 42.72 | 43.83 | 4035 | 0.0170 |
| KBD001 | 289 | Asia | Russia | 2207.8 | 42.72 | 43.83 | 4030 | 0.0185 |
| I4074 | 856 | Europe | Netherlands | 4151.8 | 5.10 | 52.73 | 4030 | 0.0212 |
| I1705 | 427 | Both | Jordan | 627.3 | 35.98 | 31.99 | 4029 | 0.0171 |
| BAS024 | 479 | Europe | Spain | 4728.9 | -1.56 | 37.76 | 4029 | 0.0184 |
| I1502 | 680 | Europe | Hungary | 2867.4 | 20.83 | 47.17 | 4026 | 0.0272 |
| I7198 | 871 | Europe | Czech Republic | 3495.2 | 14.37 | 50.05 | 4026 | 0.0152 |
| I11158 | 609 | Europe | Czech Republic | 3495.2 | 14.36 | 50.05 | 4025 | 0.0218 |
| I14675 | 668 | Europe | Italy | 3521.6 | 8.59 | 40.79 | 4021 | 0.0196 |
| SUC001 | 1093 | Europe | Italy | 3570.8 | 8.44 | 40.81 | 4020 | 0.0215 |
| I2567 | 809 | Europe | United Kingdom | 4814.3 | -2.44 | 55.97 | 4018 | 0.0218 |
| KNE001 | 971 | Europe | Czech Republic | 3495.2 | 14.27 | 50.12 | 4017 | 0.0208 |
| PDA001 | 1039 | Europe | Czech Republic | 3495.2 | 14.49 | 50.15 | 4017 | 0.0192 |
| I2477 | 795 | Europe | Italy | 3360.5 | 10.33 | 44.80 | 4015 | 0.0183 |
| I7043 | 930 | Europe | Hungary | 3004.9 | 19.05 | 47.43 | 4011 | 0.0234 |
| I7249 | 933 | Europe | Czech Republic | 3495.2 | 14.16 | 50.19 | 4011 | 0.0227 |
| I2478 | 796 | Europe | Italy | 3360.5 | 10.29 | 44.78 | 4006 | 0.0230 |
| I12510 | 627 | Europe | Moldova | 2657.9 | 28.85 | 47.92 | 4003 | 0.0167 |
| I0114 | 564 | Europe | Germany | 3710.4 | 11.68 | 51.42 | 4002 | 0.0261 |
| I0419 | 134 | Asia | Russia | 3487.5 | 50.99 | 52.91 | 4000 | 0.0223 |
| I0070 | 568 | Europe | Greece | 1797.6 | 25.83 | 35.08 | 4000 | 0.0208 |
| I0071 | 568 | Europe | Greece | 1797.6 | 25.83 | 35.08 | 4000 | 0.0223 |
| I0073 | 568 | Europe | Greece | 1797.6 | 25.83 | 35.08 | 4000 | 0.0205 |
| I12935 | 633 | Europe | United Kingdom | 4467.2 | -1.69 | 51.70 | 4000 | 0.0187 |
| OBKR_80 | 1026 | Europe | Germany | 3480.3 | 10.88 | 48.27 | 4000 | 0.0193 |
| ROU001 | 1131 | Europe | Czech Republic | 3537.1 | 14.25 | 50.42 | 4000 | 0.0156 |
| ROU006 | 1131 | Europe | Czech Republic | 3537.1 | 14.25 | 50.42 | 4000 | 0.0184 |
| ROU007 | 1131 | Europe | Czech Republic | 3537.1 | 14.25 | 50.42 | 4000 | 0.0243 |
| VLI039 | 1253 | Europe | Czech Republic | 3537.1 | 14.45 | 50.37 | 4000 | 0.0250 |

| | | | | | | | | |
|---|---|---|---|---|---|---|---|---|
| VLI061 | 1253 | Europe | Czech Republic | 3537.1 | 14.45 | 50.37 | 4000 | 0.0210 |
| BNL002 | 486 | Europe | Czech Republic | 3495.2 | 14.65 | 50.17 | 3995 | 0.0205 |
| BNL003 | 486 | Europe | Czech Republic | 3495.2 | 14.65 | 50.17 | 3995 | 0.0169 |
| BNL004 | 486 | Europe | Czech Republic | 3495.2 | 14.65 | 50.17 | 3995 | 0.0186 |
| ROU003 | 1131 | Europe | Czech Republic | 3537.1 | 14.25 | 50.42 | 3995 | 0.0156 |
| ROU004 | 1131 | Europe | Czech Republic | 3537.1 | 14.25 | 50.42 | 3995 | 0.0116 |
| I3255 | 607 | Europe | United Kingdom | 4389.8 | 0.11 | 52.17 | 3994 | 0.0199 |
| HJTM107_noUDG | 122 | Asia | China | 7746.6 | 114.02 | 33.59 | 3993 | 0.0203 |
| I2618 | 812 | Europe | United Kingdom | 4717.0 | -2.15 | 55.06 | 3993 | 0.0267 |
| I4073 | 856 | Europe | Netherlands | 4151.8 | 5.10 | 52.73 | 3992 | 0.0209 |
| I3875 | 586 | Europe | France | 3649.9 | 6.37 | 44.48 | 3985 | 0.0255 |
| I3743 | 587 | Europe | Italy | 3620.3 | 9.23 | 39.87 | 3985 | 0.0188 |
| I6561 | 912 | Europe | Ukraine | 2964.0 | 37.70 | 49.54 | 3985 | 0.0190 |
| KO7001 | 973 | Europe | Czech Republic | 3449.6 | 15.22 | 50.01 | 3985 | 0.0167 |
| BAS025 | 479 | Europe | Spain | 4728.9 | -1.56 | 37.76 | 3981 | 0.0172 |
| I0116 | 564 | Europe | Germany | 3710.4 | 11.68 | 51.42 | 3978 | 0.0248 |
| KNE007 | 971 | Europe | Czech Republic | 3495.2 | 14.27 | 50.12 | 3977 | 0.0152 |
| I14543 | 663 | Europe | United Kingdom | 4296.4 | -0.13 | 50.85 | 3975 | 0.0217 |
| I16538 | 699 | Europe | Czech Republic | 3582.7 | 13.42 | 50.40 | 3975 | 0.0237 |
| I2461 | 794 | Europe | United Kingdom | 4423.8 | -1.69 | 51.13 | 3975 | 0.0262 |
| RISE683_noUDG.SG | 368 | Asia | Russia | 5581.2 | 90.36 | 53.71 | 3973 | 0.0213 |
| KNE003 | 971 | Europe | Czech Republic | 3495.2 | 14.27 | 50.12 | 3971 | 0.0259 |
| I2600 | 794 | Europe | United Kingdom | 4423.8 | -1.69 | 51.13 | 3970 | 0.0157 |
| I3770 | 222 | Asia | Kazakhstan | 4924.4 | 81.84 | 50.22 | 3968 | 0.0246 |
| I16813 | 709 | Europe | Serbia | 2823.1 | 20.17 | 45.89 | 3966 | 0.0269 |
| I4069 | 856 | Europe | Netherlands | 4151.8 | 5.10 | 52.73 | 3966 | 0.0168 |
| SUC003 | 1093 | Europe | Italy | 3570.8 | 8.44 | 40.81 | 3963 | 0.0185 |
| POST_99 | 1052 | Europe | Germany | 3480.3 | 10.89 | 48.30 | 3962 | 0.0242 |
| BAS023 | 479 | Europe | Spain | 4728.9 | -1.56 | 37.76 | 3961 | 0.0212 |
| RISE670_noUDG.SG | 361 | Asia | Russia | 5568.9 | 90.21 | 53.16 | 3959 | 0.0230 |
| I10558 | 595 | Europe | Romania | 2451.2 | 26.68 | 45.23 | 3958 | 0.0158 |

| | | | | | | | | |
|---|---|---|---|---|---|---|---|---|
| I13025 | 637 | Europe | Netherlands | 4077.7 | 4.86 | 51.89 | 3958 | 0.0153 |
| ZPL001 | 422 | Asia | Russia | 6592.3 | 105.17 | 54.39 | 3951 | 0.0246 |
| I10420 | 593 | Europe | Moldova | 2410.9 | 28.19 | 45.85 | 3950 | 0.0161 |
| I10479 | 595 | Europe | Romania | 2451.2 | 26.68 | 45.23 | 3950 | 0.0284 |
| I13468 | 641 | Europe | Czech Republic | 3495.2 | 14.34 | 50.04 | 3950 | 0.0219 |
| I13471 | 641 | Europe | Czech Republic | 3495.2 | 14.34 | 50.04 | 3950 | 0.0273 |
| I15643 | 641 | Europe | Czech Republic | 3495.2 | 14.34 | 50.04 | 3950 | 0.0147 |
| I15644 | 641 | Europe | Czech Republic | 3495.2 | 14.34 | 50.04 | 3950 | 0.0278 |
| I15646 | 641 | Europe | Czech Republic | 3495.2 | 14.34 | 50.04 | 3950 | 0.0261 |
| I16109 | 641 | Europe | Czech Republic | 3495.2 | 14.34 | 50.04 | 3950 | 0.0204 |
| I7958 | 641 | Europe | Czech Republic | 3495.2 | 14.34 | 50.04 | 3950 | 0.0200 |
| I7959 | 641 | Europe | Czech Republic | 3495.2 | 14.34 | 50.04 | 3950 | 0.0162 |
| I7960 | 641 | Europe | Czech Republic | 3495.2 | 14.34 | 50.04 | 3950 | 0.0214 |
| I7962 | 641 | Europe | Czech Republic | 3495.2 | 14.34 | 50.04 | 3950 | 0.0206 |
| I7963 | 641 | Europe | Czech Republic | 3495.2 | 14.34 | 50.04 | 3950 | 0.0204 |
| I7964 | 641 | Europe | Czech Republic | 3495.2 | 14.34 | 50.04 | 3950 | 0.0229 |
| I4075 | 856 | Europe | Netherlands | 4151.8 | 5.10 | 52.73 | 3950 | 0.0251 |
| ALM019_merged | 461 | Europe | Spain | 4630.4 | -1.51 | 37.95 | 3946 | 0.0239 |
| I14813 | 186 | Asia | Armenia | 1927.6 | 43.95 | 40.78 | 3945 | 0.0169 |
| KPT003 | 304 | Asia | Russia | 6584.6 | 105.51 | 53.04 | 3945 | 0.0227 |
| NV3001 | 346 | Asia | Russia | 2260.5 | 41.94 | 44.73 | 3945 | 0.0227 |
| I13469 | 641 | Europe | Czech Republic | 3495.2 | 14.34 | 50.04 | 3945 | 0.0221 |
| I11527 | 157 | Asia | Kyrgyzstan | 4314.5 | 75.89 | 41.43 | 3943 | 0.0229 |
| RISE392_noUDG.SG | 355 | Asia | Russia | 3905.0 | 59.08 | 53.88 | 3943 | 0.0241 |
| I13470 | 641 | Europe | Czech Republic | 3495.2 | 14.34 | 50.04 | 3943 | 0.0250 |
| VLI045 | 1253 | Europe | Czech Republic | 3537.1 | 14.45 | 50.37 | 3943 | 0.0178 |
| I2462 | 643 | Europe | United Kingdom | 4257.1 | 1.34 | 51.36 | 3942 | 0.0227 |
| I2447 | 752 | Europe | United Kingdom | 4397.4 | -1.31 | 51.80 | 3938 | 0.0242 |
| CHL003 | 505 | Europe | Czech Republic | 3449.6 | 15.12 | 50.22 | 3937 | 0.0223 |
| OBKR_50 | 1026 | Europe | Germany | 3480.3 | 10.88 | 48.27 | 3935 | 0.0279 |
| PLTM313_noUDG | 349 | Asia | China | 7833.7 | 114.92 | 33.71 | 3934 | 0.0213 |

| | | | | | | | | |
|---|---|---|---|---|---|---|---|---|
| POST_35 | 1052 | Europe | Germany | 3480.3 | 10.89 | 48.30 | 3932 | 0.0218 |
| I10480 | 595 | Europe | Romania | 2451.2 | 26.68 | 45.23 | 3931 | 0.0169 |
| I5737 | 905 | Europe | Turkey | 1699.7 | 27.14 | 38.42 | 3931 | 0.0205 |
| MX280 | 1009 | Europe | Germany | 3550.6 | 8.87 | 47.77 | 3931 | 0.0122 |
| BNL008 | 486 | Europe | Czech Republic | 3495.2 | 14.65 | 50.17 | 3926 | 0.0222 |
| I7414 | 232 | Asia | Uzbekistan | 3504.5 | 66.83 | 37.42 | 3925 | 0.0226 |
| I0047 | 563 | Europe | Germany | 3796.2 | 11.05 | 51.90 | 3923 | 0.0228 |
| KO1014 | 972 | Europe | Czech Republic | 3449.6 | 15.17 | 50.03 | 3915 | 0.0218 |
| KO1013 | 972 | Europe | Czech Republic | 3449.6 | 15.17 | 50.03 | 3914 | 0.0174 |
| MIS001 | 1000 | Europe | Czech Republic | 3404.5 | 15.78 | 49.99 | 3908 | 0.0262 |
| I0164 | 570 | Europe | Germany | 3796.2 | 11.14 | 51.79 | 3906 | 0.0204 |
| CHL001 | 505 | Europe | Czech Republic | 3449.6 | 15.12 | 50.22 | 3905 | 0.0232 |
| OBKR_9A | 1026 | Europe | Germany | 3480.3 | 10.88 | 48.27 | 3902 | 0.0195 |
| I4161 | 227 | Asia | Uzbekistan | 3507.3 | 67.00 | 37.75 | 3900 | 0.0180 |
| I4901 | 227 | Asia | Uzbekistan | 3507.3 | 67.00 | 37.75 | 3900 | 0.0198 |
| I5608 | 227 | Asia | Uzbekistan | 3507.3 | 67.00 | 37.75 | 3900 | 0.0276 |
| L5209 | 313 | Asia | China | 5344.7 | 88.67 | 40.34 | 3900 | 0.0263 |
| L5213 | 313 | Asia | China | 5344.7 | 88.67 | 40.34 | 3900 | 0.0231 |
| L6105 | 313 | Asia | China | 5344.7 | 88.67 | 40.34 | 3900 | 0.0256 |
| LJM14_noUDG | 318 | Asia | China | 6588.7 | 101.80 | 36.51 | 3900 | 0.0317 |
| LJM2_noUDG | 318 | Asia | China | 6588.7 | 101.80 | 36.51 | 3900 | 0.0239 |
| LJM5_noUDG | 318 | Asia | China | 6588.7 | 101.80 | 36.51 | 3900 | 0.0276 |
| WD-WT5M2_noUDG | 412 | Asia | China | 7727.7 | 113.37 | 34.18 | 3900 | 0.0276 |
| I10412 | 593 | Europe | Moldova | 2410.9 | 28.19 | 45.85 | 3900 | 0.0218 |
| I10436 | 593 | Europe | Moldova | 2410.9 | 28.19 | 45.85 | 3900 | 0.0196 |
| I15824 | 609 | Europe | Czech Republic | 3495.2 | 14.36 | 50.05 | 3900 | 0.0202 |
| I23205 | 707 | Europe | Serbia | 2823.1 | 20.40 | 45.93 | 3900 | 0.0244 |
| I23206 | 707 | Europe | Serbia | 2823.1 | 20.40 | 45.93 | 3900 | 0.0300 |
| I23207 | 707 | Europe | Serbia | 2823.1 | 20.40 | 45.93 | 3900 | 0.0227 |
| I23208 | 707 | Europe | Serbia | 2823.1 | 20.40 | 45.93 | 3900 | 0.0182 |
| I23212 | 707 | Europe | Serbia | 2823.1 | 20.40 | 45.93 | 3900 | 0.0231 |

| | | | | | | | | |
|---|---|---|---|---|---|---|---|---|
| MOK10B.SG | 707 | Europe | Serbia | 2823.1 | 20.40 | 45.93 | 3900 | 0.0261 |
| MOK12.SG | 707 | Europe | Serbia | 2823.1 | 20.40 | 45.93 | 3900 | 0.0212 |
| MOK13.SG | 707 | Europe | Serbia | 2823.1 | 20.40 | 45.93 | 3900 | 0.0269 |
| MOK14.SG | 707 | Europe | Serbia | 2823.1 | 20.40 | 45.93 | 3900 | 0.0207 |
| MOK15.SG | 707 | Europe | Serbia | 2823.1 | 20.40 | 45.93 | 3900 | 0.0221 |
| MOK16A.SG | 707 | Europe | Serbia | 2823.1 | 20.40 | 45.93 | 3900 | 0.0250 |
| MOK17A.SG | 707 | Europe | Serbia | 2823.1 | 20.40 | 45.93 | 3900 | 0.0206 |
| MOK19A.SG | 707 | Europe | Serbia | 2823.1 | 20.40 | 45.93 | 3900 | 0.0221 |
| MOK20.SG | 707 | Europe | Serbia | 2823.1 | 20.40 | 45.93 | 3900 | 0.0188 |
| MOK22.SG | 707 | Europe | Serbia | 2823.1 | 20.40 | 45.93 | 3900 | 0.0258 |
| MOK23.SG | 707 | Europe | Serbia | 2823.1 | 20.40 | 45.93 | 3900 | 0.0268 |
| MOK24A.SG | 707 | Europe | Serbia | 2823.1 | 20.40 | 45.93 | 3900 | 0.0244 |
| MOK25A.SG | 707 | Europe | Serbia | 2823.1 | 20.40 | 45.93 | 3900 | 0.0228 |
| MOK26A.SG | 707 | Europe | Serbia | 2823.1 | 20.40 | 45.93 | 3900 | 0.0203 |
| MOK27.SG | 707 | Europe | Serbia | 2823.1 | 20.40 | 45.93 | 3900 | 0.0205 |
| MOK28A.SG | 707 | Europe | Serbia | 2823.1 | 20.40 | 45.93 | 3900 | 0.0228 |
| MOK29A.SG | 707 | Europe | Serbia | 2823.1 | 20.40 | 45.93 | 3900 | 0.0234 |
| MOK30.SG | 707 | Europe | Serbia | 2823.1 | 20.40 | 45.93 | 3900 | 0.0260 |
| MOK31.SG | 707 | Europe | Serbia | 2823.1 | 20.40 | 45.93 | 3900 | 0.0133 |
| MOK32.SG | 707 | Europe | Serbia | 2823.1 | 20.40 | 45.93 | 3900 | 0.0169 |
| MOK33.SG | 707 | Europe | Serbia | 2823.1 | 20.40 | 45.93 | 3900 | 0.0220 |
| MOK9B.SG | 707 | Europe | Serbia | 2823.1 | 20.40 | 45.93 | 3900 | 0.0205 |
| I17912 | 709 | Europe | Serbia | 2823.1 | 20.17 | 45.89 | 3900 | 0.0215 |
| I17913 | 709 | Europe | Serbia | 2823.1 | 20.17 | 45.89 | 3900 | 0.0224 |
| I23209 | 709 | Europe | Serbia | 2823.1 | 20.17 | 45.89 | 3900 | 0.0198 |
| I23210 | 709 | Europe | Serbia | 2823.1 | 20.17 | 45.89 | 3900 | 0.0191 |
| I23211 | 709 | Europe | Serbia | 2823.1 | 20.17 | 45.89 | 3900 | 0.0179 |
| MX254 | 1009 | Europe | Germany | 3550.6 | 8.87 | 47.77 | 3900 | 0.0185 |
| RISE97_noUDG.SG | 1129 | Europe | Sweden | 4060.0 | 13.06 | 55.56 | 3895 | 0.0198 |
| I16803 | 707 | Europe | Serbia | 2823.1 | 20.40 | 45.93 | 3894 | 0.0240 |
| KO1009 | 972 | Europe | Czech Republic | 3449.6 | 15.17 | 50.03 | 3894 | 0.0179 |

| | | | | | | | | |
|---|---|---|---|---|---|---|---|---|
| I0418 | 134 | Asia | Russia | 3487.5 | 50.99 | 52.91 | 3884 | 0.0242 |
| MIS002 | 1000 | Europe | Czech Republic | 3404.5 | 15.78 | 49.99 | 3883 | 0.0186 |
| Spiginas2 | 1141 | Europe | Lithuania | 3744.2 | 22.42 | 55.77 | 3880 | 0.0182 |
| I7494 | 232 | Asia | Uzbekistan | 3504.5 | 66.83 | 37.42 | 3875 | 0.0237 |
| WEHR_1415adult | 1260 | Europe | Germany | 3480.3 | 10.81 | 48.25 | 3872 | 0.0264 |
| I0984 | 146 | Asia | Russia | 3914.5 | 60.47 | 52.82 | 3870 | 0.0233 |
| KO1006 | 972 | Europe | Czech Republic | 3449.6 | 15.17 | 50.03 | 3868 | 0.0238 |
| R22.SG | 1093 | Europe | Italy | 3570.8 | 8.44 | 40.81 | 3868 | 0.0214 |
| TORTF.SG | 541 | Europe | France | 3895.7 | 3.06 | 43.15 | 3861 | 0.0264 |
| I3756 | 850 | Europe | Spain | 4679.9 | -2.84 | 38.58 | 3860 | 0.0200 |
| MIB028 | 1000 | Europe | Czech Republic | 3404.5 | 15.78 | 49.99 | 3855 | 0.0187 |
| I7492 | 232 | Asia | Uzbekistan | 3504.5 | 66.83 | 37.42 | 3853 | 0.0185 |
| I7495 | 232 | Asia | Uzbekistan | 3504.5 | 66.83 | 37.42 | 3853 | 0.0140 |
| ETM014 | 423 | Both | Syria | 1028.7 | 36.80 | 35.80 | 3850 | 0.0166 |
| SZ1.SG | 475 | Europe | Hungary | 2919.0 | 17.85 | 46.28 | 3850 | 0.0222 |
| I11160 | 609 | Europe | Czech Republic | 3495.2 | 14.36 | 50.05 | 3850 | 0.0217 |
| I14185 | 609 | Europe | Czech Republic | 3495.2 | 14.36 | 50.05 | 3850 | 0.0191 |
| I14186 | 609 | Europe | Czech Republic | 3495.2 | 14.36 | 50.05 | 3850 | 0.0215 |
| I14189 | 609 | Europe | Czech Republic | 3495.2 | 14.36 | 50.05 | 3850 | 0.0262 |
| I14191 | 609 | Europe | Czech Republic | 3495.2 | 14.36 | 50.05 | 3850 | 0.0248 |
| I14192 | 609 | Europe | Czech Republic | 3495.2 | 14.36 | 50.05 | 3850 | 0.0220 |
| I14193 | 609 | Europe | Czech Republic | 3495.2 | 14.36 | 50.05 | 3850 | 0.0198 |
| I14585 | 609 | Europe | Czech Republic | 3495.2 | 14.36 | 50.05 | 3850 | 0.0236 |
| N17_noUDG.SG | 1010 | Europe | Poland | 3539.2 | 18.88 | 52.66 | 3850 | 0.0222 |
| I1020 | 146 | Asia | Russia | 3914.5 | 60.47 | 52.82 | 3841 | 0.0257 |
| KDC001 | 291 | Asia | Russia | 2206.2 | 43.72 | 43.35 | 3838 | 0.0222 |
| WEHR_1586 | 1260 | Europe | Germany | 3480.3 | 10.81 | 48.25 | 3835 | 0.0257 |
| I1027 | 146 | Asia | Russia | 3914.5 | 60.47 | 52.82 | 3833 | 0.0303 |
| I1029 | 146 | Asia | Russia | 3914.5 | 60.47 | 52.82 | 3830 | 0.0257 |
| I0943 | 146 | Asia | Russia | 3914.5 | 60.47 | 52.82 | 3825 | 0.0251 |
| I1011 | 146 | Asia | Russia | 3914.5 | 60.47 | 52.82 | 3825 | 0.0244 |

| | | | | | | | | |
|---|---|---|---|---|---|---|---|---|
| I1089 | 146 | Asia | Russia | 3914.5 | 60.47 | 52.82 | 3825 | 0.0296 |
| I7480 | 146 | Asia | Russia | 3914.5 | 60.47 | 52.82 | 3825 | 0.0170 |
| ALM003 | 461 | Europe | Spain | 4630.4 | -1.51 | 37.95 | 3825 | 0.0259 |
| ALM008 | 461 | Europe | Spain | 4630.4 | -1.51 | 37.95 | 3825 | 0.0197 |
| ALM020 | 461 | Europe | Spain | 4630.4 | -1.51 | 37.95 | 3825 | 0.0240 |
| ALM021 | 461 | Europe | Spain | 4630.4 | -1.51 | 37.95 | 3825 | 0.0151 |
| ALM028_merged | 461 | Europe | Spain | 4630.4 | -1.51 | 37.95 | 3825 | 0.0230 |
| ALM036 | 461 | Europe | Spain | 4630.4 | -1.51 | 37.95 | 3825 | 0.0217 |
| ALM042 | 461 | Europe | Spain | 4630.4 | -1.51 | 37.95 | 3825 | 0.0166 |
| ALM050 | 461 | Europe | Spain | 4630.4 | -1.51 | 37.95 | 3825 | 0.0286 |
| ALM051 | 461 | Europe | Spain | 4630.4 | -1.51 | 37.95 | 3825 | 0.0207 |
| ALM058 | 461 | Europe | Spain | 4630.4 | -1.51 | 37.95 | 3825 | 0.0224 |
| ALM064 | 461 | Europe | Spain | 4630.4 | -1.51 | 37.95 | 3825 | 0.0167 |
| ALM076 | 461 | Europe | Spain | 4630.4 | -1.51 | 37.95 | 3825 | 0.0199 |
| ALM081 | 461 | Europe | Spain | 4630.4 | -1.51 | 37.95 | 3825 | 0.0152 |
| MA2205_noUDG.SG | 994 | Europe | Turkey | 1471.3 | 33.79 | 39.35 | 3825 | 0.0169 |
| ZAP002 | 1261 | Europe | Spain | 4728.9 | -1.70 | 37.67 | 3825 | 0.0175 |
| I12955 | 169 | Asia | Mongolia | 6010.8 | 96.80 | 49.70 | 3824 | 0.0249 |
| I7416 | 232 | Asia | Uzbekistan | 3504.5 | 66.83 | 37.42 | 3824 | 0.0185 |
| I3124 | 611 | Europe | Italy | 3057.3 | 13.48 | 37.91 | 3824 | 0.0190 |
| VERT113B | 1177 | Europe | Ukraine | 2837.9 | 25.85 | 48.78 | 3823 | 0.0215 |
| ALM034 | 461 | Europe | Spain | 4630.4 | -1.51 | 37.95 | 3822 | 0.0236 |
| RISE395_noUDG.SG | 356 | Asia | Russia | 3869.4 | 59.54 | 52.64 | 3815 | 0.0265 |
| I1057 | 146 | Asia | Russia | 3914.5 | 60.47 | 52.82 | 3814 | 0.0212 |
| ALM041 | 461 | Europe | Spain | 4630.4 | -1.51 | 37.95 | 3814 | 0.0214 |
| I14553 | 665 | Europe | United Kingdom | 4296.4 | -0.09 | 50.82 | 3807 | 0.0199 |
| ALM016_merged | 461 | Europe | Spain | 4630.4 | -1.51 | 37.95 | 3803 | 0.0160 |
| ALM002_merged | 461 | Europe | Spain | 4630.4 | -1.51 | 37.95 | 3800 | 0.0236 |
| yak022_noUDG.SG | 416 | Asia | Russia | 8918.7 | 152.63 | 66.80 | 3799 | 0.0263 |
| VLI050 | 1253 | Europe | Czech Republic | 3537.1 | 14.45 | 50.37 | 3796 | 0.0158 |
| I0354 | 130 | Asia | Russia | 3531.9 | 50.39 | 53.03 | 3791 | 0.0160 |

| | | | | | | | | |
|---|---|---|---|---|---|---|---|---|
| I4289 | 232 | Asia | Uzbekistan | 3504.5 | 66.83 | 37.42 | 3790 | 0.0238 |
| I7421 | 232 | Asia | Uzbekistan | 3504.5 | 66.83 | 37.42 | 3790 | 0.0193 |
| ALM086 | 461 | Europe | Spain | 4630.4 | -1.51 | 37.95 | 3788 | 0.0203 |
| R11102.SG | 1078 | Europe | Italy | 3097.9 | 11.95 | 42.03 | 3787 | 0.0198 |
| I1053 | 146 | Asia | Russia | 3914.5 | 60.47 | 52.82 | 3785 | 0.0249 |
| I3494 | 845 | Europe | Spain | 4531.7 | -0.87 | 38.85 | 3781 | 0.0243 |
| ALA095 | 459 | Europe | Turkey | 1077.6 | 36.38 | 36.24 | 3780 | 0.0177 |
| ALM006 | 461 | Europe | Spain | 4630.4 | -1.51 | 37.95 | 3780 | 0.0225 |
| XHM135.SG | 313 | Asia | China | 5344.7 | 88.67 | 40.34 | 3778 | 0.0232 |
| I12776 | 631 | Europe | United Kingdom | 4557.3 | -1.64 | 53.08 | 3778 | 0.0292 |
| AITI_72 | 455 | Europe | Germany | 3480.3 | 10.84 | 48.22 | 3776 | 0.0233 |
| I4341_noUDG | 228 | Asia | Iran | 1677.7 | 45.17 | 37.01 | 3775 | 0.0205 |
| LON001 | 956 | Europe | Italy | 3620.3 | 9.24 | 39.87 | 3775 | 0.0207 |
| KPT004 | 304 | Asia | Russia | 6584.6 | 105.51 | 53.04 | 3773 | 0.0182 |
| ALA004 | 459 | Europe | Turkey | 1077.6 | 36.38 | 36.24 | 3773 | 0.0133 |
| ALM015_merged | 461 | Europe | Spain | 4630.4 | -1.51 | 37.95 | 3773 | 0.0208 |
| I18606 | 736 | Europe | United Kingdom | 4552.7 | -0.28 | 54.10 | 3770 | 0.0196 |
| I1058 | 146 | Asia | Russia | 3914.5 | 60.47 | 52.82 | 3769 | 0.0261 |
| I4286 | 232 | Asia | Uzbekistan | 3504.5 | 66.83 | 37.42 | 3768 | 0.0198 |
| I1054 | 146 | Asia | Russia | 3914.5 | 60.47 | 52.82 | 3765 | 0.0268 |
| MIB002 | 1000 | Europe | Czech Republic | 3404.5 | 15.78 | 49.99 | 3765 | 0.0194 |
| I2602 | 643 | Europe | United Kingdom | 4257.1 | 1.34 | 51.36 | 3762 | 0.0158 |
| TU911_SX23 | 1167 | Europe | Switzerland | 3514.3 | 9.29 | 47.45 | 3762 | 0.0165 |
| I6796 | 262 | Asia | Kazakhstan | 4386.8 | 74.44 | 50.09 | 3761 | 0.0195 |
| AITI_66 | 455 | Europe | Germany | 3480.3 | 10.84 | 48.22 | 3761 | 0.0229 |
| PUC003 | 507 | Europe | Spain | 4581.0 | -0.86 | 38.63 | 3761 | 0.0207 |
| AITI_43 | 455 | Europe | Germany | 3480.3 | 10.84 | 48.22 | 3760 | 0.0210 |
| KPT001 | 304 | Asia | Russia | 6584.6 | 105.51 | 53.04 | 3759 | 0.0194 |
| I4884 | 870 | Europe | Czech Republic | 3495.2 | 14.46 | 50.12 | 3758 | 0.0258 |
| I5377 | 889 | Europe | United Kingdom | 4311.6 | -0.27 | 51.47 | 3758 | 0.0194 |
| mur004_noUDG.SG | 328 | Asia | Russia | 3737.3 | 56.85 | 52.58 | 3755 | 0.0239 |

| | | | | | | | | |
|---|---|---|---|---|---|---|---|---|
| I7419 | 232 | Asia | Uzbekistan | 3504.5 | 66.83 | 37.42 | 3754 | 0.0183 |
| kzb008_noUDG.SG | 310 | Asia | Russia | 3819.3 | 55.54 | 54.07 | 3754 | 0.0313 |
| I4892 | 870 | Europe | Czech Republic | 3495.2 | 14.46 | 50.12 | 3754 | 0.0171 |
| AITI_2 | 455 | Europe | Germany | 3480.3 | 10.84 | 48.22 | 3753 | 0.0222 |
| ALA008 | 459 | Europe | Turkey | 1077.6 | 36.38 | 36.24 | 3752 | 0.0195 |
| ALM017 | 461 | Europe | Spain | 4630.4 | -1.51 | 37.95 | 3752 | 0.0208 |
| EDM176_noUDG | 111 | Asia | China | 7815.5 | 118.95 | 42.10 | 3750 | 0.0247 |
| I7420 | 232 | Asia | Uzbekistan | 3504.5 | 66.83 | 37.42 | 3750 | 0.0211 |
| KDC002 | 291 | Asia | Russia | 2206.2 | 43.72 | 43.35 | 3750 | 0.0211 |
| I10093 | 426 | Both | Israel | 629.1 | 35.02 | 32.58 | 3750 | 0.0162 |
| I18745 | 739 | Europe | Croatia | 2823.8 | 16.43 | 43.97 | 3750 | 0.0215 |
| I18746 | 739 | Europe | Croatia | 2823.8 | 16.43 | 43.97 | 3750 | 0.0185 |
| I18752 | 739 | Europe | Croatia | 2823.8 | 16.43 | 43.97 | 3750 | 0.0151 |
| AITI_78 | 455 | Europe | Germany | 3480.3 | 10.84 | 48.22 | 3748 | 0.0158 |
| I4420_noUDG | 864 | Europe | Spain | 4583.5 | 1.42 | 38.69 | 3748 | 0.0158 |
| AITI_119 | 455 | Europe | Germany | 3480.3 | 10.84 | 48.22 | 3747 | 0.0143 |
| I5604 | 153 | Asia | Uzbekistan | 3507.3 | 67.00 | 37.67 | 3745 | 0.0126 |
| RISE150_noUDG.SG | 1125 | Europe | Poland | 3446.8 | 16.96 | 50.92 | 3744 | 0.0177 |
| I4262 | 231 | Asia | Kazakhstan | 4445.0 | 76.45 | 48.78 | 3743 | 0.0226 |
| ALA029 | 459 | Europe | Turkey | 1077.6 | 36.38 | 36.24 | 3743 | 0.0161 |
| PUC001 | 507 | Europe | Spain | 4581.0 | -0.86 | 38.63 | 3741 | 0.0200 |
| BOO001.A0101 | 489 | Europe | Russia | 5111.7 | 33.48 | 69.22 | 3740 | 0.0243 |
| I1064 | 146 | Asia | Russia | 3914.5 | 60.47 | 52.82 | 3739 | 0.0241 |
| KO1007 | 972 | Europe | Czech Republic | 3449.6 | 15.17 | 50.03 | 3737 | 0.0192 |
| ISB001 | 956 | Europe | Italy | 3620.3 | 9.24 | 39.87 | 3735 | 0.0223 |
| ALA013 | 459 | Europe | Turkey | 1077.6 | 36.38 | 36.24 | 3734 | 0.0184 |
| I6789 | 262 | Asia | Kazakhstan | 4386.8 | 74.44 | 50.09 | 3732 | 0.0204 |
| I10343 | 586 | Europe | France | 3649.9 | 6.37 | 44.48 | 3732 | 0.0158 |
| AITI_65adult | 455 | Europe | Germany | 3480.3 | 10.84 | 48.22 | 3728 | 0.0212 |
| I10342 | 586 | Europe | France | 3649.9 | 6.37 | 44.48 | 3728 | 0.0185 |
| ALM035 | 461 | Europe | Spain | 4630.4 | -1.51 | 37.95 | 3725 | 0.0125 |

| | | | | | | | | |
|---|---|---|---|---|---|---|---|---|
| BOO002.A0101 | 489 | Europe | Russia | 5111.7 | 33.48 | 69.22 | 3725 | 0.0225 |
| BOO003.A0101 | 489 | Europe | Russia | 5111.7 | 33.48 | 69.22 | 3725 | 0.0161 |
| BOO004.A0101 | 489 | Europe | Russia | 5111.7 | 33.48 | 69.22 | 3725 | 0.0255 |
| BOO005.A0101 | 489 | Europe | Russia | 5111.7 | 33.48 | 69.22 | 3725 | 0.0261 |
| BOO006.A0101 | 489 | Europe | Russia | 5111.7 | 33.48 | 69.22 | 3725 | 0.0206 |
| I6792 | 262 | Asia | Kazakhstan | 4386.8 | 74.44 | 50.09 | 3724 | 0.0206 |
| I13173 | 180 | Asia | Mongolia | 5630.5 | 91.84 | 47.05 | 3722 | 0.0266 |
| mur003_noUDG.SG | 328 | Asia | Russia | 3737.3 | 56.85 | 52.58 | 3721 | 0.0201 |
| R11104.SG | 1078 | Europe | Italy | 3097.9 | 11.95 | 42.03 | 3719 | 0.0195 |
| R11107.SG | 1078 | Europe | Italy | 3097.9 | 11.95 | 42.03 | 3719 | 0.0216 |
| I4071 | 856 | Europe | Netherlands | 4151.8 | 5.10 | 52.73 | 3717 | 0.0219 |
| I6794 | 262 | Asia | Kazakhstan | 4386.8 | 74.44 | 50.09 | 3715 | 0.0233 |
| I6799 | 263 | Asia | Kazakhstan | 4459.0 | 75.81 | 49.12 | 3715 | 0.0214 |
| JAG34_noUDG.SG | 959 | Europe | Croatia | 2878.2 | 18.51 | 45.69 | 3715 | 0.0147 |
| ALM060 | 461 | Europe | Spain | 4630.4 | -1.51 | 37.95 | 3713 | 0.0267 |
| ALA025 | 459 | Europe | Turkey | 1077.6 | 36.38 | 36.24 | 3711 | 0.0159 |
| I2981 | 833 | Europe | United Kingdom | 5265.3 | -2.42 | 59.28 | 3711 | 0.0264 |
| ALA028 | 459 | Europe | Turkey | 1077.6 | 36.38 | 36.24 | 3705 | 0.0161 |
| I1028 | 146 | Asia | Russia | 3914.5 | 60.47 | 52.82 | 3704 | 0.0284 |
| kzb002.SG | 310 | Asia | Russia | 3819.3 | 55.54 | 54.07 | 3704 | 0.0194 |
| I4070 | 856 | Europe | Netherlands | 4151.8 | 5.10 | 52.73 | 3703 | 0.0187 |
| I14759 | 670 | Europe | Turkey | 1180.9 | 37.17 | 36.69 | 3700 | 0.0143 |
| I14782 | 670 | Europe | Turkey | 1180.9 | 37.17 | 36.69 | 3700 | 0.0115 |
| I18748 | 740 | Europe | Croatia | 2714.9 | 17.00 | 43.13 | 3700 | 0.0112 |
| RISE479_noUDG.SG | 1127 | Europe | Hungary | 3004.9 | 18.90 | 47.34 | 3700 | 0.0197 |
| ALA034 | 459 | Europe | Turkey | 1077.6 | 36.38 | 36.24 | 3697 | 0.0163 |
| I6790 | 262 | Asia | Kazakhstan | 4386.8 | 74.44 | 50.09 | 3695 | 0.0255 |
| I6797 | 262 | Asia | Kazakhstan | 4386.8 | 74.44 | 50.09 | 3695 | 0.0266 |
| I14784 | 670 | Europe | Turkey | 1180.9 | 37.17 | 36.69 | 3695 | 0.0194 |
| I11541 | 159 | Asia | Kazakhstan | 3531.0 | 57.04 | 50.45 | 3684 | 0.0133 |
| I3767 | 223 | Asia | Kazakhstan | 4116.1 | 72.01 | 47.78 | 3684 | 0.0192 |

| | | | | | | | | |
|---|---|---|---|---|---|---|---|---|
| I0424 | 136 | Asia | Russia | 3454.2 | 48.37 | 53.12 | 3675 | 0.0240 |
| I0430 | 137 | Asia | Russia | 3531.9 | 50.36 | 53.08 | 3675 | 0.0214 |
| L6103 | 313 | Asia | China | 5344.7 | 88.67 | 40.34 | 3675 | 0.0263 |
| I6791 | 262 | Asia | Kazakhstan | 4386.8 | 74.44 | 50.09 | 3674 | 0.0248 |
| kzb007_noUDG.SG | 310 | Asia | Russia | 3819.3 | 55.54 | 54.07 | 3672 | 0.0240 |
| AITI_95 | 455 | Europe | Germany | 3480.3 | 10.84 | 48.22 | 3671 | 0.0224 |
| I0507 | 142 | Asia | Kazakhstan | 4598.6 | 79.36 | 45.13 | 3668 | 0.0214 |
| SUC009 | 1093 | Europe | Italy | 3570.8 | 8.44 | 40.81 | 3667 | 0.0237 |
| AITI_120 | 455 | Europe | Germany | 3480.3 | 10.84 | 48.22 | 3662 | 0.0198 |
| MIB034 | 1000 | Europe | Czech Republic | 3404.5 | 15.78 | 49.99 | 3659 | 0.0172 |
| I14789 | 670 | Europe | Turkey | 1180.9 | 37.17 | 36.69 | 3658 | 0.0200 |
| I4318 | 233 | Asia | Kazakhstan | 4519.6 | 77.00 | 48.74 | 3656 | 0.0223 |
| I4776 | 233 | Asia | Kazakhstan | 4519.6 | 77.00 | 48.74 | 3656 | 0.0244 |
| I6800 | 264 | Asia | Kazakhstan | 4006.3 | 62.69 | 52.61 | 3656 | 0.0259 |
| I3965 | 430 | Both | Israel | 629.1 | 35.20 | 32.70 | 3650 | 0.0147 |
| I3966 | 430 | Both | Israel | 629.1 | 35.20 | 32.70 | 3650 | 0.0156 |
| JAG58_noUDG.SG | 959 | Europe | Croatia | 2878.2 | 18.51 | 45.69 | 3650 | 0.0232 |
| JAG78_noUDG.SG | 959 | Europe | Croatia | 2878.2 | 18.51 | 45.69 | 3650 | 0.0232 |
| JAG82_noUDG.SG | 959 | Europe | Croatia | 2878.2 | 18.51 | 45.69 | 3650 | 0.0219 |
| JAG85_noUDG.SG | 959 | Europe | Croatia | 2878.2 | 18.51 | 45.69 | 3650 | 0.0180 |
| JAG93_noUDG.SG | 959 | Europe | Croatia | 2878.2 | 18.51 | 45.69 | 3650 | 0.0243 |
| kzb006_noUDG.SG | 310 | Asia | Russia | 3819.3 | 55.54 | 54.07 | 3648 | 0.0201 |
| PUC004 | 507 | Europe | Spain | 4581.0 | -0.86 | 38.63 | 3644 | 0.0218 |
| I7412 | 227 | Asia | Uzbekistan | 3507.3 | 67.00 | 37.75 | 3642 | 0.0177 |
| I6795 | 262 | Asia | Kazakhstan | 4386.8 | 74.44 | 50.09 | 3642 | 0.0245 |
| I10348 | 586 | Europe | France | 3649.9 | 6.37 | 44.48 | 3642 | 0.0189 |
| I3997 | 854 | Europe | Spain | 4433.7 | -0.51 | 39.53 | 3642 | 0.0213 |
| VAD001 | 1172 | Europe | Spain | 4402.7 | -2.89 | 42.44 | 3641 | 0.0250 |
| I2163 | 771 | Europe | Bulgaria | 2168.7 | 25.50 | 42.13 | 3639 | 0.0151 |
| I6348 | 160 | Asia | Mongolia | 6379.3 | 102.70 | 49.39 | 3632 | 0.0217 |
| I4779 | 233 | Asia | Kazakhstan | 4519.6 | 77.00 | 48.74 | 3631 | 0.0184 |

| | | | | | | | | |
|---|---|---|---|---|---|---|---|---|
| I6793 | 262 | Asia | Kazakhstan | 4386.8 | 74.44 | 50.09 | 3631 | 0.0266 |
| I5080 | 882 | Europe | Croatia | 2904.0 | 16.05 | 44.92 | 3631 | 0.0220 |
| RISE505_noUDG.SG | 360 | Asia | Russia | 5246.1 | 85.45 | 53.46 | 3628 | 0.0266 |
| ALA026 | 459 | Europe | Turkey | 1077.6 | 36.38 | 36.24 | 3627 | 0.0150 |
| I4323 | 235 | Asia | Kazakhstan | 4497.6 | 77.81 | 43.25 | 3622 | 0.0182 |
| ALA011 | 459 | Europe | Turkey | 1077.6 | 36.38 | 36.24 | 3620 | 0.0144 |
| I11538 | 158 | Asia | Kazakhstan | 3577.2 | 57.72 | 50.67 | 3614 | 0.0273 |
| I8506 | 167 | Asia | Uzbekistan | 3363.1 | 64.99 | 40.54 | 3614 | 0.0238 |
| I4562 | 867 | Europe | Spain | 4188.1 | 1.06 | 41.29 | 3614 | 0.0235 |
| PUC002 | 507 | Europe | Spain | 4581.0 | -0.86 | 38.63 | 3610 | 0.0252 |
| R11105.SG | 1078 | Europe | Italy | 3097.9 | 11.95 | 42.03 | 3610 | 0.0211 |
| I4312 | 227 | Asia | Uzbekistan | 3507.3 | 67.00 | 37.75 | 3609 | 0.0149 |
| I4782 | 234 | Asia | Kazakhstan | 4705.1 | 79.99 | 46.95 | 3609 | 0.0213 |
| I6470 | 909 | Europe | Spain | 4484.7 | -3.84 | 42.73 | 3609 | 0.0166 |
| I3389 | 220 | Asia | Russia | 5558.3 | 89.31 | 55.19 | 3600 | 0.0211 |
| I3390 | 220 | Asia | Russia | 5558.3 | 89.31 | 55.19 | 3600 | 0.0271 |
| I3391 | 220 | Asia | Russia | 5558.3 | 89.31 | 55.19 | 3600 | 0.0269 |
| I3392 | 220 | Asia | Russia | 5558.3 | 89.31 | 55.19 | 3600 | 0.0196 |
| I3393 | 220 | Asia | Russia | 5558.3 | 89.31 | 55.19 | 3600 | 0.0225 |
| I3394 | 220 | Asia | Russia | 5558.3 | 89.31 | 55.19 | 3600 | 0.0210 |
| I3395 | 220 | Asia | Russia | 5558.3 | 89.31 | 55.19 | 3600 | 0.0263 |
| I3396 | 220 | Asia | Russia | 5558.3 | 89.31 | 55.19 | 3600 | 0.0223 |
| ALM053 | 461 | Europe | Spain | 4630.4 | -1.51 | 37.95 | 3600 | 0.0228 |
| ALM063 | 461 | Europe | Spain | 4630.4 | -1.51 | 37.95 | 3600 | 0.0198 |
| ALM067 | 461 | Europe | Spain | 4630.4 | -1.51 | 37.95 | 3600 | 0.0212 |
| ALM068 | 461 | Europe | Spain | 4630.4 | -1.51 | 37.95 | 3600 | 0.0239 |
| ALM069 | 461 | Europe | Spain | 4630.4 | -1.51 | 37.95 | 3600 | 0.0222 |
| ALM080 | 461 | Europe | Spain | 4630.4 | -1.51 | 37.95 | 3600 | 0.0240 |
| ALM084 | 461 | Europe | Spain | 4630.4 | -1.51 | 37.95 | 3600 | 0.0225 |
| ALM088 | 461 | Europe | Spain | 4630.4 | -1.51 | 37.95 | 3600 | 0.0279 |
| BAS022 | 479 | Europe | Spain | 4728.9 | -1.56 | 37.76 | 3600 | 0.0146 |

| | | | | | | | | |
|---|---|---|---|---|---|---|---|---|
| I18712 | 737 | Europe | Croatia | 2728.3 | 16.75 | 43.46 | 3600 | 0.0196 |
| GBVPO.SG | 541 | Europe | France | 3895.7 | 3.06 | 43.15 | 3598 | 0.0151 |
| I14762 | 670 | Europe | Turkey | 1180.9 | 37.17 | 36.69 | 3583 | 0.0160 |
| ALM039 | 461 | Europe | Spain | 4630.4 | -1.51 | 37.95 | 3580 | 0.0210 |
| TU907_SX20 | 1167 | Europe | Switzerland | 3514.3 | 9.29 | 47.45 | 3576 | 0.0261 |
| I10140 | 149 | Asia | Kazakhstan | 4705.1 | 80.03 | 46.97 | 3575 | 0.0190 |
| MA2200_noUDG.SG | 994 | Europe | Turkey | 1471.3 | 33.79 | 39.35 | 3575 | 0.0197 |
| MA2203_noUDG.SG | 994 | Europe | Turkey | 1471.3 | 33.79 | 39.35 | 3575 | 0.0175 |
| I5073 | 878 | Europe | Croatia | 2810.3 | 16.55 | 43.61 | 3573 | 0.0206 |
| I6716 | 220 | Asia | Russia | 5558.3 | 89.31 | 55.19 | 3571 | 0.0286 |
| I7570 | 938 | Europe | United Kingdom | 4405.4 | -0.34 | 52.70 | 3569 | 0.0159 |
| I4778 | 231 | Asia | Kazakhstan | 4445.0 | 76.45 | 48.78 | 3562 | 0.0192 |
| I4264 | 149 | Asia | Kazakhstan | 4705.1 | 80.03 | 46.97 | 3551 | 0.0186 |
| RISE500_noUDG.SG | 360 | Asia | Russia | 5246.1 | 85.45 | 53.46 | 3550 | 0.0328 |
| I4560 | 867 | Europe | Spain | 4188.1 | 1.06 | 41.29 | 3550 | 0.0238 |
| I4561 | 867 | Europe | Spain | 4188.1 | 1.06 | 41.29 | 3550 | 0.0179 |
| TV32032extra_noUDG.SG | 880 | Europe | Portugal | 5083.7 | -8.59 | 37.20 | 3550 | 0.0217 |
| TV3831_noUDG.SG | 880 | Europe | Portugal | 5083.7 | -8.59 | 37.20 | 3550 | 0.0256 |
| RISE503_noUDG.SG | 360 | Asia | Russia | 5246.1 | 85.45 | 53.46 | 3547 | 0.0367 |
| MonteGato104_noUDG.SG | 1002 | Europe | Portugal | 4932.9 | -7.60 | 37.94 | 3546 | 0.0209 |
| I7411 | 227 | Asia | Uzbekistan | 3507.3 | 67.00 | 37.75 | 3542 | 0.0162 |
| I1856 | 210 | Asia | Russia | 5648.2 | 91.46 | 53.87 | 3540 | 0.0261 |
| I14786 | 670 | Europe | Turkey | 1180.9 | 37.17 | 36.69 | 3540 | 0.0154 |
| I10347 | 586 | Europe | France | 3649.9 | 6.37 | 44.48 | 3527 | 0.0254 |
| I11027 | 153 | Asia | Uzbekistan | 3507.3 | 67.00 | 37.67 | 3526 | 0.0138 |
| LHO003 | 989 | Europe | Spain | 4581.0 | -0.77 | 38.34 | 3524 | 0.0236 |
| I4265 | 149 | Asia | Kazakhstan | 4705.1 | 80.03 | 46.97 | 3522 | 0.0164 |
| I4321 | 234 | Asia | Kazakhstan | 4705.1 | 79.99 | 46.95 | 3522 | 0.0155 |
| I7573 | 939 | Europe | United Kingdom | 4389.8 | 0.39 | 52.31 | 3520 | 0.0204 |
| I4331 | 863 | Europe | Croatia | 2714.9 | 17.34 | 43.19 | 3519 | 0.0225 |
| I4109 | 611 | Europe | Italy | 3057.3 | 13.48 | 37.91 | 3516 | 0.0164 |

| | | | | | | | | |
|---|---|---|---|---|---|---|---|---|
| BAS002 | 479 | Europe | Spain | 4728.9 | -1.56 | 37.76 | 3515 | 0.0231 |
| ALA019 | 459 | Europe | Turkey | 1077.6 | 36.38 | 36.24 | 3514 | 0.0217 |
| I4773 | 149 | Asia | Kazakhstan | 4705.1 | 80.03 | 46.97 | 3508 | 0.0224 |
| I4784 | 235 | Asia | Kazakhstan | 4497.6 | 77.81 | 43.25 | 3508 | 0.0150 |
| I4332 | 863 | Europe | Croatia | 2714.9 | 17.34 | 43.19 | 3508 | 0.0196 |
| OTTM_151ind2_noUDG | 1033 | Europe | Germany | 3480.3 | 10.85 | 48.23 | 3507 | 0.0233 |
| I4258 | 226 | Asia | Tajikistan | 3611.2 | 68.03 | 39.45 | 3504 | 0.0188 |
| I4774 | 149 | Asia | Kazakhstan | 4705.1 | 80.03 | 46.97 | 3503 | 0.0221 |
| yak024_noUDG.SG | 416 | Asia | Russia | 8918.7 | 152.63 | 66.80 | 3500 | 0.0203 |
| I10099 | 426 | Both | Israel | 629.1 | 35.02 | 32.58 | 3500 | 0.0042 |
| I2200 | 426 | Both | Israel | 629.1 | 35.02 | 32.58 | 3500 | 0.0124 |
| LHO002 | 989 | Europe | Spain | 4581.0 | -0.77 | 38.34 | 3500 | 0.0235 |
| I10112 | 148 | Asia | Kazakhstan | 4561.2 | 75.60 | 50.82 | 3497 | 0.0197 |
| I11520 | 153 | Asia | Uzbekistan | 3507.3 | 67.00 | 37.67 | 3497 | 0.0149 |
| I6718 | 220 | Asia | Russia | 5558.3 | 89.31 | 55.19 | 3493 | 0.0178 |
| I3125 | 611 | Europe | Italy | 3057.3 | 13.48 | 37.91 | 3490 | 0.0205 |
| I4789 | 239 | Asia | Kazakhstan | 4074.3 | 73.12 | 42.69 | 3487 | 0.0213 |
| I4322 | 234 | Asia | Kazakhstan | 4705.1 | 79.99 | 46.95 | 3484 | 0.0184 |
| I12958 | 171 | Asia | Mongolia | 5630.5 | 91.83 | 47.07 | 3483 | 0.0213 |
| ALA017 | 459 | Europe | Turkey | 1077.6 | 36.38 | 36.24 | 3478 | 0.0147 |
| I0232 | 127 | Asia | Russia | 3185.5 | 54.44 | 48.10 | 3475 | 0.0253 |
| I0359 | 130 | Asia | Russia | 3531.9 | 50.39 | 53.03 | 3475 | 0.0250 |
| I0361 | 130 | Asia | Russia | 3531.9 | 50.39 | 53.03 | 3475 | 0.0214 |
| I0422 | 135 | Asia | Russia | 3443.1 | 50.50 | 52.54 | 3475 | 0.0276 |
| I4160 | 226 | Asia | Tajikistan | 3611.2 | 68.03 | 39.45 | 3475 | 0.0235 |
| I1840 | 731 | Europe | Spain | 4402.7 | -2.62 | 42.57 | 3475 | 0.0263 |
| ALA030 | 459 | Europe | Turkey | 1077.6 | 36.38 | 36.24 | 3467 | 0.0157 |
| I4315 | 227 | Asia | Uzbekistan | 3507.3 | 67.00 | 37.75 | 3461 | 0.0138 |
| LHO001 | 989 | Europe | Spain | 4581.0 | -0.77 | 38.34 | 3461 | 0.0195 |
| I4163 | 227 | Asia | Uzbekistan | 3507.3 | 67.00 | 37.75 | 3458 | 0.0152 |
| I4783 | 234 | Asia | Kazakhstan | 4705.1 | 79.99 | 46.95 | 3456 | 0.0202 |

| | | | | | | | | |
|---|---|---|---|---|---|---|---|---|
| I4236_noUDG | 228 | Asia | Iran | 1677.7 | 45.17 | 37.01 | 3450 | 0.0185 |
| I10769 | 426 | Both | Israel | 629.1 | 35.02 | 32.58 | 3450 | 0.0128 |
| I10770 | 426 | Both | Israel | 629.1 | 35.02 | 32.58 | 3450 | 0.0159 |
| CBR004 | 499 | Europe | Spain | 4581.0 | -0.90 | 38.64 | 3448 | 0.0238 |
| BRC003.SG | 491 | Europe | Italy | 3279.7 | 11.59 | 45.47 | 3447 | 0.0236 |
| I3763 | 222 | Asia | Kazakhstan | 4924.4 | 81.84 | 50.22 | 3445 | 0.0243 |
| I7060 | 239 | Asia | Kazakhstan | 4074.3 | 73.12 | 42.69 | 3439 | 0.0157 |
| I16161 | 688 | Europe | Italy | 3570.8 | 8.32 | 40.63 | 3434 | 0.0191 |
| I6707 | 261 | Asia | Kazakhstan | 4356.6 | 73.88 | 49.71 | 3432 | 0.0228 |
| I2639 | 702 | Europe | United Kingdom | 4423.8 | -1.77 | 51.17 | 3430 | 0.0155 |
| I19858 | 646 | Europe | United Kingdom | 4423.8 | -1.79 | 51.05 | 3429 | 0.0163 |
| I26830 | 820 | Europe | Netherlands | 4151.8 | 5.14 | 52.71 | 3429 | 0.0206 |
| I8136 | 948 | Europe | Spain | 4728.9 | -2.51 | 37.73 | 3426 | 0.0256 |
| I4790 | 239 | Asia | Kazakhstan | 4074.3 | 73.12 | 42.69 | 3425 | 0.0285 |
| RISE493_noUDG.SG | 357 | Asia | Russia | 5636.6 | 91.05 | 53.15 | 3423 | 0.0294 |
| I13714 | 643 | Europe | United Kingdom | 4257.1 | 1.34 | 51.36 | 3423 | 0.0238 |
| I19857 | 646 | Europe | United Kingdom | 4423.8 | -1.79 | 51.05 | 3423 | 0.0162 |
| I19915 | 643 | Europe | United Kingdom | 4257.1 | 1.34 | 51.36 | 3422 | 0.0170 |
| I4313 | 227 | Asia | Uzbekistan | 3507.3 | 67.00 | 37.75 | 3421 | 0.0212 |
| Ne61_genotyping_noUDG | 341 | Asia | China | 8288.4 | 126.22 | 45.88 | 3421 | 0.0272 |
| RISE523_noUDG.SG | 362 | Asia | Russia | 3778.5 | 57.07 | 53.04 | 3411 | 0.0264 |
| I13174 | 171 | Asia | Mongolia | 5630.5 | 91.83 | 47.07 | 3410 | 0.0197 |
| I24342 | 743 | Europe | Croatia | 2823.8 | 15.73 | 43.89 | 3410 | 0.0215 |
| I10365 | 587 | Europe | Italy | 3620.3 | 9.23 | 39.87 | 3404 | 0.0248 |
| I19859 | 646 | Europe | United Kingdom | 4423.8 | -1.79 | 51.05 | 3401 | 0.0198 |
| I11521 | 153 | Asia | Uzbekistan | 3507.3 | 67.00 | 37.67 | 3400 | 0.0143 |
| I6667 | 230 | Asia | Turkmenistan | 2630.8 | 56.25 | 38.35 | 3400 | 0.0183 |
| esp005_noUDG.SG | 533 | Europe | Spain | 4335.6 | -2.41 | 42.32 | 3400 | 0.0237 |
| I7568 | 793 | Europe | United Kingdom | 4389.8 | 0.03 | 52.34 | 3400 | 0.0186 |
| I26828 | 820 | Europe | Netherlands | 4151.8 | 5.14 | 52.71 | 3400 | 0.0202 |
| I26829 | 820 | Europe | Netherlands | 4151.8 | 5.14 | 52.71 | 3400 | 0.0253 |

| | | | | | | | | |
|---|---|---|---|---|---|---|---|---|
| I26832 | 820 | Europe | Netherlands | 4151.8 | 5.14 | 52.71 | 3400 | 0.0222 |
| I19860 | 646 | Europe | United Kingdom | 4423.8 | -1.79 | 51.05 | 3398 | 0.0157 |
| I1656 | 202 | Asia | Armenia | 1882.1 | 43.94 | 40.38 | 3397 | 0.0232 |
| I10111 | 148 | Asia | Kazakhstan | 4561.2 | 75.60 | 50.82 | 3396 | 0.0235 |
| ALA020 | 459 | Europe | Turkey | 1077.6 | 36.38 | 36.24 | 3395 | 0.0188 |
| I6717 | 220 | Asia | Russia | 5558.3 | 89.31 | 55.19 | 3391 | 0.0259 |
| I18487 | 198 | Asia | Armenia | 1990.5 | 45.20 | 40.42 | 3390 | 0.0155 |
| I19323 | 198 | Asia | Armenia | 1990.5 | 45.20 | 40.42 | 3390 | 0.0185 |
| ARS016.A0101 | 10 | Asia | Mongolia | 6157.2 | 99.73 | 49.66 | 3389 | 0.0236 |
| U1_noUDG | 402 | Asia | Nepal | 5203.2 | 83.93 | 29.04 | 3388 | 0.0192 |
| ALA002 | 459 | Europe | Turkey | 1077.6 | 36.38 | 36.24 | 3388 | 0.0181 |
| I5074 | 879 | Europe | Croatia | 2714.9 | 17.26 | 43.17 | 3385 | 0.0207 |
| ALA018 | 459 | Europe | Turkey | 1077.6 | 36.38 | 36.24 | 3382 | 0.0193 |
| I11972 | 621 | Europe | Netherlands | 4151.8 | 5.15 | 52.70 | 3381 | 0.0264 |
| I7572 | 793 | Europe | United Kingdom | 4389.8 | 0.03 | 52.34 | 3380 | 0.0211 |
| I4098_noUDG | 115 | Asia | Iran | 1677.7 | 45.46 | 37.01 | 3378 | 0.0156 |
| R11675.SG | 353 | Asia | Armenia | 1927.6 | 43.85 | 40.69 | 3372 | 0.0186 |
| I12499 | 167 | Asia | Uzbekistan | 3363.1 | 64.99 | 40.54 | 3369 | 0.0200 |
| I2573 | 810 | Europe | United Kingdom | 4846.1 | -2.90 | 55.98 | 3361 | 0.0197 |
| RISE502_noUDG.SG | 359 | Asia | Russia | 5397.1 | 88.57 | 51.91 | 3360 | 0.0389 |
| I13964 | 176 | Asia | Mongolia | 7107.6 | 111.60 | 46.77 | 3350 | 0.0219 |
| I7579 | 941 | Europe | United Kingdom | 4389.8 | -0.51 | 52.12 | 3349 | 0.0168 |
| I12972 | 176 | Asia | Mongolia | 7107.6 | 111.60 | 46.77 | 3347 | 0.0257 |
| ALA039 | 459 | Europe | Turkey | 1077.6 | 36.38 | 36.24 | 3337 | 0.0201 |
| I16170_enhanced | 688 | Europe | Italy | 3570.8 | 8.32 | 40.63 | 3337 | 0.0184 |
| I12960 | 172 | Asia | Mongolia | 6401.6 | 102.76 | 46.92 | 3332 | 0.0178 |
| I12976 | 179 | Asia | Mongolia | 5419.2 | 88.71 | 49.36 | 3332 | 0.0204 |
| JXNTM23_noUDG | 287 | Asia | China | 7690.3 | 113.15 | 35.17 | 3332 | 0.0229 |
| I2458 | 702 | Europe | United Kingdom | 4423.8 | -1.77 | 51.17 | 3331 | 0.0227 |
| I15734 | 188 | Asia | Armenia | 1981.1 | 44.50 | 40.92 | 3325 | 0.0219 |
| I16099 | 651 | Europe | Czech Republic | 3582.7 | 13.62 | 50.53 | 3325 | 0.0179 |

| | | | | | | | | |
|---|---|---|---|---|---|---|---|---|
| I16100 | 651 | Europe | Czech Republic | 3582.7 | 13.62 | 50.53 | 3325 | 0.0172 |
| I16111 | 651 | Europe | Czech Republic | 3582.7 | 13.62 | 50.53 | 3325 | 0.0253 |
| I16112 | 651 | Europe | Czech Republic | 3582.7 | 13.62 | 50.53 | 3325 | 0.0065 |
| I26773 | 819 | Europe | Croatia | 3052.4 | 13.71 | 45.07 | 3325 | 0.0233 |
| I26774 | 819 | Europe | Croatia | 3052.4 | 13.71 | 45.07 | 3325 | 0.0243 |
| I26893 | 819 | Europe | Croatia | 3052.4 | 13.71 | 45.07 | 3325 | 0.0164 |
| I15733 | 188 | Asia | Armenia | 1981.1 | 44.50 | 40.92 | 3320 | 0.0211 |
| I15732 | 188 | Asia | Armenia | 1981.1 | 44.50 | 40.92 | 3318 | 0.0215 |
| KZL004 | 311 | Asia | Kazakhstan | 4356.7 | 75.36 | 48.28 | 3318 | 0.0195 |
| SBG001 | 22 | Asia | Mongolia | 5630.5 | 91.84 | 47.06 | 3314 | 0.0201 |
| I10344 | 586 | Europe | France | 3649.9 | 6.37 | 44.48 | 3314 | 0.0235 |
| I7640 | 940 | Europe | United Kingdom | 4389.8 | 0.12 | 52.18 | 3307 | 0.0202 |
| I14037 | 41 | Asia | Mongolia | 7120.2 | 111.77 | 46.28 | 3306 | 0.0245 |
| I7571 | 793 | Europe | United Kingdom | 4389.8 | 0.03 | 52.34 | 3306 | 0.0223 |
| BUL002 | 41 | Asia | Mongolia | 7120.2 | 111.77 | 46.28 | 3300 | 0.0163 |
| I12082 | 621 | Europe | Netherlands | 4151.8 | 5.15 | 52.70 | 3300 | 0.0234 |
| I13775 | 639 | Europe | Montenegro | 2535.7 | 18.73 | 42.37 | 3300 | 0.0201 |
| I13777 | 639 | Europe | Montenegro | 2535.7 | 18.73 | 42.37 | 3300 | 0.0232 |
| BIL001 | 24 | Asia | Mongolia | 5936.2 | 96.40 | 49.35 | 3297 | 0.0235 |
| I5695 | 785 | Europe | Slovenia | 2983.9 | 15.69 | 45.85 | 3296 | 0.0242 |
| I6347 | 246 | Asia | Mongolia | 5862.3 | 95.45 | 49.31 | 3295 | 0.0212 |
| I13168 | 639 | Europe | Montenegro | 2535.7 | 18.73 | 42.37 | 3293 | 0.0252 |
| I4340_noUDG | 228 | Asia | Iran | 1677.7 | 45.17 | 37.01 | 3291 | 0.0223 |
| ULN003 | 41 | Asia | Mongolia | 7120.2 | 111.77 | 46.28 | 3289 | 0.0219 |
| I13169 | 639 | Europe | Montenegro | 2535.7 | 18.73 | 42.37 | 3287 | 0.0179 |
| I2654 | 810 | Europe | United Kingdom | 4846.1 | -2.90 | 55.98 | 3287 | 0.0200 |
| I18166 | 92 | Asia | Armenia | 1936.1 | 44.93 | 40.52 | 3285 | 0.0182 |
| I18167 | 92 | Asia | Armenia | 1936.1 | 44.93 | 40.52 | 3285 | 0.0144 |
| I18168 | 92 | Asia | Armenia | 1936.1 | 44.93 | 40.52 | 3285 | 0.0173 |
| I18267 | 92 | Asia | Armenia | 1936.1 | 44.93 | 40.52 | 3285 | 0.0199 |
| I18269 | 92 | Asia | Armenia | 1936.1 | 44.93 | 40.52 | 3285 | 0.0183 |

| | | | | | | | | |
|---|---|---|---|---|---|---|---|---|
| I18270 | 92 | Asia | Armenia | 1936.1 | 44.93 | 40.52 | 3285 | 0.0196 |
| I18271 | 92 | Asia | Armenia | 1936.1 | 44.93 | 40.52 | 3285 | 0.0195 |
| I18272 | 92 | Asia | Armenia | 1936.1 | 44.93 | 40.52 | 3285 | 0.0199 |
| I18273 | 92 | Asia | Armenia | 1936.1 | 44.93 | 40.52 | 3285 | 0.0169 |
| I18275 | 92 | Asia | Armenia | 1936.1 | 44.93 | 40.52 | 3285 | 0.0177 |
| I18276 | 92 | Asia | Armenia | 1936.1 | 44.93 | 40.52 | 3285 | 0.0166 |
| I18277 | 92 | Asia | Armenia | 1936.1 | 44.93 | 40.52 | 3285 | 0.0192 |
| I18278 | 92 | Asia | Armenia | 1936.1 | 44.93 | 40.52 | 3285 | 0.0146 |
| I18279 | 92 | Asia | Armenia | 1936.1 | 44.93 | 40.52 | 3285 | 0.0190 |
| I18280 | 92 | Asia | Armenia | 1936.1 | 44.93 | 40.52 | 3285 | 0.0160 |
| I15748 | 188 | Asia | Armenia | 1981.1 | 44.50 | 40.92 | 3285 | 0.0212 |
| I15750 | 188 | Asia | Armenia | 1981.1 | 44.50 | 40.92 | 3285 | 0.0180 |
| I18472 | 198 | Asia | Armenia | 1990.5 | 45.20 | 40.42 | 3285 | 0.0174 |
| I18486 | 198 | Asia | Armenia | 1990.5 | 45.20 | 40.42 | 3285 | 0.0198 |
| I19325 | 198 | Asia | Armenia | 1990.5 | 45.20 | 40.42 | 3285 | 0.0166 |
| I19335 | 198 | Asia | Armenia | 1990.5 | 45.20 | 40.42 | 3285 | 0.0122 |
| I19336 | 198 | Asia | Armenia | 1990.5 | 45.20 | 40.42 | 3285 | 0.0146 |
| I19341 | 198 | Asia | Armenia | 1990.5 | 45.20 | 40.42 | 3285 | 0.0133 |
| I19344 | 198 | Asia | Armenia | 1990.5 | 45.20 | 40.42 | 3285 | 0.0165 |
| I19348 | 198 | Asia | Armenia | 1990.5 | 45.20 | 40.42 | 3285 | 0.0183 |
| I19351 | 198 | Asia | Armenia | 1990.5 | 45.20 | 40.42 | 3285 | 0.0182 |
| I19352 | 198 | Asia | Armenia | 1990.5 | 45.20 | 40.42 | 3285 | 0.0178 |
| RISE496_noUDG.SG | 358 | Asia | Russia | 5557.3 | 90.19 | 52.95 | 3285 | 0.0314 |
| I11973 | 621 | Europe | Netherlands | 4151.8 | 5.15 | 52.70 | 3280 | 0.0199 |
| I19366 | 642 | Europe | Greece | 2206.5 | 21.69 | 37.03 | 3278 | 0.0148 |
| R11545.SG | 353 | Asia | Armenia | 1927.6 | 43.85 | 40.69 | 3276 | 0.0293 |
| I6264 | 245 | Asia | Mongolia | 6258.0 | 100.83 | 45.92 | 3275 | 0.0187 |
| I2470_enhanced | 731 | Europe | Spain | 4402.7 | -2.62 | 42.57 | 3275 | 0.0263 |
| I19913 | 643 | Europe | United Kingdom | 4257.1 | 1.34 | 51.36 | 3271 | 0.0134 |
| I7574 | 940 | Europe | United Kingdom | 4389.8 | 0.12 | 52.18 | 3271 | 0.0235 |
| I17183 | 206 | Asia | Armenia | 1927.6 | 43.84 | 40.87 | 3270 | 0.0129 |

| | | | | | | | | |
|---|---|---|---|---|---|---|---|---|
| I6262 | 245 | Asia | Mongolia | 6258.0 | 100.83 | 45.92 | 3270 | 0.0240 |
| I7627 | 941 | Europe | United Kingdom | 4389.8 | -0.51 | 52.12 | 3268 | 0.0276 |
| DA35_noUDG.SG | 92 | Asia | Armenia | 1936.1 | 44.93 | 40.52 | 3259 | 0.0196 |
| I12081 | 621 | Europe | Netherlands | 4151.8 | 5.15 | 52.70 | 3257 | 0.0221 |
| ARS017.A0101 | 10 | Asia | Mongolia | 6157.2 | 99.73 | 49.66 | 3256 | 0.0224 |
| I14767 | 196 | Asia | Turkey | 1745.4 | 43.75 | 38.98 | 3256 | 0.0115 |
| I6668 | 230 | Asia | Turkmenistan | 2630.8 | 56.25 | 38.35 | 3250 | 0.0227 |
| I6675 | 260 | Asia | Turkmenistan | 2799.3 | 58.43 | 37.93 | 3250 | 0.0157 |
| JXNTM2_noUDG | 287 | Asia | China | 7690.3 | 113.15 | 35.17 | 3250 | 0.0248 |
| I12208 | 625 | Europe | Spain | 4597.3 | -5.29 | 41.46 | 3250 | 0.0169 |
| I12209 | 625 | Europe | Spain | 4597.3 | -5.29 | 41.46 | 3250 | 0.0203 |
| ULN001 | 41 | Asia | Mongolia | 7120.2 | 111.77 | 46.28 | 3239 | 0.0201 |
| I10552 | 598 | Europe | Italy | 3620.3 | 9.43 | 39.68 | 3234 | 0.0236 |
| I13778 | 639 | Europe | Montenegro | 2535.7 | 18.73 | 42.37 | 3234 | 0.0232 |
| I7626 | 941 | Europe | United Kingdom | 4389.8 | -0.51 | 52.12 | 3231 | 0.0259 |
| I12973 | 177 | Asia | Mongolia | 6379.3 | 102.69 | 49.41 | 3226 | 0.0222 |
| I2653 | 810 | Europe | United Kingdom | 4846.1 | -2.90 | 55.98 | 3224 | 0.0190 |
| I13716 | 647 | Europe | United Kingdom | 4257.1 | 0.48 | 51.33 | 3223 | 0.0285 |
| I19329 | 198 | Asia | Armenia | 1990.5 | 45.20 | 40.42 | 3217 | 0.0155 |
| I16220 | 198 | Asia | Armenia | 1990.5 | 45.20 | 40.42 | 3216 | 0.0166 |
| I6900 | 208 | Asia | Pakistan | 3962.5 | 72.31 | 34.74 | 3216 | 0.0164 |
| I4787 | 238 | Asia | Kazakhstan | 3861.3 | 67.02 | 48.21 | 3216 | 0.0240 |
| I3878 | 589 | Europe | Italy | 3070.3 | 12.79 | 37.68 | 3216 | 0.0178 |
| I13766 | 185 | Asia | Mongolia | 6252.4 | 100.82 | 46.43 | 3208 | 0.0223 |
| I13172 | 639 | Europe | Montenegro | 2535.7 | 18.73 | 42.37 | 3207 | 0.0217 |
| I10373 | 589 | Europe | Italy | 3070.3 | 12.79 | 37.68 | 3200 | 0.0148 |
| I20749 | 763 | Europe | Hungary | 2958.8 | 21.50 | 47.89 | 3200 | 0.0232 |
| I20751 | 764 | Europe | Hungary | 2913.5 | 20.26 | 47.21 | 3200 | 0.0218 |
| I13767 | 187 | Asia | Mongolia | 6390.8 | 102.55 | 48.11 | 3198 | 0.0254 |
| I10372 | 589 | Europe | Italy | 3070.3 | 12.79 | 37.68 | 3198 | 0.0142 |
| I7577 | 941 | Europe | United Kingdom | 4389.8 | -0.51 | 52.12 | 3198 | 0.0173 |

| | | | | | | | | |
|---|---|---|---|---|---|---|---|---|
| ORC003 | 1032 | Europe | Italy | 3620.3 | 9.43 | 39.67 | 3181 | 0.0229 |
| I13171 | 639 | Europe | Montenegro | 2535.7 | 18.73 | 42.37 | 3175 | 0.0230 |
| I14501 | 639 | Europe | Montenegro | 2535.7 | 18.73 | 42.37 | 3175 | 0.0237 |
| ARS008.A0101 | 11 | Asia | Mongolia | 6157.2 | 99.77 | 49.69 | 3173 | 0.0237 |
| ORC007 | 1032 | Europe | Italy | 3620.3 | 9.43 | 39.67 | 3173 | 0.0250 |
| ARS007.B0101 | 11 | Asia | Mongolia | 6157.2 | 99.77 | 49.69 | 3172 | 0.0237 |
| I2062 | 428 | Both | Israel | 629.1 | 35.23 | 32.66 | 3170 | 0.0180 |
| I7231 | 931 | Europe | North Macedonia | 2286.2 | 21.94 | 41.58 | 3169 | 0.0204 |
| BER002 | 22 | Asia | Mongolia | 5630.5 | 91.84 | 47.06 | 3168 | 0.0245 |
| I13514 | 642 | Europe | Greece | 2206.5 | 21.69 | 37.03 | 3165 | 0.0203 |
| ORC001 | 1032 | Europe | Italy | 3620.3 | 9.43 | 39.67 | 3163 | 0.0221 |
| I18274 | 92 | Asia | Armenia | 1936.1 | 44.93 | 40.52 | 3150 | 0.0190 |
| I15729 | 188 | Asia | Armenia | 1981.1 | 44.50 | 40.92 | 3150 | 0.0203 |
| I15749 | 188 | Asia | Armenia | 1981.1 | 44.50 | 40.92 | 3150 | 0.0152 |
| I18479 | 198 | Asia | Armenia | 1990.5 | 45.20 | 40.42 | 3150 | 0.0176 |
| I18483 | 198 | Asia | Armenia | 1990.5 | 45.20 | 40.42 | 3150 | 0.0177 |
| I19326 | 198 | Asia | Armenia | 1990.5 | 45.20 | 40.42 | 3150 | 0.0230 |
| I3769 | 224 | Asia | Kazakhstan | 4415.5 | 76.58 | 43.37 | 3150 | 0.0220 |
| L2 | 312 | Asia | Nepal | 5215.9 | 83.75 | 28.80 | 3150 | 0.0205 |
| ORC005 | 1032 | Europe | Italy | 3620.3 | 9.43 | 39.67 | 3140 | 0.0207 |
| I20440 | 212 | Asia | Armenia | 1990.5 | 45.13 | 40.29 | 3135 | 0.0169 |
| ULI001 | 406 | Asia | Mongolia | 5632.1 | 91.93 | 45.86 | 3131 | 0.0185 |
| I6432 | 115 | Asia | Iran | 1677.7 | 45.46 | 37.01 | 3128 | 0.0151 |
| I13948 | 188 | Asia | Armenia | 1981.1 | 44.50 | 40.92 | 3128 | 0.0153 |
| I8193 | 270 | Asia | Pakistan | 3962.5 | 72.31 | 34.75 | 3128 | 0.0142 |
| I18162 | 150 | Asia | Armenia | 1927.6 | 43.92 | 40.83 | 3125 | 0.0141 |
| I18163 | 150 | Asia | Armenia | 1927.6 | 43.92 | 40.83 | 3125 | 0.0226 |
| I18164 | 150 | Asia | Armenia | 1927.6 | 43.92 | 40.83 | 3125 | 0.0172 |
| I18165 | 150 | Asia | Armenia | 1927.6 | 43.92 | 40.83 | 3125 | 0.0118 |
| I18247 | 150 | Asia | Armenia | 1927.6 | 43.92 | 40.83 | 3125 | 0.0203 |
| I18248 | 150 | Asia | Armenia | 1927.6 | 43.92 | 40.83 | 3125 | 0.0176 |

| | | | | | | | | |
|---|---|---|---|---|---|---|---|---|
| I16196 | 188 | Asia | Armenia | 1981.1 | 44.50 | 40.92 | 3125 | 0.0206 |
| I19331 | 198 | Asia | Armenia | 1990.5 | 45.20 | 40.42 | 3125 | 0.0204 |
| I19332 | 198 | Asia | Armenia | 1990.5 | 45.20 | 40.42 | 3125 | 0.0146 |
| I19333 | 198 | Asia | Armenia | 1990.5 | 45.20 | 40.42 | 3125 | 0.0131 |
| I19334 | 198 | Asia | Armenia | 1990.5 | 45.20 | 40.42 | 3125 | 0.0227 |
| I19339 | 198 | Asia | Armenia | 1990.5 | 45.20 | 40.42 | 3125 | 0.0131 |
| I19342 | 198 | Asia | Armenia | 1990.5 | 45.20 | 40.42 | 3125 | 0.0133 |
| I19345 | 198 | Asia | Armenia | 1990.5 | 45.20 | 40.42 | 3125 | 0.0159 |
| I19346 | 198 | Asia | Armenia | 1990.5 | 45.20 | 40.42 | 3125 | 0.0175 |
| I19347 | 198 | Asia | Armenia | 1990.5 | 45.20 | 40.42 | 3125 | 0.0161 |
| I19350 | 198 | Asia | Armenia | 1990.5 | 45.20 | 40.42 | 3125 | 0.0135 |
| ORC006 | 1032 | Europe | Italy | 3620.3 | 9.43 | 39.67 | 3123 | 0.0188 |
| I10554 | 598 | Europe | Italy | 3620.3 | 9.43 | 39.68 | 3122 | 0.0254 |
| I6367 | 256 | Asia | Mongolia | 5409.1 | 89.50 | 48.30 | 3116 | 0.0267 |
| I12975 | 178 | Asia | Mongolia | 6232.4 | 100.67 | 51.50 | 3110 | 0.0250 |
| I10553 | 598 | Europe | Italy | 3620.3 | 9.43 | 39.68 | 3110 | 0.0229 |
| I7575 | 941 | Europe | United Kingdom | 4389.8 | -0.51 | 52.12 | 3107 | 0.0247 |
| I10110 | 148 | Asia | Kazakhstan | 4561.2 | 75.60 | 50.82 | 3100 | 0.0249 |
| RISE495_noUDG.SG | 358 | Asia | Russia | 5557.3 | 90.19 | 52.95 | 3100 | 0.0237 |
| RISE497_noUDG.SG | 358 | Asia | Russia | 5557.3 | 90.19 | 52.95 | 3100 | 0.0276 |
| RISE499_noUDG.SG | 359 | Asia | Russia | 5397.1 | 88.57 | 51.91 | 3100 | 0.0309 |
| I12083 | 621 | Europe | Netherlands | 4151.8 | 5.15 | 52.70 | 3100 | 0.0159 |
| I14498 | 639 | Europe | Montenegro | 2535.7 | 18.73 | 42.37 | 3100 | 0.0194 |
| I18415 | 732 | Europe | Croatia | 2958.4 | 15.42 | 44.85 | 3100 | 0.0154 |
| I18416 | 732 | Europe | Croatia | 2958.4 | 15.42 | 44.85 | 3100 | 0.0163 |
| I18417 | 732 | Europe | Croatia | 2958.4 | 15.42 | 44.85 | 3100 | 0.0211 |
| I18717 | 732 | Europe | Croatia | 2958.4 | 15.42 | 44.85 | 3100 | 0.0220 |
| I18719 | 732 | Europe | Croatia | 2958.4 | 15.42 | 44.85 | 3100 | 0.0220 |
| I18721 | 732 | Europe | Croatia | 2958.4 | 15.42 | 44.85 | 3100 | 0.0197 |
| I18723 | 732 | Europe | Croatia | 2958.4 | 15.42 | 44.85 | 3100 | 0.0217 |
| I18724 | 732 | Europe | Croatia | 2958.4 | 15.42 | 44.85 | 3100 | 0.0237 |

| | | | | | | | | |
|---|---|---|---|---|---|---|---|---|
| I18725 | 732 | Europe | Croatia | 2958.4 | 15.42 | 44.85 | 3100 | 0.0163 |
| I18729 | 732 | Europe | Croatia | 2958.4 | 15.42 | 44.85 | 3100 | 0.0204 |
| I18730 | 732 | Europe | Croatia | 2958.4 | 15.42 | 44.85 | 3100 | 0.0138 |
| I18732 | 732 | Europe | Croatia | 2958.4 | 15.42 | 44.85 | 3100 | 0.0204 |
| I18737 | 732 | Europe | Croatia | 2958.4 | 15.42 | 44.85 | 3100 | 0.0197 |
| I18738 | 732 | Europe | Croatia | 2958.4 | 15.42 | 44.85 | 3100 | 0.0177 |
| I6363 | 254 | Asia | Mongolia | 5554.2 | 90.85 | 45.35 | 3095 | 0.0237 |
| I4295 | 222 | Asia | Kazakhstan | 4924.4 | 81.84 | 50.22 | 3090 | 0.0244 |
| I13167 | 639 | Europe | Montenegro | 2535.7 | 18.73 | 42.37 | 3090 | 0.0196 |
| I3741 | 598 | Europe | Italy | 3620.3 | 9.43 | 39.68 | 3088 | 0.0196 |
| I7580 | 941 | Europe | United Kingdom | 4389.8 | -0.51 | 52.12 | 3087 | 0.0145 |
| I13518 | 642 | Europe | Greece | 2206.5 | 21.69 | 37.03 | 3085 | 0.0196 |
| C1367 | 45 | Asia | China | 4909.4 | 82.64 | 43.79 | 3084 | 0.0214 |
| I16554 | 188 | Asia | Armenia | 1981.1 | 44.50 | 40.92 | 3083 | 0.0162 |
| I2656 | 815 | Europe | United Kingdom | 4846.1 | -2.90 | 55.97 | 3081 | 0.0176 |
| I7578 | 941 | Europe | United Kingdom | 4389.8 | -0.51 | 52.12 | 3080 | 0.0191 |
| I16217 | 198 | Asia | Armenia | 1990.5 | 45.20 | 40.42 | 3074 | 0.0154 |
| M9984_noUDG.SG | 320 | Asia | Russia | 9248.1 | 151.00 | 59.60 | 3068 | 0.0263 |
| I7033 | 174 | Asia | Mongolia | 5649.6 | 92.05 | 49.96 | 3066 | 0.0190 |
| I19338 | 198 | Asia | Armenia | 1990.5 | 45.20 | 40.42 | 3066 | 0.0174 |
| I7039 | 268 | Asia | Mongolia | 5631.2 | 92.23 | 47.42 | 3066 | 0.0192 |
| ORC009 | 1032 | Europe | Italy | 3620.3 | 9.43 | 39.67 | 3062 | 0.0188 |
| I11971 | 621 | Europe | Netherlands | 4151.8 | 5.15 | 52.70 | 3060 | 0.0178 |
| I15955 | 558 | Europe | Czech Republic | 3495.2 | 14.28 | 50.08 | 3059 | 0.0220 |
| I15650 | 558 | Europe | Czech Republic | 3495.2 | 14.28 | 50.08 | 3058 | 0.0188 |
| ARS002.B0101 | 10 | Asia | Mongolia | 6157.2 | 99.73 | 49.66 | 3055 | 0.0206 |
| I19354 | 198 | Asia | Armenia | 1990.5 | 45.20 | 40.42 | 3052 | 0.0156 |
| I16116 | 188 | Asia | Armenia | 1981.1 | 44.50 | 40.92 | 3050 | 0.0114 |
| I18468 | 198 | Asia | Armenia | 1990.5 | 45.20 | 40.42 | 3050 | 0.0124 |
| I18470 | 198 | Asia | Armenia | 1990.5 | 45.20 | 40.42 | 3050 | 0.0166 |
| I18481 | 198 | Asia | Armenia | 1990.5 | 45.20 | 40.42 | 3050 | 0.0165 |

| | | | | | | | | |
|---|---|---|---|---|---|---|---|---|
| I19321 | 198 | Asia | Armenia | 1990.5 | 45.20 | 40.42 | 3050 | 0.0151 |
| I2327_noUDG | 215 | Asia | Iran | 1633.2 | 45.47 | 36.99 | 3050 | 0.0183 |
| I4255 | 229 | Asia | Uzbekistan | 3966.2 | 71.86 | 40.38 | 3050 | 0.0176 |
| M0831_noUDG.SG | 320 | Asia | Russia | 9248.1 | 151.00 | 59.60 | 3050 | 0.0317 |
| I13786 | 558 | Europe | Czech Republic | 3495.2 | 14.28 | 50.08 | 3050 | 0.0200 |
| I13794 | 558 | Europe | Czech Republic | 3495.2 | 14.28 | 50.08 | 3050 | 0.0257 |
| I13798 | 558 | Europe | Czech Republic | 3495.2 | 14.28 | 50.08 | 3050 | 0.0177 |
| I13799 | 558 | Europe | Czech Republic | 3495.2 | 14.28 | 50.08 | 3050 | 0.0206 |
| I15041 | 558 | Europe | Czech Republic | 3495.2 | 14.28 | 50.08 | 3050 | 0.0170 |
| I15956 | 558 | Europe | Czech Republic | 3495.2 | 14.28 | 50.08 | 3050 | 0.0130 |
| I15957 | 558 | Europe | Czech Republic | 3495.2 | 14.28 | 50.08 | 3050 | 0.0177 |
| I15958 | 558 | Europe | Czech Republic | 3495.2 | 14.28 | 50.08 | 3050 | 0.0146 |
| I15960 | 558 | Europe | Czech Republic | 3495.2 | 14.28 | 50.08 | 3050 | 0.0157 |
| I16182 | 558 | Europe | Czech Republic | 3495.2 | 14.28 | 50.08 | 3050 | 0.0184 |
| I16488 | 695 | Europe | United Kingdom | 4580.0 | -4.21 | 51.54 | 3050 | 0.0209 |
| ULI003 | 406 | Asia | Mongolia | 5632.1 | 91.93 | 45.86 | 3046 | 0.0185 |
| I4267 | 222 | Asia | Kazakhstan | 4924.4 | 81.84 | 50.22 | 3042 | 0.0219 |
| I7576 | 941 | Europe | United Kingdom | 4389.8 | -0.51 | 52.12 | 3039 | 0.0191 |
| ARS005.A0101 | 11 | Asia | Mongolia | 6157.2 | 99.77 | 49.69 | 3038 | 0.0239 |
| I3976 | 222 | Asia | Kazakhstan | 4924.4 | 81.84 | 50.22 | 3035 | 0.0272 |
| I13795 | 651 | Europe | Czech Republic | 3582.7 | 13.62 | 50.53 | 3035 | 0.0192 |
| I19343 | 198 | Asia | Armenia | 1990.5 | 45.20 | 40.42 | 3025 | 0.0146 |
| I0099 | 563 | Europe | Germany | 3796.2 | 11.05 | 51.90 | 3024 | 0.0228 |
| ORC008 | 1032 | Europe | Italy | 3620.3 | 9.43 | 39.67 | 3023 | 0.0179 |
| I6431 | 115 | Asia | Iran | 1677.7 | 45.46 | 37.01 | 3022 | 0.0173 |
| I13783 | 558 | Europe | Czech Republic | 3495.2 | 14.28 | 50.08 | 3022 | 0.0244 |
| I15959 | 558 | Europe | Czech Republic | 3495.2 | 14.28 | 50.08 | 3022 | 0.0196 |
| I7949 | 916 | Europe | Czech Republic | 3537.1 | 13.78 | 50.55 | 3022 | 0.0233 |
| I6388 | 115 | Asia | Iran | 1677.7 | 45.46 | 37.01 | 3015 | 0.0136 |
| I3977 | 222 | Asia | Kazakhstan | 4924.4 | 81.84 | 50.22 | 3015 | 0.0174 |
| I7629 | 943 | Europe | United Kingdom | 4503.2 | -0.52 | 53.73 | 3015 | 0.0259 |

| | | | | | | | | |
|---|---|---|---|---|---|---|---|---|
| MIT001 | 323 | Asia | Mongolia | 6625.3 | 105.67 | 47.33 | 3010 | 0.0244 |
| ARS006.A0101 | 12 | Asia | Mongolia | 6157.2 | 99.76 | 49.69 | 3003 | 0.0234 |
| I1799 | 208 | Asia | Pakistan | 3962.5 | 72.31 | 34.74 | 3003 | 0.0203 |
| I12969 | 173 | Asia | Mongolia | 6941.4 | 109.19 | 47.18 | 3001 | 0.0228 |
| I13505 | 185 | Asia | Mongolia | 6252.4 | 100.82 | 46.43 | 3001 | 0.0222 |
| I13789 | 649 | Europe | Czech Republic | 3537.1 | 13.76 | 50.56 | 3000 | 0.0258 |
| I13788 | 650 | Europe | Czech Republic | 3537.1 | 13.77 | 50.51 | 3000 | 0.0259 |
| I13792 | 650 | Europe | Czech Republic | 3537.1 | 13.77 | 50.51 | 3000 | 0.0234 |
| I13793 | 650 | Europe | Czech Republic | 3537.1 | 13.77 | 50.51 | 3000 | 0.0275 |
| I13796 | 650 | Europe | Czech Republic | 3537.1 | 13.77 | 50.51 | 3000 | 0.0220 |
| I16089 | 650 | Europe | Czech Republic | 3537.1 | 13.77 | 50.51 | 3000 | 0.0211 |
| I14690 | 669 | Europe | Albania | 2438.6 | 20.39 | 42.02 | 3000 | 0.0189 |
| La364.SG | 314 | Asia | Laos | 7378.5 | 103.41 | 20.21 | 2987 | 0.0147 |
| I14618 | 188 | Asia | Armenia | 1981.1 | 44.50 | 40.92 | 2986 | 0.0234 |
| I19353 | 198 | Asia | Armenia | 1990.5 | 45.20 | 40.42 | 2986 | 0.0178 |
| I6428 | 115 | Asia | Iran | 1677.7 | 45.46 | 37.01 | 2977 | 0.0124 |
| I16537 | 188 | Asia | Armenia | 1981.1 | 44.50 | 40.92 | 2977 | 0.0196 |
| I3753 | 222 | Asia | Kazakhstan | 4924.4 | 81.84 | 50.22 | 2977 | 0.0260 |
| I13781 | 649 | Europe | Czech Republic | 3537.1 | 13.76 | 50.56 | 2977 | 0.0207 |
| I13791 | 649 | Europe | Czech Republic | 3537.1 | 13.76 | 50.56 | 2975 | 0.0260 |
| I4356_noUDG | 115 | Asia | Iran | 1677.7 | 45.46 | 37.01 | 2967 | 0.0196 |
| I13787 | 558 | Europe | Czech Republic | 3495.2 | 14.28 | 50.08 | 2958 | 0.0286 |
| ARS026.A0101 | 13 | Asia | Mongolia | 6157.2 | 99.78 | 49.69 | 2957 | 0.0201 |
| I6194 | 208 | Asia | Pakistan | 3962.5 | 72.31 | 34.74 | 2950 | 0.0206 |
| I6197 | 208 | Asia | Pakistan | 3962.5 | 72.31 | 34.74 | 2950 | 0.0171 |
| I6198 | 208 | Asia | Pakistan | 3962.5 | 72.31 | 34.74 | 2950 | 0.0162 |
| I6897 | 208 | Asia | Pakistan | 3962.5 | 72.31 | 34.74 | 2950 | 0.0167 |
| I6901 | 208 | Asia | Pakistan | 3962.5 | 72.31 | 34.74 | 2950 | 0.0247 |
| I8194 | 270 | Asia | Pakistan | 3962.5 | 72.31 | 34.75 | 2950 | 0.0173 |
| I12138 | 147 | Asia | Pakistan | 3962.5 | 72.40 | 34.75 | 2949 | 0.0254 |
| I6352 | 249 | Asia | Mongolia | 6401.6 | 102.77 | 46.90 | 2949 | 0.0225 |

| | | | | | | | | |
|---|---|---|---|---|---|---|---|---|
| I17182 | 206 | Asia | Armenia | 1927.6 | 43.84 | 40.87 | 2941 | 0.0079 |
| I1985 | 208 | Asia | Pakistan | 3962.5 | 72.31 | 34.74 | 2939 | 0.0149 |
| R11536.SG | 353 | Asia | Armenia | 1927.6 | 43.85 | 40.69 | 2932 | 0.0198 |
| KHI001 | 22 | Asia | Mongolia | 5630.5 | 91.84 | 47.06 | 2930 | 0.0179 |
| SHU001 | 383 | Asia | Mongolia | 6401.6 | 102.77 | 46.91 | 2929 | 0.0205 |
| I12457 | 147 | Asia | Pakistan | 3962.5 | 72.40 | 34.75 | 2928 | 0.0178 |
| I3876 | 589 | Europe | Italy | 3070.3 | 12.79 | 37.68 | 2928 | 0.0097 |
| I13711 | 643 | Europe | United Kingdom | 4257.1 | 1.34 | 51.36 | 2928 | 0.0267 |
| I6364 | 181 | Asia | Mongolia | 5554.2 | 90.80 | 45.39 | 2927 | 0.0231 |
| I6362 | 253 | Asia | Mongolia | 5630.9 | 92.03 | 46.06 | 2921 | 0.0178 |
| I5729 | 904 | Europe | Croatia | 3023.7 | 16.69 | 46.26 | 2920 | 0.0140 |
| I14617 | 188 | Asia | Armenia | 1981.1 | 44.50 | 40.92 | 2915 | 0.0243 |
| I14620 | 188 | Asia | Armenia | 1981.1 | 44.50 | 40.92 | 2915 | 0.0150 |
| I13712 | 643 | Europe | United Kingdom | 4257.1 | 1.34 | 51.36 | 2914 | 0.0236 |
| I13713 | 643 | Europe | United Kingdom | 4257.1 | 1.34 | 51.36 | 2912 | 0.0242 |
| I4353_noUDG | 115 | Asia | Iran | 1677.7 | 45.46 | 37.01 | 2911 | 0.0173 |
| I20439 | 212 | Asia | Armenia | 1990.5 | 45.13 | 40.29 | 2911 | 0.0151 |
| I3772 | 222 | Asia | Kazakhstan | 4924.4 | 81.84 | 50.22 | 2911 | 0.0248 |
| I3260 | 208 | Asia | Pakistan | 3962.5 | 72.31 | 34.74 | 2908 | 0.0167 |
| I6429 | 115 | Asia | Iran | 1677.7 | 45.46 | 37.01 | 2907 | 0.0192 |
| I10974 | 147 | Asia | Pakistan | 3962.5 | 72.40 | 34.75 | 2903 | 0.0233 |
| I16814 | 709 | Europe | Serbia | 2823.1 | 20.17 | 45.89 | 2903 | 0.0189 |
| ARS009.A0101 | 10 | Asia | Mongolia | 6157.2 | 99.73 | 49.66 | 2900 | 0.0270 |
| I14692 | 669 | Europe | Albania | 2438.6 | 20.39 | 42.02 | 2900 | 0.0227 |
| I4355_noUDG | 115 | Asia | Iran | 1677.7 | 45.46 | 37.01 | 2898 | 0.0168 |
| I20443 | 212 | Asia | Armenia | 1990.5 | 45.13 | 40.29 | 2896 | 0.0141 |
| I2201 | 429 | Both | Israel | 678.0 | 35.58 | 33.26 | 2889 | 0.0163 |
| I14377 | 661 | Europe | United Kingdom | 4257.1 | 1.37 | 51.33 | 2889 | 0.0289 |
| I6351 | 248 | Asia | Mongolia | 5791.2 | 93.80 | 49.70 | 2885 | 0.0239 |
| ARS001.A0101 | 10 | Asia | Mongolia | 6157.2 | 99.73 | 49.66 | 2880 | 0.0227 |
| I14603 | 188 | Asia | Armenia | 1981.1 | 44.50 | 40.92 | 2877 | 0.0205 |

| | | | | | | | | |
|---|---|---|---|---|---|---|---|---|
| I14065 | 188 | Asia | Armenia | 1981.1 | 44.50 | 40.92 | 2875 | 0.0208 |
| I14066 | 188 | Asia | Armenia | 1981.1 | 44.50 | 40.92 | 2875 | 0.0194 |
| I14588 | 188 | Asia | Armenia | 1981.1 | 44.50 | 40.92 | 2875 | 0.0144 |
| I14601 | 188 | Asia | Armenia | 1981.1 | 44.50 | 40.92 | 2875 | 0.0178 |
| I14602 | 188 | Asia | Armenia | 1981.1 | 44.50 | 40.92 | 2875 | 0.0199 |
| I14619 | 188 | Asia | Armenia | 1981.1 | 44.50 | 40.92 | 2875 | 0.0179 |
| I14621 | 188 | Asia | Armenia | 1981.1 | 44.50 | 40.92 | 2875 | 0.0249 |
| I25504 | 805 | Europe | Hungary | 3098.7 | 16.79 | 47.35 | 2875 | 0.0240 |
| I16549 | 700 | Europe | Czech Republic | 3495.2 | 14.28 | 50.06 | 2868 | 0.0107 |
| I18469 | 198 | Asia | Armenia | 1990.5 | 45.20 | 40.42 | 2865 | 0.0187 |
| I19324 | 198 | Asia | Armenia | 1990.5 | 45.20 | 40.42 | 2865 | 0.0238 |
| BR2_noUDG.SG | 490 | Europe | Hungary | 3003.9 | 20.09 | 47.73 | 2858 | 0.0160 |
| DA4_noUDG.SG | 80 | Asia | Russia | 5674.0 | 90.85 | 54.68 | 2850 | 0.0244 |
| DA6_noUDG.SG | 80 | Asia | Russia | 5674.0 | 90.85 | 54.68 | 2850 | 0.0218 |
| DA8_noUDG.SG | 80 | Asia | Russia | 5674.0 | 90.85 | 54.68 | 2850 | 0.0277 |
| DA9_noUDG.SG | 80 | Asia | Russia | 5674.0 | 90.85 | 54.68 | 2850 | 0.0273 |
| I6430 | 115 | Asia | Iran | 1677.7 | 45.46 | 37.01 | 2850 | 0.0190 |
| I10000 | 147 | Asia | Pakistan | 3962.5 | 72.40 | 34.75 | 2850 | 0.0095 |
| I10001 | 147 | Asia | Pakistan | 3962.5 | 72.40 | 34.75 | 2850 | 0.0170 |
| I12134 | 147 | Asia | Pakistan | 3962.5 | 72.40 | 34.75 | 2850 | 0.0232 |
| I12136 | 147 | Asia | Pakistan | 3962.5 | 72.40 | 34.75 | 2850 | 0.0148 |
| I12137 | 147 | Asia | Pakistan | 3962.5 | 72.40 | 34.75 | 2850 | 0.0167 |
| I12459 | 147 | Asia | Pakistan | 3962.5 | 72.40 | 34.75 | 2850 | 0.0151 |
| I12979 | 147 | Asia | Pakistan | 3962.5 | 72.40 | 34.75 | 2850 | 0.0190 |
| I12981 | 147 | Asia | Pakistan | 3962.5 | 72.40 | 34.75 | 2850 | 0.0163 |
| I12982 | 147 | Asia | Pakistan | 3962.5 | 72.40 | 34.75 | 2850 | 0.0183 |
| I12983 | 147 | Asia | Pakistan | 3962.5 | 72.40 | 34.75 | 2850 | 0.0188 |
| I12984 | 147 | Asia | Pakistan | 3962.5 | 72.40 | 34.75 | 2850 | 0.0195 |
| I12985 | 147 | Asia | Pakistan | 3962.5 | 72.40 | 34.75 | 2850 | 0.0161 |
| I12987 | 147 | Asia | Pakistan | 3962.5 | 72.40 | 34.75 | 2850 | 0.0206 |
| I12988 | 147 | Asia | Pakistan | 3962.5 | 72.40 | 34.75 | 2850 | 0.0118 |

| | | | | | | | | |
|---|---|---|---|---|---|---|---|---|
| I13220 | 147 | Asia | Pakistan | 3962.5 | 72.40 | 34.75 | 2850 | 0.0131 |
| I13223 | 147 | Asia | Pakistan | 3962.5 | 72.40 | 34.75 | 2850 | 0.0030 |
| I13224 | 147 | Asia | Pakistan | 3962.5 | 72.40 | 34.75 | 2850 | 0.0188 |
| I13227 | 147 | Asia | Pakistan | 3962.5 | 72.40 | 34.75 | 2850 | 0.0192 |
| I13228 | 147 | Asia | Pakistan | 3962.5 | 72.40 | 34.75 | 2850 | 0.0189 |
| I8997 | 147 | Asia | Pakistan | 3962.5 | 72.40 | 34.75 | 2850 | 0.0218 |
| I8998 | 147 | Asia | Pakistan | 3962.5 | 72.40 | 34.75 | 2850 | 0.0189 |
| I10523 | 151 | Asia | Pakistan | 3962.5 | 72.35 | 34.77 | 2850 | 0.0188 |
| I12146 | 151 | Asia | Pakistan | 3962.5 | 72.35 | 34.77 | 2850 | 0.0173 |
| I12445 | 151 | Asia | Pakistan | 3962.5 | 72.35 | 34.77 | 2850 | 0.0160 |
| I12451 | 151 | Asia | Pakistan | 3962.5 | 72.35 | 34.77 | 2850 | 0.0135 |
| I12454 | 151 | Asia | Pakistan | 3962.5 | 72.35 | 34.77 | 2850 | 0.0207 |
| I12460 | 151 | Asia | Pakistan | 3962.5 | 72.35 | 34.77 | 2850 | 0.0225 |
| I12462 | 151 | Asia | Pakistan | 3962.5 | 72.35 | 34.77 | 2850 | 0.0142 |
| I12464 | 151 | Asia | Pakistan | 3962.5 | 72.35 | 34.77 | 2850 | 0.0233 |
| I12470 | 151 | Asia | Pakistan | 3962.5 | 72.35 | 34.77 | 2850 | 0.0154 |
| I12477 | 151 | Asia | Pakistan | 3962.5 | 72.35 | 34.77 | 2850 | 0.0178 |
| I12968 | 151 | Asia | Pakistan | 3962.5 | 72.35 | 34.77 | 2850 | 0.0190 |
| I13219 | 151 | Asia | Pakistan | 3962.5 | 72.35 | 34.77 | 2850 | 0.0140 |
| I5398 | 151 | Asia | Pakistan | 3962.5 | 72.35 | 34.77 | 2850 | 0.0222 |
| I12971 | 175 | Asia | Mongolia | 7265.6 | 114.87 | 49.15 | 2849 | 0.0242 |
| I7032 | 267 | Asia | Mongolia | 6978.6 | 110.19 | 48.69 | 2849 | 0.0168 |
| IR1_noUDG.SG | 955 | Europe | Hungary | 3003.9 | 19.95 | 47.82 | 2849 | 0.0214 |
| I8190 | 208 | Asia | Pakistan | 3962.5 | 72.31 | 34.74 | 2847 | 0.0176 |
| I3262 | 208 | Asia | Pakistan | 3962.5 | 72.31 | 34.74 | 2844 | 0.0201 |
| I6546 | 258 | Asia | Pakistan | 3962.5 | 72.36 | 34.76 | 2841 | 0.0209 |
| I8219 | 265 | Asia | Pakistan | 3960.1 | 72.32 | 34.80 | 2841 | 0.0165 |
| I25507 | 805 | Europe | Hungary | 3098.7 | 16.79 | 47.35 | 2841 | 0.0217 |
| I14864 | 661 | Europe | United Kingdom | 4257.1 | 1.37 | 51.33 | 2839 | 0.0220 |
| I14194 | 176 | Asia | Mongolia | 7107.6 | 111.60 | 46.77 | 2834 | 0.0231 |
| I14862 | 661 | Europe | United Kingdom | 4257.1 | 1.37 | 51.33 | 2834 | 0.0166 |

| | | | | | | | | |
|---|---|---|---|---|---|---|---|---|
| F38_noUDG.SG | 115 | Asia | Iran | 1677.7 | 45.46 | 37.01 | 2833 | 0.0229 |
| R1.SG | 1061 | Europe | Italy | 2992.7 | 13.89 | 42.88 | 2832 | 0.0180 |
| I5397 | 151 | Asia | Pakistan | 3962.5 | 72.35 | 34.77 | 2829 | 0.0193 |
| I13963 | 190 | Asia | Mongolia | 6941.4 | 109.19 | 47.19 | 2829 | 0.0270 |
| I5400 | 147 | Asia | Pakistan | 3962.5 | 72.40 | 34.75 | 2824 | 0.0166 |
| I16707 | 186 | Asia | Armenia | 1927.6 | 43.95 | 40.78 | 2823 | 0.0233 |
| I4357_noUDG | 115 | Asia | Iran | 1677.7 | 45.46 | 37.01 | 2821 | 0.0231 |
| I6545 | 258 | Asia | Pakistan | 3962.5 | 72.36 | 34.76 | 2821 | 0.0135 |
| DA2_noUDG.SG | 80 | Asia | Russia | 5674.0 | 90.85 | 54.68 | 2817 | 0.0331 |
| I3451 | 221 | Asia | Kazakhstan | 4598.6 | 79.37 | 45.13 | 2816 | 0.0238 |
| I8999 | 147 | Asia | Pakistan | 3962.5 | 72.40 | 34.75 | 2813 | 0.0219 |
| I6555 | 147 | Asia | Pakistan | 3962.5 | 72.40 | 34.75 | 2811 | 0.0185 |
| I6899 | 208 | Asia | Pakistan | 3962.5 | 72.31 | 34.74 | 2811 | 0.0176 |
| I14861 | 661 | Europe | United Kingdom | 4257.1 | 1.37 | 51.33 | 2810 | 0.0201 |
| I20062 | 756 | Europe | Netherlands | 4114.8 | 5.13 | 52.43 | 2810 | 0.0191 |
| I5396 | 151 | Asia | Pakistan | 3962.5 | 72.35 | 34.77 | 2809 | 0.0147 |
| I19328 | 198 | Asia | Armenia | 1990.5 | 45.20 | 40.42 | 2809 | 0.0182 |
| I25508 | 805 | Europe | Hungary | 3098.7 | 16.79 | 47.35 | 2809 | 0.0250 |
| I3315_enhanced | 840 | Europe | Spain | 4559.8 | 3.89 | 40.00 | 2809 | 0.0202 |
| I14358 | 661 | Europe | United Kingdom | 4257.1 | 1.37 | 51.33 | 2808 | 0.0187 |
| I11855 | 163 | Asia | Armenia | 1936.1 | 44.44 | 40.15 | 2806 | 0.0207 |
| DA382_noUDG.SG | 95 | Asia | Turkmenistan | 3072.7 | 61.69 | 38.72 | 2802 | 0.0261 |
| I14379 | 661 | Europe | United Kingdom | 4257.1 | 1.37 | 51.33 | 2801 | 0.0230 |
| I7233 | 932 | Europe | North Macedonia | 2286.2 | 21.87 | 41.58 | 2800 | 0.0214 |
| R1015.SG | 1064 | Europe | Italy | 3097.9 | 12.10 | 42.02 | 2800 | 0.0211 |
| I5697 | 785 | Europe | Slovenia | 2983.9 | 15.69 | 45.85 | 2799 | 0.0207 |
| I20441 | 212 | Asia | Armenia | 1990.5 | 45.13 | 40.29 | 2793 | 0.0151 |
| I6292 | 147 | Asia | Pakistan | 3962.5 | 72.40 | 34.75 | 2781 | 0.0196 |
| I6349 | 247 | Asia | Mongolia | 7305.7 | 113.85 | 45.30 | 2781 | 0.0251 |
| I6556 | 147 | Asia | Pakistan | 3962.5 | 72.40 | 34.75 | 2776 | 0.0221 |
| KSH002 | 308 | Asia | Kazakhstan | 4370.5 | 74.71 | 48.80 | 2769 | 0.0267 |

| | | | | | | | | |
|---|---|---|---|---|---|---|---|---|
| I23911 | 784 | Europe | Croatia | 2958.4 | 15.31 | 44.56 | 2765 | 0.0242 |
| I6554 | 147 | Asia | Pakistan | 3962.5 | 72.40 | 34.75 | 2758 | 0.0138 |
| DA111_noUDG.SG | 515 | Europe | Czech Republic | 3537.1 | 14.05 | 50.51 | 2752 | 0.0218 |
| I12450 | 151 | Asia | Pakistan | 3962.5 | 72.35 | 34.77 | 2751 | 0.0122 |
| I0577 | 145 | Asia | Russia | 5746.1 | 93.71 | 52.10 | 2750 | 0.0178 |
| I14765 | 195 | Asia | Turkey | 1699.7 | 43.45 | 38.34 | 2750 | 0.0170 |
| I19613 | 195 | Asia | Turkey | 1699.7 | 43.45 | 38.34 | 2750 | 0.0168 |
| I20577 | 195 | Asia | Turkey | 1699.7 | 43.45 | 38.34 | 2750 | 0.0132 |
| I18478 | 198 | Asia | Armenia | 1990.5 | 45.20 | 40.42 | 2750 | 0.0203 |
| I20180 | 757 | Europe | Bulgaria | 2071.2 | 26.33 | 41.73 | 2750 | 0.0156 |
| I20181 | 757 | Europe | Bulgaria | 2071.2 | 26.33 | 41.73 | 2750 | 0.0204 |
| I20183 | 757 | Europe | Bulgaria | 2071.2 | 26.33 | 41.73 | 2750 | 0.0138 |
| I20184 | 757 | Europe | Bulgaria | 2071.2 | 26.33 | 41.73 | 2750 | 0.0182 |
| I20185 | 757 | Europe | Bulgaria | 2071.2 | 26.33 | 41.73 | 2750 | 0.0186 |
| I20186 | 757 | Europe | Bulgaria | 2071.2 | 26.33 | 41.73 | 2750 | 0.0174 |
| I6888 | 265 | Asia | Pakistan | 3960.1 | 72.32 | 34.80 | 2747 | 0.0141 |
| I6365 | 255 | Asia | Mongolia | 6230.4 | 99.93 | 49.66 | 2744 | 0.0199 |
| I23996 | 784 | Europe | Croatia | 2958.4 | 15.31 | 44.56 | 2744 | 0.0221 |
| R11535.SG | 353 | Asia | Armenia | 1927.6 | 43.85 | 40.69 | 2736 | 0.0168 |
| I23995 | 784 | Europe | Croatia | 2958.4 | 15.31 | 44.56 | 2729 | 0.0173 |
| CNE1_noUDG | 66 | Asia | Nepal | 5215.9 | 83.67 | 28.72 | 2725 | 0.0243 |
| I18211 | 722 | Europe | Hungary | 2868.4 | 20.02 | 46.69 | 2725 | 0.0199 |
| I18213 | 722 | Europe | Hungary | 2868.4 | 20.02 | 46.69 | 2725 | 0.0220 |
| I18245 | 722 | Europe | Hungary | 2868.4 | 20.02 | 46.69 | 2725 | 0.0207 |
| I18239 | 727 | Europe | Hungary | 3003.9 | 20.19 | 47.93 | 2725 | 0.0194 |
| I18246 | 728 | Europe | Hungary | 3003.9 | 20.48 | 47.86 | 2725 | 0.0214 |
| I14054 | 188 | Asia | Armenia | 1981.1 | 44.50 | 40.92 | 2723 | 0.0200 |
| I3313 | 839 | Europe | Croatia | 2823.8 | 15.98 | 43.89 | 2722 | 0.0215 |
| VET007 | 1179 | Europe | Italy | 3205.3 | 10.97 | 42.92 | 2721 | 0.0197 |
| VET003_4 | 1179 | Europe | Italy | 3205.3 | 10.97 | 42.92 | 2705 | 0.0151 |
| I6357 | 250 | Asia | Mongolia | 6694.1 | 106.40 | 47.70 | 2700 | 0.0220 |

| | | | | | | | | |
|---|---|---|---|---|---|---|---|---|
| I6359 | 252 | Asia | Mongolia | 6229.8 | 100.05 | 50.12 | 2700 | 0.0224 |
| Kivutkalns42 | 968 | Europe | Latvia | 3770.9 | 24.27 | 56.85 | 2697 | 0.0238 |
| I5692 | 902 | Europe | Slovenia | 3130.3 | 14.50 | 46.05 | 2693 | 0.0179 |
| I14052 | 188 | Asia | Armenia | 1981.1 | 44.50 | 40.92 | 2690 | 0.0220 |
| I14053 | 188 | Asia | Armenia | 1981.1 | 44.50 | 40.92 | 2690 | 0.0200 |
| I16376 | 188 | Asia | Armenia | 1981.1 | 44.50 | 40.92 | 2690 | 0.0139 |
| AKB001 | 4 | Asia | Kazakhstan | 4370.5 | 74.76 | 48.94 | 2677 | 0.0190 |
| I5693 | 902 | Europe | Slovenia | 3130.3 | 14.50 | 46.05 | 2674 | 0.0214 |
| I10388 | 592 | Europe | North Macedonia | 2328.0 | 20.79 | 41.11 | 2658 | 0.0293 |
| I24638 | 784 | Europe | Croatia | 2958.4 | 15.31 | 44.56 | 2658 | 0.0203 |
| I24639 | 784 | Europe | Croatia | 2958.4 | 15.31 | 44.56 | 2658 | 0.0236 |
| CSP003 | 67 | Asia | Kazakhstan | 4507.0 | 78.27 | 44.49 | 2654 | 0.0177 |
| 91KLM2_noUDG | 1 | Asia | China | 8346.2 | 125.41 | 42.50 | 2650 | 0.0236 |
| R1_noUDG | 352 | Asia | Nepal | 5215.9 | 83.83 | 28.97 | 2650 | 0.0206 |
| R2_R7_noUDG | 352 | Asia | Nepal | 5215.9 | 83.83 | 28.97 | 2650 | 0.0202 |
| R5_noUDG | 352 | Asia | Nepal | 5215.9 | 83.83 | 28.97 | 2650 | 0.0187 |
| R8_noUDG | 352 | Asia | Nepal | 5215.9 | 83.83 | 28.97 | 2650 | 0.0200 |
| I26742 | 818 | Europe | Croatia | 2823.8 | 16.52 | 43.71 | 2650 | 0.0246 |
| I23904 | 784 | Europe | Croatia | 2958.4 | 15.31 | 44.56 | 2642 | 0.0224 |
| Kivutkalns25 | 968 | Europe | Latvia | 3770.9 | 24.27 | 56.85 | 2637 | 0.0244 |
| I14980 | 677 | Europe | Czech Republic | 3537.1 | 14.06 | 50.51 | 2625 | 0.0266 |
| I14983 | 677 | Europe | Czech Republic | 3537.1 | 14.06 | 50.51 | 2625 | 0.0272 |
| I20444 | 212 | Asia | Armenia | 1990.5 | 45.13 | 40.29 | 2622 | 0.0227 |
| TAL004 | 395 | Asia | Kazakhstan | 4459.0 | 75.85 | 49.09 | 2619 | 0.0241 |
| TAL005 | 395 | Asia | Kazakhstan | 4459.0 | 75.85 | 49.09 | 2618 | 0.0238 |
| MAG001 | 997 | Europe | Italy | 3193.2 | 11.33 | 42.57 | 2617 | 0.0202 |
| KYZ002 | 309 | Asia | Kazakhstan | 4532.2 | 75.91 | 50.42 | 2616 | 0.0238 |
| Kivutkalns209 | 968 | Europe | Latvia | 3770.9 | 24.27 | 56.85 | 2616 | 0.0134 |
| I17184 | 206 | Asia | Armenia | 1927.6 | 43.84 | 40.87 | 2614 | 0.0157 |
| I8112 | 591 | Europe | North Macedonia | 2243.0 | 22.49 | 41.28 | 2614 | 0.0221 |
| I12106 | 618 | Europe | Slovakia | 3095.6 | 18.22 | 47.81 | 2614 | 0.0204 |

| | | | | | | | | |
|---|---|---|---|---|---|---|---|---|
| VET002 | 1179 | Europe | Italy | 3205.3 | 10.97 | 42.92 | 2614 | 0.0224 |
| ESZ001 | 114 | Asia | Kazakhstan | 4620.1 | 77.82 | 49.52 | 2612 | 0.0264 |
| I13682 | 644 | Europe | United Kingdom | 4439.0 | -2.41 | 51.26 | 2612 | 0.0214 |
| PRZ001 | 1057 | Europe | Italy | 3122.6 | 11.95 | 43.03 | 2610 | 0.0188 |
| I16191 | 188 | Asia | Armenia | 1981.1 | 44.50 | 40.92 | 2608 | 0.0191 |
| I10383 | 591 | Europe | North Macedonia | 2243.0 | 22.49 | 41.28 | 2607 | 0.0250 |
| I5691 | 901 | Europe | Slovenia | 3037.4 | 15.16 | 45.81 | 2605 | 0.0212 |
| KSH001 | 308 | Asia | Kazakhstan | 4370.5 | 74.71 | 48.80 | 2603 | 0.0252 |
| JK2911 | 431 | Both | Egypt | 97.7 | 31.20 | 29.90 | 2602 | 0.0240 |
| ESZ003 | 114 | Asia | Kazakhstan | 4620.1 | 77.82 | 49.52 | 2600 | 0.0244 |
| I19322 | 198 | Asia | Armenia | 1990.5 | 45.20 | 40.42 | 2600 | 0.0190 |
| I23974 | 785 | Europe | Slovenia | 2983.9 | 15.69 | 45.85 | 2600 | 0.0178 |
| Kivutkalns207 | 968 | Europe | Latvia | 3770.9 | 24.27 | 56.85 | 2600 | 0.0215 |
| R1021.SG | 1065 | Europe | Italy | 3002.1 | 12.91 | 41.64 | 2600 | 0.0172 |
| R850.SG | 1118 | Europe | Italy | 3086.5 | 12.51 | 41.60 | 2600 | 0.0225 |
| R851.SG | 1118 | Europe | Italy | 3086.5 | 12.51 | 41.60 | 2600 | 0.0295 |
| I14057 | 188 | Asia | Armenia | 1981.1 | 44.50 | 40.92 | 2599 | 0.0165 |
| I17181 | 188 | Asia | Armenia | 1981.1 | 44.50 | 40.92 | 2599 | 0.0230 |
| I12783 | 633 | Europe | United Kingdom | 4467.2 | -1.69 | 51.70 | 2599 | 0.0268 |
| I24879 | 798 | Europe | Croatia | 2997.9 | 15.32 | 45.21 | 2599 | 0.0232 |
| I24882 | 799 | Europe | Croatia | 2944.0 | 16.12 | 45.17 | 2599 | 0.0197 |
| I19867 | 646 | Europe | United Kingdom | 4423.8 | -1.79 | 51.05 | 2598 | 0.0211 |
| CAM001 | 496 | Europe | Italy | 3218.0 | 11.07 | 43.41 | 2597 | 0.0221 |
| I19861 | 646 | Europe | United Kingdom | 4423.8 | -1.79 | 51.05 | 2597 | 0.0212 |
| NUR002 | 345 | Asia | Kazakhstan | 4385.2 | 75.16 | 49.11 | 2596 | 0.0189 |
| CAM003 | 496 | Europe | Italy | 3218.0 | 11.07 | 43.41 | 2594 | 0.0177 |
| I14644 | 194 | Asia | Turkey | 1441.0 | 41.01 | 37.79 | 2593 | 0.0164 |
| I5698 | 776 | Europe | Slovenia | 3076.8 | 14.99 | 46.14 | 2593 | 0.0221 |
| BSB001 | 38 | Asia | Kazakhstan | 3427.7 | 58.94 | 48.24 | 2592 | 0.0220 |
| I5769 | 602 | Europe | Bulgaria | 2209.4 | 25.88 | 43.16 | 2592 | 0.0182 |
| I27382 | 823 | Europe | United Kingdom | 4481.2 | -2.61 | 50.71 | 2592 | 0.0209 |

| | | | | | | | | |
|---|---|---|---|---|---|---|---|---|
| I10379 | 590 | Europe | North Macedonia | 2383.7 | 21.64 | 42.03 | 2591 | 0.0217 |
| I10385 | 592 | Europe | North Macedonia | 2328.0 | 20.79 | 41.11 | 2591 | 0.0213 |
| I13688 | 646 | Europe | United Kingdom | 4423.8 | -1.79 | 51.05 | 2589 | 0.0220 |
| BKT001 | 28 | Asia | Kazakhstan | 4331.2 | 74.77 | 47.41 | 2587 | 0.0272 |
| I16117 | 188 | Asia | Armenia | 1981.1 | 44.50 | 40.92 | 2586 | 0.0110 |
| I11683 | 614 | Europe | Hungary | 3003.9 | 20.68 | 47.82 | 2586 | 0.0242 |
| I5727 | 903 | Europe | Croatia | 3023.7 | 15.70 | 45.90 | 2586 | 0.0172 |
| BIR013 | 26 | Asia | Kazakhstan | 4561.2 | 75.72 | 51.14 | 2585 | 0.0213 |
| KZL001 | 311 | Asia | Kazakhstan | 4356.7 | 75.36 | 48.28 | 2585 | 0.0189 |
| ESZ002 | 114 | Asia | Kazakhstan | 4620.1 | 77.82 | 49.52 | 2583 | 0.0221 |
| I4338_noUDG | 115 | Asia | Iran | 1677.7 | 45.46 | 37.01 | 2582 | 0.0200 |
| I14764 | 195 | Asia | Turkey | 1699.7 | 43.45 | 38.34 | 2582 | 0.0202 |
| I19868 | 646 | Europe | United Kingdom | 4423.8 | -1.79 | 51.05 | 2579 | 0.0198 |
| PRZ002 | 1057 | Europe | Italy | 3122.6 | 11.95 | 43.03 | 2579 | 0.0227 |
| I11719 | 618 | Europe | Slovakia | 3095.6 | 18.22 | 47.81 | 2576 | 0.0212 |
| I16163 | 688 | Europe | Italy | 3570.8 | 8.32 | 40.63 | 2576 | 0.0156 |
| C3345 | 55 | Asia | China | 5172.4 | 85.87 | 47.44 | 2574 | 0.0294 |
| C3350 | 55 | Asia | China | 5172.4 | 85.87 | 47.44 | 2574 | 0.0188 |
| C3351 | 55 | Asia | China | 5172.4 | 85.87 | 47.44 | 2574 | 0.0226 |
| C3353 | 55 | Asia | China | 5172.4 | 85.87 | 47.44 | 2574 | 0.0205 |
| I5690 | 900 | Europe | Slovenia | 3076.8 | 15.47 | 45.91 | 2574 | 0.0231 |
| DA195_noUDG.SG | 518 | Europe | Hungary | 2868.4 | 20.16 | 46.34 | 2572 | 0.0211 |
| I19862 | 646 | Europe | United Kingdom | 4423.8 | -1.79 | 51.05 | 2571 | 0.0213 |
| DA13_noUDG.SG | 74 | Asia | Kazakhstan | 4459.0 | 75.50 | 49.06 | 2568 | 0.0245 |
| scy009_noUDG.SG | 1134 | Europe | Ukraine | 2503.8 | 29.47 | 46.32 | 2568 | 0.0271 |
| I17180 | 188 | Asia | Armenia | 1981.1 | 44.50 | 40.92 | 2566 | 0.0139 |
| I8220 | 265 | Asia | Pakistan | 3960.1 | 72.32 | 34.80 | 2566 | 0.0191 |
| I18159 | 209 | Asia | Armenia | 1957.0 | 46.23 | 39.43 | 2565 | 0.0187 |
| I18160 | 209 | Asia | Armenia | 1957.0 | 46.23 | 39.43 | 2565 | 0.0197 |
| I18161 | 209 | Asia | Armenia | 1957.0 | 46.23 | 39.43 | 2565 | 0.0187 |
| I18236 | 209 | Asia | Armenia | 1957.0 | 46.23 | 39.43 | 2565 | 0.0192 |

| | | | | | | | | |
|---|---|---|---|---|---|---|---|---|
| I18237 | 209 | Asia | Armenia | 1957.0 | 46.23 | 39.43 | 2565 | 0.0195 |
| I18238 | 209 | Asia | Armenia | 1957.0 | 46.23 | 39.43 | 2565 | 0.0152 |
| I20224 | 758 | Europe | Turkey | 1573.9 | 28.04 | 37.34 | 2565 | 0.0113 |
| I20225 | 758 | Europe | Turkey | 1573.9 | 28.04 | 37.34 | 2565 | 0.0141 |
| I20226 | 758 | Europe | Turkey | 1573.9 | 28.04 | 37.34 | 2565 | 0.0202 |
| I20227 | 758 | Europe | Turkey | 1573.9 | 28.04 | 37.34 | 2565 | 0.0140 |
| I20228 | 758 | Europe | Turkey | 1573.9 | 28.04 | 37.34 | 2565 | 0.0163 |
| I20229 | 758 | Europe | Turkey | 1573.9 | 28.04 | 37.34 | 2565 | 0.0200 |
| I20230 | 758 | Europe | Turkey | 1573.9 | 28.04 | 37.34 | 2565 | 0.0108 |
| I20231 | 758 | Europe | Turkey | 1573.9 | 28.04 | 37.34 | 2565 | 0.0137 |
| I20232 | 758 | Europe | Turkey | 1573.9 | 28.04 | 37.34 | 2565 | 0.0168 |
| I20233 | 758 | Europe | Turkey | 1573.9 | 28.04 | 37.34 | 2565 | 0.0153 |
| I20257 | 758 | Europe | Turkey | 1573.9 | 28.04 | 37.34 | 2565 | 0.0158 |
| I20258 | 758 | Europe | Turkey | 1573.9 | 28.04 | 37.34 | 2565 | 0.0144 |
| DA221_noUDG.SG | 83 | Asia | Kazakhstan | 4328.5 | 76.41 | 43.18 | 2563 | 0.0226 |
| Kivutkalns215 | 968 | Europe | Latvia | 3770.9 | 24.27 | 56.85 | 2559 | 0.0203 |
| I1153 | 158 | Asia | Kazakhstan | 3577.2 | 57.72 | 50.67 | 2554 | 0.0209 |
| I6369 | 257 | Asia | Mongolia | 7007.6 | 110.32 | 47.38 | 2554 | 0.0215 |
| I8209 | 599 | Europe | Spain | 3994.5 | 3.11 | 42.13 | 2554 | 0.0233 |
| I12774 | 631 | Europe | United Kingdom | 4557.3 | -1.64 | 53.08 | 2554 | 0.0237 |
| I5725 | 903 | Europe | Croatia | 3023.7 | 15.70 | 45.90 | 2554 | 0.0178 |
| DA16_noUDG.SG | 75 | Asia | Kazakhstan | 4281.5 | 74.43 | 48.49 | 2553 | 0.0267 |
| KKM001 | 298 | Asia | Kazakhstan | 4668.6 | 79.27 | 48.94 | 2550 | 0.0172 |
| KSH003 | 308 | Asia | Kazakhstan | 4370.5 | 74.71 | 48.80 | 2550 | 0.0212 |
| KSH004 | 308 | Asia | Kazakhstan | 4370.5 | 74.71 | 48.80 | 2550 | 0.0205 |
| TAL003 | 395 | Asia | Kazakhstan | 4459.0 | 75.85 | 49.09 | 2550 | 0.0191 |
| I10384 | 592 | Europe | North Macedonia | 2328.0 | 20.79 | 41.11 | 2550 | 0.0224 |
| I10387 | 592 | Europe | North Macedonia | 2328.0 | 20.79 | 41.11 | 2550 | 0.0187 |
| I13170 | 640 | Europe | Montenegro | 2535.7 | 18.74 | 42.38 | 2550 | 0.0158 |
| I17260 | 710 | Europe | United Kingdom | 4352.6 | -1.49 | 51.14 | 2550 | 0.0257 |
| I20585 | 752 | Europe | United Kingdom | 4397.4 | -1.31 | 51.80 | 2550 | 0.0170 |

| | | | | | | | | |
|---|---|---|---|---|---|---|---|---|
| I25525 | 808 | Europe | Hungary | 3049.5 | 19.37 | 47.87 | 2550 | 0.0266 |
| I5689 | 900 | Europe | Slovenia | 3076.8 | 15.47 | 45.91 | 2540 | 0.0262 |
| CHK003 | 64 | Asia | Kyrgyzstan | 4490.6 | 78.38 | 42.49 | 2539 | 0.0245 |
| C1711 | 50 | Asia | China | 5177.6 | 86.36 | 48.08 | 2538 | 0.0208 |
| C1713 | 50 | Asia | China | 5177.6 | 86.36 | 48.08 | 2538 | 0.0207 |
| WAR001 | 410 | Asia | Kazakhstan | 4281.5 | 74.46 | 48.43 | 2537 | 0.0236 |
| I13689 | 646 | Europe | United Kingdom | 4423.8 | -1.79 | 51.05 | 2531 | 0.0165 |
| I8215 | 599 | Europe | Spain | 3994.5 | 3.11 | 42.13 | 2526 | 0.0227 |
| DAR002 | 104 | Asia | Mongolia | 6694.1 | 106.75 | 47.91 | 2525 | 0.0242 |
| I11717 | 618 | Europe | Slovakia | 3095.6 | 18.22 | 47.81 | 2525 | 0.0236 |
| I11721 | 618 | Europe | Slovakia | 3095.6 | 18.22 | 47.81 | 2525 | 0.0235 |
| I11722 | 618 | Europe | Slovakia | 3095.6 | 18.22 | 47.81 | 2525 | 0.0228 |
| I12097 | 618 | Europe | Slovakia | 3095.6 | 18.22 | 47.81 | 2525 | 0.0161 |
| I12099 | 618 | Europe | Slovakia | 3095.6 | 18.22 | 47.81 | 2525 | 0.0223 |
| I12103 | 618 | Europe | Slovakia | 3095.6 | 18.22 | 47.81 | 2525 | 0.0207 |
| I12105 | 618 | Europe | Slovakia | 3095.6 | 18.22 | 47.81 | 2525 | 0.0211 |
| I12107 | 618 | Europe | Slovakia | 3095.6 | 18.22 | 47.81 | 2525 | 0.0198 |
| I12110 | 618 | Europe | Slovakia | 3095.6 | 18.22 | 47.81 | 2525 | 0.0220 |
| I5287 | 618 | Europe | Slovakia | 3095.6 | 18.22 | 47.81 | 2525 | 0.0170 |
| I5288 | 618 | Europe | Slovakia | 3095.6 | 18.22 | 47.81 | 2525 | 0.0187 |
| I18227 | 726 | Europe | Hungary | 3095.6 | 18.33 | 47.73 | 2525 | 0.0289 |
| I27383 | 823 | Europe | United Kingdom | 4481.2 | -2.61 | 50.71 | 2522 | 0.0226 |
| C1192 | 44 | Asia | China | 4223.0 | 75.22 | 37.77 | 2515 | 0.0166 |
| C1194 | 44 | Asia | China | 4223.0 | 75.22 | 37.77 | 2515 | 0.0210 |
| C1203 | 44 | Asia | China | 4223.0 | 75.22 | 37.77 | 2515 | 0.0228 |
| C1207 | 44 | Asia | China | 4223.0 | 75.22 | 37.77 | 2515 | 0.0246 |
| C1220 | 44 | Asia | China | 4223.0 | 75.22 | 37.77 | 2515 | 0.0132 |
| C1221 | 44 | Asia | China | 4223.0 | 75.22 | 37.77 | 2515 | 0.0254 |
| C1222 | 44 | Asia | China | 4223.0 | 75.22 | 37.77 | 2515 | 0.0216 |
| C1229 | 44 | Asia | China | 4223.0 | 75.22 | 37.77 | 2515 | 0.0159 |
| C1230 | 44 | Asia | China | 4223.0 | 75.22 | 37.77 | 2515 | 0.0190 |

| | | | | | | | | |
|---|---|---|---|---|---|---|---|---|
| C790 | 61 | Asia | China | 4825.9 | 82.46 | 43.60 | 2515 | 0.0176 |
| C791 | 61 | Asia | China | 4825.9 | 82.46 | 43.60 | 2515 | 0.0259 |
| ALN006 | 7 | Asia | Kyrgyzstan | 3974.4 | 71.50 | 41.41 | 2507 | 0.0260 |
| I14635 | 194 | Asia | Turkey | 1441.0 | 41.01 | 37.79 | 2500 | 0.0225 |
| I14734 | 194 | Asia | Turkey | 1441.0 | 41.01 | 37.79 | 2500 | 0.0220 |
| I3322 | 842 | Europe | Spain | 4335.0 | 0.43 | 40.51 | 2500 | 0.0142 |
| I16498 | 697 | Europe | United Kingdom | 4814.3 | -2.48 | 55.99 | 2495 | 0.0214 |
| I27381 | 823 | Europe | United Kingdom | 4481.2 | -2.61 | 50.71 | 2487 | 0.0179 |
| KYZ001 | 309 | Asia | Kazakhstan | 4532.2 | 75.91 | 50.42 | 2475 | 0.0225 |
| Kivutkalns222 | 968 | Europe | Latvia | 3770.9 | 24.27 | 56.85 | 2473 | 0.0179 |
| I11154 | 607 | Europe | United Kingdom | 4389.8 | 0.11 | 52.17 | 2469 | 0.0241 |
| I23978 | 776 | Europe | Slovenia | 3076.8 | 14.99 | 46.14 | 2459 | 0.0222 |
| I11995 | 623 | Europe | United Kingdom | 4423.8 | -2.32 | 51.01 | 2457 | 0.0233 |
| RISE601_noUDG.SG | 365 | Asia | Russia | 5214.5 | 85.72 | 50.21 | 2451 | 0.0219 |
| RISE602_noUDG.SG | 366 | Asia | Russia | 5224.1 | 86.46 | 50.62 | 2451 | 0.0293 |
| I14688 | 669 | Europe | Albania | 2438.6 | 20.39 | 42.02 | 2450 | 0.0206 |
| I11149 | 605 | Europe | United Kingdom | 4389.8 | 0.18 | 52.20 | 2443 | 0.0262 |
| I14381 | 661 | Europe | United Kingdom | 4257.1 | 1.37 | 51.33 | 2443 | 0.0194 |
| I17259 | 710 | Europe | United Kingdom | 4352.6 | -1.49 | 51.14 | 2438 | 0.0176 |
| I2692 | 697 | Europe | United Kingdom | 4814.3 | -2.48 | 55.99 | 2436 | 0.0214 |
| Kivutkalns19 | 968 | Europe | Latvia | 3770.9 | 24.27 | 56.85 | 2433 | 0.0206 |
| I22938 | 776 | Europe | Slovenia | 3076.8 | 14.99 | 46.14 | 2425 | 0.0257 |
| I22940 | 776 | Europe | Slovenia | 3076.8 | 14.99 | 46.14 | 2425 | 0.0258 |
| I11033 | 601 | Europe | United Kingdom | 4586.2 | -0.77 | 53.92 | 2421 | 0.0230 |
| I17258 | 710 | Europe | United Kingdom | 4352.6 | -1.49 | 51.14 | 2415 | 0.0157 |
| I13684 | 645 | Europe | United Kingdom | 4495.0 | -2.80 | 51.18 | 2408 | 0.0234 |
| I5726 | 903 | Europe | Croatia | 3023.7 | 15.70 | 45.90 | 2407 | 0.0148 |
| I14733 | 194 | Asia | Turkey | 1441.0 | 41.01 | 37.79 | 2401 | 0.0138 |
| SRK001 | 389 | Asia | Kazakhstan | 4385.2 | 75.17 | 49.21 | 2400 | 0.0212 |
| CSN006 | 512 | Europe | Italy | 3205.3 | 11.33 | 43.03 | 2396 | 0.0212 |
| I12787 | 633 | Europe | United Kingdom | 4467.2 | -1.69 | 51.70 | 2391 | 0.0259 |

| | | | | | | | | |
|---|---|---|---|---|---|---|---|---|
| SFI-34_noUDG.SG | 436 | Both | Lebanon | 776.6 | 35.51 | 33.90 | 2385 | 0.0168 |
| SFI-44_noUDG.SG | 436 | Both | Lebanon | 776.6 | 35.51 | 33.90 | 2385 | 0.0245 |
| SFI-47_noUDG.SG | 436 | Both | Lebanon | 776.6 | 35.51 | 33.90 | 2385 | 0.0140 |
| SFI-50_noUDG.SG | 436 | Both | Lebanon | 776.6 | 35.51 | 33.90 | 2385 | 0.0203 |
| I15048 | 678 | Europe | Czech Republic | 3537.1 | 14.07 | 50.41 | 2385 | 0.0268 |
| I15950 | 678 | Europe | Czech Republic | 3537.1 | 14.07 | 50.41 | 2385 | 0.0190 |
| I8214 | 599 | Europe | Spain | 3994.5 | 3.11 | 42.13 | 2375 | 0.0205 |
| I12410 | 626 | Europe | Spain | 4140.4 | 1.67 | 41.36 | 2375 | 0.0256 |
| I5723 | 903 | Europe | Croatia | 3023.7 | 15.70 | 45.90 | 2374 | 0.0175 |
| C1708 | 49 | Asia | China | 5253.4 | 86.87 | 47.70 | 2371 | 0.0239 |
| I13504 | 184 | Asia | Mongolia | 6158.1 | 99.20 | 50.70 | 2368 | 0.0255 |
| I19863 | 646 | Europe | United Kingdom | 4423.8 | -1.79 | 51.05 | 2352 | 0.0246 |
| C1206 | 44 | Asia | China | 4223.0 | 75.22 | 37.77 | 2351 | 0.0232 |
| C3319 | 54 | Asia | China | 4827.9 | 82.51 | 43.80 | 2351 | 0.0177 |
| KM4_noUDG | 300 | Asia | Nepal | 5215.9 | 84.27 | 28.73 | 2351 | 0.0208 |
| KS20_KS25_noUDG | 300 | Asia | Nepal | 5215.9 | 84.27 | 28.73 | 2351 | 0.0178 |
| KS21_KS23_KS4_noUDG | 300 | Asia | Nepal | 5215.9 | 84.27 | 28.73 | 2351 | 0.0207 |
| KS26_noUDG | 300 | Asia | Nepal | 5215.9 | 84.27 | 28.73 | 2351 | 0.0209 |
| KS5_noUDG | 300 | Asia | Nepal | 5215.9 | 84.27 | 28.73 | 2351 | 0.0220 |
| KS8_noUDG | 300 | Asia | Nepal | 5215.9 | 84.27 | 28.73 | 2351 | 0.0210 |
| KS9_noUDG | 300 | Asia | Nepal | 5215.9 | 84.27 | 28.73 | 2351 | 0.0224 |
| SGZ002 | 376 | Asia | Kazakhstan | 3313.1 | 54.45 | 49.24 | 2350 | 0.0246 |
| CSN001 | 512 | Europe | Italy | 3205.3 | 11.33 | 43.03 | 2350 | 0.0199 |
| CSN013 | 512 | Europe | Italy | 3205.3 | 11.33 | 43.03 | 2350 | 0.0215 |
| I13729 | 607 | Europe | United Kingdom | 4389.8 | 0.11 | 52.17 | 2350 | 0.0217 |
| I16090 | 689 | Europe | Czech Republic | 3495.2 | 14.32 | 50.08 | 2350 | 0.0182 |
| R435.SG | 1110 | Europe | Italy | 3002.1 | 12.78 | 41.80 | 2350 | 0.0234 |
| I14551 | 664 | Europe | United Kingdom | 4296.4 | -0.27 | 50.84 | 2349 | 0.0206 |
| I10390 | 591 | Europe | North Macedonia | 2243.0 | 22.49 | 41.28 | 2343 | 0.0205 |
| R11540.SG | 353 | Asia | Armenia | 1927.6 | 43.85 | 40.69 | 2341 | 0.0190 |
| DA129_noUDG.SG | 73 | Asia | Kazakhstan | 4328.5 | 76.23 | 43.14 | 2333 | 0.0279 |

| | | | | | | | | |
|---|---|---|---|---|---|---|---|---|
| I12772 | 632 | Europe | United Kingdom | 4624.9 | -5.00 | 50.54 | 2329 | 0.0199 |
| I16439 | 632 | Europe | United Kingdom | 4624.9 | -5.00 | 50.54 | 2329 | 0.0211 |
| I16441 | 632 | Europe | United Kingdom | 4624.9 | -5.00 | 50.54 | 2329 | 0.0155 |
| I16442 | 632 | Europe | United Kingdom | 4624.9 | -5.00 | 50.54 | 2329 | 0.0232 |
| DA194_noUDG.SG | 518 | Europe | Hungary | 2868.4 | 20.16 | 46.34 | 2328 | 0.0256 |
| R10338.SG | 1066 | Europe | Italy | 3097.9 | 11.76 | 42.25 | 2328 | 0.0251 |
| R10340.SG | 1066 | Europe | Italy | 3097.9 | 11.76 | 42.25 | 2328 | 0.0267 |
| R10343.SG | 1066 | Europe | Italy | 3097.9 | 11.76 | 42.25 | 2328 | 0.0247 |
| R10361.SG | 1066 | Europe | Italy | 3097.9 | 11.76 | 42.25 | 2328 | 0.0201 |
| I12771 | 631 | Europe | United Kingdom | 4557.3 | -1.64 | 53.08 | 2326 | 0.0227 |
| I18220 | 723 | Europe | Hungary | 2913.5 | 20.74 | 47.10 | 2312 | 0.0185 |
| DA223_noUDG.SG | 84 | Asia | Kazakhstan | 3667.1 | 68.25 | 43.07 | 2311 | 0.0239 |
| I15952 | 678 | Europe | Czech Republic | 3537.1 | 14.07 | 50.41 | 2310 | 0.0207 |
| I18259 | 729 | Europe | Hungary | 2823.1 | 20.06 | 46.28 | 2310 | 0.0182 |
| I15954 | 678 | Europe | Czech Republic | 3537.1 | 14.07 | 50.41 | 2305 | 0.0196 |
| I14100 | 601 | Europe | United Kingdom | 4586.2 | -0.77 | 53.92 | 2302 | 0.0126 |
| DA50_noUDG.SG | 99 | Asia | Kyrgyzstan | 4229.4 | 75.33 | 41.43 | 2300 | 0.0207 |
| I0562 | 143 | Asia | Kazakhstan | 5197.6 | 86.35 | 49.34 | 2300 | 0.0193 |
| I0563 | 143 | Asia | Kazakhstan | 5197.6 | 86.35 | 49.34 | 2300 | 0.0248 |
| SHD001 | 378 | Asia | Russia | 4351.6 | 63.38 | 56.50 | 2300 | 0.0194 |
| DA139_noUDG.SG | 516 | Europe | Russia | 2641.2 | 39.55 | 47.29 | 2300 | 0.0241 |
| I11699 | 615 | Europe | Austria | 3232.5 | 15.70 | 48.24 | 2300 | 0.0234 |
| I11701 | 615 | Europe | Austria | 3232.5 | 15.70 | 48.24 | 2300 | 0.0219 |
| I11708 | 615 | Europe | Austria | 3232.5 | 15.70 | 48.24 | 2300 | 0.0238 |
| KBO001 | 290 | Asia | Kazakhstan | 4561.8 | 76.91 | 50.18 | 2297 | 0.0169 |
| SMV001 | 387 | Asia | Russia | 4284.5 | 64.86 | 55.19 | 2295 | 0.0263 |
| I18226 | 725 | Europe | Hungary | 2958.8 | 21.37 | 47.94 | 2295 | 0.0216 |
| I16119 | 163 | Asia | Armenia | 1936.1 | 44.44 | 40.15 | 2286 | 0.0162 |
| I12970 | 174 | Asia | Mongolia | 5649.6 | 92.05 | 49.96 | 2286 | 0.0305 |
| I7717 | 258 | Asia | Pakistan | 3962.5 | 72.36 | 34.76 | 2283 | 0.0157 |
| I16597 | 701 | Europe | United Kingdom | 4343.0 | -0.28 | 52.08 | 2283 | 0.0250 |

| | | | | | | | | |
|---|---|---|---|---|---|---|---|---|
| I7027 | 174 | Asia | Mongolia | 5649.6 | 92.05 | 49.96 | 2278 | 0.0173 |
| M2113_noUDG | 321 | Asia | Nepal | 5215.9 | 83.92 | 28.86 | 2275 | 0.0247 |
| M295_noUDG | 321 | Asia | Nepal | 5215.9 | 83.92 | 28.86 | 2275 | 0.0202 |
| M354_noUDG | 321 | Asia | Nepal | 5215.9 | 83.92 | 28.86 | 2275 | 0.0302 |
| M368_noUDG | 321 | Asia | Nepal | 5215.9 | 83.92 | 28.86 | 2275 | 0.0202 |
| M4580_noUDG | 321 | Asia | Nepal | 5215.9 | 83.92 | 28.86 | 2275 | 0.0195 |
| M4681_noUDG | 321 | Asia | Nepal | 5215.9 | 83.92 | 28.86 | 2275 | 0.0278 |
| M63_M339_M359_noUDG | 321 | Asia | Nepal | 5215.9 | 83.92 | 28.86 | 2275 | 0.0199 |
| I19872 | 643 | Europe | United Kingdom | 4257.1 | 1.34 | 51.36 | 2275 | 0.0158 |
| I5696 | 785 | Europe | Slovenia | 2983.9 | 15.69 | 45.85 | 2272 | 0.0220 |
| I13965 | 191 | Asia | Mongolia | 5339.3 | 88.38 | 48.68 | 2270 | 0.0227 |
| I14984 | 678 | Europe | Czech Republic | 3537.1 | 14.07 | 50.41 | 2270 | 0.0167 |
| I13732 | 643 | Europe | United Kingdom | 4257.1 | 1.34 | 51.36 | 2269 | 0.0217 |
| I21180 | 752 | Europe | United Kingdom | 4397.4 | -1.31 | 51.80 | 2267 | 0.0153 |
| I13685 | 604 | Europe | United Kingdom | 4509.5 | -2.89 | 51.31 | 2265 | 0.0167 |
| TAQ023 | 1157 | Europe | Italy | 3097.9 | 11.77 | 42.25 | 2265 | 0.0219 |
| I14378 | 661 | Europe | United Kingdom | 4257.1 | 1.37 | 51.33 | 2264 | 0.0193 |
| I21293 | 701 | Europe | United Kingdom | 4343.0 | -0.28 | 52.08 | 2263 | 0.0241 |
| I13717 | 648 | Europe | United Kingdom | 4352.6 | -1.36 | 51.15 | 2259 | 0.0149 |
| I2983 | 834 | Europe | United Kingdom | 5189.1 | -3.27 | 58.96 | 2257 | 0.0229 |
| C1368 | 45 | Asia | China | 4909.4 | 82.64 | 43.79 | 2256 | 0.0222 |
| C1369 | 45 | Asia | China | 4909.4 | 82.64 | 43.79 | 2256 | 0.0230 |
| I15046 | 678 | Europe | Czech Republic | 3537.1 | 14.07 | 50.41 | 2256 | 0.0216 |
| I20752 | 764 | Europe | Hungary | 2913.5 | 20.26 | 47.21 | 2256 | 0.0229 |
| I14986 | 678 | Europe | Czech Republic | 3537.1 | 14.07 | 50.41 | 2255 | 0.0188 |
| I14987 | 678 | Europe | Czech Republic | 3537.1 | 14.07 | 50.41 | 2255 | 0.0198 |
| I15045 | 678 | Europe | Czech Republic | 3537.1 | 14.07 | 50.41 | 2255 | 0.0177 |
| I17145 | 678 | Europe | Czech Republic | 3537.1 | 14.07 | 50.41 | 2255 | 0.0265 |
| C3363 | 57 | Asia | China | 4908.0 | 83.28 | 43.54 | 2253 | 0.0172 |
| R10363.SG | 1066 | Europe | Italy | 3097.9 | 11.76 | 42.25 | 2253 | 0.0221 |
| C3320 | 54 | Asia | China | 4827.9 | 82.51 | 43.80 | 2252 | 0.0226 |

| | | | | | | | | |
|---|---|---|---|---|---|---|---|---|
| I0575 | 144 | Asia | Russia | 3524.3 | 53.87 | 51.99 | 2250 | 0.0214 |
| I6356 | 174 | Asia | Mongolia | 5649.6 | 92.05 | 49.96 | 2250 | 0.0217 |
| I6894 | 258 | Asia | Pakistan | 3962.5 | 72.36 | 34.76 | 2250 | 0.0168 |
| I7718 | 258 | Asia | Pakistan | 3962.5 | 72.36 | 34.76 | 2250 | 0.0137 |
| I7719 | 258 | Asia | Pakistan | 3962.5 | 72.36 | 34.76 | 2250 | 0.0229 |
| I7720 | 258 | Asia | Pakistan | 3962.5 | 72.36 | 34.76 | 2250 | 0.0184 |
| I7721 | 258 | Asia | Pakistan | 3962.5 | 72.36 | 34.76 | 2250 | 0.0167 |
| I7722 | 258 | Asia | Pakistan | 3962.5 | 72.36 | 34.76 | 2250 | 0.0155 |
| I7723 | 258 | Asia | Pakistan | 3962.5 | 72.36 | 34.76 | 2250 | 0.0214 |
| LGM79_noUDG | 317 | Asia | China | 7746.6 | 113.92 | 33.87 | 2250 | 0.0257 |
| SMV002 | 387 | Asia | Russia | 4284.5 | 64.86 | 55.19 | 2250 | 0.0174 |
| I10391 | 591 | Europe | North Macedonia | 2243.0 | 22.49 | 41.28 | 2250 | 0.0219 |
| I10392 | 591 | Europe | North Macedonia | 2243.0 | 22.49 | 41.28 | 2250 | 0.0183 |
| I13780 | 609 | Europe | Czech Republic | 3495.2 | 14.36 | 50.05 | 2250 | 0.0205 |
| I16268 | 609 | Europe | Czech Republic | 3495.2 | 14.36 | 50.05 | 2250 | 0.0149 |
| I16271 | 609 | Europe | Czech Republic | 3495.2 | 14.36 | 50.05 | 2250 | 0.0197 |
| I17322 | 609 | Europe | Czech Republic | 3495.2 | 14.36 | 50.05 | 2250 | 0.0181 |
| I13615 | 643 | Europe | United Kingdom | 4257.1 | 1.34 | 51.36 | 2250 | 0.0177 |
| I19873 | 643 | Europe | United Kingdom | 4257.1 | 1.34 | 51.36 | 2250 | 0.0208 |
| I19907 | 643 | Europe | United Kingdom | 4257.1 | 1.34 | 51.36 | 2250 | 0.0147 |
| I19908 | 643 | Europe | United Kingdom | 4257.1 | 1.34 | 51.36 | 2250 | 0.0184 |
| I19910 | 643 | Europe | United Kingdom | 4257.1 | 1.34 | 51.36 | 2250 | 0.0171 |
| I19911 | 643 | Europe | United Kingdom | 4257.1 | 1.34 | 51.36 | 2250 | 0.0228 |
| I16405 | 692 | Europe | United Kingdom | 4509.5 | -3.42 | 51.41 | 2250 | 0.0237 |
| I19211 | 752 | Europe | United Kingdom | 4397.4 | -1.31 | 51.80 | 2250 | 0.0120 |
| I20586 | 752 | Europe | United Kingdom | 4397.4 | -1.31 | 51.80 | 2250 | 0.0223 |
| I20589 | 752 | Europe | United Kingdom | 4397.4 | -1.31 | 51.80 | 2250 | 0.0205 |
| I21178 | 752 | Europe | United Kingdom | 4397.4 | -1.31 | 51.80 | 2250 | 0.0222 |
| I21181 | 752 | Europe | United Kingdom | 4397.4 | -1.31 | 51.80 | 2250 | 0.0137 |
| I21182 | 752 | Europe | United Kingdom | 4397.4 | -1.31 | 51.80 | 2250 | 0.0229 |
| I19916 | 755 | Europe | France | 3717.6 | 5.27 | 43.53 | 2250 | 0.0163 |

| | | | | | | | | |
|---|---|---|---|---|---|---|---|---|
| I19917 | 755 | Europe | France | 3717.6 | 5.27 | 43.53 | 2250 | 0.0203 |
| I19918 | 755 | Europe | France | 3717.6 | 5.27 | 43.53 | 2250 | 0.0154 |
| I2982 | 834 | Europe | United Kingdom | 5189.1 | -3.27 | 58.96 | 2250 | 0.0191 |
| R437.SG | 1110 | Europe | Italy | 3002.1 | 12.78 | 41.80 | 2250 | 0.0153 |
| ORC002 | 1032 | Europe | Italy | 3620.3 | 9.43 | 39.67 | 2249 | 0.0257 |
| C3362 | 57 | Asia | China | 4908.0 | 83.28 | 43.54 | 2248 | 0.0151 |
| I13731 | 643 | Europe | United Kingdom | 4257.1 | 1.34 | 51.36 | 2246 | 0.0222 |
| C3324 | 54 | Asia | China | 4827.9 | 82.51 | 43.80 | 2245 | 0.0293 |
| I19042 | 747 | Europe | United Kingdom | 4352.6 | -1.29 | 51.07 | 2245 | 0.0219 |
| I7022 | 174 | Asia | Mongolia | 5649.6 | 92.05 | 49.96 | 2244 | 0.0249 |
| I7024 | 174 | Asia | Mongolia | 5649.6 | 92.05 | 49.96 | 2244 | 0.0253 |
| I7030 | 174 | Asia | Mongolia | 5649.6 | 92.05 | 49.96 | 2244 | 0.0181 |
| I10366 | 588 | Europe | Italy | 3620.3 | 8.84 | 39.81 | 2244 | 0.0144 |
| I20587 | 752 | Europe | United Kingdom | 4397.4 | -1.31 | 51.80 | 2244 | 0.0213 |
| I13727 | 607 | Europe | United Kingdom | 4389.8 | 0.11 | 52.17 | 2243 | 0.0210 |
| I6891 | 258 | Asia | Pakistan | 3962.5 | 72.36 | 34.76 | 2242 | 0.0123 |
| I11147 | 604 | Europe | United Kingdom | 4509.5 | -2.89 | 51.31 | 2241 | 0.0187 |
| I14807 | 672 | Europe | United Kingdom | 4397.4 | -0.99 | 51.75 | 2241 | 0.0252 |
| I15040 | 678 | Europe | Czech Republic | 3537.1 | 14.07 | 50.41 | 2240 | 0.0193 |
| I15043 | 678 | Europe | Czech Republic | 3537.1 | 14.07 | 50.41 | 2240 | 0.0210 |
| I18488 | 734 | Europe | Hungary | 3142.3 | 17.30 | 47.69 | 2240 | 0.0182 |
| I18489 | 734 | Europe | Hungary | 3142.3 | 17.30 | 47.69 | 2240 | 0.0216 |
| I18490 | 734 | Europe | Hungary | 3142.3 | 17.30 | 47.69 | 2240 | 0.0182 |
| I18491 | 734 | Europe | Hungary | 3142.3 | 17.30 | 47.69 | 2240 | 0.0214 |
| I18492 | 734 | Europe | Hungary | 3142.3 | 17.30 | 47.69 | 2240 | 0.0272 |
| I18493 | 734 | Europe | Hungary | 3142.3 | 17.30 | 47.69 | 2240 | 0.0189 |
| TAQ007 | 1157 | Europe | Italy | 3097.9 | 11.77 | 42.25 | 2240 | 0.0135 |
| I6232 | 174 | Asia | Mongolia | 5649.6 | 92.05 | 49.96 | 2239 | 0.0206 |
| I16272 | 609 | Europe | Czech Republic | 3495.2 | 14.36 | 50.05 | 2239 | 0.0140 |
| I18183 | 721 | Europe | Hungary | 2958.7 | 19.91 | 47.49 | 2239 | 0.0183 |
| I7953 | 947 | Europe | Czech Republic | 3404.5 | 15.79 | 50.11 | 2239 | 0.0203 |

| | | | | | | | | |
|---|---|---|---|---|---|---|---|---|
| I12412 | 601 | Europe | United Kingdom | 4586.2 | -0.77 | 53.92 | 2238 | 0.0234 |
| I22056 | 659 | Europe | United Kingdom | 4552.7 | -0.29 | 54.13 | 2238 | 0.0208 |
| I13730 | 643 | Europe | United Kingdom | 4257.1 | 1.34 | 51.36 | 2237 | 0.0229 |
| I22055 | 659 | Europe | United Kingdom | 4552.7 | -0.29 | 54.13 | 2237 | 0.0196 |
| scy301_noUDG.SG | 1135 | Europe | Moldova | 2552.7 | 29.80 | 46.67 | 2237 | 0.0186 |
| I14380 | 661 | Europe | United Kingdom | 4257.1 | 1.37 | 51.33 | 2235 | 0.0250 |
| I15951 | 678 | Europe | Czech Republic | 3537.1 | 14.07 | 50.41 | 2235 | 0.0178 |
| I4996 | 805 | Europe | Hungary | 3098.7 | 16.79 | 47.35 | 2235 | 0.0216 |
| AIG003 | 3 | Asia | Kazakhstan | 2660.5 | 51.33 | 43.70 | 2234 | 0.0220 |
| I19914 | 643 | Europe | United Kingdom | 4257.1 | 1.34 | 51.36 | 2233 | 0.0177 |
| I14804 | 672 | Europe | United Kingdom | 4397.4 | -0.99 | 51.75 | 2233 | 0.0175 |
| I20583 | 761 | Europe | United Kingdom | 4397.4 | -1.42 | 51.75 | 2233 | 0.0093 |
| I5724 | 903 | Europe | Croatia | 3023.7 | 15.70 | 45.90 | 2232 | 0.0249 |
| I16592 | 604 | Europe | United Kingdom | 4509.5 | -2.89 | 51.31 | 2231 | 0.0215 |
| I19656 | 622 | Europe | United Kingdom | 4495.0 | -2.74 | 50.95 | 2231 | 0.0206 |
| I14860 | 661 | Europe | United Kingdom | 4257.1 | 1.37 | 51.33 | 2231 | 0.0205 |
| I21275 | 761 | Europe | United Kingdom | 4397.4 | -1.42 | 51.75 | 2231 | 0.0155 |
| BIY003 | 27 | Asia | Russia | 4662.7 | 73.37 | 54.99 | 2230 | 0.0178 |
| I12778 | 631 | Europe | United Kingdom | 4557.3 | -1.64 | 53.08 | 2230 | 0.0230 |
| I15044 | 678 | Europe | Czech Republic | 3537.1 | 14.07 | 50.41 | 2230 | 0.0249 |
| I18530 | 719 | Europe | Hungary | 3142.3 | 17.63 | 47.67 | 2230 | 0.0179 |
| C1673 | 47 | Asia | China | 4741.8 | 81.52 | 43.08 | 2229 | 0.0183 |
| DA51_noUDG.SG | 99 | Asia | Kyrgyzstan | 4229.4 | 75.33 | 41.43 | 2229 | 0.0164 |
| I0247 | 129 | Asia | Russia | 3443.1 | 51.16 | 52.43 | 2229 | 0.0290 |
| I11156 | 608 | Europe | United Kingdom | 4405.4 | -0.18 | 52.56 | 2229 | 0.0218 |
| I21276 | 761 | Europe | United Kingdom | 4397.4 | -1.42 | 51.75 | 2229 | 0.0236 |
| VET008 | 1179 | Europe | Italy | 3205.3 | 10.97 | 42.92 | 2229 | 0.0183 |
| I14800 | 672 | Europe | United Kingdom | 4397.4 | -0.99 | 51.75 | 2228 | 0.0176 |
| I19044 | 748 | Europe | United Kingdom | 4389.8 | 0.18 | 52.21 | 2228 | 0.0219 |
| I19046 | 748 | Europe | United Kingdom | 4389.8 | 0.18 | 52.21 | 2228 | 0.0227 |
| I21179 | 752 | Europe | United Kingdom | 4397.4 | -1.31 | 51.80 | 2228 | 0.0209 |

| | | | | | | | | |
|---|---|---|---|---|---|---|---|---|
| I6896 | 258 | Asia | Pakistan | 3962.5 | 72.36 | 34.76 | 2227 | 0.0225 |
| I17014 | 604 | Europe | United Kingdom | 4509.5 | -2.89 | 51.31 | 2227 | 0.0226 |
| I17015 | 604 | Europe | United Kingdom | 4509.5 | -2.89 | 51.31 | 2227 | 0.0234 |
| I13728 | 607 | Europe | United Kingdom | 4389.8 | 0.11 | 52.17 | 2227 | 0.0202 |
| I11997 | 608 | Europe | United Kingdom | 4405.4 | -0.18 | 52.56 | 2227 | 0.0278 |
| I19909 | 643 | Europe | United Kingdom | 4257.1 | 1.34 | 51.36 | 2227 | 0.0222 |
| I14859 | 661 | Europe | United Kingdom | 4257.1 | 1.37 | 51.33 | 2227 | 0.0165 |
| BIR010 | 25 | Asia | Kazakhstan | 4561.2 | 75.70 | 51.14 | 2226 | 0.0267 |
| CHN001 | 65 | Asia | Mongolia | 5649.6 | 92.05 | 50.00 | 2226 | 0.0173 |
| I7023 | 174 | Asia | Mongolia | 5649.6 | 92.05 | 49.96 | 2226 | 0.0143 |
| SGZ001 | 376 | Asia | Kazakhstan | 3313.1 | 54.45 | 49.24 | 2226 | 0.0199 |
| I17016 | 604 | Europe | United Kingdom | 4509.5 | -2.89 | 51.31 | 2226 | 0.0237 |
| I14866 | 661 | Europe | United Kingdom | 4257.1 | 1.37 | 51.33 | 2226 | 0.0154 |
| I3758 | 851 | Europe | Spain | 4402.7 | -2.59 | 42.56 | 2226 | 0.0213 |
| VET005 | 1179 | Europe | Italy | 3205.3 | 10.97 | 42.92 | 2226 | 0.0192 |
| BDY001 | 21 | Asia | Kazakhstan | 4385.2 | 75.12 | 49.33 | 2225 | 0.0232 |
| HJTM115_noUDG | 122 | Asia | China | 7746.6 | 114.02 | 33.59 | 2225 | 0.0264 |
| I6224 | 174 | Asia | Mongolia | 5649.6 | 92.05 | 49.96 | 2225 | 0.0253 |
| I6233 | 174 | Asia | Mongolia | 5649.6 | 92.05 | 49.96 | 2225 | 0.0182 |
| I7029 | 174 | Asia | Mongolia | 5649.6 | 92.05 | 49.96 | 2225 | 0.0210 |
| I19327 | 198 | Asia | Armenia | 1990.5 | 45.20 | 40.42 | 2225 | 0.0220 |
| I6893 | 258 | Asia | Pakistan | 3962.5 | 72.36 | 34.76 | 2225 | 0.0174 |
| I12770 | 631 | Europe | United Kingdom | 4557.3 | -1.64 | 53.08 | 2225 | 0.0212 |
| I12775 | 631 | Europe | United Kingdom | 4557.3 | -1.64 | 53.08 | 2225 | 0.0189 |
| I12779 | 631 | Europe | United Kingdom | 4557.3 | -1.64 | 53.08 | 2225 | 0.0220 |
| I3014 | 631 | Europe | United Kingdom | 4557.3 | -1.64 | 53.08 | 2225 | 0.0253 |
| I14863 | 661 | Europe | United Kingdom | 4257.1 | 1.37 | 51.33 | 2225 | 0.0194 |
| I14985 | 678 | Europe | Czech Republic | 3537.1 | 14.07 | 50.41 | 2225 | 0.0187 |
| I15042 | 678 | Europe | Czech Republic | 3537.1 | 14.07 | 50.41 | 2225 | 0.0213 |
| I17264 | 712 | Europe | United Kingdom | 4352.6 | -1.53 | 51.14 | 2225 | 0.0206 |
| I17267 | 712 | Europe | United Kingdom | 4352.6 | -1.53 | 51.14 | 2225 | 0.0210 |

| | | | | | | | | |
|---|---|---|---|---|---|---|---|---|
| I20988 | 712 | Europe | United Kingdom | 4352.6 | -1.53 | 51.14 | 2225 | 0.0222 |
| I20588 | 752 | Europe | United Kingdom | 4397.4 | -1.31 | 51.80 | 2225 | 0.0175 |
| I21277 | 761 | Europe | United Kingdom | 4397.4 | -1.42 | 51.75 | 2225 | 0.0241 |
| I20623 | 762 | Europe | United Kingdom | 4557.3 | -1.74 | 53.24 | 2225 | 0.0199 |
| VET006_9 | 1179 | Europe | Italy | 3205.3 | 10.97 | 42.92 | 2225 | 0.0244 |
| BRE001 | 31 | Asia | Kazakhstan | 5197.6 | 85.79 | 49.33 | 2224 | 0.0185 |
| I14347 | 660 | Europe | United Kingdom | 4586.2 | -1.38 | 53.91 | 2224 | 0.0225 |
| I5728 | 903 | Europe | Croatia | 3023.7 | 15.70 | 45.90 | 2224 | 0.0147 |
| R10359.SG | 1066 | Europe | Italy | 3097.9 | 11.76 | 42.25 | 2224 | 0.0224 |
| BRE002 | 31 | Asia | Kazakhstan | 5197.6 | 85.79 | 49.33 | 2223 | 0.0223 |
| I13687 | 607 | Europe | United Kingdom | 4389.8 | 0.11 | 52.17 | 2222 | 0.0180 |
| I19912 | 643 | Europe | United Kingdom | 4257.1 | 1.34 | 51.36 | 2222 | 0.0232 |
| I14348 | 660 | Europe | United Kingdom | 4586.2 | -1.38 | 53.91 | 2222 | 0.0161 |
| I3759 | 851 | Europe | Spain | 4402.7 | -2.59 | 42.56 | 2222 | 0.0194 |
| C1705 | 49 | Asia | China | 5253.4 | 86.87 | 47.70 | 2221 | 0.0228 |
| I6263 | 191 | Asia | Mongolia | 5339.3 | 88.38 | 48.68 | 2221 | 0.0284 |
| I6226 | 174 | Asia | Mongolia | 5649.6 | 92.05 | 49.96 | 2220 | 0.0271 |
| I17139 | 678 | Europe | Czech Republic | 3537.1 | 14.07 | 50.41 | 2220 | 0.0149 |
| I20990 | 712 | Europe | United Kingdom | 4352.6 | -1.53 | 51.14 | 2220 | 0.0298 |
| C3333 | 54 | Asia | China | 4827.9 | 82.51 | 43.80 | 2218 | 0.0247 |
| I6231 | 174 | Asia | Mongolia | 5649.6 | 92.05 | 49.96 | 2218 | 0.0201 |
| I25524 | 807 | Europe | Hungary | 3095.6 | 18.67 | 47.73 | 2218 | 0.0248 |
| I20582 | 761 | Europe | United Kingdom | 4397.4 | -1.42 | 51.75 | 2217 | 0.0195 |
| I14801 | 672 | Europe | United Kingdom | 4397.4 | -0.99 | 51.75 | 2214 | 0.0136 |
| I18832 | 744 | Europe | Hungary | 3146.4 | 16.63 | 47.65 | 2210 | 0.0198 |
| I18834 | 744 | Europe | Hungary | 3146.4 | 16.63 | 47.65 | 2210 | 0.0214 |
| I18835 | 744 | Europe | Hungary | 3146.4 | 16.63 | 47.65 | 2210 | 0.0170 |
| I18838 | 744 | Europe | Hungary | 3146.4 | 16.63 | 47.65 | 2210 | 0.0158 |
| KEN001 | 292 | Asia | Kyrgyzstan | 4320.3 | 75.52 | 42.20 | 2202 | 0.0248 |
| I5506 | 601 | Europe | United Kingdom | 4586.2 | -0.77 | 53.92 | 2202 | 0.0193 |
| I2694 | 697 | Europe | United Kingdom | 4814.3 | -2.48 | 55.99 | 2202 | 0.0232 |

| | | | | | | | | |
|---|---|---|---|---|---|---|---|---|
| I18830 | 743 | Europe | Croatia | 2823.8 | 15.73 | 43.89 | 2201 | 0.0208 |
| I18831 | 743 | Europe | Croatia | 2823.8 | 15.73 | 43.89 | 2201 | 0.0223 |
| I11993 | 622 | Europe | United Kingdom | 4495.0 | -2.74 | 50.95 | 2200 | 0.0265 |
| I11994 | 622 | Europe | United Kingdom | 4495.0 | -2.74 | 50.95 | 2200 | 0.0247 |
| I19854 | 622 | Europe | United Kingdom | 4495.0 | -2.74 | 50.95 | 2200 | 0.0150 |
| I19855 | 622 | Europe | United Kingdom | 4495.0 | -2.74 | 50.95 | 2200 | 0.0140 |
| I12926 | 635 | Europe | United Kingdom | 4467.2 | -1.79 | 51.71 | 2200 | 0.0208 |
| I18110 | 719 | Europe | Hungary | 3142.3 | 17.63 | 47.67 | 2200 | 0.0222 |
| I18526 | 719 | Europe | Hungary | 3142.3 | 17.63 | 47.67 | 2200 | 0.0208 |
| I18527 | 719 | Europe | Hungary | 3142.3 | 17.63 | 47.67 | 2200 | 0.0229 |
| I18528 | 719 | Europe | Hungary | 3142.3 | 17.63 | 47.67 | 2200 | 0.0191 |
| I18529 | 719 | Europe | Hungary | 3142.3 | 17.63 | 47.67 | 2200 | 0.0174 |
| I18531 | 719 | Europe | Hungary | 3142.3 | 17.63 | 47.67 | 2200 | 0.0173 |
| I18839 | 719 | Europe | Hungary | 3142.3 | 17.63 | 47.67 | 2200 | 0.0153 |
| I18840 | 719 | Europe | Hungary | 3142.3 | 17.63 | 47.67 | 2200 | 0.0160 |
| I19356 | 753 | Europe | France | 3914.4 | 4.28 | 48.80 | 2200 | 0.0235 |
| I19357 | 753 | Europe | France | 3914.4 | 4.28 | 48.80 | 2200 | 0.0233 |
| I19360 | 753 | Europe | France | 3914.4 | 4.28 | 48.80 | 2200 | 0.0198 |
| I20817 | 753 | Europe | France | 3914.4 | 4.28 | 48.80 | 2200 | 0.0187 |
| I21399 | 753 | Europe | France | 3914.4 | 4.28 | 48.80 | 2200 | 0.0225 |
| I21402 | 753 | Europe | France | 3914.4 | 4.28 | 48.80 | 2200 | 0.0173 |
| I21271 | 761 | Europe | United Kingdom | 4397.4 | -1.42 | 51.75 | 2200 | 0.0223 |
| I21272 | 761 | Europe | United Kingdom | 4397.4 | -1.42 | 51.75 | 2200 | 0.0256 |
| I21274 | 761 | Europe | United Kingdom | 4397.4 | -1.42 | 51.75 | 2200 | 0.0234 |
| I14809 | 672 | Europe | United Kingdom | 4397.4 | -0.99 | 51.75 | 2199 | 0.0155 |
| I8245 | 265 | Asia | Pakistan | 3960.1 | 72.32 | 34.80 | 2197 | 0.0181 |
| I17146 | 678 | Europe | Czech Republic | 3537.1 | 14.07 | 50.41 | 2197 | 0.0155 |
| I20622 | 762 | Europe | United Kingdom | 4557.3 | -1.74 | 53.24 | 2186 | 0.0205 |
| I15039 | 678 | Europe | Czech Republic | 3537.1 | 14.07 | 50.41 | 2185 | 0.0248 |
| I15953 | 678 | Europe | Czech Republic | 3537.1 | 14.07 | 50.41 | 2185 | 0.0188 |
| I18182 | 721 | Europe | Hungary | 2958.7 | 19.91 | 47.49 | 2185 | 0.0219 |

| | | | | | | | | |
|---|---|---|---|---|---|---|---|---|
| I19657 | 622 | Europe | United Kingdom | 4495.0 | -2.74 | 50.95 | 2183 | 0.0206 |
| C2042 | 53 | Asia | China | 5552.6 | 90.75 | 46.48 | 2182 | 0.0259 |
| C1358 | 45 | Asia | China | 4909.4 | 82.64 | 43.79 | 2181 | 0.0220 |
| I11152 | 606 | Europe | United Kingdom | 4389.8 | 0.03 | 52.35 | 2179 | 0.0196 |
| I17262 | 711 | Europe | United Kingdom | 4423.8 | -1.60 | 51.14 | 2176 | 0.0219 |
| I20982 | 711 | Europe | United Kingdom | 4423.8 | -1.60 | 51.14 | 2176 | 0.0222 |
| I20983 | 711 | Europe | United Kingdom | 4423.8 | -1.60 | 51.14 | 2176 | 0.0173 |
| I20984 | 711 | Europe | United Kingdom | 4423.8 | -1.60 | 51.14 | 2176 | 0.0263 |
| I20985 | 711 | Europe | United Kingdom | 4423.8 | -1.60 | 51.14 | 2176 | 0.0213 |
| I20986 | 711 | Europe | United Kingdom | 4423.8 | -1.60 | 51.14 | 2176 | 0.0165 |
| I11034 | 601 | Europe | United Kingdom | 4586.2 | -0.77 | 53.92 | 2175 | 0.0200 |
| I12411 | 601 | Europe | United Kingdom | 4586.2 | -0.77 | 53.92 | 2175 | 0.0210 |
| I12413 | 601 | Europe | United Kingdom | 4586.2 | -0.77 | 53.92 | 2175 | 0.0260 |
| I12414 | 601 | Europe | United Kingdom | 4586.2 | -0.77 | 53.92 | 2175 | 0.0203 |
| I12415 | 601 | Europe | United Kingdom | 4586.2 | -0.77 | 53.92 | 2175 | 0.0193 |
| I13751 | 601 | Europe | United Kingdom | 4586.2 | -0.77 | 53.92 | 2175 | 0.0158 |
| I13753 | 601 | Europe | United Kingdom | 4586.2 | -0.77 | 53.92 | 2175 | 0.0181 |
| I13754 | 601 | Europe | United Kingdom | 4586.2 | -0.77 | 53.92 | 2175 | 0.0166 |
| I13756 | 601 | Europe | United Kingdom | 4586.2 | -0.77 | 53.92 | 2175 | 0.0250 |
| I13757 | 601 | Europe | United Kingdom | 4586.2 | -0.77 | 53.92 | 2175 | 0.0192 |
| I13758 | 601 | Europe | United Kingdom | 4586.2 | -0.77 | 53.92 | 2175 | 0.0148 |
| I13759 | 601 | Europe | United Kingdom | 4586.2 | -0.77 | 53.92 | 2175 | 0.0150 |
| I13760 | 601 | Europe | United Kingdom | 4586.2 | -0.77 | 53.92 | 2175 | 0.0274 |
| I14099 | 601 | Europe | United Kingdom | 4586.2 | -0.77 | 53.92 | 2175 | 0.0221 |
| I14101 | 601 | Europe | United Kingdom | 4586.2 | -0.77 | 53.92 | 2175 | 0.0204 |
| I14102 | 601 | Europe | United Kingdom | 4586.2 | -0.77 | 53.92 | 2175 | 0.0208 |
| I14104 | 601 | Europe | United Kingdom | 4586.2 | -0.77 | 53.92 | 2175 | 0.0256 |
| I14105 | 601 | Europe | United Kingdom | 4586.2 | -0.77 | 53.92 | 2175 | 0.0230 |
| I14107 | 601 | Europe | United Kingdom | 4586.2 | -0.77 | 53.92 | 2175 | 0.0225 |
| I14108 | 601 | Europe | United Kingdom | 4586.2 | -0.77 | 53.92 | 2175 | 0.0180 |
| I5505 | 601 | Europe | United Kingdom | 4586.2 | -0.77 | 53.92 | 2175 | 0.0246 |

| | | | | | | | | |
|---|---|---|---|---|---|---|---|---|
| I5508 | 601 | Europe | United Kingdom | 4586.2 | -0.77 | 53.92 | 2175 | 0.0213 |
| I20631 | 762 | Europe | United Kingdom | 4557.3 | -1.74 | 53.24 | 2175 | 0.0208 |
| I20989 | 712 | Europe | United Kingdom | 4352.6 | -1.53 | 51.14 | 2173 | 0.0216 |
| TAQ019 | 1157 | Europe | Italy | 3097.9 | 11.77 | 42.25 | 2173 | 0.0177 |
| I15047 | 678 | Europe | Czech Republic | 3537.1 | 14.07 | 50.41 | 2170 | 0.0274 |
| TAQ024 | 1157 | Europe | Italy | 3097.9 | 11.77 | 42.25 | 2167 | 0.0251 |
| I2696 | 697 | Europe | United Kingdom | 4814.3 | -2.48 | 55.99 | 2165 | 0.0196 |
| I21309 | 769 | Europe | United Kingdom | 4423.8 | -2.14 | 51.22 | 2165 | 0.0188 |
| I21313 | 770 | Europe | United Kingdom | 4439.0 | -1.84 | 51.28 | 2165 | 0.0240 |
| I0789.SG | 583 | Europe | United Kingdom | 4343.0 | 0.28 | 52.10 | 2163 | 0.0248 |
| I20584 | 761 | Europe | United Kingdom | 4397.4 | -1.42 | 51.75 | 2161 | 0.0131 |
| I13616 | 643 | Europe | United Kingdom | 4257.1 | 1.34 | 51.36 | 2153 | 0.0200 |
| TAQ004 | 1157 | Europe | Italy | 3097.9 | 11.77 | 42.25 | 2151 | 0.0232 |
| TAQ010 | 1157 | Europe | Italy | 3097.9 | 11.77 | 42.25 | 2151 | 0.0170 |
| TAQ012 | 1157 | Europe | Italy | 3097.9 | 11.77 | 42.25 | 2151 | 0.0160 |
| TAQ016 | 1157 | Europe | Italy | 3097.9 | 11.77 | 42.25 | 2151 | 0.0262 |
| BIY008 | 27 | Asia | Russia | 4662.7 | 73.37 | 54.99 | 2150 | 0.0142 |
| BIY009 | 27 | Asia | Russia | 4662.7 | 73.37 | 54.99 | 2150 | 0.0223 |
| DA227_noUDG.SG | 84 | Asia | Kazakhstan | 3667.1 | 68.25 | 43.07 | 2150 | 0.0214 |
| I8203 | 599 | Europe | Spain | 3994.5 | 3.11 | 42.13 | 2150 | 0.0348 |
| I25510 | 806 | Europe | Hungary | 3007.4 | 17.97 | 46.88 | 2150 | 0.0232 |
| I25519 | 806 | Europe | Hungary | 3007.4 | 17.97 | 46.88 | 2150 | 0.0199 |
| I25522 | 806 | Europe | Hungary | 3007.4 | 17.97 | 46.88 | 2150 | 0.0269 |
| I3320 | 841 | Europe | Spain | 4335.0 | -0.07 | 40.14 | 2150 | 0.0183 |
| I22065 | 775 | Europe | United Kingdom | 4503.2 | -0.14 | 53.73 | 2149 | 0.0208 |
| C3337 | 54 | Asia | China | 4827.9 | 82.51 | 43.80 | 2139 | 0.0165 |
| I14837 | 673 | Europe | United Kingdom | 4586.2 | -1.39 | 53.90 | 2134 | 0.0248 |
| MA2198_noUDG.SG | 994 | Europe | Turkey | 1471.3 | 33.79 | 39.35 | 2133 | 0.0208 |
| I25509 | 806 | Europe | Hungary | 3007.4 | 17.97 | 46.88 | 2131 | 0.0242 |
| I3311 | 838 | Europe | Turkey | 1663.9 | 27.44 | 37.04 | 2127 | 0.0217 |
| I3916 | 852 | Europe | Turkey | 1523.4 | 32.00 | 39.65 | 2127 | 0.0206 |

| | | | | | | | | |
|---|---|---|---|---|---|---|---|---|
| I6573 | 852 | Europe | Turkey | 1523.4 | 32.00 | 39.65 | 2127 | 0.0110 |
| I6574 | 852 | Europe | Turkey | 1523.4 | 32.00 | 39.65 | 2127 | 0.0284 |
| I13752 | 601 | Europe | United Kingdom | 4586.2 | -0.77 | 53.92 | 2121 | 0.0175 |
| I14353 | 660 | Europe | United Kingdom | 4586.2 | -1.38 | 53.91 | 2120 | 0.0167 |
| I16503 | 697 | Europe | United Kingdom | 4814.3 | -2.48 | 55.99 | 2120 | 0.0213 |
| TAQ005 | 1157 | Europe | Italy | 3097.9 | 11.77 | 42.25 | 2118 | 0.0189 |
| TAQ018 | 1157 | Europe | Italy | 3097.9 | 11.77 | 42.25 | 2118 | 0.0132 |
| I21310 | 769 | Europe | United Kingdom | 4423.8 | -2.14 | 51.22 | 2115 | 0.0197 |
| I17263 | 712 | Europe | United Kingdom | 4352.6 | -1.53 | 51.14 | 2114 | 0.0144 |
| DA38_noUDG.SG | 94 | Asia | Mongolia | 6305.4 | 101.72 | 49.27 | 2110 | 0.0172 |
| I22052 | 659 | Europe | United Kingdom | 4552.7 | -0.29 | 54.13 | 2104 | 0.0255 |
| SKT002 | 385 | Asia | Mongolia | 6305.4 | 101.42 | 49.16 | 2103 | 0.0246 |
| BRE003 | 31 | Asia | Kazakhstan | 5197.6 | 85.79 | 49.33 | 2100 | 0.0215 |
| BRE006 | 31 | Asia | Kazakhstan | 5197.6 | 85.79 | 49.33 | 2100 | 0.0266 |
| BRE010 | 31 | Asia | Kazakhstan | 5197.6 | 85.79 | 49.33 | 2100 | 0.0209 |
| ULN002 | 41 | Asia | Mongolia | 7120.2 | 111.77 | 46.28 | 2100 | 0.0169 |
| ULN006 | 41 | Asia | Mongolia | 7120.2 | 111.77 | 46.28 | 2100 | 0.0155 |
| ULN007 | 41 | Asia | Mongolia | 7120.2 | 111.77 | 46.28 | 2100 | 0.0281 |
| ULN009 | 41 | Asia | Mongolia | 7120.2 | 111.77 | 46.28 | 2100 | 0.0222 |
| ULN010 | 41 | Asia | Mongolia | 7120.2 | 111.77 | 46.28 | 2100 | 0.0218 |
| DA43_noUDG.SG | 98 | Asia | Mongolia | 6738.8 | 105.18 | 42.53 | 2100 | 0.0239 |
| LGM41_noUDG | 317 | Asia | China | 7746.6 | 113.92 | 33.87 | 2100 | 0.0239 |
| TAQ001 | 1157 | Europe | Italy | 3097.9 | 11.77 | 42.25 | 2100 | 0.0238 |
| TAQ017 | 1157 | Europe | Italy | 3097.9 | 11.77 | 42.25 | 2100 | 0.0246 |
| I21312 | 770 | Europe | United Kingdom | 4439.0 | -1.84 | 51.28 | 2096 | 0.0208 |
| TAQ015 | 1157 | Europe | Italy | 3097.9 | 11.77 | 42.25 | 2096 | 0.0174 |
| I21314 | 770 | Europe | United Kingdom | 4439.0 | -1.84 | 51.28 | 2095 | 0.0242 |
| JAG001 | 285 | Asia | Mongolia | 6845.7 | 108.53 | 47.81 | 2082 | 0.0232 |
| I16601 | 702 | Europe | United Kingdom | 4423.8 | -1.77 | 51.17 | 2082 | 0.0227 |
| I14327 | 659 | Europe | United Kingdom | 4552.7 | -0.29 | 54.13 | 2081 | 0.0210 |
| I21311 | 769 | Europe | United Kingdom | 4423.8 | -2.14 | 51.22 | 2078 | 0.0197 |

| | | | | | | | | |
|---|---|---|---|---|---|---|---|---|
| I22060 | 773 | Europe | United Kingdom | 4552.7 | -0.28 | 54.09 | 2074 | 0.0191 |
| VOL001 | 1255 | Europe | Italy | 3218.0 | 10.85 | 43.42 | 2072 | 0.0163 |
| I16499 | 696 | Europe | United Kingdom | 4846.1 | -2.72 | 56.06 | 2069 | 0.0274 |
| I27379 | 821 | Europe | United Kingdom | 4352.6 | -0.69 | 50.81 | 2068 | 0.0180 |
| DA57_noUDG.SG | 100 | Asia | Kyrgyzstan | 4317.1 | 75.80 | 41.50 | 2065 | 0.0218 |
| I5503 | 894 | Europe | United Kingdom | 4586.2 | -0.67 | 53.92 | 2065 | 0.0272 |
| DA47_noUDG.SG | 99 | Asia | Kyrgyzstan | 4229.4 | 75.33 | 41.43 | 2063 | 0.0275 |
| DA58_noUDG.SG | 100 | Asia | Kyrgyzstan | 4317.1 | 75.80 | 41.50 | 2060 | 0.0234 |
| I16495 | 696 | Europe | United Kingdom | 4846.1 | -2.72 | 56.06 | 2055 | 0.0201 |
| C3356 | 56 | Asia | China | 4908.0 | 83.26 | 43.43 | 2054 | 0.0184 |
| I5502 | 894 | Europe | United Kingdom | 4586.2 | -0.67 | 53.92 | 2054 | 0.0239 |
| R10339.SG | 1066 | Europe | Italy | 3097.9 | 11.76 | 42.25 | 2054 | 0.0211 |
| R10341.SG | 1066 | Europe | Italy | 3097.9 | 11.76 | 42.25 | 2054 | 0.0235 |
| IMA005 | 271 | Asia | Russia | 6661.1 | 106.41 | 50.50 | 2051 | 0.0273 |
| JAA001 | 285 | Asia | Mongolia | 6845.7 | 108.53 | 47.81 | 2051 | 0.0208 |
| SKT001 | 385 | Asia | Mongolia | 6305.4 | 101.42 | 49.16 | 2051 | 0.0235 |
| SKT004 | 385 | Asia | Mongolia | 6305.4 | 101.42 | 49.16 | 2051 | 0.0257 |
| SKT007 | 385 | Asia | Mongolia | 6305.4 | 101.42 | 49.16 | 2051 | 0.0195 |
| SKT008 | 385 | Asia | Mongolia | 6305.4 | 101.42 | 49.16 | 2051 | 0.0212 |
| SKT009 | 385 | Asia | Mongolia | 6305.4 | 101.42 | 49.16 | 2051 | 0.0182 |
| SKT012 | 385 | Asia | Mongolia | 6305.4 | 101.42 | 49.16 | 2051 | 0.0196 |
| I17017 | 604 | Europe | United Kingdom | 4509.5 | -2.89 | 51.31 | 2051 | 0.0179 |
| I11716 | 616 | Europe | Slovakia | 3185.9 | 17.11 | 48.14 | 2051 | 0.0266 |
| I12785 | 634 | Europe | United Kingdom | 4467.2 | -1.75 | 51.89 | 2051 | 0.0197 |
| I19870 | 643 | Europe | United Kingdom | 4257.1 | 1.34 | 51.36 | 2051 | 0.0249 |
| CHK005 | 64 | Asia | Kyrgyzstan | 4490.6 | 78.38 | 42.49 | 2050 | 0.0192 |
| I16450 | 694 | Europe | United Kingdom | 4617.3 | -5.20 | 50.20 | 2050 | 0.0190 |
| I16457 | 694 | Europe | United Kingdom | 4617.3 | -5.20 | 50.20 | 2050 | 0.0224 |
| I14351 | 660 | Europe | United Kingdom | 4586.2 | -1.38 | 53.91 | 2048 | 0.0181 |
| I14352 | 660 | Europe | United Kingdom | 4586.2 | -1.38 | 53.91 | 2048 | 0.0186 |
| DA45_noUDG.SG | 98 | Asia | Mongolia | 6738.8 | 105.18 | 42.53 | 2046 | 0.0219 |

| | | | | | | | | |
|---|---|---|---|---|---|---|---|---|
| I11713 | 617 | Europe | Slovakia | 3185.9 | 17.10 | 48.14 | 2046 | 0.0253 |
| I14106 | 601 | Europe | United Kingdom | 4586.2 | -0.77 | 53.92 | 2045 | 0.0191 |
| R10342.SG | 1066 | Europe | Italy | 3097.9 | 11.76 | 42.25 | 2044 | 0.0258 |
| DA56_noUDG.SG | 84 | Asia | Kyrgyzstan | 3667.1 | 68.25 | 43.07 | 2043 | 0.0162 |
| SFI-20_noUDG.SG | 436 | Both | Lebanon | 776.6 | 35.51 | 33.90 | 2043 | 0.0124 |
| I19869 | 643 | Europe | United Kingdom | 4257.1 | 1.34 | 51.36 | 2043 | 0.0220 |
| I6549 | 259 | Asia | Pakistan | 3962.5 | 72.37 | 34.76 | 2041 | 0.0164 |
| C3315 | 54 | Asia | China | 4827.9 | 82.51 | 43.80 | 2038 | 0.0227 |
| I3567 | 846 | Europe | United Kingdom | 5084.9 | -5.82 | 57.43 | 2037 | 0.0263 |
| I12292 | 165 | Asia | Tajikistan | 3594.5 | 68.17 | 37.42 | 2036 | 0.0195 |
| C1664 | 47 | Asia | China | 4741.8 | 81.52 | 43.08 | 2034 | 0.0254 |
| R11546.SG | 353 | Asia | Armenia | 1927.6 | 43.85 | 40.69 | 2034 | 0.0249 |
| TU905_SX18 | 1167 | Europe | Switzerland | 3514.3 | 9.29 | 47.45 | 2032 | 0.0226 |
| I17750 | 717 | Europe | Netherlands | 4114.8 | 4.64 | 52.47 | 2031 | 0.0147 |
| I3566 | 846 | Europe | United Kingdom | 5084.9 | -5.82 | 57.43 | 2031 | 0.0190 |
| C3361 | 57 | Asia | China | 4908.0 | 83.28 | 43.54 | 2028 | 0.0242 |
| R10337.SG | 1066 | Europe | Italy | 3097.9 | 11.76 | 42.25 | 2028 | 0.0206 |
| C3316 | 54 | Asia | China | 4827.9 | 82.51 | 43.80 | 2027 | 0.0203 |
| C3325 | 54 | Asia | China | 4827.9 | 82.51 | 43.80 | 2027 | 0.0163 |
| SFI-15_noUDG.SG | 436 | Both | Lebanon | 776.6 | 35.51 | 33.90 | 2027 | 0.0149 |
| C3616 | 58 | Asia | China | 5398.2 | 89.26 | 43.76 | 2025 | 0.0237 |
| I20438 | 212 | Asia | Armenia | 1990.5 | 45.13 | 40.29 | 2023 | 0.0138 |
| I20437 | 212 | Asia | Armenia | 1990.5 | 45.13 | 40.29 | 2022 | 0.0183 |
| I11145 | 603 | Europe | United Kingdom | 4495.0 | -2.76 | 50.88 | 2022 | 0.0208 |
| I11712 | 617 | Europe | Slovakia | 3185.9 | 17.10 | 48.14 | 2011 | 0.0203 |
| I14097 | 656 | Europe | United Kingdom | 4668.7 | -1.61 | 54.40 | 2002 | 0.0133 |
| I2799 | 831 | Europe | United Kingdom | 5189.1 | -3.26 | 58.98 | 2002 | 0.0218 |
| BAM001 | 16 | Asia | Mongolia | 6382.5 | 102.83 | 48.60 | 2000 | 0.0295 |
| BRV001 | 36 | Asia | Kazakhstan | 4383.9 | 70.18 | 53.05 | 2000 | 0.0206 |
| BTO001 | 40 | Asia | Mongolia | 6449.9 | 103.37 | 49.48 | 2000 | 0.0234 |
| EME002 | 112 | Asia | Mongolia | 6390.8 | 101.99 | 47.78 | 2000 | 0.0209 |

| | | | | | | | | |
|---|---|---|---|---|---|---|---|---|
| HUD001 | 124 | Asia | Mongolia | 6315.0 | 101.71 | 47.83 | 2000 | 0.0238 |
| I6551 | 259 | Asia | Pakistan | 3962.5 | 72.37 | 34.76 | 2000 | 0.0222 |
| IMA001 | 271 | Asia | Russia | 6661.1 | 106.41 | 50.50 | 2000 | 0.0248 |
| IMA002 | 271 | Asia | Russia | 6661.1 | 106.41 | 50.50 | 2000 | 0.0221 |
| IMA007 | 271 | Asia | Russia | 6661.1 | 106.41 | 50.50 | 2000 | 0.0238 |
| NAI001 | 330 | Asia | Mongolia | 6308.0 | 101.86 | 48.59 | 2000 | 0.0219 |
| NAI002 | 330 | Asia | Mongolia | 6308.0 | 101.86 | 48.59 | 2000 | 0.0227 |
| SOL001 | 388 | Asia | Mongolia | 6315.0 | 101.77 | 47.74 | 2000 | 0.0222 |
| TAK001 | 394 | Asia | Mongolia | 5631.2 | 92.10 | 47.40 | 2000 | 0.0172 |
| TEV002 | 398 | Asia | Mongolia | 6438.7 | 102.06 | 44.65 | 2000 | 0.0257 |
| TEV003 | 398 | Asia | Mongolia | 6438.7 | 102.06 | 44.65 | 2000 | 0.0254 |
| I0156.SG | 572 | Europe | United Kingdom | 4343.0 | 0.18 | 52.08 | 1981 | 0.0227 |
| DEL001 | 41 | Asia | Mongolia | 7120.2 | 111.77 | 46.28 | 1978 | 0.0225 |
| I0160.SG | 572 | Europe | United Kingdom | 4343.0 | 0.18 | 52.08 | 1973 | 0.0279 |
| I22064 | 659 | Europe | United Kingdom | 4552.7 | -0.29 | 54.13 | 1971 | 0.0215 |
| I22057 | 659 | Europe | United Kingdom | 4552.7 | -0.29 | 54.13 | 1970 | 0.0238 |
| R2201.SG | 1094 | Europe | Slovakia | 3227.8 | 18.57 | 49.21 | 1970 | 0.0302 |
| I11710 | 616 | Europe | Slovakia | 3185.9 | 17.11 | 48.14 | 1967 | 0.0202 |
| CHN010 | 65 | Asia | Mongolia | 5649.6 | 92.05 | 50.00 | 1961 | 0.0183 |
| R2200.SG | 1094 | Europe | Slovakia | 3227.8 | 18.57 | 49.21 | 1960 | 0.0252 |
| TAQ002 | 1157 | Europe | Italy | 3097.9 | 11.77 | 42.25 | 1960 | 0.0163 |
| YUR001 | 306 | Asia | Mongolia | 6669.5 | 106.67 | 49.74 | 1958 | 0.0238 |
| I1636 | 201 | Asia | Armenia | 1845.9 | 45.08 | 39.55 | 1956 | 0.0205 |
| ULN004 | 41 | Asia | Mongolia | 7120.2 | 111.77 | 46.28 | 1952 | 0.0233 |
| I2497_noUDG | 216 | Asia | Vietnam | 7593.1 | 105.80 | 19.80 | 1950 | 0.0139 |
| I8218 | 265 | Asia | Pakistan | 3960.1 | 72.32 | 34.80 | 1949 | 0.0149 |
| IMA006 | 271 | Asia | Russia | 6661.1 | 106.41 | 50.50 | 1949 | 0.0240 |
| DA30_noUDG.SG | 89 | Asia | Kazakhstan | 4077.7 | 65.67 | 51.70 | 1948 | 0.0338 |
| R11541.SG | 353 | Asia | Armenia | 1927.6 | 43.85 | 40.69 | 1947 | 0.0201 |
| I10866 | 599 | Europe | Spain | 3994.5 | 3.11 | 42.13 | 1947 | 0.0217 |
| SFI-24_noUDG.SG | 436 | Both | Lebanon | 776.6 | 35.51 | 33.90 | 1946 | 0.0182 |

| | | | | | | | | |
|---|---|---|---|---|---|---|---|---|
| I27384 | 824 | Europe | United Kingdom | 4846.1 | -3.53 | 55.99 | 1945 | 0.0242 |
| I7714 | 258 | Asia | Pakistan | 3962.5 | 72.36 | 34.76 | 1944 | 0.0194 |
| DA39_noUDG.SG | 97 | Asia | Mongolia | 6315.0 | 101.35 | 48.02 | 1942 | 0.0306 |
| DUU001 | 108 | Asia | Mongolia | 6931.2 | 109.81 | 47.40 | 1941 | 0.0159 |
| I22062 | 774 | Europe | United Kingdom | 4552.7 | -0.31 | 54.06 | 1930 | 0.0226 |
| I6550 | 259 | Asia | Pakistan | 3962.5 | 72.37 | 34.76 | 1926 | 0.0208 |
| R2202.SG | 1095 | Europe | Slovakia | 3227.8 | 18.21 | 49.07 | 1925 | 0.0147 |
| I18599 | 735 | Europe | United Kingdom | 4257.1 | 0.74 | 51.32 | 1924 | 0.0170 |
| I21302 | 623 | Europe | United Kingdom | 4423.8 | -2.32 | 51.01 | 1922 | 0.0201 |
| BUR004 | 42 | Asia | Mongolia | 6390.8 | 102.36 | 47.80 | 1919 | 0.0210 |
| I6228 | 174 | Asia | Mongolia | 5649.6 | 92.05 | 49.96 | 1919 | 0.0193 |
| I16504 | 697 | Europe | United Kingdom | 4814.3 | -2.48 | 55.99 | 1916 | 0.0188 |
| I27385 | 824 | Europe | United Kingdom | 4846.1 | -3.53 | 55.99 | 1916 | 0.0185 |
| I3568 | 846 | Europe | United Kingdom | 5084.9 | -5.82 | 57.43 | 1912 | 0.0273 |
| BRE005 | 31 | Asia | Kazakhstan | 5197.6 | 85.79 | 49.33 | 1905 | 0.0196 |
| R11651.SG | 353 | Asia | Armenia | 1927.6 | 43.85 | 40.69 | 1900 | 0.0223 |
| R11653.SG | 353 | Asia | Armenia | 1927.6 | 43.85 | 40.69 | 1900 | 0.0136 |
| R11655.SG | 353 | Asia | Armenia | 1927.6 | 43.85 | 40.69 | 1900 | 0.0223 |
| R11678.SG | 353 | Asia | Armenia | 1927.6 | 43.85 | 40.69 | 1900 | 0.0201 |
| R11713.SG | 353 | Asia | Armenia | 1927.6 | 43.85 | 40.69 | 1900 | 0.0181 |
| R11714.SG | 353 | Asia | Armenia | 1927.6 | 43.85 | 40.69 | 1900 | 0.0262 |
| I14845 | 674 | Europe | Turkey | 1828.6 | 28.69 | 40.18 | 1900 | 0.0172 |
| SAN001 | 374 | Asia | Mongolia | 7149.0 | 112.97 | 47.92 | 1890 | 0.0231 |
| BRE009 | 31 | Asia | Kazakhstan | 5197.6 | 85.79 | 49.33 | 1886 | 0.0221 |
| BRE012 | 31 | Asia | Kazakhstan | 5197.6 | 85.79 | 49.33 | 1886 | 0.0177 |
| UGU005 | 404 | Asia | Mongolia | 6600.9 | 105.38 | 49.25 | 1885 | 0.0223 |
| ATS001 | 14 | Asia | Mongolia | 6606.0 | 105.90 | 48.87 | 1878 | 0.0269 |
| KHO006 | 22 | Asia | Mongolia | 5630.5 | 91.84 | 47.06 | 1875 | 0.0223 |
| R10625.SG | 1072 | Europe | Poland | 3672.4 | 19.57 | 54.12 | 1875 | 0.0187 |
| R10626.SG | 1072 | Europe | Poland | 3672.4 | 19.57 | 54.12 | 1875 | 0.0207 |
| R10631.SG | 1072 | Europe | Poland | 3672.4 | 19.57 | 54.12 | 1875 | 0.0187 |

| | | | | | | | | |
|---|---|---|---|---|---|---|---|---|
| R10633.SG | 1072 | Europe | Poland | 3672.4 | 19.57 | 54.12 | 1875 | 0.0288 |
| R10634.SG | 1072 | Europe | Poland | 3672.4 | 19.57 | 54.12 | 1875 | 0.0285 |
| R10636.SG | 1072 | Europe | Poland | 3672.4 | 19.57 | 54.12 | 1875 | 0.0193 |
| I1639 | 201 | Asia | Armenia | 1845.9 | 45.08 | 39.55 | 1872 | 0.0287 |
| R10657.SG | 1073 | Europe | Austria | 3232.5 | 16.32 | 48.31 | 1863 | 0.0291 |
| R10659.SG | 1073 | Europe | Austria | 3232.5 | 16.32 | 48.31 | 1863 | 0.0209 |
| R6769.SG | 1116 | Europe | Serbia | 2458.4 | 21.90 | 43.33 | 1863 | 0.0258 |
| DCZ-M22IV_noUDG | 106 | Asia | China | 6677.6 | 101.96 | 36.53 | 1850 | 0.0263 |
| I10252 | 150 | Asia | Armenia | 1927.6 | 43.92 | 40.83 | 1850 | 0.0180 |
| I14550 | 664 | Europe | United Kingdom | 4296.4 | -0.27 | 50.84 | 1850 | 0.0230 |
| R111.SG | 1077 | Europe | Italy | 3097.9 | 12.49 | 41.92 | 1850 | 0.0139 |
| R113.SG | 1077 | Europe | Italy | 3097.9 | 12.49 | 41.92 | 1850 | 0.0234 |
| R114.SG | 1077 | Europe | Italy | 3097.9 | 12.49 | 41.92 | 1850 | 0.0202 |
| R115.SG | 1077 | Europe | Italy | 3097.9 | 12.49 | 41.92 | 1850 | 0.0219 |
| R116.SG | 1077 | Europe | Italy | 3097.9 | 12.49 | 41.92 | 1850 | 0.0186 |
| R131.SG | 1077 | Europe | Italy | 3097.9 | 12.49 | 41.92 | 1850 | 0.0293 |
| R75.SG | 1077 | Europe | Italy | 3097.9 | 12.49 | 41.92 | 1850 | 0.0209 |
| R76.SG | 1077 | Europe | Italy | 3097.9 | 12.49 | 41.92 | 1850 | 0.0218 |
| R78.SG | 1077 | Europe | Italy | 3097.9 | 12.49 | 41.92 | 1850 | 0.0158 |
| R436.SG | 1110 | Europe | Italy | 3002.1 | 12.78 | 41.80 | 1850 | 0.0295 |
| VK532_noUDG.SG | 1241 | Europe | Denmark | 4084.8 | 12.25 | 55.66 | 1850 | 0.0235 |
| R80.SG | 1077 | Europe | Italy | 3097.9 | 12.49 | 41.92 | 1840 | 0.0250 |
| R9674.SG | 1109 | Europe | Serbia | 2638.7 | 21.17 | 44.72 | 1836 | 0.0260 |
| I12927 | 636 | Europe | United Kingdom | 4467.2 | -2.08 | 51.95 | 1825 | 0.0224 |
| I12931 | 636 | Europe | United Kingdom | 4467.2 | -2.08 | 51.95 | 1825 | 0.0251 |
| I12932 | 636 | Europe | United Kingdom | 4467.2 | -2.08 | 51.95 | 1825 | 0.0203 |
| R1556.SG | 1087 | Europe | Italy | 3176.2 | 12.63 | 43.72 | 1825 | 0.0227 |
| R1557.SG | 1087 | Europe | Italy | 3176.2 | 12.63 | 43.72 | 1825 | 0.0161 |
| R1547.SG | 1086 | Europe | Italy | 3097.9 | 12.59 | 42.07 | 1814 | 0.0189 |
| R1548.SG | 1086 | Europe | Italy | 3097.9 | 12.59 | 42.07 | 1814 | 0.0213 |
| R1549.SG | 1086 | Europe | Italy | 3097.9 | 12.59 | 42.07 | 1814 | 0.0213 |

| | | | | | | | | |
|---|---|---|---|---|---|---|---|---|
| R1550.SG | 1086 | Europe | Italy | 3097.9 | 12.59 | 42.07 | 1814 | 0.0201 |
| R835.SG | 1117 | Europe | Italy | 3082.9 | 13.65 | 43.31 | 1814 | 0.0208 |
| R836.SG | 1117 | Europe | Italy | 3082.9 | 13.65 | 43.31 | 1814 | 0.0217 |
| C2032 | 51 | Asia | China | 5164.6 | 86.26 | 41.34 | 1811 | 0.0185 |
| SFI-33_noUDG.SG | 436 | Both | Lebanon | 776.6 | 35.51 | 33.90 | 1804 | 0.0211 |
| DA70_noUDG.SG | 96 | Asia | Kyrgyzstan | 4317.1 | 75.79 | 41.50 | 1802 | 0.0203 |
| BUR002 | 42 | Asia | Mongolia | 6390.8 | 102.36 | 47.80 | 1800 | 0.0253 |
| BUR003 | 42 | Asia | Mongolia | 6390.8 | 102.36 | 47.80 | 1800 | 0.0166 |
| TUH001 | 42 | Asia | Mongolia | 6390.8 | 102.36 | 47.80 | 1800 | 0.0199 |
| TUH002 | 42 | Asia | Mongolia | 6390.8 | 102.36 | 47.80 | 1800 | 0.0209 |
| TUK002 | 42 | Asia | Mongolia | 6390.8 | 102.36 | 47.80 | 1800 | 0.0260 |
| TUK003 | 42 | Asia | Mongolia | 6390.8 | 102.36 | 47.80 | 1800 | 0.0195 |
| I8475 | 599 | Europe | Spain | 3994.5 | 3.11 | 42.13 | 1800 | 0.0226 |
| R2204.SG | 1096 | Europe | Slovakia | 3185.9 | 16.98 | 48.32 | 1800 | 0.0257 |
| R3744.SG | 1107 | Europe | Croatia | 2905.3 | 15.28 | 44.10 | 1800 | 0.0178 |
| R9673.SG | 1109 | Europe | Serbia | 2638.7 | 21.17 | 44.72 | 1800 | 0.0233 |
| R1555.SG | 1087 | Europe | Italy | 3176.2 | 12.63 | 43.72 | 1798 | 0.0177 |
| R3743.SG | 1107 | Europe | Croatia | 2905.3 | 15.28 | 44.10 | 1798 | 0.0263 |
| R1551.SG | 1086 | Europe | Italy | 3097.9 | 12.59 | 42.07 | 1797 | 0.0184 |
| R6750.SG | 1109 | Europe | Serbia | 2638.7 | 21.17 | 44.72 | 1795 | 0.0208 |
| C3622 | 59 | Asia | China | 4674.3 | 80.19 | 37.07 | 1789 | 0.0191 |
| R10618.SG | 1072 | Europe | Poland | 3672.4 | 19.57 | 54.12 | 1789 | 0.0277 |
| R6759.SG | 1109 | Europe | Serbia | 2638.7 | 21.17 | 44.72 | 1789 | 0.0260 |
| C391_noUDG | 60 | Asia | China | 4678.7 | 79.94 | 36.97 | 1780 | 0.0194 |
| R10667.SG | 1074 | Europe | Austria | 3327.4 | 14.04 | 48.17 | 1780 | 0.0184 |
| R123.SG | 1082 | Europe | Italy | 3002.1 | 13.15 | 41.68 | 1780 | 0.0227 |
| R1554.SG | 1087 | Europe | Italy | 3176.2 | 12.63 | 43.72 | 1777 | 0.0246 |
| R3742.SG | 1107 | Europe | Croatia | 2905.3 | 15.28 | 44.10 | 1773 | 0.0230 |
| R3931.SG | 1109 | Europe | Serbia | 2638.7 | 21.17 | 44.72 | 1769 | 0.0207 |
| R6756.SG | 1109 | Europe | Serbia | 2638.7 | 21.17 | 44.72 | 1769 | 0.0173 |
| DCZ-M21II_noUDG | 106 | Asia | China | 6677.6 | 101.96 | 36.53 | 1763 | 0.0196 |

| | | | | | | | | |
|---|---|---|---|---|---|---|---|---|
| 3DT26_noUDG.SG | 439 | Europe | United Kingdom | 4586.2 | -1.08 | 53.96 | 1750 | 0.0187 |
| 6DT18_noUDG.SG | 439 | Europe | United Kingdom | 4586.2 | -1.08 | 53.96 | 1750 | 0.0197 |
| 6DT21_noUDG.SG | 439 | Europe | United Kingdom | 4586.2 | -1.08 | 53.96 | 1750 | 0.0192 |
| 6DT22_noUDG.SG | 439 | Europe | United Kingdom | 4586.2 | -1.08 | 53.96 | 1750 | 0.0199 |
| 6DT3_noUDG.SG | 439 | Europe | United Kingdom | 4586.2 | -1.08 | 53.96 | 1750 | 0.0261 |
| R125.SG | 1082 | Europe | Italy | 3002.1 | 13.15 | 41.68 | 1750 | 0.0195 |
| R128.SG | 1082 | Europe | Italy | 3002.1 | 13.15 | 41.68 | 1750 | 0.0272 |
| R1543.SG | 1085 | Europe | Italy | 3097.9 | 12.33 | 42.20 | 1750 | 0.0252 |
| R1544.SG | 1085 | Europe | Italy | 3097.9 | 12.33 | 42.20 | 1750 | 0.0203 |
| R1545.SG | 1085 | Europe | Italy | 3097.9 | 12.33 | 42.20 | 1750 | 0.0174 |
| R49.SG | 1111 | Europe | Italy | 3086.5 | 12.55 | 41.84 | 1750 | 0.0169 |
| R51.SG | 1111 | Europe | Italy | 3086.5 | 12.55 | 41.84 | 1750 | 0.0214 |
| R66.SG | 1113 | Europe | Italy | 3086.5 | 12.47 | 41.83 | 1750 | 0.0176 |
| R67.SG | 1113 | Europe | Italy | 3086.5 | 12.47 | 41.83 | 1750 | 0.0175 |
| R68.SG | 1113 | Europe | Italy | 3086.5 | 12.47 | 41.83 | 1750 | 0.0258 |
| R69.SG | 1113 | Europe | Italy | 3086.5 | 12.47 | 41.83 | 1750 | 0.0221 |
| R70.SG | 1113 | Europe | Italy | 3086.5 | 12.47 | 41.83 | 1750 | 0.0210 |
| R71.SG | 1113 | Europe | Italy | 3086.5 | 12.47 | 41.83 | 1750 | 0.0227 |
| R72.SG | 1113 | Europe | Italy | 3086.5 | 12.47 | 41.83 | 1750 | 0.0183 |
| R73.SG | 1113 | Europe | Italy | 3086.5 | 12.47 | 41.83 | 1750 | 0.0164 |
| R11563.SG | 1079 | Europe | France | 3730.5 | 7.05 | 48.73 | 1742 | 0.0197 |
| R10668.SG | 1074 | Europe | Austria | 3327.4 | 14.04 | 48.17 | 1735 | 0.0183 |
| R9669.SG | 1109 | Europe | Serbia | 2638.7 | 21.17 | 44.72 | 1732 | 0.0261 |
| I20000 | 754 | Europe | Turkey | 1573.9 | 28.03 | 37.30 | 1726 | 0.0162 |
| I20001 | 754 | Europe | Turkey | 1573.9 | 28.03 | 37.30 | 1726 | 0.0154 |
| I20139 | 754 | Europe | Turkey | 1573.9 | 28.03 | 37.30 | 1726 | 0.0161 |
| R10656.SG | 1073 | Europe | Austria | 3232.5 | 16.32 | 48.31 | 1725 | 0.0228 |
| R10658.SG | 1073 | Europe | Austria | 3232.5 | 16.32 | 48.31 | 1725 | 0.0262 |
| R10660.SG | 1073 | Europe | Austria | 3232.5 | 16.32 | 48.31 | 1725 | 0.0150 |
| I11540 | 159 | Asia | Kazakhstan | 3531.0 | 57.04 | 50.45 | 1723 | 0.0272 |
| BRE013 | 31 | Asia | Kazakhstan | 5197.6 | 85.79 | 49.33 | 1716 | 0.0181 |

| | | | | | | | | |
|---|---|---|---|---|---|---|---|---|
| ALN003 | 7 | Asia | Kyrgyzstan | 3974.4 | 71.50 | 41.41 | 1700 | 0.0219 |
| ALN008 | 7 | Asia | Kyrgyzstan | 3974.4 | 71.50 | 41.41 | 1700 | 0.0176 |
| ALN009 | 7 | Asia | Kyrgyzstan | 3974.4 | 71.50 | 41.41 | 1700 | 0.0195 |
| DA123_noUDG.SG | 72 | Asia | Kazakhstan | 3658.0 | 68.32 | 42.73 | 1700 | 0.0263 |
| I20802 | 766 | Europe | Hungary | 2913.1 | 21.53 | 47.37 | 1700 | 0.0209 |
| R3481.SG | 1099 | Europe | Montenegro | 2535.7 | 19.27 | 42.47 | 1699 | 0.0262 |
| R50.SG | 1111 | Europe | Italy | 3086.5 | 12.55 | 41.84 | 1699 | 0.0190 |
| tem003_noUDG.SG | 397 | Asia | Russia | 3780.8 | 58.12 | 52.99 | 1698 | 0.0318 |
| DA206_noUDG.SG | 82 | Asia | Kazakhstan | 3667.1 | 68.25 | 42.90 | 1697 | 0.0165 |
| R10670.SG | 1074 | Europe | Austria | 3327.4 | 14.04 | 48.17 | 1681 | 0.0174 |
| R11552.SG | 1079 | Europe | France | 3730.5 | 7.05 | 48.73 | 1675 | 0.0268 |
| R11553.SG | 1079 | Europe | France | 3730.5 | 7.05 | 48.73 | 1675 | 0.0297 |
| R11554.SG | 1079 | Europe | France | 3730.5 | 7.05 | 48.73 | 1675 | 0.0219 |
| R11555.SG | 1079 | Europe | France | 3730.5 | 7.05 | 48.73 | 1675 | 0.0239 |
| R11559.SG | 1079 | Europe | France | 3730.5 | 7.05 | 48.73 | 1675 | 0.0231 |
| R11561.SG | 1079 | Europe | France | 3730.5 | 7.05 | 48.73 | 1675 | 0.0204 |
| R132.SG | 1084 | Europe | Italy | 3097.9 | 12.50 | 41.89 | 1670 | 0.0152 |
| R3656.SG | 1104 | Europe | Croatia | 2878.2 | 18.68 | 45.55 | 1668 | 0.0240 |
| BRE007 | 31 | Asia | Kazakhstan | 5197.6 | 85.79 | 49.33 | 1667 | 0.0225 |
| DA85_noUDG.SG | 101 | Asia | Kyrgyzstan | 4232.7 | 74.99 | 41.62 | 1666 | 0.0260 |
| DA101_noUDG.SG | 69 | Asia | Kyrgyzstan | 4404.2 | 77.41 | 42.16 | 1664 | 0.0189 |
| R6701.SG | 1115 | Europe | Serbia | 2870.8 | 19.17 | 45.91 | 1661 | 0.0217 |
| R10477.SG | 1068 | Europe | Slovenia | 3130.3 | 14.51 | 46.06 | 1659 | 0.0231 |
| DA81_noUDG.SG | 101 | Asia | Kyrgyzstan | 4232.7 | 74.99 | 41.62 | 1656 | 0.0211 |
| R47.SG | 1111 | Europe | Italy | 3086.5 | 12.55 | 41.84 | 1655 | 0.0213 |
| KNT004 | 301 | Asia | Kazakhstan | 3719.4 | 68.80 | 41.41 | 1653 | 0.0212 |
| C387_noUDG | 60 | Asia | China | 4678.7 | 79.94 | 36.97 | 1651 | 0.0204 |
| C388_noUDG | 60 | Asia | China | 4678.7 | 79.94 | 36.97 | 1651 | 0.0120 |
| C390_noUDG | 60 | Asia | China | 4678.7 | 79.94 | 36.97 | 1651 | 0.0211 |
| C392_noUDG | 60 | Asia | China | 4678.7 | 79.94 | 36.97 | 1651 | 0.0171 |
| KNT002 | 301 | Asia | Kazakhstan | 3719.4 | 68.80 | 41.41 | 1650 | 0.0257 |

| | | | | | | | | |
|---|---|---|---|---|---|---|---|---|
| KNT003 | 301 | Asia | Kazakhstan | 3719.4 | 68.80 | 41.41 | 1650 | 0.0200 |
| FN2_noUDG.SG | 538 | Europe | Germany | 3480.3 | 11.41 | 48.14 | 1650 | 0.0257 |
| I3982 | 853 | Europe | Spain | 4878.1 | -3.61 | 37.18 | 1650 | 0.0250 |
| R126.SG | 1082 | Europe | Italy | 3002.1 | 13.15 | 41.68 | 1650 | 0.0231 |
| R2208.SG | 1097 | Europe | Slovakia | 3139.7 | 18.36 | 48.34 | 1650 | 0.0223 |
| VK521_noUDG.SG | 1238 | Europe | Denmark | 4084.8 | 12.24 | 55.64 | 1650 | 0.0202 |
| R2041.SG | 1090 | Europe | Croatia | 2983.9 | 16.38 | 45.46 | 1647 | 0.0260 |
| KNT001 | 301 | Asia | Kazakhstan | 3719.4 | 68.80 | 41.41 | 1641 | 0.0194 |
| ALN001 | 7 | Asia | Kyrgyzstan | 3974.4 | 71.50 | 41.41 | 1632 | 0.0162 |
| BRE014 | 31 | Asia | Kazakhstan | 5197.6 | 85.79 | 49.33 | 1630 | 0.0241 |
| BRE008 | 31 | Asia | Kazakhstan | 5197.6 | 85.79 | 49.33 | 1629 | 0.0223 |
| R10467.SG | 1068 | Europe | Slovenia | 3130.3 | 14.51 | 46.06 | 1628 | 0.0192 |
| BRE004 | 31 | Asia | Kazakhstan | 5197.6 | 85.79 | 49.33 | 1627 | 0.0217 |
| DA224_noUDG.SG | 84 | Asia | Kazakhstan | 3667.1 | 68.25 | 43.07 | 1627 | 0.0247 |
| DA71_noUDG.SG | 84 | Asia | Kyrgyzstan | 3667.1 | 68.25 | 43.07 | 1627 | 0.0135 |
| QED-7_noUDG.SG | 432 | Both | Lebanon | 776.6 | 35.78 | 34.05 | 1627 | 0.0234 |
| R2040.SG | 1090 | Europe | Croatia | 2983.9 | 16.38 | 45.46 | 1625 | 0.0110 |
| R6688.SG | 1114 | Europe | Serbia | 2732.7 | 20.07 | 45.13 | 1625 | 0.0227 |
| TUK001 | 42 | Asia | Mongolia | 6390.8 | 102.36 | 47.80 | 1623 | 0.0186 |
| KNT005 | 301 | Asia | Kazakhstan | 3719.4 | 68.80 | 41.41 | 1622 | 0.0203 |
| R10478.SG | 1068 | Europe | Slovenia | 3130.3 | 14.51 | 46.06 | 1622 | 0.0332 |
| R1874.SG | 1089 | Europe | France | 4090.8 | 2.30 | 49.61 | 1622 | 0.0213 |
| C795 | 62 | Asia | China | 5161.5 | 86.15 | 41.76 | 1620 | 0.0262 |
| R3542.SG | 1101 | Europe | Croatia | 2878.2 | 18.61 | 45.77 | 1616 | 0.0178 |
| R6764.SG | 1116 | Europe | Serbia | 2458.4 | 21.90 | 43.33 | 1616 | 0.0258 |
| BRE011 | 31 | Asia | Kazakhstan | 5197.6 | 85.79 | 49.33 | 1615 | 0.0219 |
| DA100_noUDG.SG | 69 | Asia | Kyrgyzstan | 4404.2 | 77.41 | 42.16 | 1615 | 0.0251 |
| QED-2_noUDG.SG | 432 | Both | Lebanon | 776.6 | 35.78 | 34.05 | 1610 | 0.0147 |
| I7833 | 945 | Europe | Greece | 1953.9 | 23.95 | 38.12 | 1610 | 0.0194 |
| R10474.SG | 1068 | Europe | Slovenia | 3130.3 | 14.51 | 46.06 | 1609 | 0.0292 |
| R10499.SG | 1071 | Europe | Portugal | 5047.7 | -9.37 | 38.73 | 1609 | 0.0241 |

| | | | | | | | | |
|---|---|---|---|---|---|---|---|---|
| R11550.SG | 1079 | Europe | France | 3730.5 | 7.05 | 48.73 | 1609 | 0.0145 |
| R11558.SG | 1079 | Europe | France | 3730.5 | 7.05 | 48.73 | 1609 | 0.0177 |
| R6681.SG | 1114 | Europe | Serbia | 2732.7 | 20.07 | 45.13 | 1609 | 0.0242 |
| DA96_noUDG.SG | 69 | Asia | Kyrgyzstan | 4404.2 | 77.41 | 42.16 | 1608 | 0.0266 |
| R10654.SG | 1073 | Europe | Austria | 3232.5 | 16.32 | 48.31 | 1603 | 0.0205 |
| DA104_noUDG.SG | 69 | Asia | Kyrgyzstan | 4404.2 | 77.41 | 42.16 | 1600 | 0.0204 |
| DA177_noUDG.SG | 78 | Asia | Kazakhstan | 4676.3 | 77.19 | 52.02 | 1600 | 0.0192 |
| DA80_noUDG.SG | 96 | Asia | Kyrgyzstan | 4317.1 | 75.79 | 41.50 | 1600 | 0.0266 |
| VK418_noUDG.SG | 1227 | Europe | Norway | 5144.6 | 14.98 | 67.94 | 1600 | 0.0256 |
| R11560.SG | 1079 | Europe | France | 3730.5 | 7.05 | 48.73 | 1598 | 0.0234 |
| SZLA-646.SG | 1154 | Europe | Hungary | 3049.0 | 20.45 | 48.23 | 1597 | 0.0211 |
| DA95_noUDG.SG | 102 | Asia | Kazakhstan | 4733.4 | 76.75 | 52.62 | 1596 | 0.0311 |
| ALN005 | 7 | Asia | Kyrgyzstan | 3974.4 | 71.50 | 41.41 | 1592 | 0.0245 |
| R10487.SG | 1069 | Europe | Portugal | 4910.4 | -8.49 | 40.10 | 1587 | 0.0221 |
| R10508.SG | 1071 | Europe | Portugal | 5047.7 | -9.37 | 38.73 | 1587 | 0.0188 |
| R11557.SG | 1079 | Europe | France | 3730.5 | 7.05 | 48.73 | 1587 | 0.0210 |
| POP23_noUDG.SG | 1051 | Europe | Croatia | 2878.2 | 18.57 | 45.75 | 1586 | 0.0144 |
| R10840.SG | 1076 | Europe | Lithuania | 3500.2 | 25.28 | 54.69 | 1581 | 0.0195 |
| DA72_noUDG.SG | 96 | Asia | Kyrgyzstan | 4317.1 | 75.79 | 41.50 | 1575 | 0.0245 |
| I14799 | 671 | Europe | Turkey | 1775.5 | 29.71 | 40.42 | 1575 | 0.0121 |
| I8372 | 671 | Europe | Turkey | 1775.5 | 29.71 | 40.42 | 1565 | 0.0213 |
| KRY001 | 307 | Asia | Kazakhstan | 3531.0 | 57.06 | 50.47 | 1552 | 0.0270 |
| C1652 | 46 | Asia | China | 4824.5 | 81.84 | 43.22 | 1551 | 0.0191 |
| AKG_10203.SG | 5 | Asia | South Korea | 9032.9 | 128.88 | 35.24 | 1550 | 0.0235 |
| AKG_10204.SG | 5 | Asia | South Korea | 9032.9 | 128.88 | 35.24 | 1550 | 0.0238 |
| AKG_10207.SG | 5 | Asia | South Korea | 9032.9 | 128.88 | 35.24 | 1550 | 0.0209 |
| AKG_10209.SG | 5 | Asia | South Korea | 9032.9 | 128.88 | 35.24 | 1550 | 0.0186 |
| AKG_10210.SG | 5 | Asia | South Korea | 9032.9 | 128.88 | 35.24 | 1550 | 0.0211 |
| AKG_10218.SG | 5 | Asia | South Korea | 9032.9 | 128.88 | 35.24 | 1550 | 0.0210 |
| AKG_3421.SG | 5 | Asia | South Korea | 9032.9 | 128.88 | 35.24 | 1550 | 0.0241 |
| AKG_3420.SG | 6 | Asia | South Korea | 9082.2 | 128.81 | 35.22 | 1550 | 0.0304 |

| | | | | | | | | |
|---|---|---|---|---|---|---|---|---|
| A181021 | 445 | Europe | Hungary | 2958.8 | 21.36 | 47.86 | 1550 | 0.0193 |
| A181022 | 445 | Europe | Hungary | 2958.8 | 21.36 | 47.86 | 1550 | 0.0210 |
| A181023 | 445 | Europe | Hungary | 2958.8 | 21.36 | 47.86 | 1550 | 0.0178 |
| A181024 | 445 | Europe | Hungary | 2958.8 | 21.36 | 47.86 | 1550 | 0.0223 |
| A181025 | 445 | Europe | Hungary | 2958.8 | 21.36 | 47.86 | 1550 | 0.0215 |
| A181026 | 445 | Europe | Hungary | 2958.8 | 21.36 | 47.86 | 1550 | 0.0231 |
| A181027 | 445 | Europe | Hungary | 2958.8 | 21.36 | 47.86 | 1550 | 0.0195 |
| A181028 | 445 | Europe | Hungary | 2958.8 | 21.36 | 47.86 | 1550 | 0.0225 |
| CSB-3.SG | 511 | Europe | Hungary | 2868.4 | 20.25 | 46.71 | 1550 | 0.0235 |
| HID001.B_noUDG | 556 | Europe | Germany | 3881.5 | 9.71 | 52.28 | 1550 | 0.0178 |
| HID002.A_noUDG | 556 | Europe | Germany | 3881.5 | 9.71 | 52.28 | 1550 | 0.0241 |
| HID004_noUDG | 556 | Europe | Germany | 3881.5 | 9.71 | 52.28 | 1550 | 0.0217 |
| I3983 | 853 | Europe | Spain | 4878.1 | -3.61 | 37.18 | 1550 | 0.0142 |
| R130.SG | 1084 | Europe | Italy | 3097.9 | 12.50 | 41.89 | 1550 | 0.0233 |
| R133.SG | 1084 | Europe | Italy | 3097.9 | 12.50 | 41.89 | 1550 | 0.0181 |
| R134.SG | 1084 | Europe | Italy | 3097.9 | 12.50 | 41.89 | 1550 | 0.0264 |
| R136.SG | 1084 | Europe | Italy | 3097.9 | 12.50 | 41.89 | 1550 | 0.0198 |
| R137.SG | 1084 | Europe | Italy | 3097.9 | 12.50 | 41.89 | 1550 | 0.0229 |
| R2209.SG | 1097 | Europe | Slovakia | 3139.7 | 18.36 | 48.34 | 1550 | 0.0246 |
| R2210.SG | 1097 | Europe | Slovakia | 3139.7 | 18.36 | 48.34 | 1550 | 0.0239 |
| R2211.SG | 1097 | Europe | Slovakia | 3139.7 | 18.36 | 48.34 | 1550 | 0.0244 |
| VK522_noUDG.SG | 1239 | Europe | Sweden | 4093.6 | 16.61 | 56.54 | 1550 | 0.0196 |
| R6730.SG | 1108 | Europe | Serbia | 2743.4 | 19.61 | 44.97 | 1547 | 0.0201 |
| R31.SG | 1098 | Europe | Italy | 3097.9 | 12.47 | 41.91 | 1538 | 0.0206 |
| ASZK-1.SG | 473 | Europe | Hungary | 3098.7 | 17.48 | 47.63 | 1525 | 0.0247 |
| R10488.SG | 1069 | Europe | Portugal | 4910.4 | -8.49 | 40.10 | 1525 | 0.0176 |
| R10500.SG | 1071 | Europe | Portugal | 5047.7 | -9.37 | 38.73 | 1525 | 0.0136 |
| R10501.SG | 1071 | Europe | Portugal | 5047.7 | -9.37 | 38.73 | 1525 | 0.0198 |
| R10502.SG | 1071 | Europe | Portugal | 5047.7 | -9.37 | 38.73 | 1525 | 0.0208 |
| R10506.SG | 1071 | Europe | Portugal | 5047.7 | -9.37 | 38.73 | 1525 | 0.0183 |
| R10507.SG | 1071 | Europe | Portugal | 5047.7 | -9.37 | 38.73 | 1525 | 0.0237 |

| | | | | | | | | |
|---|---|---|---|---|---|---|---|---|
| SEI-1.SG | 1136 | Europe | Hungary | 2868.4 | 20.13 | 46.49 | 1513 | 0.0212 |
| SEI-5.SG | 1136 | Europe | Hungary | 2868.4 | 20.13 | 46.49 | 1513 | 0.0229 |
| SEI-6.SG | 1136 | Europe | Hungary | 2868.4 | 20.13 | 46.49 | 1513 | 0.0183 |
| R10830.SG | 1075 | Europe | Lithuania | 3582.1 | 23.90 | 54.90 | 1512 | 0.0226 |
| R2053.SG | 1091 | Europe | Croatia | 3052.4 | 14.54 | 45.20 | 1512 | 0.0175 |
| DA69_noUDG.SG | 96 | Asia | Kyrgyzstan | 4317.1 | 75.79 | 41.50 | 1509 | 0.0240 |
| KMT-2785.SG | 970 | Europe | Hungary | 2913.5 | 19.84 | 47.03 | 1509 | 0.0190 |
| DA27_noUDG.SG | 88 | Asia | Kazakhstan | 4006.3 | 62.89 | 52.84 | 1508 | 0.0196 |
| A181014 | 444 | Europe | Hungary | 2960.2 | 19.72 | 46.86 | 1500 | 0.0232 |
| A181015 | 444 | Europe | Hungary | 2960.2 | 19.72 | 46.86 | 1500 | 0.0170 |
| A181018 | 444 | Europe | Hungary | 2960.2 | 19.72 | 46.86 | 1500 | 0.0200 |
| A181019 | 444 | Europe | Hungary | 2960.2 | 19.72 | 46.86 | 1500 | 0.0214 |
| A181020 | 444 | Europe | Hungary | 2960.2 | 19.72 | 46.86 | 1500 | 0.0255 |
| A181029 | 446 | Europe | Hungary | 3098.7 | 17.41 | 47.51 | 1500 | 0.0185 |
| Alh1_noUDG.SG | 460 | Europe | Germany | 3465.3 | 12.20 | 48.58 | 1500 | 0.0230 |
| R11868.SG | 1081 | Europe | Germany | 3715.8 | 11.00 | 51.11 | 1500 | 0.0221 |
| R11872.SG | 1081 | Europe | Germany | 3715.8 | 11.00 | 51.11 | 1500 | 0.0255 |
| R11873.SG | 1081 | Europe | Germany | 3715.8 | 11.00 | 51.11 | 1500 | 0.0248 |
| R11875.SG | 1081 | Europe | Germany | 3715.8 | 11.00 | 51.11 | 1500 | 0.0190 |
| R2206.SG | 1097 | Europe | Slovakia | 3139.7 | 18.36 | 48.34 | 1500 | 0.0244 |
| R2207.SG | 1097 | Europe | Slovakia | 3139.7 | 18.36 | 48.34 | 1500 | 0.0197 |
| VZ-12673.SG | 1259 | Europe | Hungary | 3004.9 | 19.15 | 47.64 | 1500 | 0.0182 |
| DA74_noUDG.SG | 96 | Asia | Kyrgyzstan | 4317.1 | 75.79 | 41.50 | 1492 | 0.0260 |
| I13175 | 181 | Asia | Mongolia | 5554.2 | 90.80 | 45.39 | 1483 | 0.0173 |
| I0773 | 582 | Europe | United Kingdom | 4389.8 | 0.07 | 52.26 | 1479 | 0.0219 |
| Alh10_noUDG.SG | 460 | Europe | Germany | 3465.3 | 12.20 | 48.58 | 1477 | 0.0236 |
| MSG-1.SG | 1004 | Europe | Romania | 2684.8 | 24.67 | 46.69 | 1476 | 0.0270 |
| R10832.SG | 1075 | Europe | Lithuania | 3582.1 | 23.90 | 54.90 | 1475 | 0.0279 |
| R10836.SG | 1075 | Europe | Lithuania | 3582.1 | 23.90 | 54.90 | 1475 | 0.0242 |
| R10838.SG | 1075 | Europe | Lithuania | 3582.1 | 23.90 | 54.90 | 1475 | 0.0206 |
| R2050.SG | 1091 | Europe | Croatia | 3052.4 | 14.54 | 45.20 | 1475 | 0.0250 |

| | | | | | | | | |
|---|---|---|---|---|---|---|---|---|
| R2051.SG | 1091 | Europe | Croatia | 3052.4 | 14.54 | 45.20 | 1475 | 0.0145 |
| R11544.SG | 353 | Asia | Armenia | 1927.6 | 43.85 | 40.69 | 1470 | 0.0204 |
| I0774 | 582 | Europe | United Kingdom | 4389.8 | 0.07 | 52.26 | 1469 | 0.0213 |
| R11543.SG | 353 | Asia | Armenia | 1927.6 | 43.85 | 40.69 | 1468 | 0.0163 |
| I3576 | 847 | Europe | Spain | 4878.1 | -4.01 | 37.32 | 1468 | 0.0234 |
| SZ27 | 475 | Europe | Hungary | 2919.0 | 17.85 | 46.28 | 1467 | 0.0232 |
| I14095 | 656 | Europe | United Kingdom | 4668.7 | -1.61 | 54.40 | 1466 | 0.0196 |
| I14839 | 671 | Europe | Turkey | 1775.5 | 29.71 | 40.42 | 1466 | 0.0214 |
| R3545.SG | 1102 | Europe | Croatia | 2728.3 | 16.71 | 43.61 | 1466 | 0.0217 |
| R11538.SG | 353 | Asia | Armenia | 1927.6 | 43.85 | 40.69 | 1465 | 0.0209 |
| R11542.SG | 353 | Asia | Armenia | 1927.6 | 43.85 | 40.69 | 1465 | 0.0145 |
| R12229.SG | 433 | Both | Lebanon | 727.3 | 35.83 | 33.85 | 1465 | 0.0163 |
| I0777 | 582 | Europe | United Kingdom | 4389.8 | 0.07 | 52.26 | 1465 | 0.0157 |
| SZ13 | 475 | Europe | Hungary | 2919.0 | 17.85 | 46.28 | 1462 | 0.0171 |
| I16113 | 651 | Europe | Czech Republic | 3582.7 | 13.62 | 50.53 | 1462 | 0.0159 |
| R2055.SG | 1092 | Europe | France | 3819.7 | 6.17 | 49.12 | 1461 | 0.0246 |
| DA385_noUDG.SG | 96 | Asia | Kyrgyzstan | 4317.1 | 75.79 | 41.50 | 1458 | 0.0255 |
| MDM002.SG | 998 | Europe | Netherlands | 4188.7 | 5.44 | 53.18 | 1458 | 0.0192 |
| SZ19 | 475 | Europe | Hungary | 2919.0 | 17.85 | 46.28 | 1456 | 0.0178 |
| R3660.SG | 1105 | Europe | Croatia | 2983.9 | 16.12 | 45.77 | 1456 | 0.0190 |
| R3477.SG | 435 | Both | Lebanon | 727.3 | 35.50 | 33.62 | 1450 | 0.0264 |
| I11585 | 612 | Europe | United Kingdom | 4586.2 | -0.61 | 54.17 | 1450 | 0.0291 |
| I11587 | 612 | Europe | United Kingdom | 4586.2 | -0.61 | 54.17 | 1450 | 0.0227 |
| I11589 | 612 | Europe | United Kingdom | 4586.2 | -0.61 | 54.17 | 1450 | 0.0172 |
| I11590 | 612 | Europe | United Kingdom | 4586.2 | -0.61 | 54.17 | 1450 | 0.0113 |
| I20640 | 612 | Europe | United Kingdom | 4586.2 | -0.61 | 54.17 | 1450 | 0.0176 |
| I20641 | 612 | Europe | United Kingdom | 4586.2 | -0.61 | 54.17 | 1450 | 0.0223 |
| I20644 | 612 | Europe | United Kingdom | 4586.2 | -0.61 | 54.17 | 1450 | 0.0182 |
| I20646 | 612 | Europe | United Kingdom | 4586.2 | -0.61 | 54.17 | 1450 | 0.0206 |
| I20649 | 612 | Europe | United Kingdom | 4586.2 | -0.61 | 54.17 | 1450 | 0.0170 |
| I20650 | 612 | Europe | United Kingdom | 4586.2 | -0.61 | 54.17 | 1450 | 0.0185 |

| | | | | | | | | |
|---|---|---|---|---|---|---|---|---|
| I20652 | 612 | Europe | United Kingdom | 4586.2 | -0.61 | 54.17 | 1450 | 0.0203 |
| I20654 | 612 | Europe | United Kingdom | 4586.2 | -0.61 | 54.17 | 1450 | 0.0127 |
| I20655 | 612 | Europe | United Kingdom | 4586.2 | -0.61 | 54.17 | 1450 | 0.0223 |
| I20656 | 612 | Europe | United Kingdom | 4586.2 | -0.61 | 54.17 | 1450 | 0.0240 |
| I20658 | 612 | Europe | United Kingdom | 4586.2 | -0.61 | 54.17 | 1450 | 0.0186 |
| I20659 | 612 | Europe | United Kingdom | 4586.2 | -0.61 | 54.17 | 1450 | 0.0250 |
| I20660 | 612 | Europe | United Kingdom | 4586.2 | -0.61 | 54.17 | 1450 | 0.0268 |
| I20661 | 612 | Europe | United Kingdom | 4586.2 | -0.61 | 54.17 | 1450 | 0.0225 |
| I20668 | 612 | Europe | United Kingdom | 4586.2 | -0.61 | 54.17 | 1450 | 0.0219 |
| I20670 | 612 | Europe | United Kingdom | 4586.2 | -0.61 | 54.17 | 1450 | 0.0160 |
| I20671 | 612 | Europe | United Kingdom | 4586.2 | -0.61 | 54.17 | 1450 | 0.0200 |
| I20673 | 612 | Europe | United Kingdom | 4586.2 | -0.61 | 54.17 | 1450 | 0.0174 |
| I20674 | 612 | Europe | United Kingdom | 4586.2 | -0.61 | 54.17 | 1450 | 0.0126 |
| I20677 | 612 | Europe | United Kingdom | 4586.2 | -0.61 | 54.17 | 1450 | 0.0182 |
| I20678 | 612 | Europe | United Kingdom | 4586.2 | -0.61 | 54.17 | 1450 | 0.0241 |
| I20679 | 612 | Europe | United Kingdom | 4586.2 | -0.61 | 54.17 | 1450 | 0.0209 |
| I20680 | 612 | Europe | United Kingdom | 4586.2 | -0.61 | 54.17 | 1450 | 0.0178 |
| I3981 | 853 | Europe | Spain | 4878.1 | -3.61 | 37.18 | 1450 | 0.0216 |
| R105.SG | 1067 | Europe | Italy | 3097.9 | 12.48 | 41.89 | 1450 | 0.0244 |
| R106.SG | 1067 | Europe | Italy | 3097.9 | 12.48 | 41.89 | 1450 | 0.0179 |
| R107.SG | 1067 | Europe | Italy | 3097.9 | 12.48 | 41.89 | 1450 | 0.0202 |
| R108.SG | 1067 | Europe | Italy | 3097.9 | 12.48 | 41.89 | 1450 | 0.0245 |
| R110.SG | 1067 | Europe | Italy | 3097.9 | 12.48 | 41.89 | 1450 | 0.0220 |
| R117.SG | 1080 | Europe | Italy | 3086.5 | 12.29 | 41.76 | 1450 | 0.0215 |
| R118.SG | 1080 | Europe | Italy | 3086.5 | 12.29 | 41.76 | 1450 | 0.0198 |
| R120.SG | 1080 | Europe | Italy | 3086.5 | 12.29 | 41.76 | 1450 | 0.0199 |
| R121.SG | 1080 | Europe | Italy | 3086.5 | 12.29 | 41.76 | 1450 | 0.0183 |
| R122.SG | 1080 | Europe | Italy | 3086.5 | 12.29 | 41.76 | 1450 | 0.0221 |
| R30.SG | 1098 | Europe | Italy | 3097.9 | 12.47 | 41.91 | 1450 | 0.0207 |
| R32.SG | 1098 | Europe | Italy | 3097.9 | 12.47 | 41.91 | 1450 | 0.0286 |
| R33.SG | 1098 | Europe | Italy | 3097.9 | 12.47 | 41.91 | 1450 | 0.0178 |

| | | | | | | | | |
|---|---|---|---|---|---|---|---|---|
| R34.SG | 1098 | Europe | Italy | 3097.9 | 12.47 | 41.91 | 1450 | 0.0222 |
| R35.SG | 1100 | Europe | Italy | 3097.9 | 12.49 | 41.89 | 1450 | 0.0237 |
| R36.SG | 1100 | Europe | Italy | 3097.9 | 12.49 | 41.89 | 1450 | 0.0207 |
| R3685.SG | 1106 | Europe | Croatia | 2728.3 | 16.80 | 43.62 | 1450 | 0.0220 |
| VK390_noUDG.SG | 1221 | Europe | Norway | 4538.9 | 9.59 | 59.22 | 1450 | 0.0262 |
| I0769 | 582 | Europe | United Kingdom | 4389.8 | 0.07 | 52.26 | 1449 | 0.0214 |
| SZ11.SG | 475 | Europe | Hungary | 2919.0 | 17.85 | 46.28 | 1442 | 0.0218 |
| SZ12 | 475 | Europe | Hungary | 2919.0 | 17.85 | 46.28 | 1442 | 0.0168 |
| SZ15.SG | 475 | Europe | Hungary | 2919.0 | 17.85 | 46.28 | 1442 | 0.0227 |
| SZ16 | 475 | Europe | Hungary | 2919.0 | 17.85 | 46.28 | 1442 | 0.0172 |
| SZ18 | 475 | Europe | Hungary | 2919.0 | 17.85 | 46.28 | 1442 | 0.0158 |
| SZ2.SG | 475 | Europe | Hungary | 2919.0 | 17.85 | 46.28 | 1442 | 0.0172 |
| SZ22 | 475 | Europe | Hungary | 2919.0 | 17.85 | 46.28 | 1442 | 0.0199 |
| SZ23 | 475 | Europe | Hungary | 2919.0 | 17.85 | 46.28 | 1442 | 0.0177 |
| SZ28 | 475 | Europe | Hungary | 2919.0 | 17.85 | 46.28 | 1442 | 0.0221 |
| SZ3.SG | 475 | Europe | Hungary | 2919.0 | 17.85 | 46.28 | 1442 | 0.0192 |
| SZ32 | 475 | Europe | Hungary | 2919.0 | 17.85 | 46.28 | 1442 | 0.0236 |
| SZ36.SG | 475 | Europe | Hungary | 2919.0 | 17.85 | 46.28 | 1442 | 0.0229 |
| SZ38 | 475 | Europe | Hungary | 2919.0 | 17.85 | 46.28 | 1442 | 0.0214 |
| SZ4.SG | 475 | Europe | Hungary | 2919.0 | 17.85 | 46.28 | 1442 | 0.0195 |
| SZ40 | 475 | Europe | Hungary | 2919.0 | 17.85 | 46.28 | 1442 | 0.0152 |
| SZ42 | 475 | Europe | Hungary | 2919.0 | 17.85 | 46.28 | 1442 | 0.0255 |
| SZ45.SG | 475 | Europe | Hungary | 2919.0 | 17.85 | 46.28 | 1442 | 0.0174 |
| SZ5.SG | 475 | Europe | Hungary | 2919.0 | 17.85 | 46.28 | 1442 | 0.0227 |
| SZ7 | 475 | Europe | Hungary | 2919.0 | 17.85 | 46.28 | 1442 | 0.0213 |
| SZ9 | 475 | Europe | Hungary | 2919.0 | 17.85 | 46.28 | 1442 | 0.0223 |
| R3543.SG | 1102 | Europe | Croatia | 2728.3 | 16.71 | 43.61 | 1438 | 0.0270 |
| R3476.SG | 435 | Both | Lebanon | 727.3 | 35.50 | 33.62 | 1431 | 0.0206 |
| KV-3369.SG | 982 | Europe | Hungary | 2915.5 | 19.44 | 46.62 | 1417 | 0.0247 |
| MDM005.SG | 998 | Europe | Netherlands | 4188.7 | 5.44 | 53.18 | 1417 | 0.0201 |
| PV-125.SG | 1060 | Europe | Hungary | 2776.0 | 20.86 | 46.30 | 1417 | 0.0261 |

| | | | | | | | | |
|---|---|---|---|---|---|---|---|---|
| MM-240.SG | 1001 | Europe | Hungary | 2823.1 | 20.52 | 46.22 | 1410 | 0.0198 |
| QED-4_noUDG.SG | 432 | Both | Lebanon | 776.6 | 35.78 | 34.05 | 1406 | 0.0196 |
| RISE174_noUDG.SG | 1126 | Europe | Sweden | 4060.0 | 13.10 | 55.55 | 1405 | 0.0228 |
| R10503.SG | 1071 | Europe | Portugal | 5047.7 | -9.37 | 38.73 | 1401 | 0.0167 |
| CHN007 | 65 | Asia | Mongolia | 5649.6 | 92.05 | 50.00 | 1400 | 0.0286 |
| CHN008 | 65 | Asia | Mongolia | 5649.6 | 92.05 | 50.00 | 1400 | 0.0189 |
| CHN012 | 65 | Asia | Mongolia | 5649.6 | 92.05 | 50.00 | 1400 | 0.0247 |
| CHN016 | 65 | Asia | Mongolia | 5649.6 | 92.05 | 50.00 | 1400 | 0.0254 |
| I14842 | 671 | Europe | Turkey | 1775.5 | 29.71 | 40.42 | 1400 | 0.0211 |
| I14843 | 671 | Europe | Turkey | 1775.5 | 29.71 | 40.42 | 1400 | 0.0220 |
| I17272 | 713 | Europe | United Kingdom | 4635.1 | -1.31 | 54.59 | 1400 | 0.0195 |
| I17273 | 713 | Europe | United Kingdom | 4635.1 | -1.31 | 54.59 | 1400 | 0.0223 |
| I17275 | 713 | Europe | United Kingdom | 4635.1 | -1.31 | 54.59 | 1400 | 0.0211 |
| I17276 | 713 | Europe | United Kingdom | 4635.1 | -1.31 | 54.59 | 1400 | 0.0178 |
| MDM001.A | 998 | Europe | Netherlands | 4188.7 | 5.44 | 53.18 | 1400 | 0.0228 |
| OAI016 | 1024 | Europe | United Kingdom | 4389.8 | 0.10 | 52.20 | 1400 | 0.0276 |
| SZ43.SG | 475 | Europe | Hungary | 2919.0 | 17.85 | 46.28 | 1393 | 0.0229 |
| I12031 | 624 | Europe | Spain | 3994.5 | 2.81 | 42.02 | 1392 | 0.0293 |
| I12034 | 624 | Europe | Spain | 3994.5 | 2.81 | 42.02 | 1392 | 0.0224 |
| I14091 | 656 | Europe | United Kingdom | 4668.7 | -1.61 | 54.40 | 1392 | 0.0172 |
| MM-245.SG | 1001 | Europe | Hungary | 2823.1 | 20.52 | 46.22 | 1391 | 0.0256 |
| R3547.SG | 1103 | Europe | Croatia | 3052.4 | 13.66 | 45.41 | 1381 | 0.0225 |
| R3544.SG | 1102 | Europe | Croatia | 2728.3 | 16.71 | 43.61 | 1374 | 0.0246 |
| BandaKD11 | 17 | Asia | China | 7617.5 | 107.50 | 24.05 | 1372 | 0.0184 |
| LadaKH01 | 316 | Asia | China | 7617.5 | 107.60 | 24.00 | 1372 | 0.0239 |
| KFP-7.SG | 965 | Europe | Hungary | 2960.2 | 19.38 | 47.22 | 1367 | 0.0242 |
| FGD-4.SG | 536 | Europe | Hungary | 2915.5 | 18.98 | 46.48 | 1365 | 0.0217 |
| AV1 | 475 | Europe | Hungary | 2919.0 | 17.85 | 46.28 | 1360 | 0.0173 |
| CSPF-114.SG | 513 | Europe | Hungary | 2868.4 | 19.89 | 46.44 | 1357 | 0.0255 |
| DA228_noUDG.SG | 84 | Asia | Kazakhstan | 3667.1 | 68.25 | 43.07 | 1350 | 0.0218 |
| I4613 | 237 | Asia | Turkey | 1449.8 | 41.37 | 37.42 | 1350 | 0.0193 |

| | | | | | | | | |
|---|---|---|---|---|---|---|---|---|
| CSB-9.SG | 511 | Europe | Hungary | 2868.4 | 20.25 | 46.71 | 1350 | 0.0245 |
| I14534 | 662 | Europe | United Kingdom | 4296.4 | 0.08 | 50.79 | 1350 | 0.0189 |
| I14535 | 662 | Europe | United Kingdom | 4296.4 | 0.08 | 50.79 | 1350 | 0.0245 |
| I14536 | 662 | Europe | United Kingdom | 4296.4 | 0.08 | 50.79 | 1350 | 0.0236 |
| I14538 | 662 | Europe | United Kingdom | 4296.4 | 0.08 | 50.79 | 1350 | 0.0225 |
| I14541 | 662 | Europe | United Kingdom | 4296.4 | 0.08 | 50.79 | 1350 | 0.0254 |
| I14542 | 662 | Europe | United Kingdom | 4296.4 | 0.08 | 50.79 | 1350 | 0.0200 |
| I8366 | 671 | Europe | Turkey | 1775.5 | 29.71 | 40.42 | 1350 | 0.0270 |
| I8367 | 671 | Europe | Turkey | 1775.5 | 29.71 | 40.42 | 1350 | 0.0271 |
| I8369 | 671 | Europe | Turkey | 1775.5 | 29.71 | 40.42 | 1350 | 0.0244 |
| I8370 | 671 | Europe | Turkey | 1775.5 | 29.71 | 40.42 | 1350 | 0.0232 |
| I8371 | 671 | Europe | Turkey | 1775.5 | 29.71 | 40.42 | 1350 | 0.0213 |
| I8373 | 671 | Europe | Turkey | 1775.5 | 29.71 | 40.42 | 1350 | 0.0207 |
| I18174 | 720 | Europe | Hungary | 2913.5 | 20.32 | 47.20 | 1350 | 0.0213 |
| I18184 | 720 | Europe | Hungary | 2913.5 | 20.32 | 47.20 | 1350 | 0.0133 |
| IND001.A_noUDG | 954 | Europe | Germany | 3957.6 | 6.36 | 50.85 | 1350 | 0.0171 |
| IND002.A_noUDG | 954 | Europe | Germany | 3957.6 | 6.36 | 50.85 | 1350 | 0.0217 |
| IND003.A_noUDG | 954 | Europe | Germany | 3957.6 | 6.36 | 50.85 | 1350 | 0.0168 |
| IND004.A_noUDG | 954 | Europe | Germany | 3957.6 | 6.36 | 50.85 | 1350 | 0.0185 |
| IND005.A_noUDG | 954 | Europe | Germany | 3957.6 | 6.36 | 50.85 | 1350 | 0.0193 |
| IND006.A_noUDG | 954 | Europe | Germany | 3957.6 | 6.36 | 50.85 | 1350 | 0.0175 |
| IND007.A_noUDG | 954 | Europe | Germany | 3957.6 | 6.36 | 50.85 | 1350 | 0.0241 |
| IND008.A_noUDG | 954 | Europe | Germany | 3957.6 | 6.36 | 50.85 | 1350 | 0.0157 |
| IND009.A_noUDG | 954 | Europe | Germany | 3957.6 | 6.36 | 50.85 | 1350 | 0.0232 |
| IND010.A_noUDG | 954 | Europe | Germany | 3957.6 | 6.36 | 50.85 | 1350 | 0.0216 |
| IND012.A_noUDG | 954 | Europe | Germany | 3957.6 | 6.36 | 50.85 | 1350 | 0.0177 |
| IND013.A_noUDG | 954 | Europe | Germany | 3957.6 | 6.36 | 50.85 | 1350 | 0.0229 |
| IND014.A_noUDG | 954 | Europe | Germany | 3957.6 | 6.36 | 50.85 | 1350 | 0.0153 |
| IND017.A_noUDG | 954 | Europe | Germany | 3957.6 | 6.36 | 50.85 | 1350 | 0.0293 |
| AV2 | 475 | Europe | Hungary | 2919.0 | 17.85 | 46.28 | 1348 | 0.0160 |
| I20259 | 758 | Europe | Turkey | 1573.9 | 28.04 | 37.34 | 1346 | 0.0149 |

| | | | | | | | | |
|---|---|---|---|---|---|---|---|---|
| I20260 | 758 | Europe | Turkey | 1573.9 | 28.04 | 37.34 | 1346 | 0.0137 |
| I20264 | 758 | Europe | Turkey | 1573.9 | 28.04 | 37.34 | 1346 | 0.0215 |
| I20265 | 758 | Europe | Turkey | 1573.9 | 28.04 | 37.34 | 1346 | 0.0152 |
| I20266 | 758 | Europe | Turkey | 1573.9 | 28.04 | 37.34 | 1346 | 0.0148 |
| CL121 | 506 | Europe | Italy | 3505.4 | 7.58 | 45.08 | 1345 | 0.0158 |
| CL145 | 506 | Europe | Italy | 3505.4 | 7.58 | 45.08 | 1345 | 0.0241 |
| CL146 | 506 | Europe | Italy | 3505.4 | 7.58 | 45.08 | 1345 | 0.0219 |
| CL151 | 506 | Europe | Italy | 3505.4 | 7.58 | 45.08 | 1345 | 0.0162 |
| CL23 | 506 | Europe | Italy | 3505.4 | 7.58 | 45.08 | 1345 | 0.0231 |
| CL25 | 506 | Europe | Italy | 3505.4 | 7.58 | 45.08 | 1345 | 0.0140 |
| CL30 | 506 | Europe | Italy | 3505.4 | 7.58 | 45.08 | 1345 | 0.0197 |
| CL36 | 506 | Europe | Italy | 3505.4 | 7.58 | 45.08 | 1345 | 0.0207 |
| CL38 | 506 | Europe | Italy | 3505.4 | 7.58 | 45.08 | 1345 | 0.0161 |
| CL47 | 506 | Europe | Italy | 3505.4 | 7.58 | 45.08 | 1345 | 0.0133 |
| CL57 | 506 | Europe | Italy | 3505.4 | 7.58 | 45.08 | 1345 | 0.0217 |
| CL63 | 506 | Europe | Italy | 3505.4 | 7.58 | 45.08 | 1345 | 0.0197 |
| CL83 | 506 | Europe | Italy | 3505.4 | 7.58 | 45.08 | 1345 | 0.0200 |
| CL84 | 506 | Europe | Italy | 3505.4 | 7.58 | 45.08 | 1345 | 0.0168 |
| CL87 | 506 | Europe | Italy | 3505.4 | 7.58 | 45.08 | 1345 | 0.0229 |
| CL92 | 506 | Europe | Italy | 3505.4 | 7.58 | 45.08 | 1345 | 0.0227 |
| CL93 | 506 | Europe | Italy | 3505.4 | 7.58 | 45.08 | 1345 | 0.0208 |
| CL94 | 506 | Europe | Italy | 3505.4 | 7.58 | 45.08 | 1345 | 0.0187 |
| NIEcap1 | 1020 | Europe | Germany | 3533.3 | 10.23 | 48.54 | 1345 | 0.0235 |
| NIEcap3b | 1020 | Europe | Germany | 3533.3 | 10.23 | 48.54 | 1345 | 0.0247 |
| R9818.SG | 435 | Both | Lebanon | 727.3 | 35.50 | 33.62 | 1340 | 0.0233 |
| MM-61.SG | 1001 | Europe | Hungary | 2823.1 | 20.52 | 46.22 | 1333 | 0.0169 |
| MM-80.SG | 1001 | Europe | Hungary | 2823.1 | 20.52 | 46.22 | 1328 | 0.0272 |
| R9823.SG | 435 | Both | Lebanon | 727.3 | 35.50 | 33.62 | 1327 | 0.0236 |
| A1824 | 450 | Europe | Hungary | 2919.0 | 18.70 | 45.95 | 1325 | 0.0152 |
| I16744 | 703 | Europe | Hungary | 2958.8 | 21.47 | 47.67 | 1325 | 0.0205 |
| I16750 | 704 | Europe | Hungary | 2958.8 | 21.50 | 47.70 | 1325 | 0.0162 |

| | | | | | | | | |
|---|---|---|---|---|---|---|---|---|
| I16812 | 708 | Europe | Hungary | 2913.1 | 21.54 | 47.35 | 1325 | 0.0256 |
| I20800 | 766 | Europe | Hungary | 2913.1 | 21.53 | 47.37 | 1325 | 0.0180 |
| I20801 | 766 | Europe | Hungary | 2913.1 | 21.53 | 47.37 | 1325 | 0.0182 |
| MM-131.SG | 1001 | Europe | Hungary | 2823.1 | 20.52 | 46.22 | 1325 | 0.0262 |
| MM-151.SG | 1001 | Europe | Hungary | 2823.1 | 20.52 | 46.22 | 1325 | 0.0241 |
| MM-83.SG | 1001 | Europe | Hungary | 2823.1 | 20.52 | 46.22 | 1325 | 0.0193 |
| SZOD1-127.SG | 1149 | Europe | Hungary | 2868.4 | 20.32 | 46.72 | 1325 | 0.0222 |
| SZOD1-76.SG | 1149 | Europe | Hungary | 2868.4 | 20.32 | 46.72 | 1325 | 0.0213 |
| ZAA002 | 419 | Asia | Mongolia | 6542.4 | 104.05 | 47.85 | 1324 | 0.0211 |
| I16510 | 698 | Europe | United Kingdom | 4397.4 | -0.82 | 52.06 | 1324 | 0.0191 |
| MS-45.SG | 1003 | Europe | Hungary | 2870.8 | 19.53 | 46.33 | 1322 | 0.0253 |
| R104.SG | 1067 | Europe | Italy | 3097.9 | 12.48 | 41.89 | 1319 | 0.0237 |
| ALT-77.SG | 462 | Europe | Hungary | 2958.7 | 20.12 | 47.51 | 1314 | 0.0189 |
| I18743 | 738 | Europe | Hungary | 3004.9 | 19.64 | 47.25 | 1313 | 0.0250 |
| I16509 | 698 | Europe | United Kingdom | 4397.4 | -0.82 | 52.06 | 1311 | 0.0217 |
| KPM-14.SG | 979 | Europe | Hungary | 2915.5 | 19.44 | 46.75 | 1310 | 0.0216 |
| ShenxianKP09 | 379 | Asia | China | 7636.3 | 107.59 | 23.31 | 1308 | 0.0231 |
| I4474 | 866 | Europe | Turkey | 1394.0 | 40.95 | 37.18 | 1308 | 0.0135 |
| AN-286.SG | 463 | Europe | Hungary | 2823.1 | 20.63 | 46.25 | 1305 | 0.0266 |
| AN-376.SG | 463 | Europe | Hungary | 2823.1 | 20.63 | 46.25 | 1305 | 0.0190 |
| CS-465.SG | 510 | Europe | Hungary | 3004.9 | 19.10 | 47.46 | 1305 | 0.0226 |
| DK-701.SG | 521 | Europe | Hungary | 2960.2 | 19.03 | 47.03 | 1305 | 0.0259 |
| KFP-30a.SG | 965 | Europe | Hungary | 2960.2 | 19.38 | 47.22 | 1305 | 0.0169 |
| KFP-6.SG | 965 | Europe | Hungary | 2960.2 | 19.38 | 47.22 | 1305 | 0.0235 |
| MS-43.SG | 1003 | Europe | Hungary | 2870.8 | 19.53 | 46.33 | 1305 | 0.0192 |
| SSD-17.SG | 1144 | Europe | Hungary | 2870.8 | 19.10 | 46.29 | 1305 | 0.0256 |
| SSD-58.SG | 1144 | Europe | Hungary | 2870.8 | 19.10 | 46.29 | 1305 | 0.0219 |
| DA162_noUDG.SG | 76 | Asia | Russia | 2206.0 | 44.54 | 43.22 | 1300 | 0.0143 |
| TAV005 | 396 | Asia | Mongolia | 7305.7 | 113.14 | 45.36 | 1300 | 0.0185 |
| TAV011 | 396 | Asia | Mongolia | 7305.7 | 113.14 | 45.36 | 1300 | 0.0226 |
| A1801 | 440 | Europe | Hungary | 3004.9 | 18.96 | 47.28 | 1300 | 0.0192 |

| | | | | | | | | |
|---|---|---|---|---|---|---|---|---|
| A1802 | 441 | Europe | Hungary | 2960.2 | 19.22 | 47.02 | 1300 | 0.0215 |
| A1804 | 442 | Europe | Hungary | 2913.5 | 20.56 | 46.86 | 1300 | 0.0180 |
| A1805 | 442 | Europe | Hungary | 2913.5 | 20.56 | 46.86 | 1300 | 0.0195 |
| A1806 | 442 | Europe | Hungary | 2913.5 | 20.56 | 46.86 | 1300 | 0.0249 |
| A1807 | 442 | Europe | Hungary | 2913.5 | 20.56 | 46.86 | 1300 | 0.0209 |
| A1812 | 443 | Europe | Hungary | 2868.4 | 19.74 | 46.75 | 1300 | 0.0223 |
| A1816 | 447 | Europe | Hungary | 2960.2 | 19.24 | 47.08 | 1300 | 0.0203 |
| A1817 | 447 | Europe | Hungary | 2960.2 | 19.24 | 47.08 | 1300 | 0.0214 |
| A1818 | 447 | Europe | Hungary | 2960.2 | 19.24 | 47.08 | 1300 | 0.0198 |
| A1821 | 447 | Europe | Hungary | 2960.2 | 19.24 | 47.08 | 1300 | 0.0202 |
| A1822 | 447 | Europe | Hungary | 2960.2 | 19.24 | 47.08 | 1300 | 0.0201 |
| A1819 | 448 | Europe | Hungary | 2868.4 | 19.83 | 46.63 | 1300 | 0.0282 |
| A1823 | 449 | Europe | Hungary | 2960.2 | 19.70 | 46.91 | 1300 | 0.0252 |
| CSPF-37.SG | 513 | Europe | Hungary | 2868.4 | 19.89 | 46.44 | 1300 | 0.0220 |
| KFP-31.SG | 965 | Europe | Hungary | 2960.2 | 19.38 | 47.22 | 1300 | 0.0198 |
| SZOD1-187.SG | 1149 | Europe | Hungary | 2868.4 | 20.32 | 46.72 | 1300 | 0.0191 |
| SZF-43.SG | 1151 | Europe | Hungary | 2868.4 | 20.12 | 46.34 | 1300 | 0.0215 |
| SZK-102.SG | 1152 | Europe | Hungary | 2823.1 | 20.26 | 46.23 | 1300 | 0.0243 |
| SZK-213.SG | 1152 | Europe | Hungary | 2823.1 | 20.26 | 46.23 | 1300 | 0.0198 |
| SZM-259.SG | 1155 | Europe | Hungary | 2868.4 | 20.27 | 46.40 | 1300 | 0.0229 |
| SZM-38.SG | 1155 | Europe | Hungary | 2868.4 | 20.27 | 46.40 | 1300 | 0.0194 |
| SZRV-54.SG | 1156 | Europe | Hungary | 2913.5 | 20.56 | 46.98 | 1300 | 0.0249 |
| MT-29.SG | 1005 | Europe | Hungary | 2870.8 | 19.38 | 46.10 | 1290 | 0.0179 |
| HH-102.SG | 555 | Europe | Hungary | 2915.5 | 19.13 | 46.54 | 1288 | 0.0238 |
| SZOD1-554.SG | 1149 | Europe | Hungary | 2868.4 | 20.32 | 46.72 | 1288 | 0.0208 |
| SZOD1-829.SG | 1149 | Europe | Hungary | 2868.4 | 20.32 | 46.72 | 1288 | 0.0232 |
| SZF-371.SG | 1151 | Europe | Hungary | 2868.4 | 20.12 | 46.34 | 1288 | 0.0190 |
| ACG-19.SG | 451 | Europe | Hungary | 2958.8 | 20.99 | 47.83 | 1286 | 0.0218 |
| MT-74.SG | 1005 | Europe | Hungary | 2870.8 | 19.38 | 46.10 | 1286 | 0.0211 |
| HC-168.SG | 551 | Europe | Hungary | 2915.5 | 19.17 | 46.49 | 1285 | 0.0185 |
| R3473.SG | 435 | Both | Lebanon | 727.3 | 35.50 | 33.62 | 1279 | 0.0245 |

| | | | | | | | | |
|---|---|---|---|---|---|---|---|---|
| I16508 | 698 | Europe | United Kingdom | 4397.4 | -0.82 | 52.06 | 1276 | 0.0186 |
| S10_S13_noUDG | 372 | Asia | Nepal | 5203.2 | 84.03 | 29.28 | 1275 | 0.0213 |
| S143_S173_noUDG | 372 | Asia | Nepal | 5203.2 | 84.03 | 29.28 | 1275 | 0.0208 |
| S153_S183_noUDG | 372 | Asia | Nepal | 5203.2 | 84.03 | 29.28 | 1275 | 0.0253 |
| S18_S20_S21_S22_noUDG | 372 | Asia | Nepal | 5203.2 | 84.03 | 29.28 | 1275 | 0.0179 |
| S8_noUDG | 372 | Asia | Nepal | 5203.2 | 84.03 | 29.28 | 1275 | 0.0254 |
| SZK-180.SG | 1152 | Europe | Hungary | 2823.1 | 20.26 | 46.23 | 1275 | 0.0252 |
| SZK-83.SG | 1152 | Europe | Hungary | 2823.1 | 20.26 | 46.23 | 1275 | 0.0284 |
| SZRV-147.SG | 1156 | Europe | Hungary | 2913.5 | 20.56 | 46.98 | 1275 | 0.0252 |
| TMH-388.SG | 1161 | Europe | Hungary | 2958.8 | 20.75 | 47.74 | 1275 | 0.0264 |
| KDA-485.SG | 963 | Europe | Hungary | 2868.4 | 20.09 | 46.38 | 1270 | 0.0207 |
| KDA-517.SG | 963 | Europe | Hungary | 2868.4 | 20.09 | 46.38 | 1270 | 0.0208 |
| KDA-520.SG | 963 | Europe | Hungary | 2868.4 | 20.09 | 46.38 | 1270 | 0.0229 |
| KK1-252.SG | 963 | Europe | Hungary | 2868.4 | 20.09 | 46.38 | 1270 | 0.0226 |
| KK1-541.SG | 963 | Europe | Hungary | 2868.4 | 20.09 | 46.38 | 1270 | 0.0203 |
| MS-50.SG | 1003 | Europe | Hungary | 2870.8 | 19.53 | 46.33 | 1270 | 0.0244 |
| SZM-24.SG | 1155 | Europe | Hungary | 2868.4 | 20.27 | 46.40 | 1270 | 0.0261 |
| SZF-181.SG | 1151 | Europe | Hungary | 2868.4 | 20.12 | 46.34 | 1267 | 0.0222 |
| CSPF-182.SG | 513 | Europe | Hungary | 2868.4 | 19.89 | 46.44 | 1265 | 0.0275 |
| MT-17.SG | 1005 | Europe | Hungary | 2870.8 | 19.38 | 46.10 | 1265 | 0.0248 |
| MT-23.SG | 1005 | Europe | Hungary | 2870.8 | 19.38 | 46.10 | 1265 | 0.0262 |
| SSD-144.SG | 1144 | Europe | Hungary | 2870.8 | 19.10 | 46.29 | 1265 | 0.0196 |
| SSD-151.SG | 1144 | Europe | Hungary | 2870.8 | 19.10 | 46.29 | 1265 | 0.0232 |
| SSD-198.SG | 1144 | Europe | Hungary | 2870.8 | 19.10 | 46.29 | 1265 | 0.0245 |
| SZF-26.SG | 1151 | Europe | Hungary | 2868.4 | 20.12 | 46.34 | 1263 | 0.0181 |
| SZKT-311.SG | 1153 | Europe | Hungary | 2868.4 | 20.57 | 46.56 | 1263 | 0.0262 |
| SZRV-168.SG | 1156 | Europe | Hungary | 2913.5 | 20.56 | 46.98 | 1263 | 0.0227 |
| TMH-1273.SG | 1161 | Europe | Hungary | 2958.8 | 20.75 | 47.74 | 1263 | 0.0236 |
| KV-3450.SG | 982 | Europe | Hungary | 2915.5 | 19.44 | 46.62 | 1260 | 0.0251 |
| SSD-35.SG | 1144 | Europe | Hungary | 2870.8 | 19.10 | 46.29 | 1259 | 0.0292 |
| KKB002 | 297 | Asia | Kazakhstan | 5049.9 | 84.51 | 49.21 | 1250 | 0.0273 |

| | | | | | | | | |
|---|---|---|---|---|---|---|---|---|
| NOM001 | 344 | Asia | Mongolia | 6472.4 | 103.15 | 47.45 | 1250 | 0.0182 |
| TUM001 | 401 | Asia | Mongolia | 6542.4 | 104.64 | 47.96 | 1250 | 0.0214 |
| A1813 | 443 | Europe | Hungary | 2868.4 | 19.74 | 46.75 | 1250 | 0.0248 |
| ALT-224.SG | 462 | Europe | Hungary | 2958.7 | 20.12 | 47.51 | 1250 | 0.0209 |
| ALT-442.SG | 462 | Europe | Hungary | 2958.7 | 20.12 | 47.51 | 1250 | 0.0287 |
| CSPF-213.SG | 513 | Europe | Hungary | 2868.4 | 19.89 | 46.44 | 1250 | 0.0252 |
| HH-22.SG | 555 | Europe | Hungary | 2915.5 | 19.13 | 46.54 | 1250 | 0.0233 |
| I10546 | 594 | Europe | Turkey | 1775.5 | 29.30 | 40.48 | 1250 | 0.0152 |
| I18223 | 724 | Europe | Hungary | 3004.9 | 19.63 | 47.24 | 1250 | 0.0212 |
| I18224 | 724 | Europe | Hungary | 3004.9 | 19.63 | 47.24 | 1250 | 0.0214 |
| I18744 | 724 | Europe | Hungary | 3004.9 | 19.63 | 47.24 | 1250 | 0.0153 |
| I18742 | 738 | Europe | Hungary | 3004.9 | 19.64 | 47.25 | 1250 | 0.0174 |
| JHT-154.SG | 960 | Europe | Hungary | 2958.7 | 20.14 | 47.52 | 1250 | 0.0242 |
| KPM-27.SG | 979 | Europe | Hungary | 2915.5 | 19.44 | 46.75 | 1250 | 0.0180 |
| OBT-106.SG | 1028 | Europe | Hungary | 2868.4 | 20.72 | 46.65 | 1250 | 0.0226 |
| OBT-108.SG | 1028 | Europe | Hungary | 2868.4 | 20.72 | 46.65 | 1250 | 0.0235 |
| TMH-756.SG | 1161 | Europe | Hungary | 2958.8 | 20.75 | 47.74 | 1250 | 0.0232 |
| VEN013 | 1176 | Europe | Italy | 2723.3 | 15.83 | 40.97 | 1250 | 0.0174 |
| VEN015 | 1176 | Europe | Italy | 2723.3 | 15.83 | 40.97 | 1250 | 0.0158 |
| ARK-41.SG | 469 | Europe | Hungary | 2913.1 | 21.05 | 47.59 | 1244 | 0.0238 |
| KDA-188.SG | 963 | Europe | Hungary | 2868.4 | 20.09 | 46.38 | 1244 | 0.0174 |
| VK485_noUDG.SG | 1235 | Europe | Estonia | 3974.5 | 22.25 | 58.17 | 1244 | 0.0273 |
| KK1-245.SG | 963 | Europe | Hungary | 2868.4 | 20.09 | 46.38 | 1243 | 0.0200 |
| KK1-251.SG | 963 | Europe | Hungary | 2868.4 | 20.09 | 46.38 | 1243 | 0.0243 |
| KD-29.SG | 962 | Europe | Hungary | 2913.1 | 21.39 | 47.40 | 1241 | 0.0208 |
| R10666.SG | 1074 | Europe | Austria | 3327.4 | 14.04 | 48.17 | 1241 | 0.0252 |
| SZM-255.SG | 1155 | Europe | Hungary | 2868.4 | 20.27 | 46.40 | 1240 | 0.0207 |
| VK506_noUDG.SG | 1235 | Europe | Estonia | 3974.5 | 22.25 | 58.17 | 1237 | 0.0292 |
| syr013 | 393 | Asia | Syria | 1403.1 | 41.46 | 36.83 | 1235 | 0.0147 |
| R10665.SG | 1074 | Europe | Austria | 3327.4 | 14.04 | 48.17 | 1235 | 0.0230 |
| I0159 | 572 | Europe | United Kingdom | 4343.0 | 0.18 | 52.08 | 1234 | 0.0244 |

| | | | | | | | | |
|---|---|---|---|---|---|---|---|---|
| ARK-38.SG | 469 | Europe | Hungary | 2913.1 | 21.05 | 47.59 | 1232 | 0.0239 |
| I15742 | 683 | Europe | Croatia | 2810.3 | 16.26 | 43.53 | 1232 | 0.0253 |
| TSB001 | 347 | Asia | Mongolia | 6401.6 | 102.08 | 47.11 | 1231 | 0.0281 |
| HH-10.SG | 555 | Europe | Hungary | 2915.5 | 19.13 | 46.54 | 1231 | 0.0236 |
| KK1-368.SG | 963 | Europe | Hungary | 2868.4 | 20.09 | 46.38 | 1230 | 0.0250 |
| OBT-51.SG | 1028 | Europe | Hungary | 2868.4 | 20.72 | 46.65 | 1230 | 0.0231 |
| RISE569_noUDG.SG | 933 | Europe | Czech Republic | 3495.2 | 14.16 | 50.19 | 1229 | 0.0194 |
| I15744 | 683 | Europe | Croatia | 2810.3 | 16.26 | 43.53 | 1228 | 0.0137 |
| I4475 | 866 | Europe | Turkey | 1394.0 | 40.95 | 37.18 | 1228 | 0.0254 |
| KPM-23.SG | 979 | Europe | Hungary | 2915.5 | 19.44 | 46.75 | 1228 | 0.0167 |
| OBH-37.SG | 1025 | Europe | Hungary | 2868.4 | 20.74 | 46.60 | 1228 | 0.0204 |
| ZAA007 | 419 | Asia | Mongolia | 6542.4 | 104.05 | 47.85 | 1227 | 0.0207 |
| FU-193.SG | 539 | Europe | Hungary | 2868.4 | 20.20 | 46.77 | 1227 | 0.0237 |
| FU-215.SG | 539 | Europe | Hungary | 2868.4 | 20.20 | 46.77 | 1227 | 0.0194 |
| I0157 | 572 | Europe | United Kingdom | 4343.0 | 0.18 | 52.08 | 1227 | 0.0179 |
| SZKT-89.SG | 1153 | Europe | Hungary | 2868.4 | 20.57 | 46.56 | 1227 | 0.0260 |
| VK490_noUDG.SG | 1235 | Europe | Estonia | 3974.5 | 22.25 | 58.17 | 1227 | 0.0132 |
| ARK-36.SG | 469 | Europe | Hungary | 2913.1 | 21.05 | 47.59 | 1226 | 0.0242 |
| PV-205.SG | 1060 | Europe | Hungary | 2776.0 | 20.86 | 46.30 | 1226 | 0.0228 |
| UGU001 | 404 | Asia | Mongolia | 6600.9 | 105.38 | 49.25 | 1225 | 0.0182 |
| I16759 | 706 | Europe | Hungary | 2919.0 | 18.72 | 46.21 | 1225 | 0.0171 |
| I18222 | 724 | Europe | Hungary | 3004.9 | 19.63 | 47.24 | 1225 | 0.0201 |
| I18225 | 724 | Europe | Hungary | 3004.9 | 19.63 | 47.24 | 1225 | 0.0196 |
| SZM-332.SG | 1155 | Europe | Hungary | 2868.4 | 20.27 | 46.40 | 1225 | 0.0202 |
| ARK-24.SG | 469 | Europe | Hungary | 2913.1 | 21.05 | 47.59 | 1224 | 0.0218 |
| PV-200.SG | 1060 | Europe | Hungary | 2776.0 | 20.86 | 46.30 | 1223 | 0.0220 |
| TMH-199.SG | 1161 | Europe | Hungary | 2958.8 | 20.75 | 47.74 | 1223 | 0.0272 |
| I4532 | 866 | Europe | Turkey | 1394.0 | 40.95 | 37.18 | 1222 | 0.0221 |
| ARK-29.SG | 469 | Europe | Hungary | 2913.1 | 21.05 | 47.59 | 1220 | 0.0248 |
| I10430 | 594 | Europe | Turkey | 1775.5 | 29.30 | 40.48 | 1218 | 0.0290 |
| ARK-50.SG | 469 | Europe | Hungary | 2913.1 | 21.05 | 47.59 | 1217 | 0.0279 |

| | | | | | | | | |
|---|---|---|---|---|---|---|---|---|
| I15462 | 683 | Europe | Croatia | 2810.3 | 16.26 | 43.53 | 1216 | 0.0190 |
| R3359.SG | 434 | Both | Syria | 1158.9 | 40.56 | 34.97 | 1209 | 0.0196 |
| ARK-43.SG | 469 | Europe | Hungary | 2913.1 | 21.05 | 47.59 | 1205 | 0.0212 |
| A1810 | 443 | Europe | Hungary | 2868.4 | 19.74 | 46.75 | 1200 | 0.0216 |
| A1811 | 443 | Europe | Hungary | 2868.4 | 19.74 | 46.75 | 1200 | 0.0194 |
| A1815 | 443 | Europe | Hungary | 2868.4 | 19.74 | 46.75 | 1200 | 0.0216 |
| ARK-17.SG | 469 | Europe | Hungary | 2913.1 | 21.05 | 47.59 | 1200 | 0.0189 |
| ARK-6.SG | 469 | Europe | Hungary | 2913.1 | 21.05 | 47.59 | 1200 | 0.0255 |
| DRU001.A_noUDG | 525 | Europe | Germany | 4011.8 | 8.19 | 52.81 | 1200 | 0.0212 |
| DRU002.A_noUDG | 525 | Europe | Germany | 4011.8 | 8.19 | 52.81 | 1200 | 0.0157 |
| DRU003.A_noUDG | 525 | Europe | Germany | 4011.8 | 8.19 | 52.81 | 1200 | 0.0170 |
| DRU004.A_noUDG | 525 | Europe | Germany | 4011.8 | 8.19 | 52.81 | 1200 | 0.0149 |
| DRU006.A_noUDG | 525 | Europe | Germany | 4011.8 | 8.19 | 52.81 | 1200 | 0.0225 |
| DRU007.A_noUDG | 525 | Europe | Germany | 4011.8 | 8.19 | 52.81 | 1200 | 0.0219 |
| DRU011.A_noUDG | 525 | Europe | Germany | 4011.8 | 8.19 | 52.81 | 1200 | 0.0136 |
| DRU012.A_noUDG | 525 | Europe | Germany | 4011.8 | 8.19 | 52.81 | 1200 | 0.0192 |
| DRU013.A_noUDG | 525 | Europe | Germany | 4011.8 | 8.19 | 52.81 | 1200 | 0.0248 |
| DRU014.A_noUDG | 525 | Europe | Germany | 4011.8 | 8.19 | 52.81 | 1200 | 0.0287 |
| DRU015.A_noUDG | 525 | Europe | Germany | 4011.8 | 8.19 | 52.81 | 1200 | 0.0162 |
| DRU016.A_noUDG | 525 | Europe | Germany | 4011.8 | 8.19 | 52.81 | 1200 | 0.0205 |
| DRU017.A_noUDG | 525 | Europe | Germany | 4011.8 | 8.19 | 52.81 | 1200 | 0.0169 |
| DRU018.A_noUDG | 525 | Europe | Germany | 4011.8 | 8.19 | 52.81 | 1200 | 0.0144 |
| EAS001.SG | 527 | Europe | United Kingdom | 4225.2 | 1.30 | 51.20 | 1200 | 0.0247 |
| EAS003.SG | 527 | Europe | United Kingdom | 4225.2 | 1.30 | 51.20 | 1200 | 0.0230 |
| EAS006_noUDG | 527 | Europe | United Kingdom | 4225.2 | 1.30 | 51.20 | 1200 | 0.0280 |
| ELY002.SG | 532 | Europe | United Kingdom | 4389.8 | 0.30 | 52.40 | 1200 | 0.0206 |
| ELY003.SG | 532 | Europe | United Kingdom | 4389.8 | 0.30 | 52.40 | 1200 | 0.0202 |
| ELY005.SG | 532 | Europe | United Kingdom | 4389.8 | 0.30 | 52.40 | 1200 | 0.0249 |
| GRO023.A | 546 | Europe | Netherlands | 4095.9 | 6.57 | 53.22 | 1200 | 0.0200 |
| HAD001.SG | 548 | Europe | United Kingdom | 4389.8 | 0.20 | 52.20 | 1200 | 0.0231 |
| HAD002.SG | 548 | Europe | United Kingdom | 4389.8 | 0.20 | 52.20 | 1200 | 0.0278 |

| | | | | | | | | |
|---|---|---|---|---|---|---|---|---|
| HAD004.SG | 548 | Europe | United Kingdom | 4389.8 | 0.20 | 52.20 | 1200 | 0.0265 |
| HAD005.SG | 548 | Europe | United Kingdom | 4389.8 | 0.20 | 52.20 | 1200 | 0.0247 |
| HAD008.SG | 548 | Europe | United Kingdom | 4389.8 | 0.20 | 52.20 | 1200 | 0.0240 |
| HAD009 | 548 | Europe | United Kingdom | 4389.8 | 0.20 | 52.20 | 1200 | 0.0192 |
| HAD010.SG | 548 | Europe | United Kingdom | 4389.8 | 0.20 | 52.20 | 1200 | 0.0204 |
| HAD011 | 548 | Europe | United Kingdom | 4389.8 | 0.20 | 52.20 | 1200 | 0.0195 |
| HAD012 | 548 | Europe | United Kingdom | 4389.8 | 0.20 | 52.20 | 1200 | 0.0233 |
| HAD013.SG | 548 | Europe | United Kingdom | 4389.8 | 0.20 | 52.20 | 1200 | 0.0215 |
| HAD014 | 548 | Europe | United Kingdom | 4389.8 | 0.20 | 52.20 | 1200 | 0.0210 |
| HAD015.SG | 548 | Europe | United Kingdom | 4389.8 | 0.20 | 52.20 | 1200 | 0.0207 |
| HAD016 | 548 | Europe | United Kingdom | 4389.8 | 0.20 | 52.20 | 1200 | 0.0221 |
| HAD017 | 548 | Europe | United Kingdom | 4389.8 | 0.20 | 52.20 | 1200 | 0.0205 |
| HAD018_noUDG | 548 | Europe | United Kingdom | 4389.8 | 0.20 | 52.20 | 1200 | 0.0218 |
| I15463 | 683 | Europe | Croatia | 2810.3 | 16.26 | 43.53 | 1200 | 0.0216 |
| I16751 | 705 | Europe | Hungary | 3003.9 | 20.02 | 47.77 | 1200 | 0.0242 |
| I16752 | 705 | Europe | Hungary | 3003.9 | 20.02 | 47.77 | 1200 | 0.0222 |
| I16753 | 705 | Europe | Hungary | 3003.9 | 20.02 | 47.77 | 1200 | 0.0195 |
| I17268 | 713 | Europe | United Kingdom | 4635.1 | -1.31 | 54.59 | 1200 | 0.0163 |
| I17270 | 713 | Europe | United Kingdom | 4635.1 | -1.31 | 54.59 | 1200 | 0.0165 |
| I17271 | 713 | Europe | United Kingdom | 4635.1 | -1.31 | 54.59 | 1200 | 0.0219 |
| I20798 | 765 | Europe | Hungary | 2913.1 | 21.51 | 47.35 | 1200 | 0.0195 |
| I20799 | 765 | Europe | Hungary | 2913.1 | 21.51 | 47.35 | 1200 | 0.0247 |
| I4534 | 866 | Europe | Turkey | 1394.0 | 40.95 | 37.18 | 1200 | 0.0147 |
| KD-16.SG | 962 | Europe | Hungary | 2913.1 | 21.39 | 47.40 | 1200 | 0.0251 |
| LAK001_noUDG | 983 | Europe | United Kingdom | 4320.7 | 0.50 | 52.40 | 1200 | 0.0219 |
| LAK002_noUDG | 983 | Europe | United Kingdom | 4320.7 | 0.50 | 52.40 | 1200 | 0.0255 |
| LAK003_noUDG | 983 | Europe | United Kingdom | 4320.7 | 0.50 | 52.40 | 1200 | 0.0222 |
| LAK004_noUDG | 983 | Europe | United Kingdom | 4320.7 | 0.50 | 52.40 | 1200 | 0.0268 |
| LAK005_noUDG | 983 | Europe | United Kingdom | 4320.7 | 0.50 | 52.40 | 1200 | 0.0285 |
| LAK006_noUDG | 983 | Europe | United Kingdom | 4320.7 | 0.50 | 52.40 | 1200 | 0.0220 |
| LAK007_noUDG | 983 | Europe | United Kingdom | 4320.7 | 0.50 | 52.40 | 1200 | 0.0157 |

| | | | | | | | | |
|---|---|---|---|---|---|---|---|---|
| LAK008_noUDG | 983 | Europe | United Kingdom | 4320.7 | 0.50 | 52.40 | 1200 | 0.0216 |
| LAK009_noUDG | 983 | Europe | United Kingdom | 4320.7 | 0.50 | 52.40 | 1200 | 0.0271 |
| LAK010_noUDG | 983 | Europe | United Kingdom | 4320.7 | 0.50 | 52.40 | 1200 | 0.0214 |
| LAK011_noUDG | 983 | Europe | United Kingdom | 4320.7 | 0.50 | 52.40 | 1200 | 0.0245 |
| LAK012_noUDG | 983 | Europe | United Kingdom | 4320.7 | 0.50 | 52.40 | 1200 | 0.0184 |
| LAK013_noUDG | 983 | Europe | United Kingdom | 4320.7 | 0.50 | 52.40 | 1200 | 0.0279 |
| LAK014_noUDG | 983 | Europe | United Kingdom | 4320.7 | 0.50 | 52.40 | 1200 | 0.0220 |
| LAK015_noUDG | 983 | Europe | United Kingdom | 4320.7 | 0.50 | 52.40 | 1200 | 0.0283 |
| OAI001.SG | 1024 | Europe | United Kingdom | 4389.8 | 0.10 | 52.20 | 1200 | 0.0260 |
| OAI003.SG | 1024 | Europe | United Kingdom | 4389.8 | 0.10 | 52.20 | 1200 | 0.0279 |
| OAI004.SG | 1024 | Europe | United Kingdom | 4389.8 | 0.10 | 52.20 | 1200 | 0.0209 |
| OAI005.SG | 1024 | Europe | United Kingdom | 4389.8 | 0.10 | 52.20 | 1200 | 0.0247 |
| OAI007.SG | 1024 | Europe | United Kingdom | 4389.8 | 0.10 | 52.20 | 1200 | 0.0202 |
| OAI008 | 1024 | Europe | United Kingdom | 4389.8 | 0.10 | 52.20 | 1200 | 0.0225 |
| OAI009 | 1024 | Europe | United Kingdom | 4389.8 | 0.10 | 52.20 | 1200 | 0.0195 |
| OAI010 | 1024 | Europe | United Kingdom | 4389.8 | 0.10 | 52.20 | 1200 | 0.0218 |
| OAI011 | 1024 | Europe | United Kingdom | 4389.8 | 0.10 | 52.20 | 1200 | 0.0201 |
| OAI012 | 1024 | Europe | United Kingdom | 4389.8 | 0.10 | 52.20 | 1200 | 0.0183 |
| OAI013 | 1024 | Europe | United Kingdom | 4389.8 | 0.10 | 52.20 | 1200 | 0.0195 |
| OAI015 | 1024 | Europe | United Kingdom | 4389.8 | 0.10 | 52.20 | 1200 | 0.0211 |
| OBH-52.SG | 1025 | Europe | Hungary | 2868.4 | 20.74 | 46.60 | 1200 | 0.0253 |
| OBT-3.SG | 1028 | Europe | Hungary | 2868.4 | 20.72 | 46.65 | 1200 | 0.0234 |
| OBT-56.SG | 1028 | Europe | Hungary | 2868.4 | 20.72 | 46.65 | 1200 | 0.0235 |
| POH002.SG | 1049 | Europe | United Kingdom | 4311.6 | 0.20 | 51.30 | 1200 | 0.0193 |
| POH003 | 1049 | Europe | United Kingdom | 4311.6 | 0.20 | 51.30 | 1200 | 0.0215 |
| POH004 | 1049 | Europe | United Kingdom | 4311.6 | 0.20 | 51.30 | 1200 | 0.0225 |
| POH005 | 1049 | Europe | United Kingdom | 4311.6 | 0.20 | 51.30 | 1200 | 0.0250 |
| POH007 | 1049 | Europe | United Kingdom | 4311.6 | 0.20 | 51.30 | 1200 | 0.0174 |
| POH008 | 1049 | Europe | United Kingdom | 4311.6 | 0.20 | 51.30 | 1200 | 0.0271 |
| POH009 | 1049 | Europe | United Kingdom | 4311.6 | 0.20 | 51.30 | 1200 | 0.0145 |
| PV-12.SG | 1060 | Europe | Hungary | 2776.0 | 20.86 | 46.30 | 1200 | 0.0215 |

| | | | | | | | | |
|---|---|---|---|---|---|---|---|---|
| SED004 | 1138 | Europe | United Kingdom | 4369.6 | 0.50 | 52.90 | 1200 | 0.0151 |
| SED005.SG | 1138 | Europe | United Kingdom | 4369.6 | 0.50 | 52.90 | 1200 | 0.0221 |
| SED008.SG | 1138 | Europe | United Kingdom | 4369.6 | 0.50 | 52.90 | 1200 | 0.0309 |
| SED013 | 1138 | Europe | United Kingdom | 4369.6 | 0.50 | 52.90 | 1200 | 0.0221 |
| SED014 | 1138 | Europe | United Kingdom | 4369.6 | 0.50 | 52.90 | 1200 | 0.0186 |
| SED018 | 1138 | Europe | United Kingdom | 4369.6 | 0.50 | 52.90 | 1200 | 0.0172 |
| SED020 | 1138 | Europe | United Kingdom | 4369.6 | 0.50 | 52.90 | 1200 | 0.0246 |
| SZK-130.SG | 1152 | Europe | Hungary | 2823.1 | 20.26 | 46.23 | 1200 | 0.0209 |
| SZKT-265.SG | 1153 | Europe | Hungary | 2868.4 | 20.57 | 46.56 | 1200 | 0.0230 |
| SZKT-62.SG | 1153 | Europe | Hungary | 2868.4 | 20.57 | 46.56 | 1200 | 0.0197 |
| SZKT-70.SG | 1153 | Europe | Hungary | 2868.4 | 20.57 | 46.56 | 1200 | 0.0207 |
| SZRV-212.SG | 1156 | Europe | Hungary | 2913.5 | 20.56 | 46.98 | 1200 | 0.0191 |
| SZRV-277.SG | 1156 | Europe | Hungary | 2913.5 | 20.56 | 46.98 | 1200 | 0.0165 |
| SZRV-316.SG | 1156 | Europe | Hungary | 2913.5 | 20.56 | 46.98 | 1200 | 0.0225 |
| SZRV-67.SG | 1156 | Europe | Hungary | 2913.5 | 20.56 | 46.98 | 1200 | 0.0172 |
| VK480_noUDG.SG | 1235 | Europe | Estonia | 3974.5 | 22.25 | 58.17 | 1200 | 0.0184 |
| VK481_noUDG.SG | 1235 | Europe | Estonia | 3974.5 | 22.25 | 58.17 | 1200 | 0.0208 |
| VK482_noUDG.SG | 1235 | Europe | Estonia | 3974.5 | 22.25 | 58.17 | 1200 | 0.0248 |
| VK484_noUDG.SG | 1235 | Europe | Estonia | 3974.5 | 22.25 | 58.17 | 1200 | 0.0243 |
| VK486_noUDG.SG | 1235 | Europe | Estonia | 3974.5 | 22.25 | 58.17 | 1200 | 0.0278 |
| VK487_noUDG.SG | 1235 | Europe | Estonia | 3974.5 | 22.25 | 58.17 | 1200 | 0.0282 |
| VK488_noUDG.SG | 1235 | Europe | Estonia | 3974.5 | 22.25 | 58.17 | 1200 | 0.0189 |
| VK489_noUDG.SG | 1235 | Europe | Estonia | 3974.5 | 22.25 | 58.17 | 1200 | 0.0217 |
| VK491_noUDG.SG | 1235 | Europe | Estonia | 3974.5 | 22.25 | 58.17 | 1200 | 0.0246 |
| VK492_noUDG.SG | 1235 | Europe | Estonia | 3974.5 | 22.25 | 58.17 | 1200 | 0.0293 |
| VK493_noUDG.SG | 1235 | Europe | Estonia | 3974.5 | 22.25 | 58.17 | 1200 | 0.0250 |
| VK495_noUDG.SG | 1235 | Europe | Estonia | 3974.5 | 22.25 | 58.17 | 1200 | 0.0160 |
| VK496_noUDG.SG | 1235 | Europe | Estonia | 3974.5 | 22.25 | 58.17 | 1200 | 0.0180 |
| VK497_noUDG.SG | 1235 | Europe | Estonia | 3974.5 | 22.25 | 58.17 | 1200 | 0.0259 |
| VK498_noUDG.SG | 1235 | Europe | Estonia | 3974.5 | 22.25 | 58.17 | 1200 | 0.0218 |
| VK504_noUDG.SG | 1235 | Europe | Estonia | 3974.5 | 22.25 | 58.17 | 1200 | 0.0281 |

| | | | | | | | | |
|---|---|---|---|---|---|---|---|---|
| VK505_noUDG.SG | 1235 | Europe | Estonia | 3974.5 | 22.25 | 58.17 | 1200 | 0.0192 |
| VK508_noUDG.SG | 1235 | Europe | Estonia | 3974.5 | 22.25 | 58.17 | 1200 | 0.0213 |
| VK509_noUDG.SG | 1235 | Europe | Estonia | 3974.5 | 22.25 | 58.17 | 1200 | 0.0239 |
| VK510_noUDG.SG | 1235 | Europe | Estonia | 3974.5 | 22.25 | 58.17 | 1200 | 0.0239 |
| VK511_noUDG.SG | 1235 | Europe | Estonia | 3974.5 | 22.25 | 58.17 | 1200 | 0.0166 |
| VK512_noUDG.SG | 1235 | Europe | Estonia | 3974.5 | 22.25 | 58.17 | 1200 | 0.0184 |
| VK549_noUDG.SG | 1235 | Europe | Estonia | 3974.5 | 22.25 | 58.17 | 1200 | 0.0188 |
| VK550_noUDG.SG | 1235 | Europe | Estonia | 3974.5 | 22.25 | 58.17 | 1200 | 0.0225 |
| VK551_noUDG.SG | 1235 | Europe | Estonia | 3974.5 | 22.25 | 58.17 | 1200 | 0.0197 |
| VK552_noUDG.SG | 1235 | Europe | Estonia | 3974.5 | 22.25 | 58.17 | 1200 | 0.0287 |
| VK553_noUDG.SG | 1235 | Europe | Estonia | 3974.5 | 22.25 | 58.17 | 1200 | 0.0212 |
| VK555_noUDG.SG | 1235 | Europe | Estonia | 3974.5 | 22.25 | 58.17 | 1200 | 0.0199 |
| VK514_noUDG.SG | 1236 | Europe | Norway | 5144.6 | 15.13 | 67.91 | 1200 | 0.0244 |
| I3585 | 848 | Europe | Spain | 4927.4 | -4.12 | 36.96 | 1195 | 0.0195 |
| PV-116.SG | 1060 | Europe | Hungary | 2776.0 | 20.86 | 46.30 | 1191 | 0.0215 |
| VK483_noUDG.SG | 1235 | Europe | Estonia | 3974.5 | 22.25 | 58.17 | 1191 | 0.0230 |
| VK301_noUDG.SG | 1200 | Europe | Denmark | 4158.0 | 10.61 | 55.43 | 1185 | 0.0220 |
| VK545_noUDG.SG | 1249 | Europe | Ireland | 4778.9 | -6.27 | 53.34 | 1185 | 0.0202 |
| VK329_noUDG.SG | 1202 | Europe | Denmark | 4199.9 | 8.76 | 55.33 | 1183 | 0.0217 |
| R10494.SG | 1070 | Europe | Portugal | 4864.7 | -7.16 | 38.88 | 1178 | 0.0194 |
| A1809 | 443 | Europe | Hungary | 2868.4 | 19.74 | 46.75 | 1175 | 0.0190 |
| A1814 | 443 | Europe | Hungary | 2868.4 | 19.74 | 46.75 | 1175 | 0.0211 |
| KK2-429.SG | 963 | Europe | Hungary | 2868.4 | 20.09 | 46.38 | 1175 | 0.0210 |
| KK2-441.SG | 963 | Europe | Hungary | 2868.4 | 20.09 | 46.38 | 1175 | 0.0279 |
| KK2-445.SG | 963 | Europe | Hungary | 2868.4 | 20.09 | 46.38 | 1175 | 0.0199 |
| KK2-670.SG | 963 | Europe | Hungary | 2868.4 | 20.09 | 46.38 | 1175 | 0.0249 |
| I3352 | 218 | Asia | India | 4788.1 | 79.74 | 30.25 | 1166 | 0.0170 |
| SZRV-266.SG | 1156 | Europe | Hungary | 2913.5 | 20.56 | 46.98 | 1162 | 0.0234 |
| I0161 | 572 | Europe | United Kingdom | 4343.0 | 0.18 | 52.08 | 1157 | 0.0319 |
| ARK-14.SG | 469 | Europe | Hungary | 2913.1 | 21.05 | 47.59 | 1156 | 0.0254 |
| ADN001_noUDG | 452 | Europe | Germany | 3881.5 | 9.87 | 52.36 | 1150 | 0.0178 |

| | | | | | | | | |
|---|---|---|---|---|---|---|---|---|
| ADN002_noUDG | 452 | Europe | Germany | 3881.5 | 9.87 | 52.36 | 1150 | 0.0194 |
| ADN003_noUDG | 452 | Europe | Germany | 3881.5 | 9.87 | 52.36 | 1150 | 0.0166 |
| ADN005_noUDG | 452 | Europe | Germany | 3881.5 | 9.87 | 52.36 | 1150 | 0.0211 |
| ADN006_noUDG | 452 | Europe | Germany | 3881.5 | 9.87 | 52.36 | 1150 | 0.0244 |
| ADN007_noUDG | 452 | Europe | Germany | 3881.5 | 9.87 | 52.36 | 1150 | 0.0195 |
| ADN008_noUDG | 452 | Europe | Germany | 3881.5 | 9.87 | 52.36 | 1150 | 0.0246 |
| ADN009_noUDG | 452 | Europe | Germany | 3881.5 | 9.87 | 52.36 | 1150 | 0.0193 |
| ADN010_noUDG | 452 | Europe | Germany | 3881.5 | 9.87 | 52.36 | 1150 | 0.0211 |
| ADN011_noUDG | 452 | Europe | Germany | 3881.5 | 9.87 | 52.36 | 1150 | 0.0176 |
| ADN013_noUDG | 452 | Europe | Germany | 3881.5 | 9.87 | 52.36 | 1150 | 0.0240 |
| ADN014_noUDG | 452 | Europe | Germany | 3881.5 | 9.87 | 52.36 | 1150 | 0.0169 |
| ADN015_noUDG | 452 | Europe | Germany | 3881.5 | 9.87 | 52.36 | 1150 | 0.0211 |
| ALT-369.SG | 462 | Europe | Hungary | 2958.7 | 20.12 | 47.51 | 1150 | 0.0198 |
| ALT-412.SG | 462 | Europe | Hungary | 2958.7 | 20.12 | 47.51 | 1150 | 0.0204 |
| ALT-414.SG | 462 | Europe | Hungary | 2958.7 | 20.12 | 47.51 | 1150 | 0.0229 |
| ALT-596.SG | 462 | Europe | Hungary | 2958.7 | 20.12 | 47.51 | 1150 | 0.0208 |
| ARK-11.SG | 469 | Europe | Hungary | 2913.1 | 21.05 | 47.59 | 1150 | 0.0212 |
| ARK-16.SG | 469 | Europe | Hungary | 2913.1 | 21.05 | 47.59 | 1150 | 0.0216 |
| ARK-19.SG | 469 | Europe | Hungary | 2913.1 | 21.05 | 47.59 | 1150 | 0.0246 |
| ARK-20.SG | 469 | Europe | Hungary | 2913.1 | 21.05 | 47.59 | 1150 | 0.0246 |
| ARK-48.SG | 469 | Europe | Hungary | 2913.1 | 21.05 | 47.59 | 1150 | 0.0199 |
| JHT-130.SG | 960 | Europe | Hungary | 2958.7 | 20.14 | 47.52 | 1150 | 0.0233 |
| SRS003.A_noUDG | 1143 | Europe | Germany | 4088.8 | 7.94 | 53.54 | 1150 | 0.0160 |
| SRS004.A_noUDG | 1143 | Europe | Germany | 4088.8 | 7.94 | 53.54 | 1150 | 0.0178 |
| TCS-2.SG | 1158 | Europe | Hungary | 3003.9 | 20.57 | 47.67 | 1150 | 0.0214 |
| TMH-509.SG | 1161 | Europe | Hungary | 2958.8 | 20.75 | 47.74 | 1150 | 0.0267 |
| TMH-798.SG | 1161 | Europe | Hungary | 2958.8 | 20.75 | 47.74 | 1150 | 0.0238 |
| VPB-307.SG | 1257 | Europe | Hungary | 3055.1 | 17.31 | 46.80 | 1150 | 0.0230 |
| VPB-31.SG | 1257 | Europe | Hungary | 3055.1 | 17.31 | 46.80 | 1150 | 0.0202 |
| KV-3456.SG | 982 | Europe | Hungary | 2915.5 | 19.44 | 46.62 | 1137 | 0.0300 |
| VK350_noUDG.SG | 1205 | Europe | Sweden | 4173.2 | 16.72 | 56.73 | 1134 | 0.0251 |

| | | | | | | | | |
|---|---|---|---|---|---|---|---|---|
| R3478.SG | 1099 | Europe | Montenegro | 2535.7 | 19.27 | 42.47 | 1133 | 0.0238 |
| TTSZ-43.SG | 1165 | Europe | Hungary | 2960.2 | 19.41 | 47.20 | 1132 | 0.0240 |
| CHK004 | 64 | Asia | Kyrgyzstan | 4490.6 | 78.38 | 42.49 | 1129 | 0.0245 |
| DA93_noUDG.SG | 84 | Asia | Kazakhstan | 3667.1 | 68.25 | 43.07 | 1126 | 0.0186 |
| RISE504_noUDG.SG | 360 | Asia | Russia | 5246.1 | 85.45 | 53.46 | 1126 | 0.0280 |
| I14622 | 666 | Europe | Albania | 2438.6 | 20.40 | 42.04 | 1121 | 0.0175 |
| JHT-30.SG | 960 | Europe | Hungary | 2958.7 | 20.14 | 47.52 | 1121 | 0.0187 |
| VK507_noUDG.SG | 1235 | Europe | Estonia | 3974.5 | 22.25 | 58.17 | 1121 | 0.0226 |
| VK554_noUDG.SG | 1235 | Europe | Estonia | 3974.5 | 22.25 | 58.17 | 1120 | 0.0216 |
| R10491.SG | 1070 | Europe | Portugal | 4864.7 | -7.16 | 38.88 | 1119 | 0.0219 |
| I4533 | 866 | Europe | Turkey | 1394.0 | 40.95 | 37.18 | 1116 | 0.0198 |
| R10496.SG | 1070 | Europe | Portugal | 4864.7 | -7.16 | 38.88 | 1116 | 0.0240 |
| VK349_noUDG.SG | 1212 | Europe | Sweden | 4093.6 | 16.43 | 56.46 | 1110 | 0.0216 |
| I6934 | 218 | Asia | India | 4788.1 | 79.74 | 30.25 | 1109 | 0.0242 |
| KKB001 | 297 | Asia | Kazakhstan | 5049.9 | 84.51 | 49.21 | 1109 | 0.0220 |
| TAQ003 | 1157 | Europe | Italy | 3097.9 | 11.77 | 42.25 | 1109 | 0.0231 |
| COR002 | 508 | Europe | Italy | 3570.9 | 8.79 | 40.52 | 1106 | 0.0222 |
| VK328_noUDG.SG | 1202 | Europe | Denmark | 4199.9 | 8.76 | 55.33 | 1104 | 0.0240 |
| DA222_noUDG.SG | 81 | Asia | Kazakhstan | 4411.1 | 76.98 | 43.20 | 1100 | 0.0234 |
| MS20.SG | 326 | Asia | Russia | 4229.0 | 61.55 | 55.85 | 1100 | 0.0209 |
| GRO021.A | 546 | Europe | Netherlands | 4095.9 | 6.57 | 53.22 | 1100 | 0.0192 |
| VK289_noUDG.SG | 1196 | Europe | Denmark | 4076.7 | 10.69 | 54.80 | 1100 | 0.0160 |
| VK291_noUDG.SG | 1196 | Europe | Denmark | 4076.7 | 10.69 | 54.80 | 1100 | 0.0263 |
| VK392_noUDG.SG | 1222 | Europe | Norway | 4615.1 | 7.92 | 59.58 | 1100 | 0.0190 |
| VK417_noUDG.SG | 1226 | Europe | Norway | 4641.8 | 10.57 | 60.36 | 1100 | 0.0240 |
| VK543_noUDG.SG | 1247 | Europe | Ireland | 5063.6 | -10.13 | 53.51 | 1100 | 0.0215 |
| VK544_noUDG.SG | 1248 | Europe | Ireland | 4778.9 | -6.30 | 53.39 | 1100 | 0.0234 |
| VK548_noUDG.SG | 1250 | Europe | Norway | 4905.8 | 11.22 | 63.44 | 1100 | 0.0264 |
| VK69_noUDG.SG | 1251 | Europe | Denmark | 4134.3 | 12.08 | 55.87 | 1100 | 0.0288 |
| VK70_noUDG.SG | 1251 | Europe | Denmark | 4134.3 | 12.08 | 55.87 | 1100 | 0.0224 |
| VK442_noUDG.SG | 1203 | Europe | Sweden | 4093.6 | 16.43 | 56.42 | 1097 | 0.0197 |

| | | | | | | | | |
|---|---|---|---|---|---|---|---|---|
| VK444_noUDG.SG | 1203 | Europe | Sweden | 4093.6 | 16.43 | 56.42 | 1097 | 0.0295 |
| VK355_noUDG.SG | 1214 | Europe | Sweden | 4093.6 | 16.41 | 56.28 | 1097 | 0.0245 |
| I15741 | 683 | Europe | Croatia | 2810.3 | 16.26 | 43.53 | 1096 | 0.0183 |
| R1283.SG | 1083 | Europe | Italy | 3097.9 | 12.47 | 41.90 | 1095 | 0.0257 |
| VK358_noUDG.SG | 1206 | Europe | Sweden | 4173.2 | 16.73 | 56.81 | 1092 | 0.0261 |
| VK336_noUDG.SG | 1206 | Europe | Sweden | 4173.2 | 16.73 | 56.81 | 1091 | 0.0182 |
| ARK-49.SG | 469 | Europe | Hungary | 2913.1 | 21.05 | 47.59 | 1090 | 0.0237 |
| C3619 | 58 | Asia | China | 5398.2 | 89.26 | 43.76 | 1089 | 0.0240 |
| POP001 | 1050 | Europe | Italy | 3205.3 | 11.02 | 42.91 | 1089 | 0.0167 |
| ETR007 | 535 | Europe | Italy | 3122.6 | 11.95 | 43.02 | 1088 | 0.0257 |
| VK332_noUDG.SG | 1203 | Europe | Sweden | 4093.6 | 16.43 | 56.42 | 1085 | 0.0250 |
| OLN001 | 347 | Asia | Mongolia | 6401.6 | 102.08 | 47.11 | 1075 | 0.0251 |
| OLN002 | 347 | Asia | Mongolia | 6401.6 | 102.08 | 47.11 | 1075 | 0.0200 |
| OLN004 | 347 | Asia | Mongolia | 6401.6 | 102.08 | 47.11 | 1075 | 0.0239 |
| OLN005 | 347 | Asia | Mongolia | 6401.6 | 102.08 | 47.11 | 1075 | 0.0187 |
| OLN007 | 347 | Asia | Mongolia | 6401.6 | 102.08 | 47.11 | 1075 | 0.0257 |
| OLN008 | 347 | Asia | Mongolia | 6401.6 | 102.08 | 47.11 | 1075 | 0.0255 |
| OLN009 | 347 | Asia | Mongolia | 6401.6 | 102.08 | 47.11 | 1075 | 0.0190 |
| OLN010 | 347 | Asia | Mongolia | 6401.6 | 102.08 | 47.11 | 1075 | 0.0189 |
| OLN011 | 347 | Asia | Mongolia | 6401.6 | 102.08 | 47.11 | 1075 | 0.0223 |
| OLN012 | 347 | Asia | Mongolia | 6401.6 | 102.08 | 47.11 | 1075 | 0.0157 |
| ZAA001 | 419 | Asia | Mongolia | 6542.4 | 104.05 | 47.85 | 1075 | 0.0233 |
| ZAA004 | 419 | Asia | Mongolia | 6542.4 | 104.05 | 47.85 | 1075 | 0.0205 |
| GRO013.A | 546 | Europe | Netherlands | 4095.9 | 6.57 | 53.22 | 1075 | 0.0211 |
| GRO014.A_noUDG | 546 | Europe | Netherlands | 4095.9 | 6.57 | 53.22 | 1075 | 0.0221 |
| VK294_noUDG.SG | 1199 | Europe | Denmark | 4116.6 | 11.25 | 55.56 | 1075 | 0.0240 |
| VK369_noUDG.SG | 1199 | Europe | Denmark | 4116.6 | 11.25 | 55.56 | 1075 | 0.0215 |
| VK384_noUDG.SG | 1215 | Europe | Denmark | 4197.3 | 10.20 | 55.95 | 1075 | 0.0231 |
| VK84_noUDG.SG | 1215 | Europe | Denmark | 4197.3 | 10.20 | 55.95 | 1075 | 0.0253 |
| VK87_noUDG.SG | 1215 | Europe | Denmark | 4197.3 | 10.20 | 55.95 | 1075 | 0.0196 |
| VK385_noUDG.SG | 1216 | Europe | Denmark | 4084.8 | 11.95 | 55.60 | 1075 | 0.0201 |

| | | | | | | | | |
|---|---|---|---|---|---|---|---|---|
| ARK-21.SG | 469 | Europe | Hungary | 2913.1 | 21.05 | 47.59 | 1069 | 0.0192 |
| VK337_noUDG.SG | 1207 | Europe | Sweden | 4093.6 | 16.58 | 56.48 | 1069 | 0.0218 |
| VK333_noUDG.SG | 1204 | Europe | Sweden | 4142.5 | 16.46 | 56.58 | 1052 | 0.0299 |
| DA243_noUDG.SG | 86 | Asia | Russia | 2207.8 | 42.59 | 43.96 | 1050 | 0.0191 |
| KLK002 | 299 | Asia | Kazakhstan | 4684.0 | 80.24 | 45.65 | 1050 | 0.0236 |
| DUN002.A_noUDG | 526 | Europe | Germany | 4088.8 | 7.65 | 53.60 | 1050 | 0.0279 |
| DUN007.A_noUDG | 526 | Europe | Germany | 4088.8 | 7.65 | 53.60 | 1050 | 0.0157 |
| DUN010.A_noUDG | 526 | Europe | Germany | 4088.8 | 7.65 | 53.60 | 1050 | 0.0202 |
| DUN011.A_noUDG | 526 | Europe | Germany | 4088.8 | 7.65 | 53.60 | 1050 | 0.0184 |
| GRO012.A | 546 | Europe | Netherlands | 4095.9 | 6.57 | 53.22 | 1050 | 0.0174 |
| GRO017.A_noUDG | 546 | Europe | Netherlands | 4095.9 | 6.57 | 53.22 | 1050 | 0.0164 |
| VK221_noUDG.SG | 1185 | Europe | Russia | 4101.4 | 32.30 | 60.00 | 1050 | 0.0228 |
| VK170_noUDG.SG | 1188 | Europe | Isle of Man | 4788.8 | -4.68 | 54.08 | 1050 | 0.0294 |
| VK415_noUDG.SG | 1225 | Europe | Norway | 4614.0 | 10.63 | 60.29 | 1050 | 0.0211 |
| VK420_noUDG.SG | 1228 | Europe | Norway | 4661.2 | 11.10 | 60.80 | 1050 | 0.0224 |
| VK448_noUDG.SG | 1233 | Europe | Norway | 4661.2 | 10.87 | 60.80 | 1050 | 0.0224 |
| R59.SG | 1112 | Europe | Italy | 3002.1 | 13.09 | 41.69 | 1045 | 0.0171 |
| R60.SG | 1112 | Europe | Italy | 3002.1 | 13.09 | 41.69 | 1045 | 0.0219 |
| I10895 | 600 | Europe | Spain | 3994.5 | 2.87 | 42.05 | 1038 | 0.0238 |
| VPB-167.SG | 1257 | Europe | Hungary | 3055.1 | 17.31 | 46.80 | 1038 | 0.0225 |
| I15743 | 683 | Europe | Croatia | 2810.3 | 16.26 | 43.53 | 1029 | 0.0214 |
| VK326_noUDG.SG | 1202 | Europe | Denmark | 4199.9 | 8.76 | 55.33 | 1026 | 0.0269 |
| K1-1.SG | 961 | Europe | Hungary | 2960.4 | 21.81 | 48.46 | 1025 | 0.0214 |
| K1-10.SG | 961 | Europe | Hungary | 2960.4 | 21.81 | 48.46 | 1025 | 0.0263 |
| K1-3286.SG | 961 | Europe | Hungary | 2960.4 | 21.81 | 48.46 | 1025 | 0.0278 |
| K2-16.SG | 961 | Europe | Hungary | 2960.4 | 21.81 | 48.46 | 1025 | 0.0248 |
| K2-52.SG | 961 | Europe | Hungary | 2960.4 | 21.81 | 48.46 | 1025 | 0.0266 |
| K2-61.SG | 961 | Europe | Hungary | 2960.4 | 21.81 | 48.46 | 1025 | 0.0266 |
| LB-1432.SG | 984 | Europe | Hungary | 2960.2 | 19.48 | 47.17 | 1025 | 0.0194 |
| HMSZ-43.SG | 555 | Europe | Hungary | 2915.5 | 19.13 | 46.54 | 1022 | 0.0217 |
| HMSZ-50.SG | 555 | Europe | Hungary | 2915.5 | 19.13 | 46.54 | 1021 | 0.0212 |

| | | | | | | | | |
|---|---|---|---|---|---|---|---|---|
| Anapa-11.SG | 464 | Europe | Russia | 2444.0 | 37.50 | 44.97 | 1020 | 0.0255 |
| DA204_noUDG.SG | 81 | Asia | Kazakhstan | 4411.1 | 76.98 | 43.20 | 1017 | 0.0229 |
| COR001 | 508 | Europe | Italy | 3570.9 | 8.79 | 40.52 | 1017 | 0.0240 |
| Anapa-10.SG | 464 | Europe | Russia | 2444.0 | 37.50 | 44.97 | 1015 | 0.0254 |
| SE-16.SG | 1136 | Europe | Hungary | 2868.4 | 20.13 | 46.49 | 1015 | 0.0188 |
| I13839 | 654 | Europe | Albania | 2367.5 | 20.71 | 40.43 | 1014 | 0.0225 |
| I2525 | 801 | Europe | Bulgaria | 2257.6 | 25.61 | 43.14 | 1014 | 0.0214 |
| I2868 | 218 | Asia | India | 4788.1 | 79.74 | 30.25 | 1011 | 0.0175 |
| SZA-154.SG | 1149 | Europe | Hungary | 2868.4 | 20.32 | 46.72 | 1011 | 0.0267 |
| VK143_noUDG.SG | 1184 | Europe | United Kingdom | 4397.4 | -1.26 | 51.76 | 1010 | 0.0280 |
| VK145_noUDG.SG | 1184 | Europe | United Kingdom | 4397.4 | -1.26 | 51.76 | 1010 | 0.0260 |
| VK146_noUDG.SG | 1184 | Europe | United Kingdom | 4397.4 | -1.26 | 51.76 | 1010 | 0.0153 |
| VK147_noUDG.SG | 1184 | Europe | United Kingdom | 4397.4 | -1.26 | 51.76 | 1010 | 0.0168 |
| VK150_noUDG.SG | 1184 | Europe | United Kingdom | 4397.4 | -1.26 | 51.76 | 1010 | 0.0193 |
| VK151_noUDG.SG | 1184 | Europe | United Kingdom | 4397.4 | -1.26 | 51.76 | 1010 | 0.0222 |
| VK165_noUDG.SG | 1184 | Europe | United Kingdom | 4397.4 | -1.26 | 51.76 | 1010 | 0.0231 |
| VK166_noUDG.SG | 1184 | Europe | United Kingdom | 4397.4 | -1.26 | 51.76 | 1010 | 0.0212 |
| VK167_noUDG.SG | 1184 | Europe | United Kingdom | 4397.4 | -1.26 | 51.76 | 1010 | 0.0264 |
| VK168_noUDG.SG | 1184 | Europe | United Kingdom | 4397.4 | -1.26 | 51.76 | 1010 | 0.0229 |
| VK172_noUDG.SG | 1184 | Europe | United Kingdom | 4397.4 | -1.26 | 51.76 | 1010 | 0.0246 |
| VK173_noUDG.SG | 1184 | Europe | United Kingdom | 4397.4 | -1.26 | 51.76 | 1010 | 0.0226 |
| VK174_noUDG.SG | 1184 | Europe | United Kingdom | 4397.4 | -1.26 | 51.76 | 1010 | 0.0198 |
| VK175_noUDG.SG | 1184 | Europe | United Kingdom | 4397.4 | -1.26 | 51.76 | 1010 | 0.0208 |
| VK176_noUDG.SG | 1184 | Europe | United Kingdom | 4397.4 | -1.26 | 51.76 | 1010 | 0.0191 |
| VK177_noUDG.SG | 1184 | Europe | United Kingdom | 4397.4 | -1.26 | 51.76 | 1010 | 0.0233 |
| VK178_noUDG.SG | 1184 | Europe | United Kingdom | 4397.4 | -1.26 | 51.76 | 1010 | 0.0200 |
| AGY-92.SG | 454 | Europe | Hungary | 2868.4 | 20.35 | 46.48 | 1009 | 0.0216 |
| NTH-2.SG | 1023 | Europe | Hungary | 3004.9 | 19.43 | 47.61 | 1008 | 0.0255 |
| VPB-310.SG | 1257 | Europe | Hungary | 3055.1 | 17.31 | 46.80 | 1008 | 0.0222 |
| PLE-200.SG | 1042 | Europe | Hungary | 2913.1 | 21.12 | 47.37 | 1007 | 0.0231 |
| AGY-49.SG | 454 | Europe | Hungary | 2868.4 | 20.35 | 46.48 | 1005 | 0.0268 |

| | | | | | | | | |
|---|---|---|---|---|---|---|---|---|
| PLE-38.SG | 1042 | Europe | Hungary | 2913.1 | 21.12 | 47.37 | 1003 | 0.0254 |
| AGY-75.SG | 454 | Europe | Hungary | 2868.4 | 20.35 | 46.48 | 1000 | 0.0173 |
| AGY-87.SG | 454 | Europe | Hungary | 2868.4 | 20.35 | 46.48 | 1000 | 0.0214 |
| CSU-11.SG | 514 | Europe | Hungary | 2868.4 | 20.18 | 46.37 | 1000 | 0.0150 |
| DA142_noUDG.SG | 517 | Europe | Russia | 2641.2 | 39.54 | 47.28 | 1000 | 0.0231 |
| ETR010 | 535 | Europe | Italy | 3122.6 | 11.95 | 43.02 | 1000 | 0.0130 |
| GRO002.A_noUDG | 546 | Europe | Netherlands | 4095.9 | 6.57 | 53.22 | 1000 | 0.0153 |
| IBE-154.SG | 952 | Europe | Hungary | 2960.4 | 21.81 | 48.22 | 1000 | 0.0273 |
| IBE-161.SG | 952 | Europe | Hungary | 2960.4 | 21.81 | 48.22 | 1000 | 0.0217 |
| IBE-206.SG | 952 | Europe | Hungary | 2960.4 | 21.81 | 48.22 | 1000 | 0.0252 |
| MH1-14.SG | 999 | Europe | Hungary | 2867.4 | 21.68 | 47.07 | 1000 | 0.0255 |
| MH1-4.SG | 999 | Europe | Hungary | 2867.4 | 21.68 | 47.07 | 1000 | 0.0257 |
| MH1-9.SG | 999 | Europe | Hungary | 2867.4 | 21.68 | 47.07 | 1000 | 0.0179 |
| NK-2.SG | 1022 | Europe | Hungary | 2913.5 | 19.93 | 47.07 | 1000 | 0.0278 |
| R1291.SG | 1083 | Europe | Italy | 3097.9 | 12.47 | 41.90 | 1000 | 0.0190 |
| R1294.SG | 1083 | Europe | Italy | 3097.9 | 12.47 | 41.90 | 1000 | 0.0148 |
| SEO-3.SG | 1139 | Europe | Hungary | 2868.4 | 20.20 | 46.40 | 1000 | 0.0198 |
| SEO-4.SG | 1139 | Europe | Hungary | 2868.4 | 20.20 | 46.40 | 1000 | 0.0319 |
| SH-103.SG | 1140 | Europe | Hungary | 2913.1 | 21.21 | 47.31 | 1000 | 0.0247 |
| SH-106.SG | 1140 | Europe | Hungary | 2913.1 | 21.21 | 47.31 | 1000 | 0.0241 |
| SH-12.SG | 1140 | Europe | Hungary | 2913.1 | 21.21 | 47.31 | 1000 | 0.0237 |
| SH-143.SG | 1140 | Europe | Hungary | 2913.1 | 21.21 | 47.31 | 1000 | 0.0208 |
| SH-175.SG | 1140 | Europe | Hungary | 2913.1 | 21.21 | 47.31 | 1000 | 0.0256 |
| SH-182.SG | 1140 | Europe | Hungary | 2913.1 | 21.21 | 47.31 | 1000 | 0.0207 |
| SH-98.SG | 1140 | Europe | Hungary | 2913.1 | 21.21 | 47.31 | 1000 | 0.0201 |
| SO-5.SG | 1140 | Europe | Hungary | 2913.1 | 21.21 | 47.31 | 1000 | 0.0200 |
| SZA-20.SG | 1149 | Europe | Hungary | 2868.4 | 20.32 | 46.72 | 1000 | 0.0194 |
| SZA-29.SG | 1149 | Europe | Hungary | 2868.4 | 20.32 | 46.72 | 1000 | 0.0191 |
| SZA-44.SG | 1149 | Europe | Hungary | 2868.4 | 20.32 | 46.72 | 1000 | 0.0208 |
| SZA-52.SG | 1149 | Europe | Hungary | 2868.4 | 20.32 | 46.72 | 1000 | 0.0251 |
| SZA-7.SG | 1149 | Europe | Hungary | 2868.4 | 20.32 | 46.72 | 1000 | 0.0267 |

| | | | | | | | | |
|---|---|---|---|---|---|---|---|---|
| SZOD-394.SG | 1149 | Europe | Hungary | 2868.4 | 20.32 | 46.72 | 1000 | 0.0234 |
| SZOD-566.SG | 1149 | Europe | Hungary | 2868.4 | 20.32 | 46.72 | 1000 | 0.0187 |
| SZAK-4.SG | 1150 | Europe | Hungary | 3098.7 | 16.81 | 47.45 | 1000 | 0.0218 |
| SZAK-6.SG | 1150 | Europe | Hungary | 3098.7 | 16.81 | 47.45 | 1000 | 0.0230 |
| SZAK-7.SG | 1150 | Europe | Hungary | 3098.7 | 16.81 | 47.45 | 1000 | 0.0164 |
| VK141_noUDG.SG | 1183 | Europe | Denmark | 4158.0 | 10.33 | 55.50 | 1000 | 0.0254 |
| VK279_noUDG.SG | 1183 | Europe | Denmark | 4158.0 | 10.33 | 55.50 | 1000 | 0.0218 |
| VK370_noUDG.SG | 1183 | Europe | Denmark | 4158.0 | 10.33 | 55.50 | 1000 | 0.0200 |
| VK372_noUDG.SG | 1183 | Europe | Denmark | 4158.0 | 10.33 | 55.50 | 1000 | 0.0205 |
| VK373_noUDG.SG | 1183 | Europe | Denmark | 4158.0 | 10.33 | 55.50 | 1000 | 0.0306 |
| VK446_noUDG.SG | 1183 | Europe | Denmark | 4158.0 | 10.33 | 55.50 | 1000 | 0.0192 |
| VK265_noUDG.SG | 1190 | Europe | Sweden | 4191.5 | 13.93 | 57.18 | 1000 | 0.0173 |
| VK274_noUDG.SG | 1192 | Europe | Denmark | 4076.7 | 10.78 | 54.85 | 1000 | 0.0222 |
| VK317_noUDG.SG | 1192 | Europe | Denmark | 4076.7 | 10.78 | 54.85 | 1000 | 0.0212 |
| VK281_noUDG.SG | 1193 | Europe | Denmark | 4035.3 | 11.95 | 55.13 | 1000 | 0.0269 |
| VK284_noUDG.SG | 1194 | Europe | Denmark | 4084.8 | 12.14 | 55.65 | 1000 | 0.0268 |
| VK286_noUDG.SG | 1195 | Europe | Denmark | 4076.7 | 10.72 | 54.87 | 1000 | 0.0267 |
| VK288_noUDG.SG | 1195 | Europe | Denmark | 4076.7 | 10.72 | 54.87 | 1000 | 0.0211 |
| VK320_noUDG.SG | 1195 | Europe | Denmark | 4076.7 | 10.72 | 54.87 | 1000 | 0.0223 |
| VK361_noUDG.SG | 1195 | Europe | Denmark | 4076.7 | 10.72 | 54.87 | 1000 | 0.0239 |
| VK363_noUDG.SG | 1195 | Europe | Denmark | 4076.7 | 10.72 | 54.87 | 1000 | 0.0279 |
| VK364_noUDG.SG | 1195 | Europe | Denmark | 4076.7 | 10.72 | 54.87 | 1000 | 0.0201 |
| VK365_noUDG.SG | 1195 | Europe | Denmark | 4076.7 | 10.72 | 54.87 | 1000 | 0.0200 |
| VK367_noUDG.SG | 1195 | Europe | Denmark | 4076.7 | 10.72 | 54.87 | 1000 | 0.0196 |
| VK368_noUDG.SG | 1195 | Europe | Denmark | 4076.7 | 10.72 | 54.87 | 1000 | 0.0258 |
| VK290_noUDG.SG | 1198 | Europe | Denmark | 4076.7 | 10.74 | 54.89 | 1000 | 0.0217 |
| VK316_noUDG.SG | 1201 | Europe | Denmark | 4158.0 | 10.45 | 55.51 | 1000 | 0.0181 |
| VK323_noUDG.SG | 1202 | Europe | Denmark | 4199.9 | 8.76 | 55.33 | 1000 | 0.0226 |
| VK335_noUDG.SG | 1205 | Europe | Sweden | 4173.2 | 16.72 | 56.73 | 1000 | 0.0271 |
| VK345_noUDG.SG | 1205 | Europe | Sweden | 4173.2 | 16.72 | 56.73 | 1000 | 0.0234 |
| VK346_noUDG.SG | 1205 | Europe | Sweden | 4173.2 | 16.72 | 56.73 | 1000 | 0.0282 |

| | | | | | | | | |
|---|---|---|---|---|---|---|---|---|
| VK533_noUDG.SG | 1206 | Europe | Sweden | 4173.2 | 16.73 | 56.81 | 1000 | 0.0239 |
| VK342_noUDG.SG | 1208 | Europe | Sweden | 4093.6 | 16.57 | 56.44 | 1000 | 0.0229 |
| VK343_noUDG.SG | 1209 | Europe | Sweden | 4093.6 | 16.62 | 56.55 | 1000 | 0.0259 |
| VK344_noUDG.SG | 1210 | Europe | Sweden | 4093.6 | 16.41 | 56.33 | 1000 | 0.0222 |
| VK348_noUDG.SG | 1211 | Europe | Sweden | 4173.2 | 16.72 | 56.88 | 1000 | 0.0237 |
| VK352_noUDG.SG | 1213 | Europe | Sweden | 4173.2 | 16.71 | 56.72 | 1000 | 0.0227 |
| VK386_noUDG.SG | 1217 | Europe | Norway | 4854.6 | 9.05 | 62.09 | 1000 | 0.0211 |
| VK389_noUDG.SG | 1220 | Europe | Norway | 4501.9 | 9.61 | 59.22 | 1000 | 0.0207 |
| VK393_noUDG.SG | 1223 | Europe | Norway | 4661.2 | 11.00 | 60.84 | 1000 | 0.0230 |
| VK422_noUDG.SG | 1229 | Europe | Norway | 4708.5 | 11.34 | 61.15 | 1000 | 0.0276 |
| VK443_noUDG.SG | 1231 | Europe | Sweden | 4221.6 | 17.02 | 57.34 | 1000 | 0.0174 |
| VK445_noUDG.SG | 1232 | Europe | Denmark | 4084.8 | 11.97 | 55.62 | 1000 | 0.0195 |
| VPB-561.SG | 1257 | Europe | Hungary | 3055.1 | 17.31 | 46.80 | 1000 | 0.0145 |
| TAQ011 | 1157 | Europe | Italy | 3097.9 | 11.77 | 42.25 | 991 | 0.0263 |
| ETR013 | 535 | Europe | Italy | 3122.6 | 11.95 | 43.02 | 982 | 0.0199 |
| VK327_noUDG.SG | 1202 | Europe | Denmark | 4199.9 | 8.76 | 55.33 | 979 | 0.0211 |
| BK-2.SG | 484 | Europe | Hungary | 3004.9 | 19.31 | 47.36 | 975 | 0.0227 |
| KH-500.SG | 963 | Europe | Hungary | 2868.4 | 20.09 | 46.38 | 975 | 0.0252 |
| KH-596.SG | 963 | Europe | Hungary | 2868.4 | 20.09 | 46.38 | 975 | 0.0228 |
| NTH-1.SG | 1023 | Europe | Hungary | 3004.9 | 19.43 | 47.61 | 975 | 0.0194 |
| NTH-19.SG | 1023 | Europe | Hungary | 3004.9 | 19.43 | 47.61 | 975 | 0.0219 |
| NTH-20.SG | 1023 | Europe | Hungary | 3004.9 | 19.43 | 47.61 | 975 | 0.0212 |
| PLE-216.SG | 1042 | Europe | Hungary | 2913.1 | 21.12 | 47.37 | 975 | 0.0163 |
| PLE-23.SG | 1042 | Europe | Hungary | 2913.1 | 21.12 | 47.37 | 975 | 0.0222 |
| PLE-28.SG | 1042 | Europe | Hungary | 2913.1 | 21.12 | 47.37 | 975 | 0.0269 |
| PLE-57.SG | 1042 | Europe | Hungary | 2913.1 | 21.12 | 47.37 | 975 | 0.0254 |
| SH-251.SG | 1140 | Europe | Hungary | 2913.1 | 21.21 | 47.31 | 975 | 0.0119 |
| SH-41.SG | 1140 | Europe | Hungary | 2913.1 | 21.21 | 47.31 | 975 | 0.0270 |
| SH-81.SG | 1140 | Europe | Hungary | 2913.1 | 21.21 | 47.31 | 975 | 0.0237 |
| VK429_noUDG.SG | 1230 | Europe | Sweden | 4252.3 | 18.19 | 57.34 | 975 | 0.0240 |
| VK433_noUDG.SG | 1230 | Europe | Sweden | 4252.3 | 18.19 | 57.34 | 975 | 0.0247 |

| | | | | | | | | |
|---|---|---|---|---|---|---|---|---|
| VK455_noUDG.SG | 1230 | Europe | Sweden | 4252.3 | 18.19 | 57.34 | 975 | 0.0206 |
| VK456_noUDG.SG | 1230 | Europe | Sweden | 4252.3 | 18.19 | 57.34 | 975 | 0.0261 |
| VK56_noUDG.SG | 1230 | Europe | Sweden | 4252.3 | 18.19 | 57.34 | 975 | 0.0232 |
| VK58_noUDG.SG | 1230 | Europe | Sweden | 4252.3 | 18.19 | 57.34 | 975 | 0.0263 |
| VK60_noUDG.SG | 1230 | Europe | Sweden | 4252.3 | 18.19 | 57.34 | 975 | 0.0279 |
| VK64_noUDG.SG | 1230 | Europe | Sweden | 4252.3 | 18.19 | 57.34 | 975 | 0.0310 |
| VK468_noUDG.SG | 1234 | Europe | Sweden | 4300.0 | 18.28 | 57.63 | 975 | 0.0200 |
| VK473_noUDG.SG | 1234 | Europe | Sweden | 4300.0 | 18.28 | 57.63 | 975 | 0.0240 |
| VK474_noUDG.SG | 1234 | Europe | Sweden | 4300.0 | 18.28 | 57.63 | 975 | 0.0231 |
| VK475_noUDG.SG | 1234 | Europe | Sweden | 4300.0 | 18.28 | 57.63 | 975 | 0.0243 |
| VK477_noUDG.SG | 1234 | Europe | Sweden | 4300.0 | 18.28 | 57.63 | 975 | 0.0192 |
| VK478_noUDG.SG | 1234 | Europe | Sweden | 4300.0 | 18.28 | 57.63 | 975 | 0.0229 |
| VK479_noUDG.SG | 1234 | Europe | Sweden | 4300.0 | 18.28 | 57.63 | 975 | 0.0201 |
| VK50_noUDG.SG | 1234 | Europe | Sweden | 4300.0 | 18.28 | 57.63 | 975 | 0.0215 |
| TAQ009 | 1157 | Europe | Italy | 3097.9 | 11.77 | 42.25 | 972 | 0.0227 |
| K2-18.SG | 961 | Europe | Hungary | 2960.4 | 21.81 | 48.46 | 970 | 0.0278 |
| K2-29.SG | 961 | Europe | Hungary | 2960.4 | 21.81 | 48.46 | 970 | 0.0216 |
| K2-33.SG | 961 | Europe | Hungary | 2960.4 | 21.81 | 48.46 | 970 | 0.0270 |
| K3-12.SG | 961 | Europe | Hungary | 2960.4 | 21.81 | 48.46 | 970 | 0.0253 |
| K3-13.SG | 961 | Europe | Hungary | 2960.4 | 21.81 | 48.46 | 970 | 0.0266 |
| KeF1-10936.SG | 964 | Europe | Hungary | 3004.4 | 21.66 | 48.20 | 970 | 0.0219 |
| SE-114.SG | 1136 | Europe | Hungary | 2868.4 | 20.13 | 46.49 | 970 | 0.0184 |
| SE-23.SG | 1136 | Europe | Hungary | 2868.4 | 20.13 | 46.49 | 970 | 0.0185 |
| SE-64.SG | 1136 | Europe | Hungary | 2868.4 | 20.13 | 46.49 | 970 | 0.0230 |
| SP-10.SG | 1140 | Europe | Hungary | 2913.1 | 21.21 | 47.31 | 970 | 0.0222 |
| SP-2.SG | 1140 | Europe | Hungary | 2913.1 | 21.21 | 47.31 | 970 | 0.0321 |
| SP-9.SG | 1140 | Europe | Hungary | 2913.1 | 21.21 | 47.31 | 970 | 0.0251 |
| TCS-5.SG | 1158 | Europe | Hungary | 3003.9 | 20.57 | 47.67 | 970 | 0.0236 |
| TCS-18.SG | 1158 | Europe | Hungary | 3003.9 | 20.57 | 47.67 | 963 | 0.0255 |
| R9919.SG | 1099 | Europe | Montenegro | 2535.7 | 19.27 | 42.47 | 960 | 0.0279 |
| R63.SG | 1112 | Europe | Italy | 3002.1 | 13.09 | 41.69 | 960 | 0.0282 |

| | | | | | | | | |
|---|---|---|---|---|---|---|---|---|
| VK256_noUDG.SG | 1189 | Europe | United Kingdom | 4409.4 | -2.47 | 50.67 | 953 | 0.0259 |
| VK257_noUDG.SG | 1189 | Europe | United Kingdom | 4409.4 | -2.47 | 50.67 | 953 | 0.0209 |
| VK258_noUDG.SG | 1189 | Europe | United Kingdom | 4409.4 | -2.47 | 50.67 | 953 | 0.0230 |
| VK259_noUDG.SG | 1189 | Europe | United Kingdom | 4409.4 | -2.47 | 50.67 | 953 | 0.0280 |
| VK260_noUDG.SG | 1189 | Europe | United Kingdom | 4409.4 | -2.47 | 50.67 | 953 | 0.0228 |
| VK261_noUDG.SG | 1189 | Europe | United Kingdom | 4409.4 | -2.47 | 50.67 | 953 | 0.0164 |
| VK262_noUDG.SG | 1189 | Europe | United Kingdom | 4409.4 | -2.47 | 50.67 | 953 | 0.0259 |
| VK263_noUDG.SG | 1189 | Europe | United Kingdom | 4409.4 | -2.47 | 50.67 | 953 | 0.0241 |
| VK264_noUDG.SG | 1189 | Europe | United Kingdom | 4409.4 | -2.47 | 50.67 | 953 | 0.0247 |
| VK449_noUDG.SG | 1189 | Europe | United Kingdom | 4409.4 | -2.47 | 50.67 | 953 | 0.0242 |
| GRO007.A | 546 | Europe | Netherlands | 4095.9 | 6.57 | 53.22 | 950 | 0.0203 |
| GRO010.A | 546 | Europe | Netherlands | 4095.9 | 6.57 | 53.22 | 950 | 0.0226 |
| GRO015.A_noUDG | 546 | Europe | Netherlands | 4095.9 | 6.57 | 53.22 | 950 | 0.0302 |
| GRO016.A_noUDG | 546 | Europe | Netherlands | 4095.9 | 6.57 | 53.22 | 950 | 0.0188 |
| HMSZ-157.SG | 555 | Europe | Hungary | 2915.5 | 19.13 | 46.54 | 950 | 0.0223 |
| HMSZ-229.SG | 555 | Europe | Hungary | 2915.5 | 19.13 | 46.54 | 950 | 0.0255 |
| HMSZ-231.SG | 555 | Europe | Hungary | 2915.5 | 19.13 | 46.54 | 950 | 0.0235 |
| HMSZ-245.SG | 555 | Europe | Hungary | 2915.5 | 19.13 | 46.54 | 950 | 0.0224 |
| HMSZ-86.SG | 555 | Europe | Hungary | 2915.5 | 19.13 | 46.54 | 950 | 0.0195 |
| HMSZ-88.SG | 555 | Europe | Hungary | 2915.5 | 19.13 | 46.54 | 950 | 0.0267 |
| I10892 | 600 | Europe | Spain | 3994.5 | 2.87 | 42.05 | 950 | 0.0234 |
| KIL001.B_noUDG | 967 | Europe | Ireland | 4945.5 | -8.20 | 54.01 | 950 | 0.0199 |
| KIL003.B_noUDG | 967 | Europe | Ireland | 4945.5 | -8.20 | 54.01 | 950 | 0.0210 |
| KIL004.B_noUDG | 967 | Europe | Ireland | 4945.5 | -8.20 | 54.01 | 950 | 0.0148 |
| KIL006.B_noUDG | 967 | Europe | Ireland | 4945.5 | -8.20 | 54.01 | 950 | 0.0265 |
| KIL009.B_noUDG | 967 | Europe | Ireland | 4945.5 | -8.20 | 54.01 | 950 | 0.0223 |
| KIL012.A_noUDG | 967 | Europe | Ireland | 4945.5 | -8.20 | 54.01 | 950 | 0.0252 |
| KIL013.B_noUDG | 967 | Europe | Ireland | 4945.5 | -8.20 | 54.01 | 950 | 0.0216 |
| KIL014.B_noUDG | 967 | Europe | Ireland | 4945.5 | -8.20 | 54.01 | 950 | 0.0190 |
| KIL016.B_noUDG | 967 | Europe | Ireland | 4945.5 | -8.20 | 54.01 | 950 | 0.0270 |
| KIL022.B_noUDG | 967 | Europe | Ireland | 4945.5 | -8.20 | 54.01 | 950 | 0.0195 |

| | | | | | | | | |
|---|---|---|---|---|---|---|---|---|
| KIL023.B_noUDG | 967 | Europe | Ireland | 4945.5 | -8.20 | 54.01 | 950 | 0.0240 |
| KIL025.B_noUDG | 967 | Europe | Ireland | 4945.5 | -8.20 | 54.01 | 950 | 0.0215 |
| KIL026.B_noUDG | 967 | Europe | Ireland | 4945.5 | -8.20 | 54.01 | 950 | 0.0198 |
| KIL027.B_noUDG | 967 | Europe | Ireland | 4945.5 | -8.20 | 54.01 | 950 | 0.0163 |
| KIL028.B_noUDG | 967 | Europe | Ireland | 4945.5 | -8.20 | 54.01 | 950 | 0.0199 |
| KIL032.B_noUDG | 967 | Europe | Ireland | 4945.5 | -8.20 | 54.01 | 950 | 0.0138 |
| KIL033.B_noUDG | 967 | Europe | Ireland | 4945.5 | -8.20 | 54.01 | 950 | 0.0181 |
| KIL034.A_noUDG | 967 | Europe | Ireland | 4945.5 | -8.20 | 54.01 | 950 | 0.0208 |
| KIL035.B_noUDG | 967 | Europe | Ireland | 4945.5 | -8.20 | 54.01 | 950 | 0.0247 |
| KIL037.A_noUDG | 967 | Europe | Ireland | 4945.5 | -8.20 | 54.01 | 950 | 0.0222 |
| KIL041.B_noUDG | 967 | Europe | Ireland | 4945.5 | -8.20 | 54.01 | 950 | 0.0190 |
| KIL042.A_noUDG | 967 | Europe | Ireland | 4945.5 | -8.20 | 54.01 | 950 | 0.0201 |
| KIL043.B_noUDG | 967 | Europe | Ireland | 4945.5 | -8.20 | 54.01 | 950 | 0.0174 |
| KIL044.B_noUDG | 967 | Europe | Ireland | 4945.5 | -8.20 | 54.01 | 950 | 0.0225 |
| KIL047.B_noUDG | 967 | Europe | Ireland | 4945.5 | -8.20 | 54.01 | 950 | 0.0207 |
| VK108_noUDG.SG | 1181 | Europe | Sweden | 4060.0 | 13.05 | 55.54 | 950 | 0.0170 |
| VK219_noUDG.SG | 1185 | Europe | Russia | 4101.4 | 32.30 | 60.00 | 950 | 0.0210 |
| VK154_noUDG.SG | 1186 | Europe | Poland | 3539.2 | 18.90 | 52.70 | 950 | 0.0223 |
| VK156_noUDG.SG | 1186 | Europe | Poland | 3539.2 | 18.90 | 52.70 | 950 | 0.0192 |
| VK157_noUDG.SG | 1186 | Europe | Poland | 3539.2 | 18.90 | 52.70 | 950 | 0.0291 |
| VK273_noUDG.SG | 1191 | Europe | Russia | 3548.0 | 31.87 | 54.78 | 950 | 0.0238 |
| VK387_noUDG.SG | 1218 | Europe | Norway | 4641.8 | 10.43 | 60.31 | 950 | 0.0249 |
| VK414_noUDG.SG | 1224 | Europe | Norway | 4835.7 | 8.40 | 61.88 | 950 | 0.0227 |
| VPB-118.SG | 1257 | Europe | Hungary | 3055.1 | 17.31 | 46.80 | 950 | 0.0227 |
| VPB-588.SG | 1257 | Europe | Hungary | 3055.1 | 17.31 | 46.80 | 950 | 0.0276 |
| VPB-600.SG | 1257 | Europe | Hungary | 3055.1 | 17.31 | 46.80 | 950 | 0.0248 |
| IBE-176.SG | 952 | Europe | Hungary | 2960.4 | 21.81 | 48.22 | 944 | 0.0288 |
| I10495 | 596 | Europe | Romania | 2402.8 | 26.01 | 44.90 | 943 | 0.0198 |
| I14791 | 670 | Europe | Turkey | 1180.9 | 37.17 | 36.69 | 942 | 0.0194 |
| R1285.SG | 1083 | Europe | Italy | 3097.9 | 12.47 | 41.90 | 935 | 0.0172 |
| VK324_noUDG.SG | 1202 | Europe | Denmark | 4199.9 | 8.76 | 55.33 | 935 | 0.0231 |

| | | | | | | | | |
|---|---|---|---|---|---|---|---|---|
| VK322_noUDG.SG | 1202 | Europe | Denmark | 4199.9 | 8.76 | 55.33 | 934 | 0.0261 |
| GRO008.A | 546 | Europe | Netherlands | 4095.9 | 6.57 | 53.22 | 925 | 0.0181 |
| PLE-115.SG | 1042 | Europe | Hungary | 2913.1 | 21.12 | 47.37 | 925 | 0.0229 |
| PLE-95.SG | 1042 | Europe | Hungary | 2913.1 | 21.12 | 47.37 | 925 | 0.0198 |
| I2530 | 802 | Europe | North Macedonia | 2328.0 | 21.33 | 41.03 | 915 | 0.0175 |
| R3918.SG | 1108 | Europe | Serbia | 2743.4 | 19.61 | 44.97 | 904 | 0.0242 |
| GRO001.A_noUDG | 546 | Europe | Netherlands | 4095.9 | 6.57 | 53.22 | 900 | 0.0301 |
| GRO005.A_noUDG | 546 | Europe | Netherlands | 4095.9 | 6.57 | 53.22 | 900 | 0.0201 |
| GRO024.A | 546 | Europe | Netherlands | 4095.9 | 6.57 | 53.22 | 900 | 0.0251 |
| I7498 | 935 | Europe | Spain | 4728.9 | -2.54 | 37.81 | 900 | 0.0213 |
| I7499 | 935 | Europe | Spain | 4728.9 | -2.54 | 37.81 | 900 | 0.0184 |
| IBE-106.SG | 952 | Europe | Hungary | 2960.4 | 21.81 | 48.22 | 900 | 0.0315 |
| IBE-107.SG | 952 | Europe | Hungary | 2960.4 | 21.81 | 48.22 | 900 | 0.0231 |
| IBE-116.SG | 952 | Europe | Hungary | 2960.4 | 21.81 | 48.22 | 900 | 0.0172 |
| IBE-90.SG | 952 | Europe | Hungary | 2960.4 | 21.81 | 48.22 | 900 | 0.0214 |
| KPN001.A_noUDG | 980 | Europe | Denmark | 4134.3 | 12.57 | 55.68 | 900 | 0.0255 |
| KPN002.A_noUDG | 980 | Europe | Denmark | 4134.3 | 12.57 | 55.68 | 900 | 0.0165 |
| KPN003.A_noUDG | 980 | Europe | Denmark | 4134.3 | 12.57 | 55.68 | 900 | 0.0164 |
| KPN004.A_noUDG | 980 | Europe | Denmark | 4134.3 | 12.57 | 55.68 | 900 | 0.0123 |
| KPN005.A_noUDG | 980 | Europe | Denmark | 4134.3 | 12.57 | 55.68 | 900 | 0.0126 |
| KPN006.A_noUDG | 980 | Europe | Denmark | 4134.3 | 12.57 | 55.68 | 900 | 0.0212 |
| KPN007.A_noUDG | 980 | Europe | Denmark | 4134.3 | 12.57 | 55.68 | 900 | 0.0174 |
| KPN008.A_noUDG | 980 | Europe | Denmark | 4134.3 | 12.57 | 55.68 | 900 | 0.0201 |
| KPN011.A_noUDG | 980 | Europe | Denmark | 4134.3 | 12.57 | 55.68 | 900 | 0.0216 |
| KPN012.A_noUDG | 980 | Europe | Denmark | 4134.3 | 12.57 | 55.68 | 900 | 0.0205 |
| KPN013.A_noUDG | 980 | Europe | Denmark | 4134.3 | 12.57 | 55.68 | 900 | 0.0238 |
| KPN014.A_noUDG | 980 | Europe | Denmark | 4134.3 | 12.57 | 55.68 | 900 | 0.0203 |
| MH-106.SG | 999 | Europe | Hungary | 2867.4 | 21.68 | 47.07 | 900 | 0.0145 |
| MH-107.SG | 999 | Europe | Hungary | 2867.4 | 21.68 | 47.07 | 900 | 0.0293 |
| MH-153.SG | 999 | Europe | Hungary | 2867.4 | 21.68 | 47.07 | 900 | 0.0193 |
| MH-88.SG | 999 | Europe | Hungary | 2867.4 | 21.68 | 47.07 | 900 | 0.0229 |

| | | | | | | | | |
|---|---|---|---|---|---|---|---|---|
| PLE-327.SG | 1042 | Europe | Hungary | 2913.1 | 21.12 | 47.37 | 900 | 0.0169 |
| PLE-441.SG | 1042 | Europe | Hungary | 2913.1 | 21.12 | 47.37 | 900 | 0.0161 |
| SZOD-426.SG | 1149 | Europe | Hungary | 2868.4 | 20.32 | 46.72 | 900 | 0.0161 |
| VK15_noUDG.SG | 1185 | Europe | Russia | 4101.4 | 32.30 | 60.00 | 900 | 0.0171 |
| VK18_noUDG.SG | 1185 | Europe | Russia | 4101.4 | 32.30 | 60.00 | 900 | 0.0254 |
| VK220_noUDG.SG | 1185 | Europe | Russia | 4101.4 | 32.30 | 60.00 | 900 | 0.0222 |
| VK29_noUDG.SG | 1197 | Europe | Sweden | 4366.6 | 13.65 | 58.38 | 900 | 0.0241 |
| VK303_noUDG.SG | 1197 | Europe | Sweden | 4366.6 | 13.65 | 58.38 | 900 | 0.0219 |
| VK306_noUDG.SG | 1197 | Europe | Sweden | 4366.6 | 13.65 | 58.38 | 900 | 0.0177 |
| VK308_noUDG.SG | 1197 | Europe | Sweden | 4366.6 | 13.65 | 58.38 | 900 | 0.0249 |
| VK33_noUDG.SG | 1197 | Europe | Sweden | 4366.6 | 13.65 | 58.38 | 900 | 0.0248 |
| VK34_noUDG.SG | 1197 | Europe | Sweden | 4366.6 | 13.65 | 58.38 | 900 | 0.0231 |
| VK35_noUDG.SG | 1197 | Europe | Sweden | 4366.6 | 13.65 | 58.38 | 900 | 0.0246 |
| VK395_noUDG.SG | 1197 | Europe | Sweden | 4366.6 | 13.65 | 58.38 | 900 | 0.0153 |
| VK396_noUDG.SG | 1197 | Europe | Sweden | 4366.6 | 13.65 | 58.38 | 900 | 0.0256 |
| VK397_noUDG.SG | 1197 | Europe | Sweden | 4366.6 | 13.65 | 58.38 | 900 | 0.0270 |
| VK398_noUDG.SG | 1197 | Europe | Sweden | 4366.6 | 13.65 | 58.38 | 900 | 0.0246 |
| VK399_noUDG.SG | 1197 | Europe | Sweden | 4366.6 | 13.65 | 58.38 | 900 | 0.0190 |
| VK40_noUDG.SG | 1197 | Europe | Sweden | 4366.6 | 13.65 | 58.38 | 900 | 0.0359 |
| VK400_noUDG.SG | 1197 | Europe | Sweden | 4366.6 | 13.65 | 58.38 | 900 | 0.0248 |
| VK401_noUDG.SG | 1197 | Europe | Sweden | 4366.6 | 13.65 | 58.38 | 900 | 0.0204 |
| VK402_noUDG.SG | 1197 | Europe | Sweden | 4366.6 | 13.65 | 58.38 | 900 | 0.0206 |
| VK403_noUDG.SG | 1197 | Europe | Sweden | 4366.6 | 13.65 | 58.38 | 900 | 0.0268 |
| VK404_noUDG.SG | 1197 | Europe | Sweden | 4366.6 | 13.65 | 58.38 | 900 | 0.0244 |
| VK405_noUDG.SG | 1197 | Europe | Sweden | 4366.6 | 13.65 | 58.38 | 900 | 0.0231 |
| VK406_noUDG.SG | 1197 | Europe | Sweden | 4366.6 | 13.65 | 58.38 | 900 | 0.0267 |
| VK42_noUDG.SG | 1197 | Europe | Sweden | 4366.6 | 13.65 | 58.38 | 900 | 0.0225 |
| VK517_noUDG.SG | 1237 | Europe | Sweden | 4442.5 | 17.65 | 59.86 | 900 | 0.0242 |
| VK527_noUDG.SG | 1237 | Europe | Sweden | 4442.5 | 17.65 | 59.86 | 900 | 0.0216 |
| VK539_noUDG.SG | 1244 | Europe | Ukraine | 3165.9 | 31.18 | 51.37 | 900 | 0.0278 |
| VK540_noUDG.SG | 1244 | Europe | Ukraine | 3165.9 | 31.18 | 51.37 | 900 | 0.0226 |

| | | | | | | | | |
|---|---|---|---|---|---|---|---|---|
| VK542_noUDG.SG | 1246 | Europe | Ukraine | 3165.9 | 31.31 | 51.49 | 900 | 0.0220 |
| VK330_noUDG.SG | 1202 | Europe | Denmark | 4199.9 | 8.76 | 55.33 | 896 | 0.0231 |
| VK353_noUDG.SG | 1203 | Europe | Sweden | 4093.6 | 16.43 | 56.42 | 893 | 0.0209 |
| R9918.SG | 1099 | Europe | Montenegro | 2535.7 | 19.27 | 42.47 | 892 | 0.0198 |
| VK357_noUDG.SG | 1203 | Europe | Sweden | 4093.6 | 16.43 | 56.42 | 887 | 0.0319 |
| DA230_noUDG.SG | 81 | Asia | Kazakhstan | 4411.1 | 76.98 | 43.20 | 883 | 0.0232 |
| SZOD-376.SG | 1149 | Europe | Hungary | 2868.4 | 20.32 | 46.72 | 875 | 0.0156 |
| PLE-384.SG | 1042 | Europe | Hungary | 2913.1 | 21.12 | 47.37 | 870 | 0.0243 |
| PLE-418.SG | 1042 | Europe | Hungary | 2913.1 | 21.12 | 47.37 | 870 | 0.0204 |
| R3482.SG | 1099 | Europe | Montenegro | 2535.7 | 19.27 | 42.47 | 864 | 0.0262 |
| TAQ022 | 1157 | Europe | Italy | 3097.9 | 11.77 | 42.25 | 859 | 0.0144 |
| Anapa-9.SG | 464 | Europe | Russia | 2444.0 | 37.50 | 44.97 | 857 | 0.0260 |
| I3044 | 836 | Europe | United Kingdom | 4453.8 | -0.54 | 53.23 | 856 | 0.0248 |
| I12514 | 628 | Europe | Spain | 4384.3 | -0.20 | 39.93 | 854 | 0.0194 |
| I6890 | 258 | Asia | Pakistan | 3962.5 | 72.36 | 34.76 | 853 | 0.0152 |
| DA118_noUDG.SG | 71 | Asia | Kyrgyzstan | 4404.2 | 76.97 | 42.07 | 851 | 0.0219 |
| SI-53_noUDG.SG | 437 | Both | Lebanon | 727.3 | 35.40 | 33.56 | 851 | 0.0208 |
| I14648 | 667 | Europe | Turkey | 1180.9 | 37.54 | 36.87 | 851 | 0.0176 |
| DA164_noUDG.SG | 77 | Asia | Russia | 2253.8 | 44.18 | 43.33 | 850 | 0.0240 |
| DA179_noUDG.SG | 79 | Asia | Kazakhstan | 4326.4 | 73.62 | 49.04 | 850 | 0.0262 |
| ZAA005 | 419 | Asia | Mongolia | 6542.4 | 104.05 | 47.85 | 850 | 0.0260 |
| I20140 | 754 | Europe | Turkey | 1573.9 | 28.03 | 37.30 | 850 | 0.0112 |
| I20141 | 754 | Europe | Turkey | 1573.9 | 28.03 | 37.30 | 850 | 0.0172 |
| R9920.SG | 1099 | Europe | Montenegro | 2535.7 | 19.27 | 42.47 | 850 | 0.0139 |
| R58.SG | 1112 | Europe | Italy | 3002.1 | 13.09 | 41.69 | 850 | 0.0237 |
| SWG004.A | 1148 | Europe | Germany | 4122.1 | 9.56 | 54.52 | 850 | 0.0220 |
| SWG005.A | 1148 | Europe | Germany | 4122.1 | 9.56 | 54.52 | 850 | 0.0218 |
| SWG006.A_noUDG | 1148 | Europe | Germany | 4122.1 | 9.56 | 54.52 | 850 | 0.0216 |
| SWG007.A_noUDG | 1148 | Europe | Germany | 4122.1 | 9.56 | 54.52 | 850 | 0.0148 |
| SWG010.A_noUDG | 1148 | Europe | Germany | 4122.1 | 9.56 | 54.52 | 850 | 0.0246 |
| SWG011.A | 1148 | Europe | Germany | 4122.1 | 9.56 | 54.52 | 850 | 0.0154 |

| | | | | | | | | |
|---|---|---|---|---|---|---|---|---|
| SWG012.A | 1148 | Europe | Germany | 4122.1 | 9.56 | 54.52 | 850 | 0.0196 |
| SWG015.A | 1148 | Europe | Germany | 4122.1 | 9.56 | 54.52 | 850 | 0.0209 |
| VK160_noUDG.SG | 1187 | Europe | Russia | 3984.8 | 36.16 | 58.78 | 850 | 0.0184 |
| ULA001 | 405 | Asia | Mongolia | 6542.4 | 104.52 | 47.93 | 846 | 0.0215 |
| DA23_noUDG.SG | 85 | Asia | Kazakhstan | 4006.3 | 62.65 | 52.69 | 840 | 0.0234 |
| R64.SG | 1112 | Europe | Italy | 3002.1 | 13.09 | 41.69 | 840 | 0.0182 |
| R65.SG | 1112 | Europe | Italy | 3002.1 | 13.09 | 41.69 | 840 | 0.0225 |
| I10548 | 597 | Europe | Bulgaria | 2257.6 | 25.68 | 43.12 | 825 | 0.0164 |
| I2959 | 208 | Asia | Pakistan | 3962.5 | 72.31 | 34.74 | 822 | 0.0171 |
| Sunghir6_noUDG.SG | 1147 | Europe | Russia | 3576.6 | 40.50 | 56.18 | 813 | 0.0237 |
| SWG001.A | 1148 | Europe | Germany | 4122.1 | 9.56 | 54.52 | 800 | 0.0202 |
| SWG003.A | 1148 | Europe | Germany | 4122.1 | 9.56 | 54.52 | 800 | 0.0272 |
| SWG008.A_noUDG | 1148 | Europe | Germany | 4122.1 | 9.56 | 54.52 | 800 | 0.0215 |
| VK388_noUDG.SG | 1219 | Europe | Norway | 5190.9 | 15.66 | 68.34 | 800 | 0.0284 |
| VK536_noUDG.SG | 1243 | Europe | Italy | 2820.7 | 15.26 | 41.34 | 800 | 0.0205 |
| VK537_noUDG.SG | 1243 | Europe | Italy | 2820.7 | 15.26 | 41.34 | 800 | 0.0229 |
| VK538_noUDG.SG | 1243 | Europe | Italy | 2820.7 | 15.26 | 41.34 | 800 | 0.0259 |
| I17980 | 597 | Europe | Bulgaria | 2257.6 | 25.68 | 43.12 | 797 | 0.0189 |
| SWG002.A | 1148 | Europe | Germany | 4122.1 | 9.56 | 54.52 | 775 | 0.0172 |
| SWG013.A | 1148 | Europe | Germany | 4122.1 | 9.56 | 54.52 | 775 | 0.0239 |
| VK534_noUDG.SG | 1242 | Europe | Italy | 2820.7 | 15.58 | 41.42 | 766 | 0.0201 |
| DA116_noUDG.SG | 70 | Asia | Kyrgyzstan | 4404.2 | 76.96 | 42.07 | 761 | 0.0210 |
| Yana_Young_noUDG.SG | 417 | Asia | Russia | 8066.8 | 135.42 | 70.72 | 761 | 0.0245 |
| DA99_noUDG.SG | 70 | Asia | Kyrgyzstan | 4404.2 | 76.96 | 42.07 | 757 | 0.0239 |
| I1630 | 199 | Asia | Armenia | 1912.7 | 46.20 | 38.87 | 750 | 0.0199 |
| I1660 | 199 | Asia | Armenia | 1912.7 | 46.20 | 38.87 | 750 | 0.0241 |
| I12515 | 628 | Europe | Spain | 4384.3 | -0.20 | 39.93 | 750 | 0.0176 |
| I12644 | 629 | Europe | Spain | 4433.7 | -0.38 | 39.47 | 750 | 0.0251 |
| I12647 | 629 | Europe | Spain | 4433.7 | -0.38 | 39.47 | 750 | 0.0213 |
| I14645 | 667 | Europe | Turkey | 1180.9 | 37.54 | 36.87 | 750 | 0.0187 |
| I14651 | 667 | Europe | Turkey | 1180.9 | 37.54 | 36.87 | 750 | 0.0210 |

| | | | | | | | | |
|---|---|---|---|---|---|---|---|---|
| VK118_noUDG.SG | 1182 | Europe | Norway | 4956.4 | 10.39 | 63.43 | 750 | 0.0204 |
| R62.SG | 1112 | Europe | Italy | 3002.1 | 13.09 | 41.69 | 735 | 0.0237 |
| C1654 | 46 | Asia | China | 4824.5 | 81.84 | 43.22 | 733 | 0.0209 |
| I7716 | 258 | Asia | Pakistan | 3962.5 | 72.36 | 34.76 | 728 | 0.0142 |
| R3361.SG | 434 | Both | Syria | 1158.9 | 40.56 | 34.97 | 728 | 0.0241 |
| SHU002 | 383 | Asia | Mongolia | 6401.6 | 102.77 | 46.91 | 726 | 0.0174 |
| ULN011 | 41 | Asia | Mongolia | 7120.2 | 111.77 | 46.28 | 719 | 0.0193 |
| SI-41_noUDG.SG | 437 | Both | Lebanon | 727.3 | 35.40 | 33.56 | 710 | 0.0147 |
| ARG001 | 8 | Asia | Mongolia | 6618.3 | 105.25 | 47.87 | 700 | 0.0206 |
| ARG003 | 8 | Asia | Mongolia | 6618.3 | 105.25 | 47.87 | 700 | 0.0219 |
| BAY001 | 19 | Asia | Mongolia | 6694.1 | 106.41 | 47.72 | 700 | 0.0230 |
| BAZ001 | 20 | Asia | Mongolia | 6395.3 | 101.04 | 43.20 | 700 | 0.0254 |
| ERD001 | 20 | Asia | Mongolia | 6395.3 | 101.04 | 43.20 | 700 | 0.0209 |
| TAH002 | 22 | Asia | Mongolia | 5630.5 | 91.84 | 47.06 | 700 | 0.0175 |
| BRG002 | 32 | Asia | Mongolia | 6527.0 | 104.88 | 49.22 | 700 | 0.0204 |
| BRG005 | 32 | Asia | Mongolia | 6527.0 | 104.88 | 49.22 | 700 | 0.0243 |
| CHD001 | 63 | Asia | Mongolia | 7007.6 | 110.53 | 47.27 | 700 | 0.0184 |
| DAS001 | 105 | Asia | Mongolia | 7305.7 | 113.86 | 45.35 | 700 | 0.0262 |
| GAN002 | 118 | Asia | Mongolia | 6478.0 | 102.13 | 43.15 | 700 | 0.0272 |
| GTO001 | 120 | Asia | Mongolia | 7305.7 | 114.02 | 45.22 | 700 | 0.0248 |
| KGK001 | 293 | Asia | Mongolia | 6395.9 | 102.64 | 47.55 | 700 | 0.0262 |
| KHL001 | 294 | Asia | Mongolia | 5708.1 | 92.79 | 46.69 | 700 | 0.0305 |
| KHN001 | 295 | Asia | Mongolia | 7172.7 | 112.06 | 46.95 | 700 | 0.0210 |
| KNN001 | 295 | Asia | Mongolia | 7172.7 | 112.06 | 46.95 | 700 | 0.0207 |
| ZAY001 | 295 | Asia | Mongolia | 7172.7 | 112.06 | 46.95 | 700 | 0.0210 |
| KNU001 | 302 | Asia | Mongolia | 6239.2 | 100.54 | 48.09 | 700 | 0.0244 |
| KRN001 | 306 | Asia | Mongolia | 6669.5 | 106.67 | 49.74 | 700 | 0.0159 |
| KRN002 | 306 | Asia | Mongolia | 6669.5 | 106.67 | 49.74 | 700 | 0.0246 |
| MRI001 | 325 | Asia | Mongolia | 6305.4 | 101.46 | 49.44 | 700 | 0.0230 |
| RAH001 | 354 | Asia | Mongolia | 6941.4 | 108.97 | 47.22 | 700 | 0.0216 |
| SHA001 | 377 | Asia | Mongolia | 6538.8 | 103.61 | 44.08 | 700 | 0.0177 |

| | | | | | | | | |
|---|---|---|---|---|---|---|---|---|
| SHG001 | 380 | Asia | Mongolia | 7263.1 | 113.49 | 46.37 | 700 | 0.0219 |
| SHG002 | 380 | Asia | Mongolia | 7263.1 | 113.49 | 46.37 | 700 | 0.0252 |
| SHG003 | 380 | Asia | Mongolia | 7263.1 | 113.49 | 46.37 | 700 | 0.0277 |
| SHR001 | 381 | Asia | Mongolia | 7172.7 | 112.05 | 46.97 | 700 | 0.0251 |
| TSA001 | 400 | Asia | Mongolia | 7265.6 | 114.88 | 49.14 | 700 | 0.0193 |
| TSA002 | 400 | Asia | Mongolia | 7265.6 | 114.88 | 49.14 | 700 | 0.0212 |
| TSA004 | 400 | Asia | Mongolia | 7265.6 | 114.88 | 49.14 | 700 | 0.0250 |
| TSA005 | 400 | Asia | Mongolia | 7265.6 | 114.88 | 49.14 | 700 | 0.0256 |
| TSA006 | 400 | Asia | Mongolia | 7265.6 | 114.88 | 49.14 | 700 | 0.0210 |
| UGO002 | 403 | Asia | Mongolia | 7149.0 | 112.94 | 47.90 | 700 | 0.0185 |
| ZAM002 | 420 | Asia | Mongolia | 6580.2 | 104.07 | 45.53 | 700 | 0.0198 |
| VK535_noUDG.SG | 1242 | Europe | Italy | 2820.7 | 15.58 | 41.42 | 700 | 0.0228 |
| VK541_noUDG.SG | 1245 | Europe | Ukraine | 3070.3 | 25.32 | 50.74 | 700 | 0.0225 |
| SI-45_noUDG.SG | 437 | Both | Lebanon | 727.3 | 35.40 | 33.56 | 698 | 0.0182 |
| I20187 | 754 | Europe | Turkey | 1573.9 | 28.03 | 37.30 | 660 | 0.0178 |
| I1659 | 199 | Asia | Armenia | 1912.7 | 46.20 | 38.87 | 655 | 0.0247 |
| I19539 | 754 | Europe | Turkey | 1573.9 | 28.03 | 37.30 | 650 | 0.0131 |
| I20142 | 754 | Europe | Turkey | 1573.9 | 28.03 | 37.30 | 650 | 0.0162 |
| I20143 | 754 | Europe | Turkey | 1573.9 | 28.03 | 37.30 | 650 | 0.0105 |
| I20145 | 754 | Europe | Turkey | 1573.9 | 28.03 | 37.30 | 650 | 0.0240 |
| I20146 | 754 | Europe | Turkey | 1573.9 | 28.03 | 37.30 | 650 | 0.0121 |
| I20147 | 754 | Europe | Turkey | 1573.9 | 28.03 | 37.30 | 650 | 0.0184 |
| I20574 | 754 | Europe | Turkey | 1573.9 | 28.03 | 37.30 | 650 | 0.0186 |
| I7715 | 258 | Asia | Pakistan | 3962.5 | 72.36 | 34.76 | 619 | 0.0195 |
| R2065.SG | 1092 | Europe | France | 3819.7 | 6.17 | 49.12 | 613 | 0.0168 |
| BSK002 | 39 | Asia | Kyrgyzstan | 4245.8 | 74.69 | 42.81 | 612 | 0.0258 |
| I6889 | 258 | Asia | Pakistan | 3962.5 | 72.36 | 34.76 | 605 | 0.0188 |
| UUS002 | 408 | Asia | Mongolia | 6231.6 | 100.41 | 49.40 | 604 | 0.0159 |
| R2066.SG | 1092 | Europe | France | 3819.7 | 6.17 | 49.12 | 604 | 0.0277 |
| R1290.SG | 1083 | Europe | Italy | 3097.9 | 12.47 | 41.90 | 602 | 0.0254 |
| BRL001 | 33 | Asia | Mongolia | 6600.9 | 105.91 | 49.31 | 601 | 0.0214 |

| | | | | | | | | |
|---|---|---|---|---|---|---|---|---|
| UUS001 | 408 | Asia | Mongolia | 6231.6 | 100.41 | 49.40 | 600 | 0.0232 |
